# Enhancer locus in ch14q23.1 modulates brain asymmetric temporal regions involved in language processing

**DOI:** 10.1101/539189

**Authors:** Yann Le Guen, François Leroy, Cathy Philippe, IMAGEN consortium, Jean-François Mangin, Ghislaine Dehaene-Lambertz, Vincent Frouin

## Abstract

Identifying the genes that contribute to the variability in brain regions involved in language processing may shed light on the evolution of brain structures essential to the emergence of language in Homo sapiens. The superior temporal asymmetrical pit (STAP), which is not observed in chimpanzees, represents an ideal phenotype to investigate the genetic variations that support human communication. The left STAP depth was significantly associated with a predicted enhancer annotation located in the 14q23.1 locus, between *DACT1* and *KIAA0586*, in the UK Biobank British discovery sample (N=16,515). This association was replicated in the IMAGEN cohort (N=1,726) and the UK Biobank non-British validation sample (N=2,161). This genomic region was also associated to a lesser extent with the right STAP depth and the formation of sulcal interruptions, *plis de passage*, in the bilateral STAP but not with other structural brain MRI phenotypes, highlighting its notable association with the superior temporal regions. Diffusion MRI emphasized an association with the fractional anisotropy of the left auditory fibers of the corpus callosum and with networks involved in linguistic processing in resting-state functional MRI. Overall, this evidence demonstrates a specific relationship between this locus and the establishment of the superior temporal regions that support human communication.

## Introduction

Which genes participate in the biological foundations of human language processing? This question remains unanswered. Initial approaches sought to identify rare monogenic mutations in families with language and speech disorders, which revealed the roles of *FOXP2* (Fisher et al. 1998) and *KIAA0319* (Francks et al. 2004), for example. The discovery of these genes led to productive research in humans and animals in the following aspects: First, which brain areas do these genes modulate either structurally or functionally in humans; and second, how do animal variants and even human variants in animals affect neural networks and animal abilities? A complementary approach, which is now possible due to genotyping and neuroimaging in large cohorts, is to investigate the relationship between certain brain structures and allelic variations in the normal human population (Cai et al. 2014). Although all participants have language ability, this type of research can pinpoint particular genetic and neural architectures, opening new windows to understanding language. For example, Dediu and Ladd (Dediu and Ladd 2007) proposed that typological features of languages may depend on certain allele frequencies in human groups, and they explored the relationship between tonal languages and the distribution of allelic variants of *ASPM* and *Microcephalin*. Functional and structural asymmetries may be one of the human brain phenotypes whose genetic study may shed light on this issue. Indeed, human communication is based on the retrieval of two crucial types of information in speech, who speaks and what is said. In most adults, the message and the messenger are processed in parallel in the superior temporal region, with the linguistic message (Mazoyer et al. 1993) and voice identification (Belin et al. 2000) processed in the left and right hemispheres, respectively. This channeling is present early in human infants (Dehaene-Lambertz et al. 2002; Bristow et al. 2009). At six months of gestation, activations to syllables are differently lateralized when preterm infants discriminate a phonetic change (/ba/ vs /ga/) or a voice change (male vs female) (Mahmoudzadeh et al. 2013). Thus, the left and right peri-sylvian regions are distinctly functionally organized from the onset of thalamocortical connectivity. Therefore, this organization occurs before sensory input can shape these regions, suggesting genetically determined protomaps.

These functional asymmetries should be discussed in relation to the notable asymmetries visible at the macroscopic level and reported near the Sylvian fissure. The Sylvian fissure is more dorsal and oblique in the right hemisphere, the *planum temporale* is larger in the left hemisphere (Geschwind and Levitsky 1968) and the right superior temporal sulcus (STS) is deeper than the left (Leroy et al. 2015). Some asymmetries, such as the shape and location of the Sylvian fissure and the left-right size difference in the *planum*, are observed in apes (Hopkins et al. 2000), but the STS depth asymmetry is the first macroscopic brain asymmetry that has been observed only in humans. A 45-mm-long segment located under Heschl’s gyrus, superior temporal asymmetrical pit (STAP), is deeper in the right hemisphere than in the left in 96% of human infants, children, and adults but not in chimpanzees (Leroy et al. 2015). During gestation, the right STS appears earlier than the left, similar to many other sulci (Fontes 1944; Dubois et al. 2008; Kasprian et al. 2011; Habas et al. 2012); however other left sulci catch up, while the left STS remains shallower than the right STS and is more often interrupted by one or more *plis de passage* (PPs) (Leroy et al. 2015; Le Guen, Leroy, et al. 2018). The observation of these structural and functional differences before term age suggests that genetically driven mechanisms guide the development of peri-sylvian regions to endorse the complexity of verbal and nonverbal communication in the human species. Because of the sharp distinction between chimpanzees and humans, Leroy et al. (2015) suggested that the STAP may represent an important macroscopic marker of the recent cytoarchitectonic changes (Cantalupo and Hopkins 2001) in the primate lineage supporting the communication function in humans. We have previously examined the heritability (i.e., the genetic proportion in phenotypic variance across individuals) of STS characteristics. Because sulcal pits correspond to the deeper parts of sulci where the initial or primary folds occur, they show less intersubject variability (Meng et al. 2014; Le Guen, Auzias, et al. 2018). In the left STAP area, the two sulcal pit regions (STS b and c in Le Guen, Auzias, et al. (2018)) are highly heritable, in contrast to the right STS, yielding the largest asymmetric estimate of heritability for all brain sulci. In addition, the PPs that interrupt some sulci are more frequent and heritable in the left STS (h^2^ = 0.53) than in the right (h^2^ = 0.27) (Le Guen, Leroy, et al. 2018). In most brain regions, this interhemispheric difference in heritability estimates was not detected, therefore suggesting a distinctive genetic modulation in the superior temporal lobes.

Therefore, to clarify the genetic influence on the development of the human communication system, we studied the genetic variants that are significantly associated with STAP phenotypes (i.e., the right and left sulcal depths, their asymmetry and the formation of PPs). We took advantage of the most recent release of the UK Biobank, which includes approximately 22,000 individuals genotyped and scanned with the same MRI scanner and imaging protocol (Miller et al. 2016). We used 16,515 genetically confirmed individuals of British ancestry as our discovery sample and the remaining 2,161 non-British ancestry subjects as a first replication sample. In addition, we used the IMAGEN cohort of 1,726 adolescents (14-15 years) as the second replication sample. We found that an intergenic region upstream of *DACT1* located in 14q23.1 modulates STS phenotypes. To further understand the functional role of this region, we first used databases of information on gene tissue expression and chromatin states in various brain tissues at different stages of development. Second, we investigated the effect of this region on brain phenotypes by taking advantage of the multimodal sequences acquired in the same UK Biobank subjects (T1w, diffusion tensor imaging (DTI) and resting state). We observed that this gene has a localized effect on the superior temporal regions, particularly on the left STS, which suggests a role in the establishment of the linguistic network.

## Methods

### Subjects

In this study, we used the UK Biobank data released in October 2018, including 22,392 subjects with T1-weighted MRI who were aged 44 to 81 years at the time of the imaging visit. The genetic data have already been subject to rigorous quality control (Bycroft et al. 2018). We limited our analysis to individuals identified by the UK Biobank as belonging to the main subset of white British ancestry. Subjects with high genotyped missingness, high heterozygosity or sex mismatches were excluded from our analysis (Bycroft et al. 2018). Briefly, the UK Biobank contains genotypes of 488,377 participants who were genotyped on the following arrays: UK BiLEVE Axiom Array (807,411 markers; 49,950 participants) and UK Biobank Axiom Array (825,927 markers; 438,427 participants). The two arrays are very similar, sharing approximately 95% of the marker content. Genotypes were called from the array intensity data in 106 batches of approximately 4700 samples each using a custom genotype-calling pipeline.

Among individuals with the STAP phenotype, 13,956 individuals of British ancestry were scanned at Cheadle and 2,559 at Nottingham, while, 1,935 individuals of non-British ancestry were scanned at Cheadle and 226 at Nottingham.

### Ethical approval

UK Biobank has approval from the North West Multi-centre Research Ethics Committee (MREC), which covers the UK. The present study was approved by the UK Biobank ethics committee under application number 25251.

### Imputation

Phasing and imputation were carried out by the UK Biobank and are described in detail elsewhere (Bycroft et al. 2018). Briefly, phasing on the autosomes was performed using SHAPEIT3 (O’Connell et al. 2016) with the 1000 Genomes Phase 3 dataset as a reference panel. For imputation, both the Haplotype Reference Consortium (HRC) reference panel (McCarthy et al. 2016) and a merged UK10K/1000 Genomes Phase 3 panel were used. This imputation resulted in a dataset with 92,693,895 autosomal SNPs, short indels and large structural variants. This customized imputation represents the most reliable high-resolution imputation to date and enables fine-mapping to identify causal genetic variants (Schaid et al. 2018). Using PLINK v1.9 (Chang et al. 2015), we filtered variants from the UK Biobank imputed data. We removed variants with more than 5% missing genotype data, variants with Hardy-Weinberg exact test < 10^−6^ and variants with minor allele frequency < 0.01. 8,011,919 imputed variants remained after filtering. In addition, we excluded all individuals who fulfilled at least one of the following criteria: discrepancy between genetically inferred sex (Data field 22001) and self-reported sex (Data field 31), outliers in heterozygosity and missing rates (Data field 22027), and individuals not flagged by UK Biobank as belonging to a white British ancestry subset (Data field 22006) (i.e., identified as British by principal component analysis and self-reported ‘British’).

### MRI data acquisition

The MRI data acquisition parameters and preprocessing are described in detail in the UK Biobank Brain Imaging Documentation (v.1.5, August 2018) and in previous UK Biobank publications describing imaging phenotypes and QC (Miller et al. 2016; Alfaro-Almagro et al. 2018; Elliott et al. 2018).

Briefly:

- T1 images were acquired with 1 mm isotropic resolution (key parameters: 3D MPRAGE, sagittal, R=2, T1/TR = 880/2000 ms, duration = 4 min:54 s);
- T2 FLAIR images were acquired with a 1.05*1.0*1.0 mm resolution (key parameters: FLAIR, 3D SPACE, sagittal, R = 2, PF 7/8, fat sat, T1/TR = 1800/5000 ms, elliptical, duration 5 min:52 s);
- diffusion MRI (dMRI) images were acquired with 2.0 mm isotropic resolution (key parameters: MB = 3, R = 1, TE/TR = 92/3600 ms, PF 6/8, fat sat, b = 0 s/mm^2^ (5x+ 3*phase-encoding reversed), b = 1000 s/mm^2^ (50*), b = 2000 s/mm^2^ (50*), duration = 7 min:8s);
- resting-state functional MRI (rfMRI) images were acquired with 2.4 mm isotropic resolution (key parameters: TE/TR = 39/735 ms, MB = 8, R = 1, flip angle = 52°, fat sat, duration = 6 min:10 s).

### STAP depth extraction

The STAP section was extracted using the automated method described in Le Guen, Leroy, et al. (2018). Briefly, we first extracted the sulcal pits (primary cortical folds) using a watershed algorithm (Auzias et al. 2015). Then, we summed all the individual pit densities on a symmetric template using surface-based interhemispheric registration (Greve et al. 2013). We used a watershed algorithm on this group pit density, which yielded a group parcellation of the sulcal pits. We projected these group areas onto the individual meshes and identified areas denoted STS b, c, d (Le Guen, Auzias, et al. 2018). Then, we extracted the geodesic path of the vertices that follow the sulcal fundus on the white mesh between the deepest pits STS b and the STS c/d border, which approximately correspond to the STAP Talaraich coordinates defined by Leroy et al. (2015). From this set of vertices, we obtained individual profiles of the geodesic depth and depth potential function (PF), which considers both sulcal depth and convexity. We averaged these values along the STAP for each subject to obtain our depth phenotypes. We also applied the PP detection method described in Le Guen, Leroy, et al. (2018) and benchmarked on manually labeled data from Leroy et al. (2015) to identify subjects with at least one sulcal interruption in their depth profile. This process led to the binary phenotype “presence or absence of PP”. Among our British discovery sample that completed the pipeline (16,515 subjects), 9,501 subjects had a left PP (57.53%), and 2,587 had a right PP (15.66%). We defined the asymmetry index as 2(L-R)/(L+R).

### Phenome-wide analysis

We processed the T1w and T2w images with the Freesurfer v6 pipeline (Fischl 2012), and extracted individual measures from the following output files: ?h.aparc.stats (Desikan parcellation (Desikan et al. 2006)), ?h.aparc.a2009s.stats (Destrieux parcellation (Destrieux et al. 2010)), and aseg.stats (subcortical volumes). We processed the T1w images with Brainvisa Morphologist pipeline (Perrot et al. 2011) and obtained individual sulcus measures per subject. We obtained diffusion MRI and resting-state MRI phenotypes from the UK Biobank (Miller et al. 2016; Elliott et al. 2018). Briefly, diffusion MRI measures were extracted using TBSS (Smith et al. 2006) from the ICBM 48 white matter tracts atlas (Mori et al. 2008). For resting-state fMRI (rfMRI), independent component analysis (ICA) was performed by the UK Biobank on the first 4,181 participants (Miller et al. 2016). A total of 21 and 55 good components were identified for 25 and 100 component analyses, respectively (Elliott et al. 2018). These components were subsequently used for rfMRI acquired in the following subjects. The rfMRI phenotypes include the amplitude of the ICA weighted rfMRI BOLD signal, referred to as ICA amplitude, the Pearson correlation and partial correlations between pairs of ICA time series, which are also referred to as functional connectivity.

### Quantitative genetics

In our discovery sample, we used genome complex trait analysis (GCTA), (Yang et al. 2011) which provides an estimate of heritability in population studies with unrelated genotyped participants. To compute the kinship matrix of the population, we selected specific SNPs with PLINK v1.9 (Purcell et al. 2007) using the following thresholds: missing genotype = 5% (70,737 variants excluded), Hardy-Weinberg equilibrium test (hwe) = 10^−6^ (18,462), and minor allele frequency (maf) = 1.0% (101,839). We kept the SNPs in moderate linkage disequilibrium with a variation inflation factor of 10 within a window of 50 SNPs (91,496 variants excluded). Then, we computed the genetic relationship matrix with GCTA 1.25.3 using the 501,722 SNPs left. The amount of phenotypic variance captured by this matrix is estimated using a mixed-effects linear model. As covariates, we included gender, genotyping array type, age at the MRI session and the 10 genetic principal components (PCs) provided by the UK Biobank to account for population stratification.

### Heritability replication in the HCP cohort

The heritability estimates for the Human Connectome Project followed the steps described in our previous study (Le Guen, Leroy, et al. 2018). Briefly, these estimates are computed using the Sequential Oligogenic Linkage Analysis Routines (SOLAR) (Almasy and Blangero 1998) on an extended pedigree of 820 Caucasian individuals, which includes twins and siblings. To replicate heritability estimates of the STAP phenotypes, we included 820 subjects (383/437 M/F) labeled as Caucasian with 69 individuals from the Hispanic ethnicity. The pedigree is composed of 191 twin pairs (127 monozygotic (MZ) with 123 siblings and 64 dizygotic (DZ) with 64 siblings and 1 half sibling), 190 siblings, 1 half sibling and 59 unpaired individuals, aged between 22 and 36 years old (μ ± σ = 29.0 ± 3.6 years). The pedigree was provided by the HCP, and most sibling pairs were genetically confirmed. Note that in (Le Guen, Leroy, et al. 2018), we only reported in the main text the heritability of the PPs. However, these were extracted in each individual based on their STAP geodesic depth and depth PF; thus, to obtain the other STAP features, we averaged these depth profiles and computed their asymmetry index in the same set of subjects and used the same methodology to estimate the heritability.

### Genome-wide associations in the UK Biobank

The analyses of genotype-phenotype associations were performed with PLINKv1.9 (Purcell et al. 2007) using the imputed genotype data. The same thresholds as above were applied (missing genotype: 5%, hwe: 10^−6^, and maf: 0.01). We included the same covariates as those in the quantitative genetics section.

### GWAs replication in the IMAGEN cohort

For the replication of GWAS, we included 1,726 subjects with T1 MR Images (ADNI-MPRAGE whose parameters are available in (Schumann et al. 2010)) and genotyping data. The SNPs common to the Illumina 610-Quad and 660-Quad arrays were considered and we used the imputed data available from the IMAGEN project (https://imagen2.cea.fr). Briefly, imputation was performed using MaCH using the 1000 genome Phase 3 dataset as a reference (v3.20101123).

### Functional annotation

The functional annotations of loci were provided by FUMA (Watanabe et al. 2017), which notably annotates SNP location, expression quantitative trait loci (eQTL) querying the Braineac (Ramasamy et al. 2014), GTEx (GTEx Consortium 2017), Common Mind (CMC) (Fromer et al. 2016), and eQTLGen (Võsa et al. 2018) databases, and splicing QTL (sQTL) in GTEx. GTEx, Braineac and eQTLGen were further manually queried. Additionally, we queried the Brain-eMeta eQTL summary data (Qi et al. 2018), which is a set of eQTL data from a meta-analysis of GTEx brain (GTEx Consortium 2017), CMC (Fromer et al. 2016), and ROSMAP (Ng et al. 2017). The estimated effective sample size is n = 1,194. Only SNPs within a 1Mb distance from each probe are available. This dataset is available on the SMR website (cnsgenomics.com/software/smr). The chromatin state was from the Roadmap Epigenomics Consortium (Roadmap Epigenomics Consortium et al. 2015) and inferred by ChromHMM (Ernst and Kellis 2012) in various brain tissues.

## Results

### Heritability and genome-wide association

The first two columns in **Table 1** present the heritability estimates of STAP phenotypes in the following populations: the extended pedigree of the Human Connectome Project (HCP) using SOLAR (Almasy and Blangero 1998) and unrelated individuals in the UK Biobank using GCTA (Yang et al. 2011). Although weaker due to the different cohort design, the higher heritability estimates in the left hemisphere of the geodesic depth and presence of PP previously obtained in HCP (Le Guen, Leroy, et al. 2018) were reproduced in UK Biobank individuals (details in **Table S1**). Note that heritability estimates are generally higher in a pedigree than in unrelated individuals, when considering the same phenotype, due to shared environmental effects, non-additive genetic variations and/or epigenetic factors (Yang et al. 2017). The depth asymmetry index was also heritable. We computed the genome-wide association (GWA) with these phenotypes in the UK Biobank discovery sample (see **Fig. S1-S3** for the Manhattan and QQ plots of these association tests). A region upstream (26.5 kb) of the *DACT1* gene in the cytogenetic band 14q23.1 contains SNPs whose associated p-values passed the Bonferroni-corrected genomic threshold (p < 5**·**10^−8^/8 = 6.25**·**10^−9^) (**Fig. 1A**). **Fig. S4** presents the associated region corresponding to 2 MB around the locus. The strongest association in imputed SNPs was with rs160458 (p =2.0**·**10^−12^, N = 16,203, base pair (bp) position = 59074878 (GRCh37)) and in genotyped SNPs with rs160459 (p = 4.2**·**10^−12^, N=16,515, base pair position = 59074136), which are both annotated in Ensembl (Zerbino et al. 2018) as “human-specific bases” (**Fig. S5**). Suggestive associations (10^−6^ < p < 10^−8^) were observed for other left/right STAP phenotypes, while the asymmetry index of the geodesic depth and DPF were significant at p < 0.05 (uncorrected). In the following section, we focus on the genotyped SNP rs160459 because its chromatin state in the brain appears more consistent with a regulatory role (**Fig. 1B**), and both SNPs are in strong linkage disequilibrium (r^2^ = 0.936 in British ancestry, distance in build GRCh37 = 742 bp). **Table S2** provides estimate of the proportion of variance in phenotype explained for rs160459. We note that rs160459 dosage only explains 0.3% of the left STAP depth variability and 1.4% of this trait heritability in the UK Biobank. For association with the lead SNP, we controlled *a posteriori* for the MRI scanner center and brain volume (**Tables S3-S4**). These two covariates do not affect the main result presented here.

**Table 1.** P-values of the association of rs160459 with the STAP phenotypes in three populations.

| Phenotype/Cohort | Heritability | | Association - $\beta$ (p-val) and OR (p-val) for PPs | | |
| --- | --- | --- | --- | --- | --- |
|  | HCP | UKB | UKB discovery<br>N=16,515 British | IMAGEN<br>replication<br>N=1,726 | UKB replication<br>N=2,161 Non-British |
| Left STAP geodesic depth | 48%* | 21%* | **0.2683 ($4.2 \cdot 10^{-12}$)** | 0.4614 ($8.0 \cdot 10^{-5}$) | 0.242 (0.0133) |
| Right STAP geodesic depth | 46%* | 15%* | 0.179 ($1.3 \cdot 10^{-7}$) | 0.3204 ($2.2 \cdot 10^{-4}$) | 0.04269 (0.1563) |
| AI of the STAP geodesic depth | 30%* | 9%* | 0.008638 (0.0017) | 0.01226 (0.0406) | 0.01355 (0.1675) |
| Left STAP depth PF | 25%* | 18%* | 0.009865 ($3.8 \cdot 10^{-8}$) | 0.01265 (0.01) | 0.009936 (0.1004) |
| Right STAP depth PF | 38%* | 4% | 0.004759 ($4.1 \cdot 10^{-6}$) | 0.005239 (0.0971) | 0.004668 (0.0249) |
| AI of the STAP depth PF | 22%* | 11%* | 0.004949 (0.0053) | 0.007321 (0.0909) | 0.005525 (0.5414) |
| PP in the left STAP | 53%* | 16%* | 0.8901 ($2.7 \cdot 10^{-7}$) | 0.8025 (0.0022) | 0.9084 (0.2447) |
| PP in the right STAP | 27% | 0% | 0.8533 ($2.6 \cdot 10^{-7}$) | 0.7506 (0.0034) | 0.8724 (0.2067) |

**Fig. 1.**
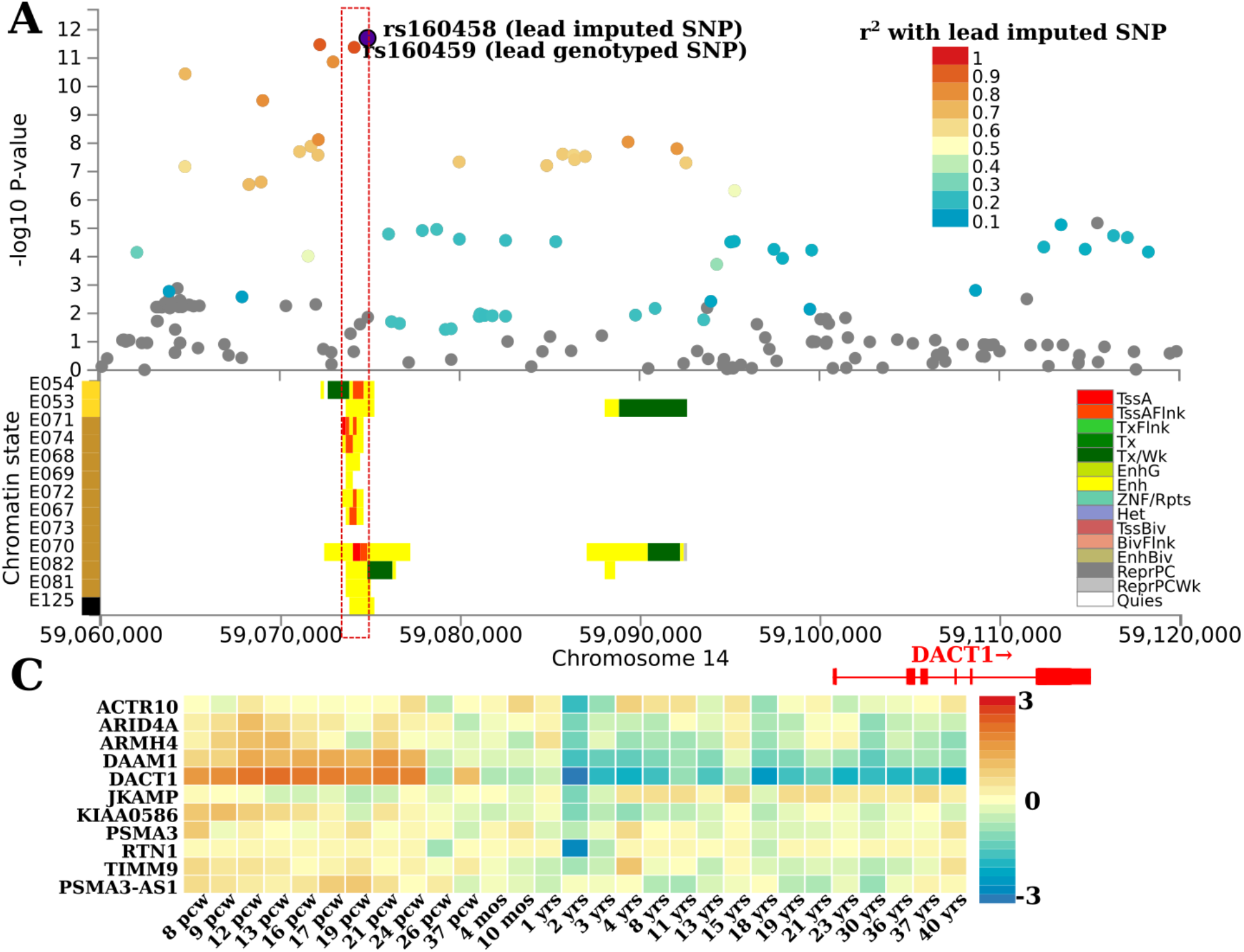
**(A) Manhattan plot of the geodesic depth in the left STAP zoomed in on the significant associations upstream of the** *DACT1* **gene (14q23.1). (B) Chromatin state of the genomic region near lead SNPs rs160458 (imputed) and rs160459 (genotyped).** The *DACT1* position and exons are highlighted in red on the axis showing the base pair position (GRCh37) on chromosome 14. Yellow corresponds to the enhancer (Enh) state, red-orange to the flanking active transcription start site (TssAFlnk), green to weak transcription (Tx/Wk) and white to quiescent (Quies) (see ref. (Ernst and Kellis 2012; Roadmap Epigenomics Consortium et al. 2015) for the complete legend). **(C) Average normalized gene expression (zero mean across samples) of genes +/−1 MB of rs160459 (Fig. S4, Fig. S7) at 29 developmental ages from Brainspan** (Miller et al. 2014). Brain tissue name abbreviations in **(B)**: E054: Ganglion Eminence-derived primary cultured neurospheres, E053: Cortex-derived primary cultured neurospheres, E071: Brain Hippocampus Middle, E074: Brain Substantia Nigra, E068: Brain Anterior Caudate, E069: Brain Cingulate Gyrus, E072: Brain Inferior Temporal Lobe, E067: Brain Angular Gyrus, E073: Brain Dorsolateral Prefrontal Cortex, E070: Brain Germinal Matrix, E082: Fetal Brain Female, E081: Fetal Brain Male, E125: Astrocyte Primary Cells.

### Replication of the association in two other populations

(**Table 1** for rs160459 and **Table S5** for rs160458). The association with the STAP left geodesic depth was replicated in the two samples above the nominal significance level (p < 0.05), with p = 8.0**·**10^−5^ in the IMAGEN cohort and p = 0.041 in the non-British UK Biobank sample. On average, the allelic repartition across the three population samples is 21.6% ‘CC’, 49.9% ‘CA’, and 28.5% ‘AA’. **Fig. S6** illustrates the effect size of the rs160459 allelic configuration on the geodesic depth and depth PF in the three populations revealing shallower STS in the ‘AA’ allelic configuration. **Fig. 2A** shows the influence of rs160459 along the STAP geodesic depth profile. Its allelic configuration exerts a strong control on the posterior part of the left STAP notably at the average location of the posterior PP when it exists.

**Fig. 2.**
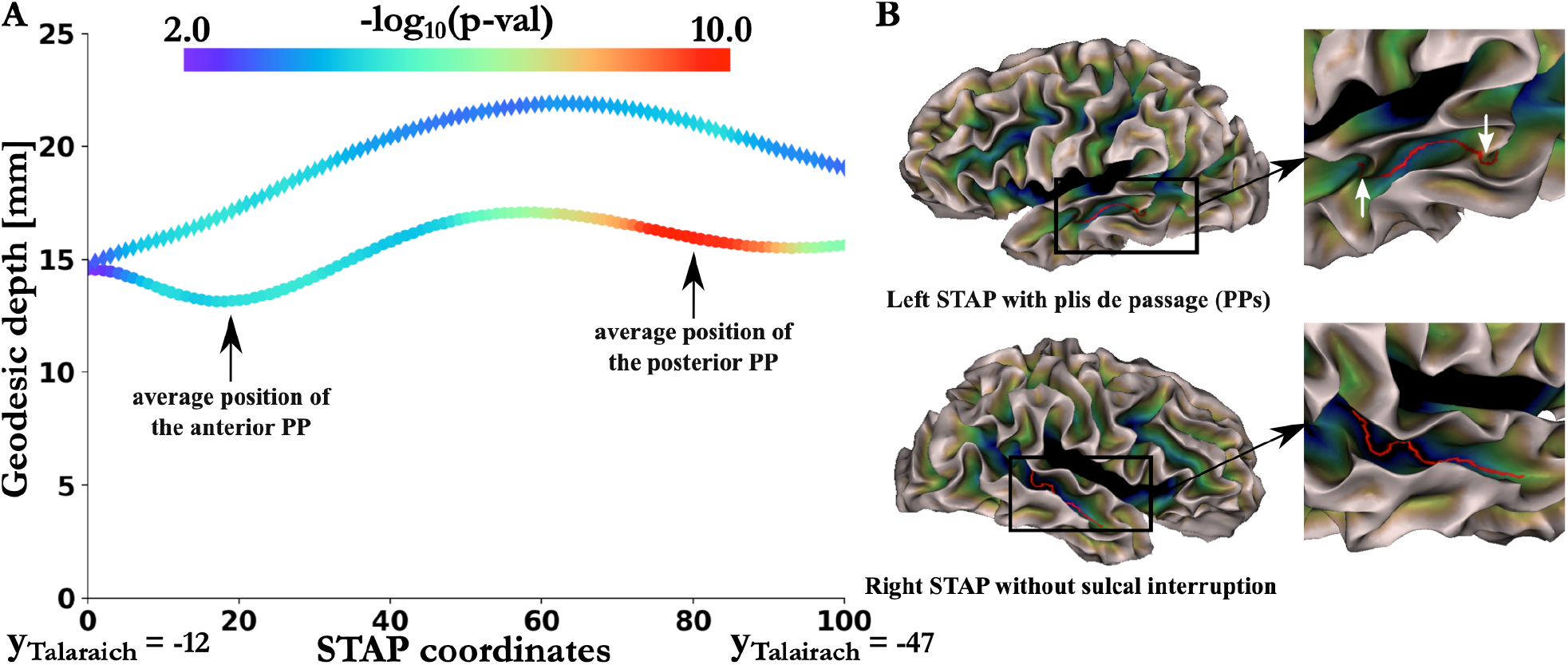
**(A) Mapping of the p-values along the STAP profile averaged over 16,515 British subjects of the association between rs160459 and geodesic depth.** Left values are represented by disks and right values by diamonds (for more details, see **Fig. S16**). **(B) Example of asymmetries in an individual subject with anterior and posterior PPs in the left STAP but no PP in the right STAP.** Note that most subjects have only one PP either anterior or posterior in the left STAP and no sulcal interruption in the right STAP (Le Guen, Leroy, et al. 2018).

### Role of the rs160459 region in the regulation of nearby genes

**Fig. 1B** presents the chromatin state functional annotation of the genomic region upstream of *DACT1* (i.e., the region significantly associated with the left STAP geodesic depth). The subregion around rs160459 has its chromatin state in brain tissues annotated as enhancer (Enh) and/or flanking active transcription start site (TssAFlnk) (Ernst and Kellis 2012; Roadmap Epigenomics Consortium et al. 2015), thus revealing its active chromatin state in all tested brain areas, with the notable exception of the dorso-lateral prefrontal cortex. In contrast, this region has a quiescent chromatin state in most other body tissues except stem cells and epithelial cells, suggesting a tropism for neuro-ectoderm derived cell lines (**Fig. S7**). **Table S6** provides the eQTL query results in Brain-eMeta for the genome-wide significant (GWS) hits. We did not find any significant eQTLs in brain tissues among our GWS SNPs in Brain-eMeta, except rs10782438 which eQTL of *PSMA3-AS1* (p_uncorrected_ = 7**·**10^−5^, p_FDR_ = 0.05). Querying eQTLGen, which includes a large number of subjects with gene expression in blood, rs160459 is an eQTL of *DACT1* (p_uncorrected_ = 9**·**10^−5^, N = 30,757) but not of other protein coding genes, with a decrease in *DACT1* expression with rs160459-C. Additionally, one of the GWS SNPs, rs10782438, is an eQTL FDR significant of *DACT1* (p_uncorrected_ = 5**·**10^−7^, N = 31,086, p_FDR_ = 0.01) (**Table S7**). In British ancestry, according to LDlink, rs160459 and rs10782438 are in strong linkage disequilibrium with r^2^ = 0.9132 and D’ = 0.9772. Additionally, we note in GTEx that rs160459 and several SNPs of the GWS locus block are splicing QTLs (sQTLs) of *KIAA0586* and *PSMA3-AS1* in several non-brain tissues (**Table S8**). Finally, we obtained the CADD (Combined Annotation Dependent Depletion) scores (Rentzsch et al. 2019) of GWS variants and observed that rs160459 has the highest score among these variants (CADD = 14.66, **Table S9**). **Table S10** shows the predicted motif features of rs160459 site. It includes as predicted consequences a modification of the region considered as transcription factors binding sites, which would impact the putative regulation of the following genes: *CUX1, ETV2:PAX5, ELK1:PAX1, ELK1:PAX5, ELK1:PAX9, ETV2:BHLHA15, FLI1:BHLHA15, FOXG1, FOXK1, GCM2:SOX15, HOXB2:SOX15.* **Fig. S8** presents the LD structure around the regulatory including rs160459 in the 1000 Genomes British panel.

### Nearby gene expression across the lifespan

**Fig. S9** presents the expression levels of genes within +/−1 MB of the GWS locus, across brain developmental stages, from Brainspan (Miller et al. 2014). Among these genes, *ACTR10*, *RTN1* and *DACT1* showed the highest expression level, with *RTN1* mainly expressed in brain tissues (GTEx data not shown). The normalized expression of the genes at least moderately expressed in the brain is reported in **Fig. 1C**. We note that *DACT1* and *DAAM1* are mainly expressed during early brain development in the following distinct periods: high expression from the early to late mid-prenatal period (8 to 24 weeks post-conception (pcw), no data before 8 pcw), followed by a decrease from the late prenatal period (26 pcw) until the end of the first year, and minimal expression after early childhood (from 2 years of age). *KIAA0586* is mainly expressed during the early prenatal period (8 to 12 pcw), while *PSMA3-AS1* has a similar trend with an expression level peak during the late-mid prenatal period (16-17 pcw).

### Association of multimodality MRI phenotypes association with rs160459

To characterize the potential role of this enhancer genomic region in shaping the human brain, we performed a phenome-wide association (PheWA) of rs160459 with various brain phenotypes computed on structural, diffusion and functional MRI exams available from the UK Biobank. We extracted (i) the brain sulci features with Morphologist (Perrot et al. 2011); (ii) cortical thickness, surface area and gray matter volumes with Freesurfer on the Desikan (Desikan et al. 2006) and Destrieux (Destrieux et al. 2010) atlases; and (iii) the subcortical volumes with Freesurfer. From the UK Biobank, we obtained (iv) tract-based spatial statistics (TBSS) (Smith et al. 2006) on white matter tracts from the ICBM atlas (Mori et al. 2008) and (v) resting-state functional MRI (rfMRI) phenotypes (see Methods).

**Table 2** summarizes the 20 phenotypes with the strongest associations (smallest p-values) with rs160459 (**Table S11** for rs160458, **Table S12** with all phenotypes for rs160459). The Bonferroni significance threshold was p < 9.7**·**10^−6^ (0.05/5179 phenotypes, strict correction not accounting for correlations between phenotypes). A majority of phenotypes are directly related to lateral and superior parts of the temporal cortex, notably the left STS (depth, gray matter volume, surface area and thickness of its banks).

**Table 2.** Top 20 MRI phenotypes associated with rs160459 (significant after Bonferroni correction p < 9·10^−6^, 0.05/5179). (GM: gray matter, DKT: Desikan-Killiany atlas, DST: Destrieux atlas, rfMRI: resting-state functional MRI). Color code: orange: Brainvisa phenotypes, green: Freesurfer phenotypes, blue: diffusion MRI phenotypes, red: rfMRI phenotypes.

| Phenotypes | Pvals | N subjects (British) |
| --- | --- | --- |
| **Mean depth of the left STS** | **$1.4 \cdot 10^{-13}$** | **18101** |
| Surface area of the left temporal pole - DST | $5.8 \cdot 10^{-12}$ | 17127 |
| GM volume of the left banks STS - DKT | $7.6 \cdot 10^{-8}$ | 17127 |
| Surface area of the right cuneus gyrus - DST | $2.3 \cdot 10^{-7}$ | 17127 |
| Surface area of the left banks STS - DKT | $2.8 \cdot 10^{-7}$ | 17127 |
| Fractional anisotropy of the left tapetum of corpus callosum | $5.7 \cdot 10^{-7}$ | 16541 |
| Surface area of the right cuneus - DKT | $6.6 \cdot 10^{-7}$ | 17127 |
| GM thickness of the right middle temporal gyrus - DKT | $6.9 \cdot 10^{-7}$ | 17127 |
| GM thickness of the left banks STS - DKT | $8.8 \cdot 10^{-7}$ | 17127 |
| GM thickness of the left middle temporal - DKT | $1.1 \cdot 10^{-6}$ | 17127 |
| **Maximum depth of the left anterior inferior temporal sulcus** | **$1.5 \cdot 10^{-6}$** | **18101** |
| GM thickness of the left middle temporal gyrus - DST | $1.9 \cdot 10^{-6}$ | 17127 |
| **rfMRI correlation between ICA100: 29 and 41 (edge 1146)** | **$2.1 \cdot 10^{-6}$** | **15859** |
| **Mean depth of the right STS** | **$2.3 \cdot 10^{-6}$** | **18100** |
| Surface area of the right temporal pole - DST | $2.4 \cdot 10^{-6}$ | 17126 |
| GM thickness of the right banks STS - DKT | $6.0 \cdot 10^{-6}$ | 17127 |
| **rfMRI correlation between ICA25: 5 and 18 (edge 87)** | **$7.6 \cdot 10^{-6}$** | **15864** |
| GM volume of the left Jensen sulcus - DST | $7.6 \cdot 10^{-6}$ | 17108 |
| GM volume of the left transverse temporal - DKT | $7.9 \cdot 10^{-6}$ | 17126 |
| **Surface area of the left anterior inferior temporal sulcus** | **$8.6 \cdot 10^{-6}$** | **18101** |

**Fig. S10** shows the mapping of the p-values of the association between rs160459 and brain sulci features (maximum depth, mean depth, sulcal width, surface, gray matter thickness along the sulcus, and sulcus length). This genetic variant clearly has a localized effect on the superior temporal region curvature, with an effect on both the left and right STS as well as on the anterior inferior temporal sulcus. Similarly, regarding cortical features obtained from Freesurfer parcellations (gray volume, surface area, and cortical thickness), the effect is localized to the temporal regions, notably the left banks of the STS in the Desikan atlas (**Fig. S11**) and the left temporal pole and the middle temporal gyrus in the Destrieux atlas (**Fig. S12**). Two main associations were uncovered with diffusion MRI analysis. The most significant association was with the fractional anisotropy (FA) measured in the left tapetum of the corpus callosum, and the second corresponded to the mode of anisotropy (MO) of the left cerebellar peduncle.

Finally, we analyzed resting-state activity. rs160459 modulates several components that comprise linguistic regions (**Fig. S13**, p < 0.01, uncorrected), such as the superior temporal regions, the insula, the inferior frontal region and the basal ganglia (component 18 for ICA25; and components 9 and 38 for ICA100). Associations with ICA100 amplitudes included the left fusiform and associative visual areas (components 52 and 8, respectively). Correlations between resting-state networks revealed associations with components 29-41 and 8-31 for ICA100 and 5-18 for ICA25 (**Fig. S14**), which are implicated in language processing (**Fig. S15**). In particular, **Fig. 3A** presents the most associated pair, 29-41, with rs160459. Notably, component 29 bilaterally comprises the angular and supra-marginal gyri, the STAP and the posterior cingulate. Component 41 includes a string of areas from Brodmann area 19 and the fusiform gyri to the posterior STS.

**Fig. 3.**
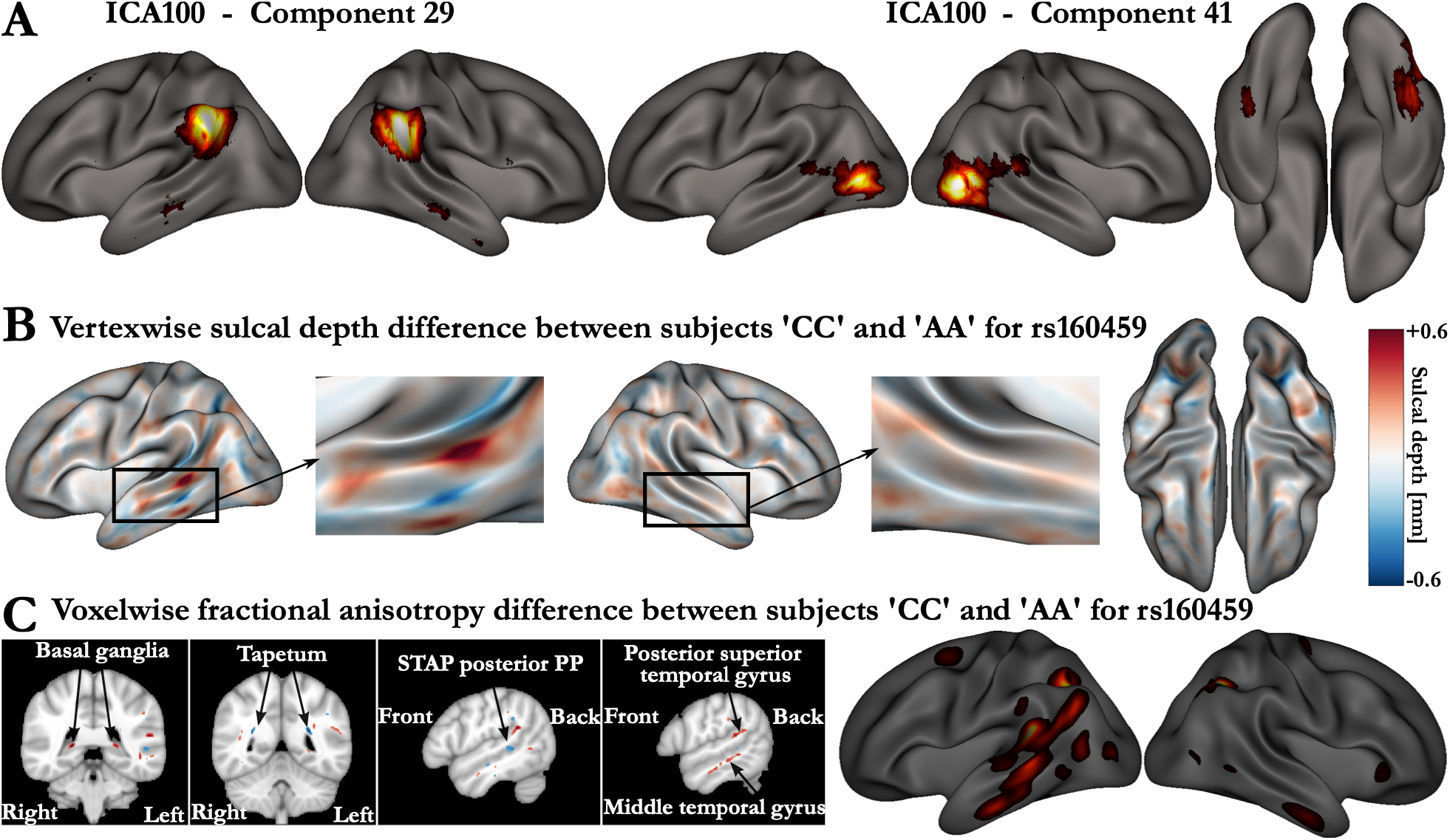
**(A) ICA100 pair (29-41) for which their full correlation has the most significant association with rs160459. (B) Vertexwise sulcal depth difference in rs160459 between ‘CC’ and ‘AA’ subjects**. Red values show higher depth values for CC subjects and blue for AA subjects. **(C) Voxelwise fractional anisotropy difference in rs160459 in the MNI space between ‘CC’ and ‘AA’ subjects**. These figures are thresholded at 0.005 [unitless fractional measure] to underline the main differences. In the right column, these differences in white matter are projected onto the cortex.

### Characterizing the local influence of rs160459 on brain anatomy

The ‘CC’ and ‘AA’ allelic configurations were observed in a roughly similar number of subjects. We compared brain anatomy (DTI FA values, sulcal depth and thickness values) in ‘CC’ vs ‘AA’ subjects. A voxelwise difference in FA values in the MNI space was detected between the 4,742 British subjects with the ‘AA’ allele and the 3,470 subjects with the ‘CC’ allele (**Fig. 3C**, left). These regions are projected on the cortex in **Fig. 3C** (right). The tapetum, which comprises the auditory callosal fibers, is delineated bilaterally, in addition to a region close to the hippocampus. We also noted a difference in FA in the left STS region at the average boundary between gray and white matter, which corresponds to the average location of the posterior PP in the STAP when present. Similarly, **Fig. 3B** highlights the vertexwise difference in sulcal depth between ‘AA’ and ‘CC’ individuals (other configuration differences are shown in **Fig. S16**). The left STAP clearly appears as the region with the most substantial difference in depth between these two groups of individuals. Two other areas display reverse effects relative to the STAP; the first area is in the left intraparietal sulcus, and the second is in the left inferior temporal gyrus along the STAP. In the posterior part of the left STAP relative to ‘CC’ subjects, ‘AA’ subjects have a shallower sulcus surrounded by more pronounced gyri, whereas the white matter is more isotropic under the sulcus and anisotropic under the *planum temporale* and the middle temporal gyrus. The vertexwise gray matter thickness difference between ‘AA’ and ‘CC’ subjects did not produce remarkable differences, although a small cluster in the left inferior temporal gyrus along the STAP is notable (**Fig. S17**).

## Discussion

### A genomic effect focused on the language comprehension network

We explored the genetic associations of a recent evolutionary brain feature, the asymmetry of the depth of the STS, and found that the depth of the left STS was heritable in two different population samples (HCP, UK Biobank from British ancestors) and associated with a genomic region upstream of *DACT1*. This genomic region modulates several other features of the left STS (the gray matter volume, thickness and surface areas of its banks) and of close brain parcels within the left auditory/linguistic network (i.e., Jensen sulcus at the border between the supra-marginal and angular gyri, Heschl’s gyrus volume and thickness of the left middle temporal lobe), as well as, to a lesser extent, their right homologues (i.e., STS depth and thickness). Notably, although we started from an asymmetric feature, rs160459 associations with brain phenotypes were often significant in both hemispheres. However, the most apparent effects were always observed in the left hemisphere. As summarized in **Fig. 3C**, this SNP mainly modulated the macro- and microstructure of the temporal and supra-marginal components of the linguistic network with an effect limited to the left STS for some phenotypes, such as sulcal depth (**Fig. S10-S13**). This tropism for the auditory/linguistic regions was confirmed by resting-state analyses. rs160459 modulated the amplitude of components comprising the classical language areas (superior temporal and inferior frontal regions and the basal ganglia), as indicated by the correlation between two resting-state components (ICA100: 29-41, p ~ 10^−6^). The ICA recovers bilateral components, making it difficult to disentangle the contribution of each hemisphere. Nonetheless, in the left hemisphere, the correlation between components 29-41 is reminiscent of the ventral and dorsal pathway of the reading network. This interpretation related to reading would also be consistent with the significant modulation of the amplitude of the left fusiform gyrus (ICA100: component 52). The relationship between this genomic region and language is also supported by two patients with a 14q23.1 deletion who had moderate to severe expressive language delay (Jiang et al. 2008; Martínez-Frías et al. 2014).

An interesting structure was also highlighted in our analyses: the left tapetum, i.e., the auditory fibers of the corpus callosum. This result may be associated with the observation that patients with corpus callosum agenesis are also the only group with no significant STS depth asymmetry in Leroy et al. (2015). The authors speculated that the dense local connectivity useful for phonetic encoding (DeWitt and Rauschecker 2012) intertwined with many long-range bundles (Turken and Dronkers 2011) (the callosal auditory fibers but also the arcuate fasciculus, inferior occipito-frontal fasciculus, and middle and inferior longitudinal fasciculi), which underlie the left STS and create different tensions and tissue stiffness compared with those in the right hemisphere, explaining the depth asymmetry of this sulcus. Therefore, the elevation of the bottom of the left sulcus, particularly in its posterior part, could be related to a localized higher fiber density, which may eventually result in a PP (**Fig. 2**, **3B**). Among these tracts, the arcuate fasciculus, which is the most left-lateralized tract (Dubois et al. 2010), was not directly extracted in our analyses, but the voxelwise differences between allelic variants below the left STAP could correspond to it.

It is important to note that this locus explains only about 1% of the heritability of the STAP phenotypes. Thus, there are probably many other genetic regions that contribute to this polygenic trait variability. This observation is not unique to the STAP traits and is general to other GWAS in the literature. Common genetic variants identified by GWAS usually explain only a small percentage of the heritability and this issue is known as the missing heritability (Yang et al. 2011). The main current hypothesis assumes that most complex traits are highly polygenic and regulated additively by many variants of small effect size. We believe that this hypothesis also applies to the STAP variability.

### A role in establishing the auditory/linguistic network

The locus uncovered by our genome-wide association is nearest to *DACT1*, a gene mainly active, in human brain tissues, during gestation until the age of two years, a crucial period for language acquisition and social communication. The protein encoded by *DACT1* belongs to the dapper family and is involved in regulating intracellular signaling pathways during development. Among its related pathways is the noncanonical Wnt/planar cell polarity pathway, which is activated in the neuronal differentiation process (Jiao et al. 2018) and synaptogenesis (Rosso et al. 2013). *DACT1* is reportedly involved in excitatory synapse organization, dendrite and synapse formation, establishment of spines in hippocampal pyramidal neurons (Okerlund et al. 2010), and in migration of cortical interneurons (Arguello et al. 2013). Thus, *DACT1* plays a major role in the morphology of pyramidal cells and interneurons. Our results suggest that the uncovered locus regulates *DACT1* expression in blood. Since, blood cis-eQTLs often support cis-eQTLs in brain tissues, providing a sufficient sample size (Qi et al. 2018), this locus may be an eQTL early in brain development when *DACT1* is most highly expressed. Interestingly, the right STS that appears one or two weeks earlier than the left becomes visible around 23-26 wGA (Fontes 1944; Chi et al. 1977; Kasprian et al. 2011; Habas et al. 2012) at the time when the *DACT1* expression level significantly decreases (**Fig. 1C**). Several genes, including some with different levels of expression in humans and chimpanzees, show also differences in expression in the left and right temporal lobes in early gestation, but their expression becomes symmetrical after 19 wGA (Sun et al. 2006) suggesting that many anatomical features are determined long before they are observed. As the brain begins to process external stimuli during the last trimester of gestation, such as sounds and voices, along distinctive pathways (Mahmoudzadeh et al. 2013), and stimulation becomes richer, dendrite development and axonal arborization might complexify the gyrification pattern, and accentuate the initial hemispheric biases.

Furthermore, the uncovered locus regulates *PSMA3-AS1* and *KIAA0586* splicing. *PSMA3-AS1* is a *PSMA3* antisense, the antisense roles are not well characterized, but they may be involved in the regulation of gene expression through RNA hybrids that lead to transcriptional interference (Wight and Werner 2013). For instance, *PSMA3-AS1* can form an RNA duplex with pre-PSMA3, which promotes *PSMA3* and increases its stability (Xu et al. 2019). *KIAA0586*, also known as Talpid3, is required for ciliogenesis (Yin et al. 2009), sonic Hedgehog signaling (Davey et al. 2006), and the asymmetrical localization of CEP210 to daughter centrioles (Kobayashi et al. 2014). Additionally, mutations in *KIAA0586* have been linked to Joubert Syndrome (Bachmann-Gagescu et al. 2015), which is characterized by underdevelopment of the cerebellar vermis, mental retardation with notably motor and language deficit.

In conclusion, the genomic region in linkage disequilibrium with rs160459 modulates the STS depth and other related MRI phenotypes in the temporal lobe. Overall, the identified locus contributes to human variability in the formation of superior temporal networks, notably those involved in language. The link between this locus and the following three candidate genes should be further investigated: *DACT1*, whose gene expression in the brain corresponds with the STS formation timeframes; *KIAA0586*, which is required for sonic Hedgehog signaling and essential for primary ciliogenesis, and whose loss of function causes ciliopathies; and *PSMA3-AS1* whose function remains largely unexplored.

## ACKNOWLEDGMENTS

The present analyses were conducted under UK Biobank data application number 25251. The acknowledgments of IMAGEN contributing consortium authors can be found in the Supplementary Notes.

