## Supplementary Materials for "Enhancer locus in ch14q23.1 modulates brain asymmetric temporal regions involved in language processing"

**This PDF file includes:**

Supplementary Notes

Supplementary Tables (Tables S1 to S5 and S7 to S11)

Supplementary Figures (Figs. S1 to S18)

**Other supplementary materials for this manuscript include the following:**

Table S6

Table S12

### Supplementary Notes

#### Acknowledgments for the IMAGEN consortium

**Principal Investigators.** Tobias Banaschewski M.D., Ph.D.[1]; Gareth J. Barker* Ph.D.[2]; Arun L.W. Bokde Ph.D.[3]; Uli Bromberg Ph.D. [4]; Christian Büchel M.D.[4]; Erin Burke Quinlan, Ph.D.[5]; Sylvane Desrivières Ph.D.[5]; Herta Flor Ph.D.[6,7]; Antoine Grigis Ph.D.[8]; Hugh Garavan Ph.D.[9]; Penny Gowland Ph.D.[10]; Andreas Heinz M.D., Ph.D.[11]; Bernd Ittermann Ph.D.[12]; Jean-Luc Martinot M.D., Ph.D.[13]; Frauke Nees Ph.D.[1,6]; Dimitri Papadopoulos Orfanos Ph.D.[8]; Tomáš Paus M.D., Ph.D.[17]; Luise Poustka M.D.[18] , Sarah Hohmann M.D.[1]; Juliane H. Fröhner Dipl.-Psych.[19]; Michael N. Smolka M.D.[19]; Henrik Walter M.D., Ph.D.[11]; Robert Whelan Ph.D.[20]; Gunter Schumann M.D.[5];

[1]Department of Child and Adolescent Psychiatry and Psychotherapy, Central Institute of Mental Health, Medical Faculty Mannheim, Heidelberg University, Square J5, 68159 Mannheim, Germany;
[2]Department of Neuroimaging, Institute of Psychiatry, Psychology & Neuroscience, King's College London, United Kingdom;
[3]Discipline of Psychiatry, School of Medicine and Trinity College Institute of Neuroscience, Trinity College Dublin;
[4]University Medical Centre Hamburg-Eppendorf, House W34, 3.OG, Martinistr. 52, 20246, Hamburg, Germany;
[5]Medical Research Council - Social, Genetic and Developmental Psychiatry Centre, Institute of Psychiatry, Psychology & Neuroscience, King's College London, United Kingdom;
[6]Department of Cognitive and Clinical Neuroscience, Central Institute of Mental Health, Medical Faculty Mannheim, Heidelberg University, Square J5, Mannheim, Germany;
[7] Department of Psychology, School of Social Sciences, University of Mannheim, 68131 Mannheim, Germany;
[8] NeuroSpin, CEA, Université Paris-Saclay, F-91191 Gif-sur-Yvette, France;
[9]Departments of Psychiatry and Psychology, University of Vermont, 05405 Burlington, Vermont, USA;
[10]Sir Peter Mansfield Imaging Centre School of Physics and Astronomy, University of Nottingham, University Park, Nottingham, United Kingdom;
[11]Charité – Universitätsmedizin Berlin, corporate member of Freie Universität Berlin, Humboldt-Universität zu Berlin, and Berlin Institute of Health, Department of Psychiatry and Psychotherapy, Campus Charité Mitte, Charitéplatz 1, Berlin, Germany; or depending on journal requirements can be Charité – Universitätsmedizin Berlin, Department of Psychiatry and Psychotherapy, Campus Charité Mitte, Charitéplatz 1, Berlin, Germany;
[12]Physikalisch-Technische Bundesanstalt (PTB), Braunschweig and Berlin, Germany [or depending on journal requirements can be: Physikalisch-Technische Bundesanstalt (PTB), Abbestr. 2 - 12, Berlin, Germany;
[13]Institut National de la Santé et de la Recherche Médicale, INSERM Unit 1000 "Neuroimaging & Psychiatry", University Paris Sud, University Paris Descartes - Sorbonne Paris Cité; and Maison de Solenn, Paris, France;
[14]Institut National de la Santé et de la Recherche Médicale, INSERM Unit 1000 "Neuroimaging & Psychiatry", University Paris Sud, University Paris Descartes; Sorbonne Université; and AP-HP, Department of Child and Adolescent Psychiatry, Pitié-Salpêtrière Hospital, Paris, France;
[15]Institut National de la Santé et de la Recherche Médicale, INSERM Unit 1000 "Neuroimaging & Psychiatry", University Paris Sud, University Paris Descartes - Sorbonne Paris Cité; and Psychiatry Department 91G16, Orsay Hospital, France;

[16]Institut National de la Santé et de la Recherche Médicale, INSERM Unit 1000 "Neuroimaging & Psychiatry", Faculté de médecine, Université Paris-Sud, Le Kremlin-Bicêtre; and Université Paris Descartes, Sorbonne Paris Cité, Paris, France;
[17]Bloorview Research Institute, Holland Bloorview Kids Rehabilitation Hospital and Departments of Psychology and Psychiatry, University of Toronto, Toronto, Ontario, M6A 2E1, Canada;
[18]Department of Child and Adolescent Psychiatry and Psychotherapy, University Medical Centre Göttingen, von-Siebold-Str. 5, 37075, Göttingen, Germany;
[19]Department of Psychiatry and Neuroimaging Center, Technische Universität Dresden, Dresden, Germany;
[20]School of Psychology and Global Brain Health Institute, Trinity College Dublin, Ireland;

**Disclosures.** Dr. Banaschewski has served as an advisor or consultant to Bristol-Myers Squibb, Desitin Arzneimittel, Eli Lilly, Medice, Novartis, Pfizer, Shire, UCB, and Vifor Pharma; he has received conference attendance support, conference support, or speaking fees from Eli Lilly, Janssen McNeil, Medice, Novartis, Shire, and UCB; he is involved in clinical trials conducted by Eli Lilly, Novartis, and Shire; and the present work is unrelated to these relationships. Dr. Barker has received honoraria from General Electric Healthcare for teaching scanner programming courses and acts as a consultant for IXICO. The other authors report no biomedical financial interests or potential conflicts of interest.

**Funding supporting the IMAGEN consortium.** This work received support from the following sources: the European Union-funded FP6 Integrated Project IMAGEN (Reinforcement-related behaviour in normal brain function and psychopathology) (LSHM-CT- 2007-037286), the Horizon 2020 funded ERC Advanced Grant 'STRATIFY' (Brain network based stratification of reinforcement-related disorders) (695313), ERANID (Understanding the Interplay between Cultural, Biological and Subjective Factors in Drug Use Pathways) (PR-ST-0416-10004), BRIDGET (JPND: BRain Imaging, cognition Dementia and next generation GEnomics) (MR/N027558/1), the FP7 projects IMAGEMEND(602450; IMAging GEnetics for MENtal Disorders) and MATRICS (603016), the Innovative Medicine Initiative Project EU-AIMS (115300-2), the Medical Research Council Grant 'c-VEDA' (Consortium on Vulnerability to Externalizing Disorders and Addictions) (MR/N000390/1), the Swedish Research Council FORMAS, the Medical Research Council, the National Institute for Health Research (NIHR) Biomedical Research Centre at South London and Maudsley NHS Foundation Trust and King's College London, the Bundesministeriumfür Bildung und Forschung (BMBF grants 01GS08152; 01EV0711; eMED SysAlc01ZX1311A; Forschungsnetz AERIAL 01EE1406A, 01EE1406B), the Deutsche Forschungsgemeinschaft (DFG grants SM 80/7-2, SFB 940/2), the Medical Research Foundation and Medical research council (grant MR/R00465X/1). Further support was provided by grants from: ANR (project AF12-NEUR0008-01 - WM2NA, and ANR-12-SAMA-0004), the Fondation de France, the Fondation pour la Recherche Médicale, the Mission Interministérielle de Lutte-contre-les-Drogues-et-les-Conduites-Addictives (MILDECA), the Assistance-Publique-Hôpitaux-de-Paris and INSERM (interface grant), Paris Sud University IDEX 2012; the National Institutes of Health, Science Foundation Ireland (16/ERCD/3797), U.S.A. (Axon, Testosterone and Mental Health during Adolescence; RO1 MH085772-01A1), and by NIH Consortium grant U54 EB020403, supported by a cross-NIH alliance that funds Big Data to Knowledge Centres of Excellence.

### Supplementary Tables

**Table S1. Heritability estimates of the STAP phenotypes in three different populations.** NB: The Human Connectome Project is composed of related twins and siblings constituting an extended pedigree of 820 individuals, while the UK Biobank is composed of unrelated individuals. STAP = superior temporal asymmetrical pit, AI = asymmetry index, depth PF = depth potential function, PP = *pli de passage*, p-values assessing the significance of heritability estimates are written within parentheses.

| Phenotype/Cohort | HCP (820 Caucasian subjects, extended pedigree) | UKB (16,515 British subjects, unrelated individuals) | IMAGEN (1,726 European adolescents, unrelated individuals) |
|---|---|---|---|
| Left STAP geodesic depth | $h^2 = 0.48\pm0.06$ ($1.8\cdot10^{-15}$) | $h^2 = 0.21\pm0.03$ ($8.9\cdot10^{-11}$) | $h^2 = 0.24\pm0.21$ (0.13) |
| Right STAP geodesic depth | $h^2 = 0.46\pm0.06$ ($2.2\cdot10^{-13}$) | $h^2 = 0.15\pm0.03$ ($1.8\cdot10^{-6}$) | $h^2 = 0.00\pm0.21$ (0.5) |
| AI of the STAP geodesic depth | $h^2 = 0.30\pm0.06$ ($5\cdot10^{-7}$) | $h^2 = 0.09\pm0.03$ (0.0021) | $h^2 = 0.48\pm0.21$ (0.012) |
| Left STAP depth PF | $h^2 = 0.25\pm0.06$ ($3.2\cdot10^{-5}$) | $h^2 = 0.18\pm0.03$ ($4.9\cdot10^{-8}$) | $h^2 = 0.35\pm0.21$ (0.050) |
| Right STAP depth PF | $h^2 = 0.38\pm0.06$ ($4.7\cdot10^{-10}$) | $h^2 = 0.04\pm0.03$ (0.1198) | $h^2 = 0.16\pm0.21$ (0.23) |
| AI of the STAP depth PF | $h^2 = 0.22\pm0.06$ ($1.8\cdot10^{-4}$) | $h^2 = 0.11\pm0.03$ (0.0004) | $h^2 = 0.44\pm0.21$ (0.022) |
| Presence of PP in the left STAP | $h^2 = 0.53\pm0.09$ ($1\cdot10^{-7}$) | $h^2 = 0.16\pm0.03$ ($5.2\cdot10^{-7}$) | $h^2 = 0.15\pm0.21$ (0.24) |
| Presence of PP in the right STAP | $h^2 = 0.27\pm0.15$ (0.03) | $h^2 = 0.01\pm0.03$ (0.4308) | $h^2 = 0.00\pm0.21$ (0.5) |

**Table S2. Proportion of phenotypic variance explained and proportion of heritability explained by rs160459 allelic dosage in UK Biobank British.** Significant heritability estimates $p < 6.25\cdot10^{-3}$ (0.05/8) are indicated by *. (AI: asymmetry index, depth PF: depth potential function, PP: *pli de passage*)

| Phenotype | Heritability | Proportion of variance explained in phenotype | Proportion of heritability explained |
|---|---|---|---|
| Left STAP geodesic depth | 21%* | 0.29% | 1.4% |
| Right STAP geodesic depth | 15%* | 0.17% | 1.1% |
| AI of the STAP geodesic depth | 9% | 0.06% | Non-significant $h^2$ |
| Left STAP depth PF | 18%* | 0.18% | 1.0% |
| Right STAP depth PF | 4% | 0.13% | Non-significant $h^2$ |
| AI of the STAP depth PF | 11% | 0.05% | Non-significant $h^2$ |
| Presence of PP in the left STAP | 16%* | 0.16% | 1.0% |
| Presence of PP in the right STAP | 1% | 0.16% | Non-significant $h^2$ |

**Table S3. Association details between rs160459 and left STAP geodesic depth with the MRI center added as a covariate.** *Subjects included in the analysis were scanned at two different sites using two identical scanners and the same protocol. When controlling for the center the association at the main SNP remained unchanged, even though the center covariate was associated with the phenotype. This result is due to the association of this phenotype with age and sex, which accounts for the difference in phenotype between centers. Indeed, the second center was set up later than the first one, and subjects with a similar age were recruited at the first time point of the UK Biobank. The subjects included in the two centers should have the same average birth year, but because the first center scanned a major proportion of the subjects earlier, the subjects were younger at the time of the MRI scan.*

| CHR | SNP | BP | A1 | TEST | NMISS | BETA | STAT | P |
|---|---|---|---|---|---|---|---|---|
| 14 | rs160459 | 59074136 | C | ADD | 16515 | 0.2673 | 6.923 | 4.573E-12 |
| 14 | rs160459 | 59074136 | C | Sex | 16515 | -0.4127 | -7.569 | 3.964E-14 |
| 14 | rs160459 | 59074136 | C | Age | 16515 | -0.023 | -6.264 | 3.834E-10 |
| 14 | rs160459 | 59074136 | C | PC1 | 16515 | 0.02285 | 1.304 | 0.1923 |
| 14 | rs160459 | 59074136 | C | PC2 | 16515 | 0.02042 | 1.114 | 0.2653 |
| 14 | rs160459 | 59074136 | C | PC3 | 16515 | 0.01127 | 0.6384 | 0.5232 |
| 14 | rs160459 | 59074136 | C | PC4 | 16515 | 0.0004496 | 0.03371 | 0.9731 |
| 14 | rs160459 | 59074136 | C | PC5 | 16515 | -0.007925 | -1.341 | 0.1798 |
| 14 | rs160459 | 59074136 | C | PC6 | 16515 | -0.006324 | -0.3754 | 0.7073 |
| 14 | rs160459 | 59074136 | C | PC7 | 16515 | 0.007391 | 0.4848 | 0.6279 |
| 14 | rs160459 | 59074136 | C | PC8 | 16515 | -0.009206 | -0.6063 | 0.5443 |
| 14 | rs160459 | 59074136 | C | PC9 | 16515 | 0.00415 | 0.5818 | 0.5607 |
| 14 | rs160459 | 59074136 | C | PC10 | 16515 | -0.03203 | -2.421 | 0.01551 |
| 14 | rs160459 | 59074136 | C | Array | 16515 | 0.1291 | 1.423 | 0.1549 |
| 14 | rs160459 | 59074136 | C | MRI Center | 16515 | 0.6519 | 8.269 | 1.454E-16 |

**Table S4. Association details between rs160459 and the left STAP geodesic depth with eTIV added as covariate.**

| CHR | SNP | BP | A1 | TEST | NMISS | BETA | STAT | P |
|---|---|---|---|---|---|---|---|---|
| 14 | rs160459 | 59074136 | C | ADD | 16515 | 0.2672 | 6.924 | 4.56E-12 |
| 14 | rs160459 | 59074136 | C | Sex | 16515 | -0.4663 | -8.03 | 1.039E-15 |
| 14 | rs160459 | 59074136 | C | Age | 16515 | -0.02255 | -6.137 | 8.623E-10 |
| 14 | rs160459 | 59074136 | C | PC1 | 16515 | 0.02269 | 1.295 | 0.1952 |
| 14 | rs160459 | 59074136 | C | PC2 | 16515 | 0.01977 | 1.079 | 0.2808 |
| 14 | rs160459 | 59074136 | C | PC3 | 16515 | 0.01193 | 0.6758 | 0.4992 |
| 14 | rs160459 | 59074136 | C | PC4 | 16515 | 0.0009621 | 0.07214 | 0.9425 |
| 14 | rs160459 | 59074136 | C | PC5 | 16515 | -0.008107 | -1.372 | 0.17 |
| 14 | rs160459 | 59074136 | C | PC6 | 16515 | -0.006904 | -0.4099 | 0.6819 |
| 14 | rs160459 | 59074136 | C | PC7 | 16515 | 0.007103 | 0.4659 | 0.6413 |
| 14 | rs160459 | 59074136 | C | PC8 | 16515 | -0.009703 | -0.6391 | 0.5228 |
| 14 | rs160459 | 59074136 | C | PC9 | 16515 | 0.00415 | 0.582 | 0.5606 |
| 14 | rs160459 | 59074136 | C | PC10 | 16515 | -0.032 | -2.418 | 0.0156 |
| 14 | rs160459 | 59074136 | C | Array | 16515 | 0.1252 | 1.38 | 0.1676 |
| 14 | rs160459 | 59074136 | C | MRI center | 16515 | 0.6728 | 8.494 | 2.161E-17 |
| 14 | rs160459 | 59074136 | C | eTIV | 16515 | 3.54E-07 | 2.68 | 0.00737 |

**Table S5. P-values of the association of rs160458 with STAP phenotypes in three populations.** Significant heritability estimates $p < 6.25 \cdot 10^{-3}$ (0.05/8) are indicated by *. (AI: asymmetry index, depth PF: depth potential function, PP: *pli de passage*)

| **Phenotype/Cohort** | **Heritability** | | **UKB (16,203 subjects, British)** | **IMAGEN (1,726 subjects)** | **UKB (2,097 subjects, not British)** |
|---|---|---|---|---|---|
| | **HCP** | **UKB** | | | |
| Left STAP geodesic depth | 48%* | 20%* | $p = 2.0 \cdot 10^{-12}$ | $p = 1.8 \cdot 10^{-5}$ | p = 0.0308 |
| Right STAP geodesic depth | 46%* | 19%* | $p = 6.9 \cdot 10^{-8}$ | $p = 5.2 \cdot 10^{-5}$ | p = 0.1846 |
| AI of the STAP geodesic depth | 30%* | 11% | p = 0.0026 | p = 0.0371 | p = 0.2586 |
| Left STAP depth PF | 25%* | 18%* | $p = 1.5 \cdot 10^{-7}$ | p = 0.0181 | p = 0.1777 |
| Right STAP depth PF | 38%* | 8% | $p = 4.0 \cdot 10^{-6}$ | p = 0.1636 | p = 0.0511 |
| AI of the STAP depth PF | 22%* | 12% | p = 0.0105 | p = 0.1366 | p = 0.6358 |
| Presence of PP in the left STAP | 53%* | 17%* | $p = 2.3 \cdot 10^{-7}$ | p = 0.0060 | p = 0.3669 |
| Presence of PP in the right STAP | 27% | 0% | $p = 1.2 \cdot 10^{-7}$ | p = 0.0019 | p = 0.3229 |

**Table S6 (separate file). eQTL summary from Brain-eMeta for the genome wide significant SNPs in the ch14q23.1 locus.**

SNPs list: ['rs170239', 'rs221326', 'rs468213', 'rs149142', 'rs17094771', 'rs10782438', 'rs186347', 'rs160460', 'rs160459', 'rs160458', 'rs4898962', 'rs311814', 'rs543709795', 'rs311813', 'rs1742882', 'rs5808978', 'rs191103', 'rs128016']

Brain-eMeta eQTL summary data (Qi et al. 2018), which is a set of eQTL data from a meta-analysis of GTEx brain(GTEx Consortium 2017), CMC(Fromer et al. 2016), and ROSMAP (Ng et al. 2017). The estimated effective is n = 1,194 samples. Only SNPs within 1 Mb distance from each probe are available. This dataset is available on the SMR website (cnsgenomics.com/software/smr).

**Table S7. eQTL association between genome-wide significant SNPs and *DACT1* in eQTLGen.**

| SNP | Chr | SNP Pos | Assessed Allele | Other Allele | Gene Symbol | Nb Cohorts | Nb Samples | Pvalue | FDR |
|---|---|---|---|---|---|---|---|---|---|
| rs10782438 | 14 | 59072144 | C | T | DACT1 | 35 | 31086 | 5.4E-06 | 0.02 |
| rs160459 | 14 | 59074136 | C | A | DACT1 | 34 | 30757 | 9.89E-05 | 0.22 |
| rs160458 | 14 | 59074878 | C | T | DACT1 | 32 | 30304 | 0.0001093 | 0.24 |
| rs186347 | 14 | 59072226 | T | G | DACT1 | 33 | 30641 | 0.0001764 | 0.35 |
| rs149142 | 14 | 59071725 | C | T | DACT1 | 36 | 31470 | 0.0002091 | 0.39 |
| rs311813 | 14 | 59086392 | A | G | DACT1 | 36 | 31470 | 0.0002251 | 0.42 |
| rs191103 | 14 | 59092075 | T | C | DACT1 | 36 | 31459 | 0.0002316 | 0.42 |
| rs4898962 | 14 | 59079971 | C | T | DACT1 | 36 | 31470 | 0.0002666 | 0.46 |
| rs1742882 | 14 | 59086975 | A | G | DACT1 | 36 | 31470 | 0.0003078 | 0.51 |
| rs468213 | 14 | 59071100 | G | C | DACT1 | 35 | 31347 | 0.0003286 | 0.53 |
| rs311814 | 14 | 59085726 | T | C | DACT1 | 35 | 31355 | 0.0004863 | 0.66 |
| rs221326 | 14 | 59069053 | T | G | DACT1 | 32 | 30378 | 0.000494 | 0.66 |
| rs128016 | 14 | 59092581 | C | G | DACT1 | 35 | 31341 | 0.0005614 | 0.7 |
| rs170239 | 14 | 59064739 | T | C | DACT1 | 34 | 31025 | 0.0007291 | 0.78 |
| rs160460 | 14 | 59072964 | G | A | DACT1 | 34 | 31021 | 0.0010307 | 0.87 |

**Table S8. Significant splicing QTLs among genome-wide significant SNPs found on GTEx (October 2019).**

| Gene Symbol | SNP Id | Intron Id | P-Value | NES | Tissue |
|---|---|---|---|---|---|
| PSMA3-AS1 | rs170239 | 58295811:58297955:clu_20725 | 3.8E-10 | 0.33 | Artery - Tibial |
| PSMA3-AS1 | rs170239 | 58295811:58297955:clu_18042 | 2.8E-07 | 0.52 | Ovary |
| KIAA0586 | rs170239 | 58543967:58547781:clu_23844 | 3.7E-07 | -0.31 | Thyroid |
| PSMA3-AS1 | rs170239 | 58295811:58297955:clu_22647 | 4.8E-07 | 0.33 | Breast - Mammary Tissue |
| PSMA3-AS1 | rs170239 | 58295811:58297955:clu_19747 | 5.7E-07 | 0.35 | Heart - Atrial Appendage |
| PSMA3-AS1 | rs170239 | 58295811:58297955:clu_22507 | 6.1E-07 | 0.27 | Adipose - Subcutaneous |
| KIAA0586 | rs170239 | 58543967:58547781:clu_23033 | 1.7E-06 | -0.28 | Skin - Sun Exposed (Lower Leg) |
| PSMA3-AS1 | rs221326 | 58295811:58297955:clu_22507 | 3.3E-06 | 0.26 | Adipose - Subcutaneous |
| PSMA3-AS1 | rs468213 | 58295811:58297955:clu_22647 | 9E-09 | 0.38 | Breast - Mammary Tissue |
| KIAA0586 | rs468213 | 58543967:58547781:clu_23033 | 1.3E-08 | -0.34 | Skin - Sun Exposed (Lower Leg) |
| KIAA0586 | rs468213 | 58540136:58543883:clu_20751 | 5.3E-08 | -0.31 | Artery - Tibial |
| PSMA3-AS1 | rs468213 | 58295811:58297955:clu_20725 | 1E-07 | 0.29 | Artery - Tibial |
| KIAA0586 | rs468213 | 58543967:58547781:clu_22532 | 5.1E-07 | -0.29 | Adipose - Subcutaneous |

| | | | | | |
|---|---|---|---|---|---|
| PSMA3-AS1 | rs468213 | 58295811:58297955:clu_20802 | 5.3E-07 | 0.31 | Esophagus - Mucosa |
| KIAA0586 | rs468213 | 58543967:58547781:clu_23844 | 1.4E-06 | -0.3 | Thyroid |
| KIAA0586 | rs468213 | 58543967:58547781:clu_38797 | 5.7E-06 | -0.37 | Testis |
| KIAA0586 | rs149142 | 58543967:58547781:clu_23033 | 7.7E-09 | -0.34 | Skin - Sun Exposed (Lower Leg) |
| PSMA3-AS1 | rs149142 | 58295811:58297955:clu_20725 | 1.1E-08 | 0.31 | Artery - Tibial |
| KIAA0586 | rs149142 | 58540136:58543883:clu_20751 | 9.1E-08 | -0.31 | Artery - Tibial |
| PSMA3-AS1 | rs149142 | 58295811:58297955:clu_22647 | 1.1E-07 | 0.35 | Breast - Mammary Tissue |
| PSMA3-AS1 | rs149142 | 58295811:58297955:clu_18042 | 1.7E-07 | 0.54 | Ovary |
| PSMA3-AS1 | rs149142 | 58295811:58297955:clu_20802 | 6.3E-07 | 0.31 | Esophagus - Mucosa |
| KIAA0586 | rs149142 | 58543967:58547781:clu_23844 | 1.2E-06 | -0.3 | Thyroid |
| PSMA3-AS1 | rs149142 | 58295811:58297955:clu_23817 | 2.6E-06 | 0.26 | Thyroid |
| PSMA3-AS1 | rs149142 | 58295811:58297955:clu_22507 | 3.2E-06 | 0.26 | Adipose - Subcutaneous |
| KIAA0586 | rs17094771 | 58543967:58547781:clu_23033 | 2.4E-09 | -0.36 | Skin - Sun Exposed (Lower Leg) |
| PSMA3-AS1 | rs17094771 | 58295811:58297955:clu_20725 | 2.4E-08 | 0.3 | Artery - Tibial |
| PSMA3-AS1 | rs17094771 | 58295811:58297955:clu_18042 | 3E-08 | 0.57 | Ovary |
| PSMA3-AS1 | rs17094771 | 58295811:58297955:clu_22647 | 6.9E-08 | 0.37 | Breast - Mammary Tissue |
| KIAA0586 | rs17094771 | 58540136:58543883:clu_20751 | 1.5E-07 | -0.3 | Artery - Tibial |
| PSMA3-AS1 | rs17094771 | 58295811:58297955:clu_20802 | 4.1E-07 | 0.32 | Esophagus - Mucosa |
| PSMA3-AS1 | rs17094771 | 58295811:58297955:clu_22507 | 5.6E-07 | 0.28 | Adipose - Subcutaneous |
| KIAA0586 | rs17094771 | 58543967:58547781:clu_23844 | 6.9E-07 | -0.31 | Thyroid |
| PSMA3-AS1 | rs10782438 | 58295811:58297955:clu_18042 | 6.4E-08 | 0.57 | Ovary |
| KIAA0586 | rs10782438 | 58543967:58547781:clu_23033 | 1.3E-07 | -0.32 | Skin - Sun Exposed (Lower Leg) |
| PSMA3-AS1 | rs10782438 | 58295811:58297955:clu_22647 | 6.2E-07 | 0.35 | Breast - Mammary Tissue |
| PSMA3-AS1 | rs10782438 | 58295811:58297955:clu_20725 | 1.3E-06 | 0.27 | Artery - Tibial |
| KIAA0586 | rs160460 | 58543967:58547781:clu_23033 | 6.3E-07 | -0.3 | Skin - Sun Exposed (Lower Leg) |
| PSMA3-AS1 | rs160460 | 58295811:58297955:clu_18042 | 6.5E-07 | 0.51 | Ovary |
| PSMA3-AS1 | rs160460 | 58295811:58297955:clu_20725 | 2.3E-06 | 0.26 | Artery - Tibial |
| PSMA3-AS1 | rs160459 | 58295811:58297955:clu_23817 | 1.9E-06 | 0.27 | Thyroid |
| PSMA3-AS1 | rs160459 | 58295811:58297955:clu_22507 | 2.2E-06 | 0.27 | Adipose - Subcutaneous |
| KIAA0586 | rs160459 | 58543967:58547781:clu_23033 | 3.1E-06 | -0.29 | Skin - Sun Exposed (Lower Leg) |

**Table S9. CADD score and annotation of genome wide significant variants in the *DACT1* locus.**

| SNP | Location | Allele | Consequence | Feature_type | Feature | BIOTYPE | Ancestral Allele | CADD |
|---|---|---|---|---|---|---|---|---|
| rs170239 | 14:59064739 | T | intergenic_variant | - | - | - | T | 1.832 |
| rs221326 | 14:59069053 | A | intergenic_variant | - | - | - | T | 1.551 |
| rs468213 | 14:59071100 | G | intergenic_variant | - | - | - | C | 0.061 |
| rs149142 | 14:59071725 | C | intergenic_variant | - | - | - | T | 2.913 |
| rs17094771 | 14:59072121 | A | intergenic_variant | - | - | - | A | 0.072 |
| rs10782438 | 14:59072144 | C | intergenic_variant | - | - | - | C | 6.361 |
| rs186347 | 14:59072226 | T | regulatory_region_variant | RegulatoryFeature | ENSR00001071726 | enhancer | T | 1.569 |
| rs160460 | 14:59072964 | G | intergenic_variant | - | - | - | G | 0.6 |
| rs160459 | 14:59074136 | C | regulatory_region_variant | RegulatoryFeature | ENSR00001071727 | promoter_flanking_region | C | 14.66 |
| rs160458 | 14:59074878 | C | regulatory_region_variant | RegulatoryFeature | ENSR00001071727 | promoter_flanking_region | C | 0.852 |
| rs4898962 | 14:59079971 | C | intergenic_variant | - | - | - | T | 2.881 |
| rs311814 | 14:59085726 | T | intergenic_variant | - | - | - | C | 1.026 |
| rs543709795 | 14:59086337 | A | intergenic_variant | - | - | - | - | 1.452 |
| rs311813 | 14:59086392 | A | intergenic_variant | - | - | - | A | 0.151 |
| rs1742882 | 14:59086975 | A | regulatory_region_variant | RegulatoryFeature | ENSR00001071729 | enhancer | A | 2.368 |
| rs5808978 | 14:59089363 | - | regulatory_region_variant | RegulatoryFeature | ENSR00001071729 | enhancer | AG | - |
| rs191103 | 14:59092075 | T | regulatory_region_variant | RegulatoryFeature | ENSR00001071730 | promoter_flanking_region | T | - |
| rs128016 | 14:59092581 | C | regulatory_region_variant | RegulatoryFeature | ENSR00001071730 | promoter_flanking_region | G | 6.602 |

**Table S10. Motif feature predicted consequences for rs160459, available from Ensembl.**

http://uswest.ensembl.org/Homo_sapiens/Variation/Mappings?db=core;r=14:58606918-58607918;v=rs160459;vdb=variation;vf=76598670

**Motif feature consequences**

| Binding matrix | Allele | Consequence type | Transcription factors | Motif position | High information position | Motif score change |
|---|---|---|---|---|---|---|
| ENSPFM0043 | C | TF binding site | CUX1 | 17 (out of 17) | No | ▼ |
| ENSPFM0104 | C | TF binding site | ETV2::PAX5, ELK1::PAX1, ELK1::PAX5, ELK1::PAX9 | 20 (out of 20) | No | ▼ |
| ENSPFM0133 | C | TF binding site | ETV2::BHLHA15, FLI1::BHLHA15 | 12 (out of 15) | No | ▼ |
| ENSPFM0202 | C | TF binding site | FOXG1, FOXK1 | 9 (out of 10) | Yes | ▼ |
| ENSPFM0264 | C | TF binding site | GCM2::SOX15 | 4 (out of 21) | No | ▲ |
| ENSPFM0329 | C | TF binding site | HOXB2::SOX15 | 4 (out of 18) | No | ▼ |

**Table S11. Top 20 MRI phenotypes associated with rs160458 (significant hits after Bonferroni correction with $p < 9 \cdot 10^{-6}$, 0.05/5179).** (GM: gray matter, DKT: Desikan-Killiany atlas, DST: Destrieux atlas, rfMRI: resting-state functional MRI). Color code: orange: Brainvisa phenotypes, green: Freesurfer phenotypes, blue: diffusion MRI phenotypes, red: rfMRI phenotypes.

| Phenotypes | Pvals | N subjects (British) |
|---|---|---|
| **Mean depth of the left STS** | $2.3 \cdot 10^{-13}$ | 17763 |
| **Surface area of the left pole temporal - DST** | $2.2 \cdot 10^{-10}$ | 16800 |
| **GM volume of the left banks STS - DKT** | $6.7 \cdot 10^{-8}$ | 16800 |
| **Fractional anisotropy of the left tapetum of corpus callosum** | $1.7 \cdot 10^{-7}$ | 16228 |
| **Surface area of the left banks STS - DKT** | $2.0 \cdot 10^{-7}$ | 16800 |
| **Maximum depth of the left anterior inferior temporal sulcus** | $2.0 \cdot 10^{-6}$ | 17763 |
| **GM thickness of the left banks STS - DKT** | $2.3 \cdot 10^{-6}$ | 16800 |
| **Surface area of the right cuneus - DST** | $3.3 \cdot 10^{-6}$ | 16800 |
| **Surface area of the right cuneus - DKT** | $5.9 \cdot 10^{-6}$ | 16800 |
| **rfMRI correlation between ICA100: 29 and 41 (edge 1146)** | $7.6 \cdot 10^{-6}$ | 15556 |
| **rfMRI correlation between ICA100: 8 and 36 (edge 385)** | $9.2 \cdot 10^{-6}$ | 15560 |
| **rfMRI correlation between ICA25: 5 and 18 (edge 87)** | $9.5 \cdot 10^{-6}$ | 15561 |
| **GM volume of the right supramarginal - DKT** | $1.0 \cdot 10^{-5}$ | 16800 |
| **Surface area of the right parietal inferior supramarginal gyrus - DST** | $1.2 \cdot 10^{-5}$ | 16800 |
| **GM volume of the right parietal inferior supramarginal gyrus - DST** | $1.2 \cdot 10^{-5}$ | 16800 |
| **GM volume of the left interm primary Jensen sulcus - DST** | $1.4 \cdot 10^{-5}$ | 16781 |
| **GM thickness of the left middle temporal - DKT** | $1.4 \cdot 10^{-5}$ | 16800 |
| **GM volume of the left transverse temporal - DKT** | $1.6 \cdot 10^{-5}$ | 16799 |
| **Surface area of the right temporal pole - DST** | $1.7 \cdot 10^{-5}$ | 16799 |
| **GM thickness of the left middle temporal gyrus - DST** | $1.8 \cdot 10^{-5}$ | 16800 |

**Table S12 (separate file). P-values of rs160459 phenome-wide association (PheWAs) with the 5179 MRI phenotypes.** Significant heritability estimates have $p < 9.7 \cdot 10^{-6}$ (0.05/5179).

### Supplementary Figures

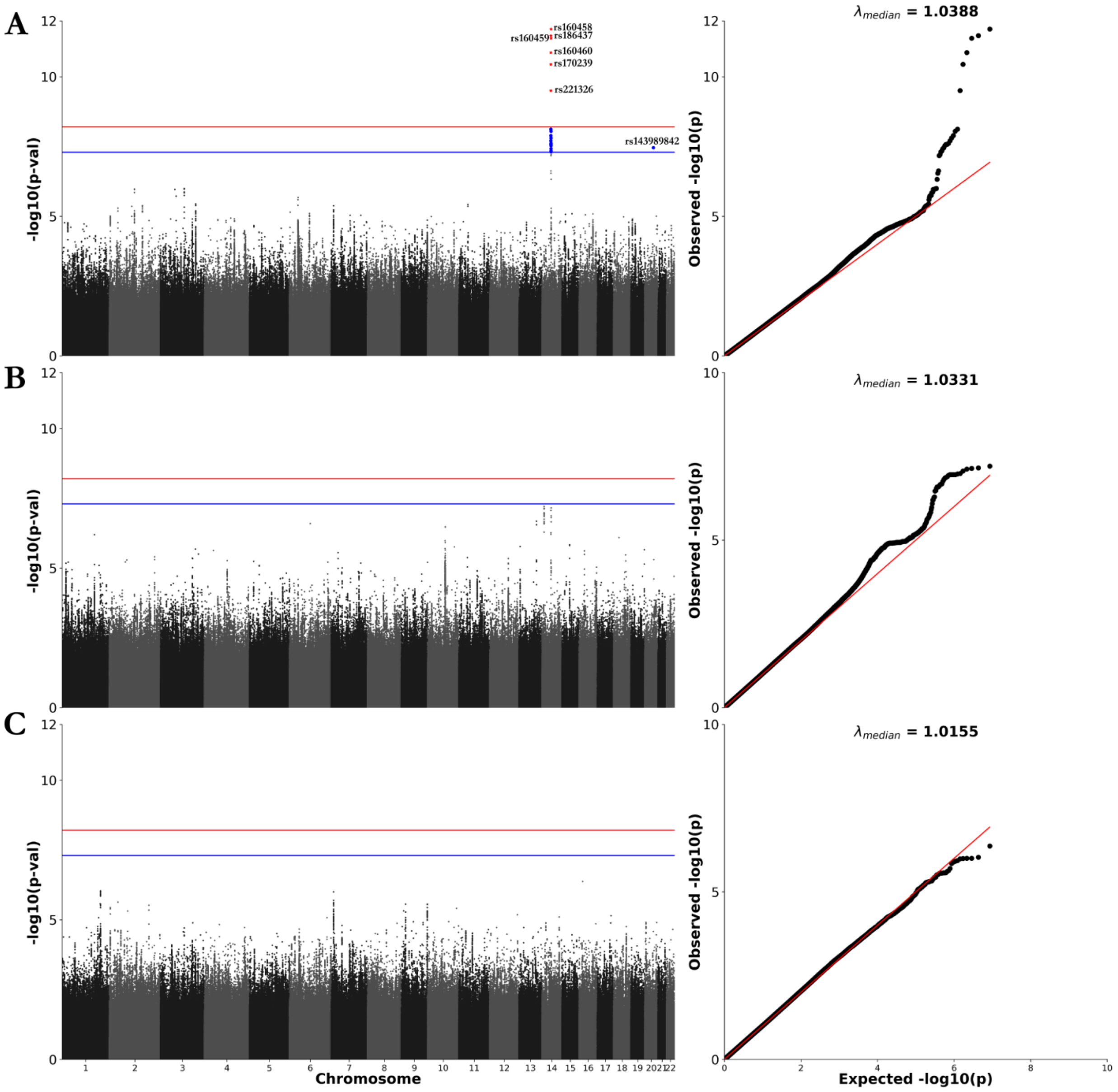


**Fig. S1. Manhattan and QQ plots for the left STAP geodesic depth (A), the right STAP geodesic depth (B) and the asymmetry index(C) of the STAP geodesic depth.** Blue lines correspond to the genomic thresholds $p = 5{\cdot}10^{-8}$ and red lines to the Bonferroni corrected thresholds accounting for the number of phenotypes ($p < 6.25{\cdot}10^{-9} = 5{\cdot}10^{-8}/8$).

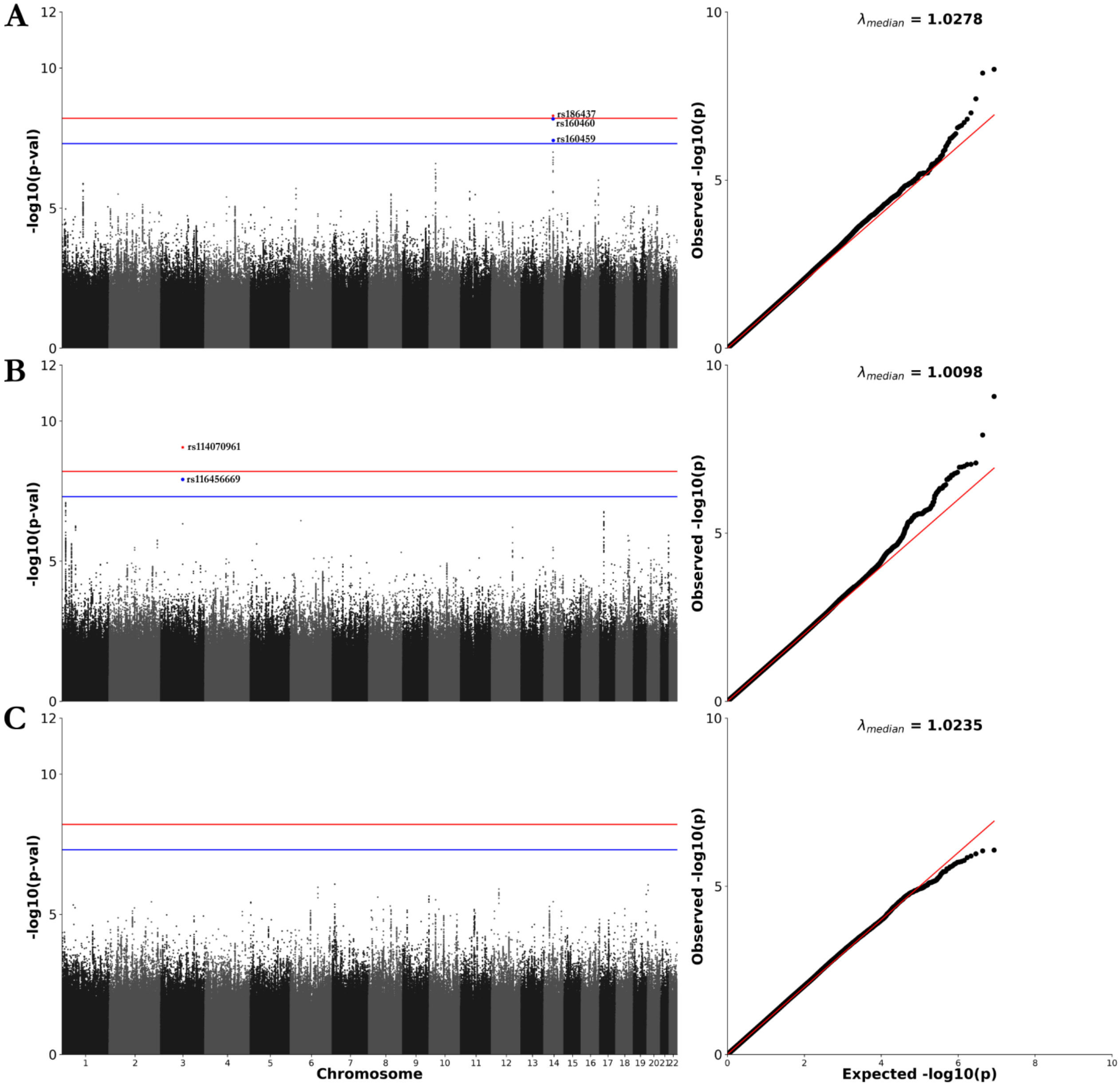


**Fig. S2. Manhattan and QQ plots for the left STAP depth PF (A), the right STAP depth PF (B) and the asymmetry index(C) of the STAP depth PF. (depth PF = depth potential function accounting for both sulcal depth and convexity.)** Blue lines correspond to the genomic thresholds $p = 5{\cdot}10^{-8}$ and red lines to the Bonferroni corrected thresholds accounting for the number of phenotypes ($p < 6.25{\cdot}10^{-9} = 5{\cdot}10^{-8}/8$).

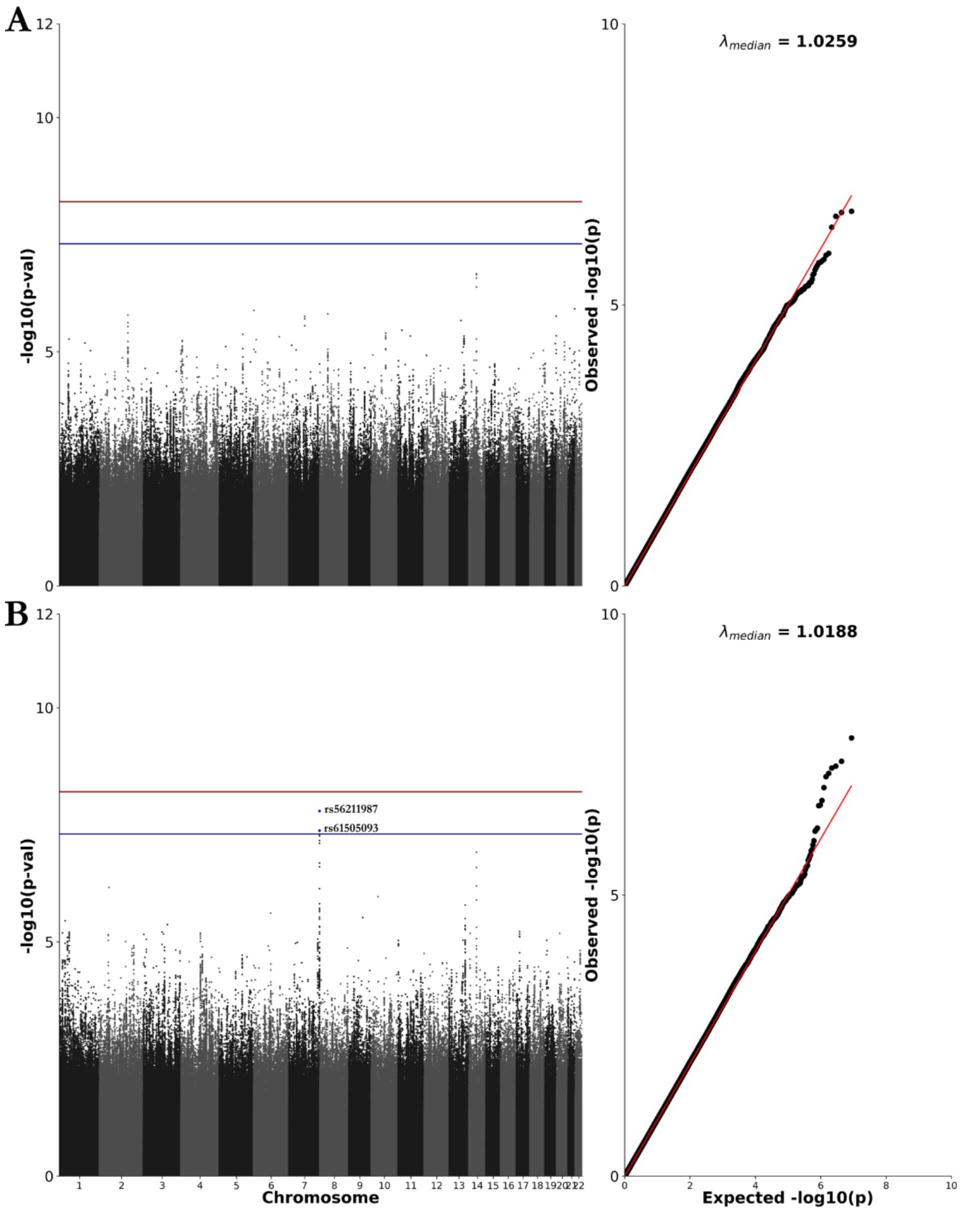


**Fig. S3. Manhattan and QQ plots for the presence of *pli de passage* in the left STAP (A) and the right STAP (B).** Blue lines correspond to the genomic thresholds $p = 5{\cdot}10^{-8}$ and red lines to the Bonferroni corrected thresholds accounting for the number of phenotypes ($p < 6.25{\cdot}10^{-9} = 5{\cdot}10^{-8}/8$).

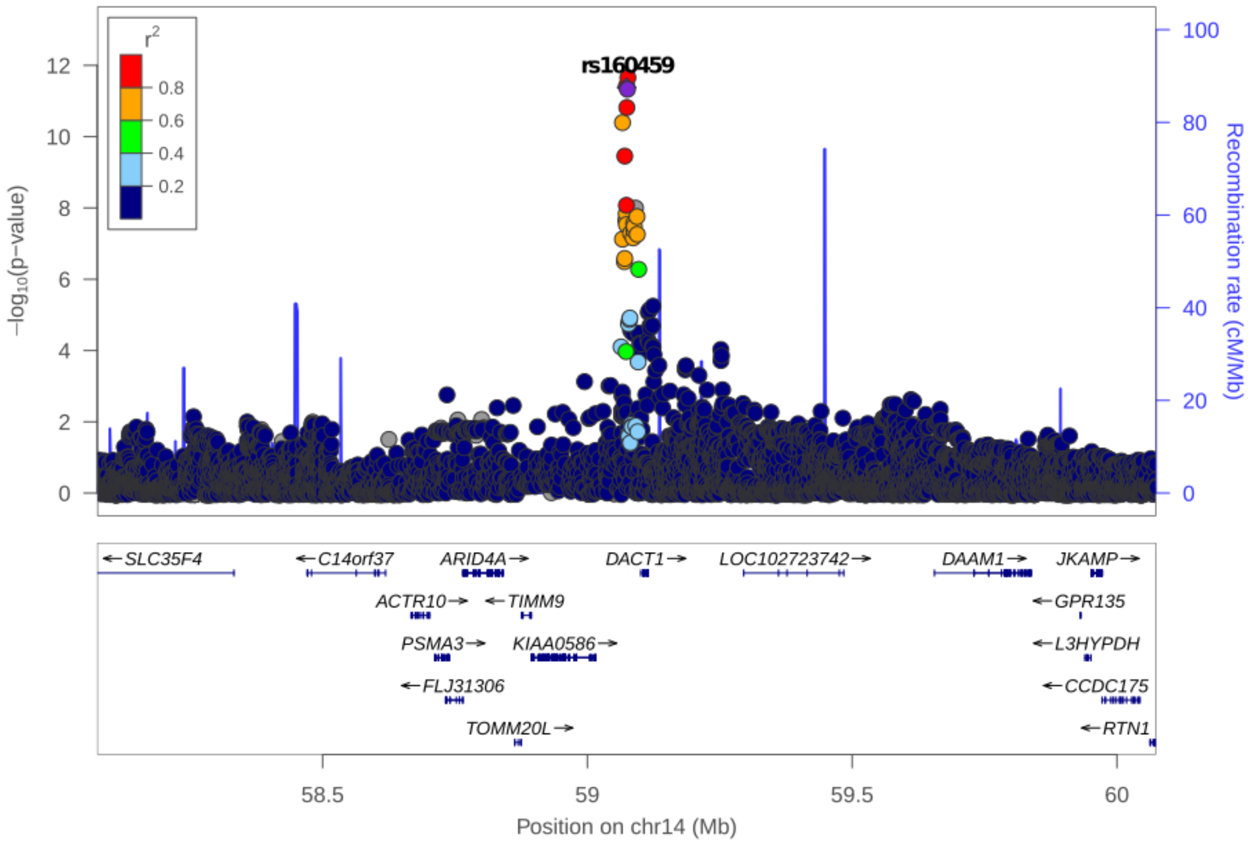


**Fig. S4. Locus plot association of the left STAP geodesic depth in the UKB British ancestry (+/- 1MB around rs160459).**

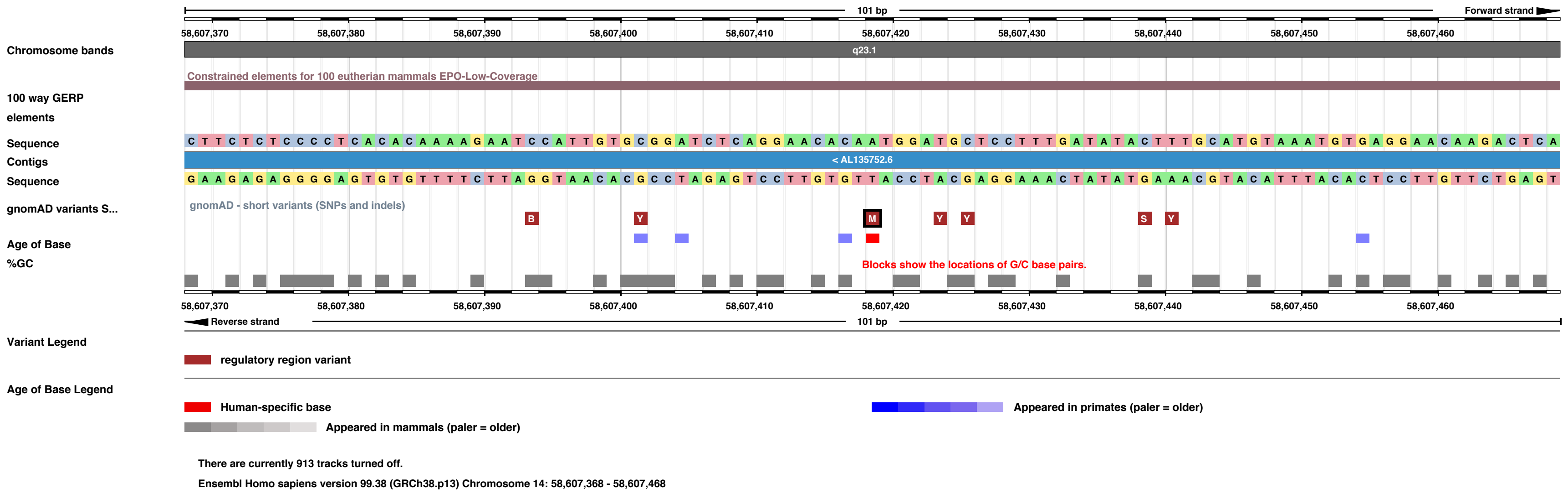


**Fig. S5 Human-specific base annotation in Ensembl for rs160459.** "Timing of the most recent mutation as determined by inter-species whole genome alignments. Each base pair in which the human reference genome differs by substitution from one of its inferred ancestral genomes is colored in either grey (event prior to the primate branch), blue (primate specific), red (human specific (fixed variant) or human specific segregating variant, i.e. SNP). Clicking on a mutation position reveals the sub-tree of species which have inherited the same mutation from their common ancestor. It also reveals a score that represents the age of the mutation in arbitrary units, and determines the intensity of the coloring. The more recent the mutation, the lower the score and the darker the color."

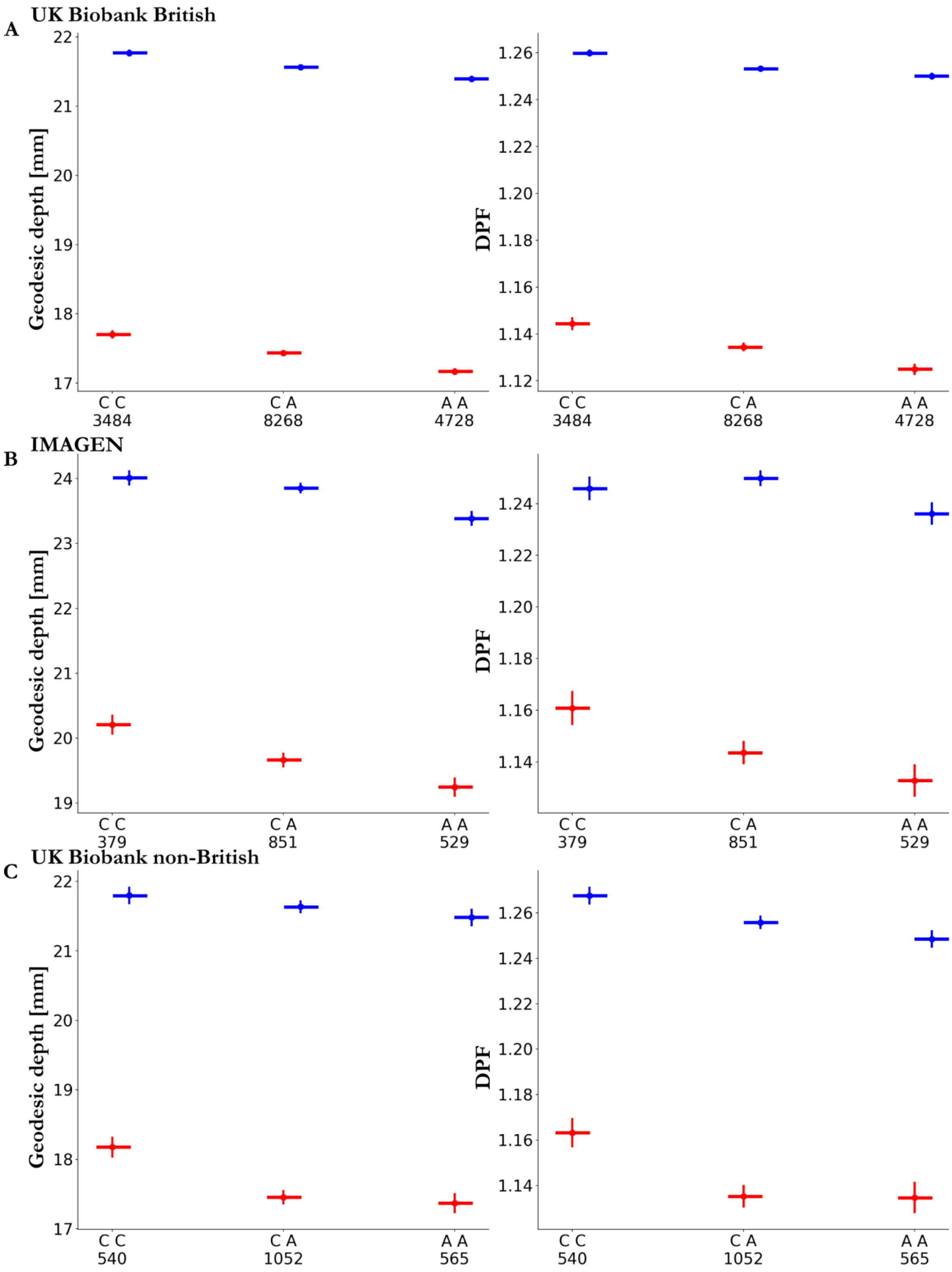


**Fig. S6. Effect sizes of rs160459 configuration on the geodesic depth and DPF in three populations (UKB British (A); IMAGEN (B); UKB non-British (C)).** Left hemisphere values in red and right hemisphere values in blue.

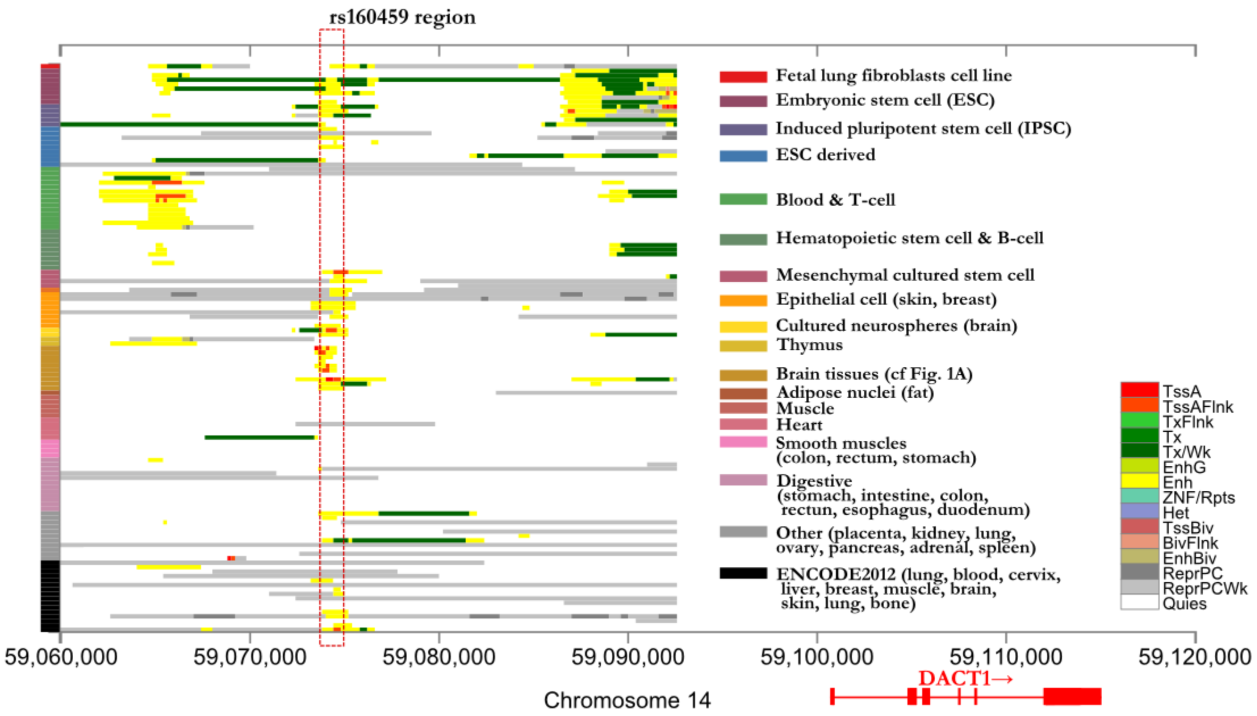


**Fig. S7. Chromatin state annotation of the genomic region around rs160459 (upstream of DACT1) in various tissues.** Red highlighting corresponds to the rs160459 location.

<u>RoadMap Core 15 chromatin state model abbreviations:</u> TssA: Active Transcription Start Site (TSS), TssFlnk: Flanking Active TSS, TxFlnk: Transcription at gene 5' and 3', Tx: Strong transcription, TxWk: Weak transcription, EnhG: Genic Enhancers, Enh: Enhancers, ZNF/Rpts: ZNF genes and repeats, Het: Heterochromatin, TssBiv: Bivalent/Poised TSS, BivFlnk: Flanking Bivalent TSS/Enhancer, EnhBiv: Bivalent Enhancer ReprPC: Repressed PolyComb, ReprPCWk: Weak Repressed PolyComb, Quies: Quiescent/Low.

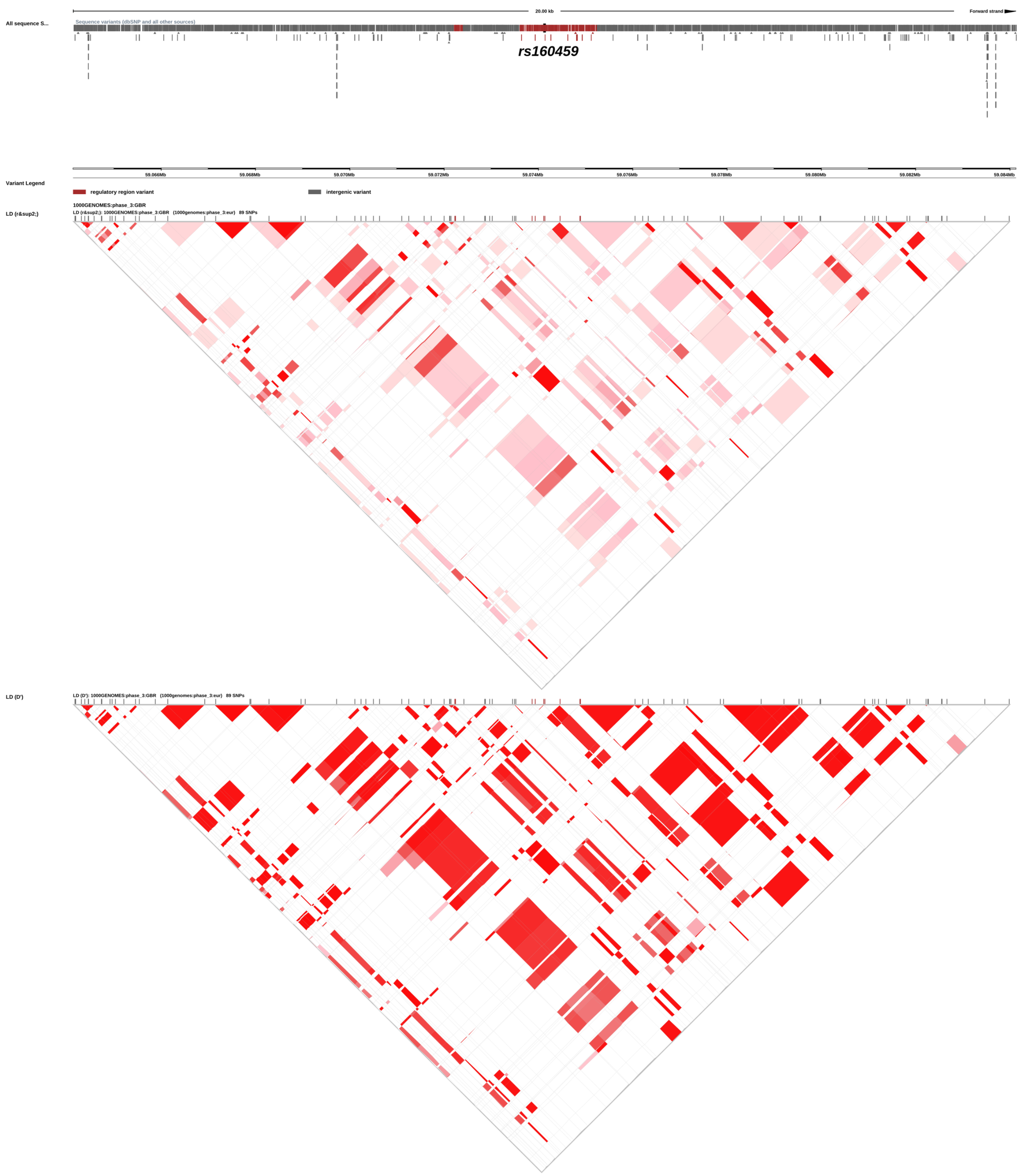


**Fig. S8. LD structure around the regulatory region including rs160459 in the 1000 genomes GBR (British) sample.** The first LD triangle correspond to LD calculated with $r^2$ and the second LD triangle correspond to LD calculated with D'.

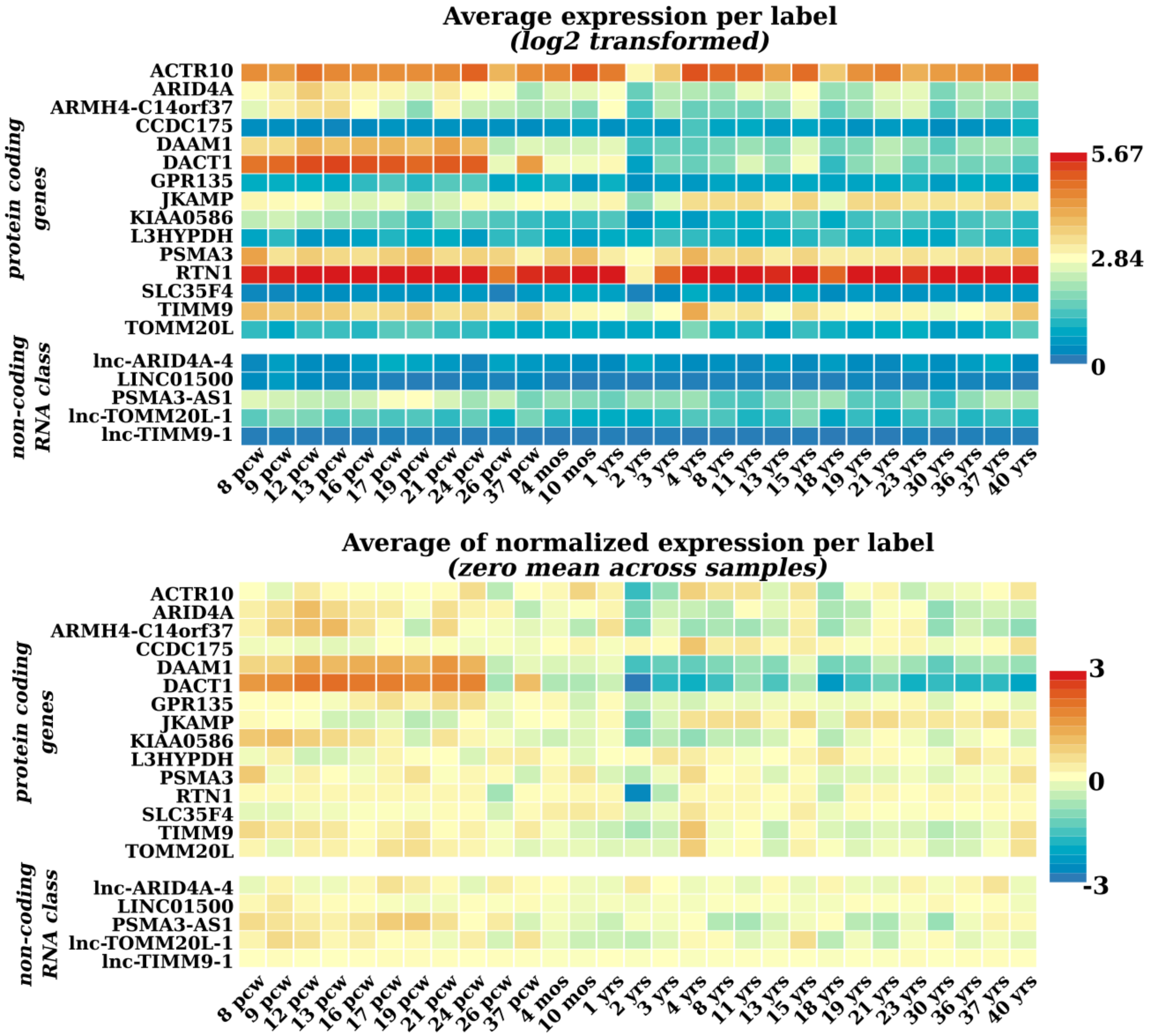


**Fig. S9. Gene expression of the genes within 1 MB of rs160459 across brain developmental stages from the Brainspan, of the genes.**

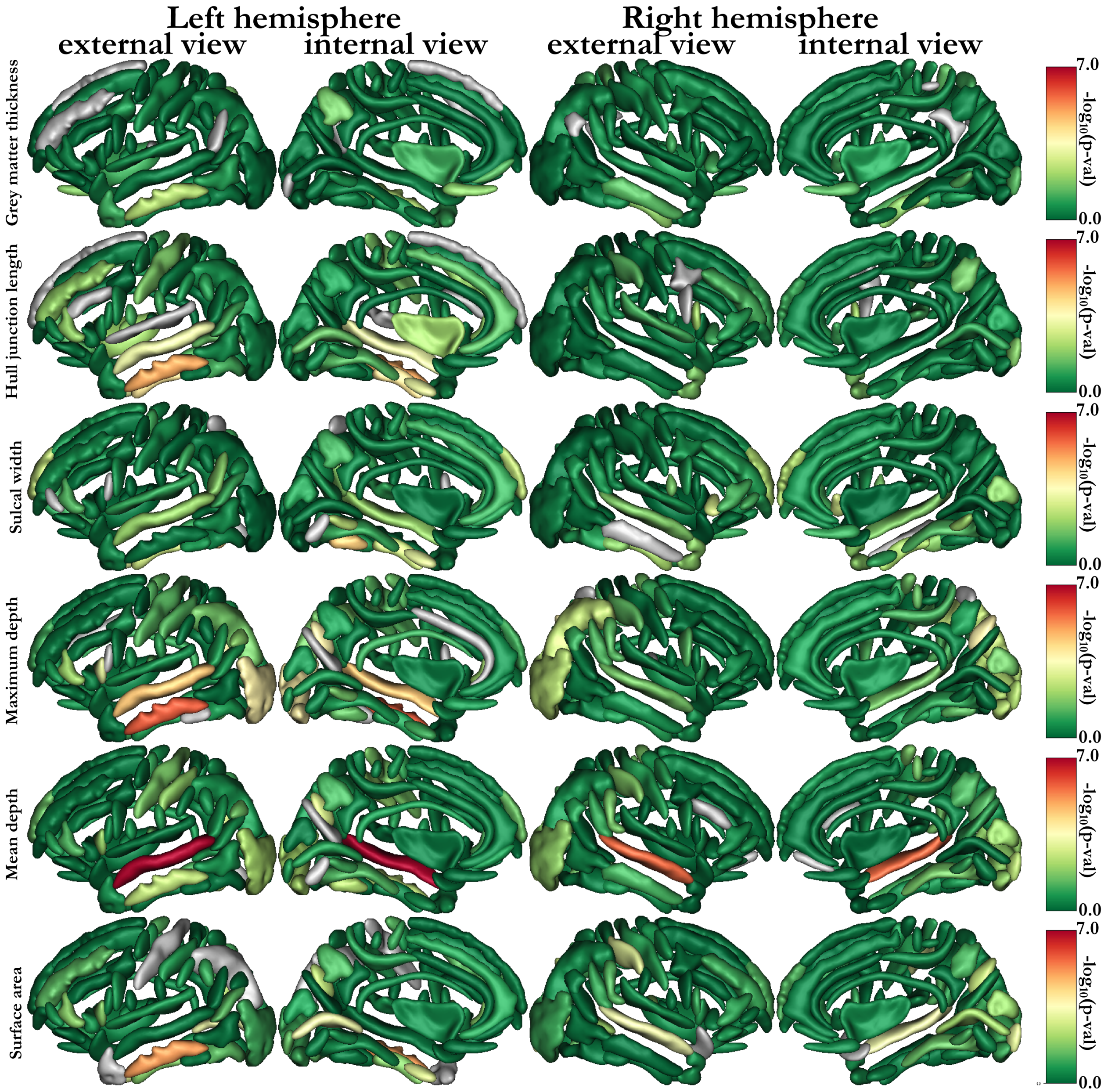


**Fig. S10. Mapping of the log$_{10}$ p-values of the linear association between rs160459 configuration and sulci features (in the following order: gray matter thickness, hull junction length, sulcal width, maximum depth, mean depth, and surface area).** The sulci are displayed using the Statistical Probability Anatomy Map (SPAM) representation, which represents the average sulci shape and position on the reference base of the Brainvisa sulci extraction pipeline (Perrot *et al*. 2011).

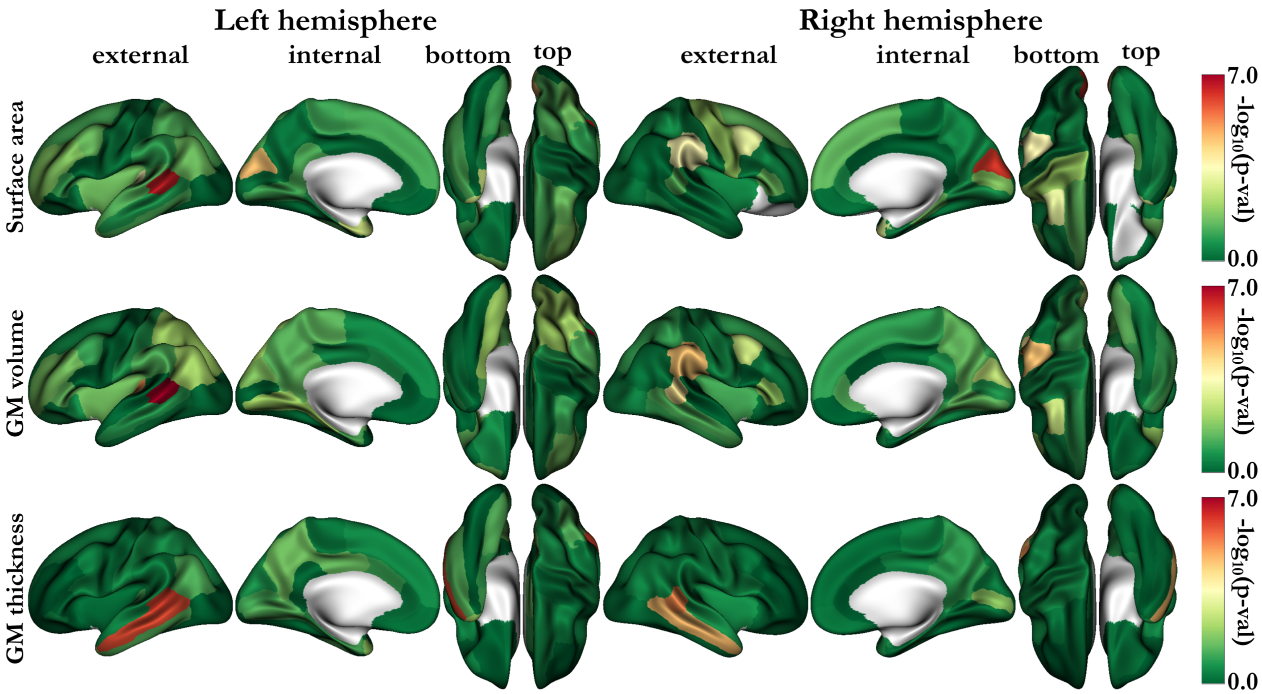


**Fig. S11. Mapping of the $\log_{10}$ p-values of the linear association between rs160459 configuration and Freesurfer features on Desikan-Killiany parcellation (in the following order: surface area, gray matter volume and gray matter thickness).**

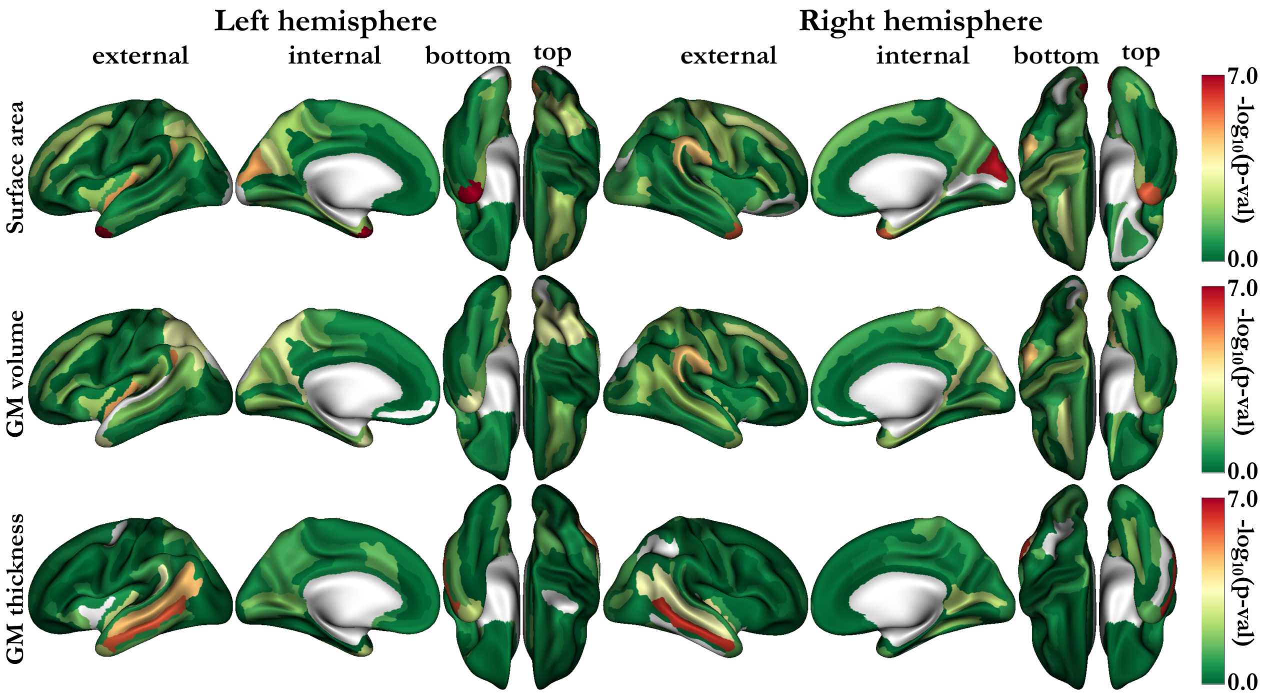


**Fig. S12. Mapping of the $\log_{10}$ p-values of the linear association between rs160459 configuration and Freesurfer features on Destrieux parcellation (in the following order: surface area, gray matter volume and gray matter thickness).** Note that the Destrieux atlas has more parcels than the Desikan atlas. In particular, the gyri and sulci separation is better delineated in the Destrieux atlas.

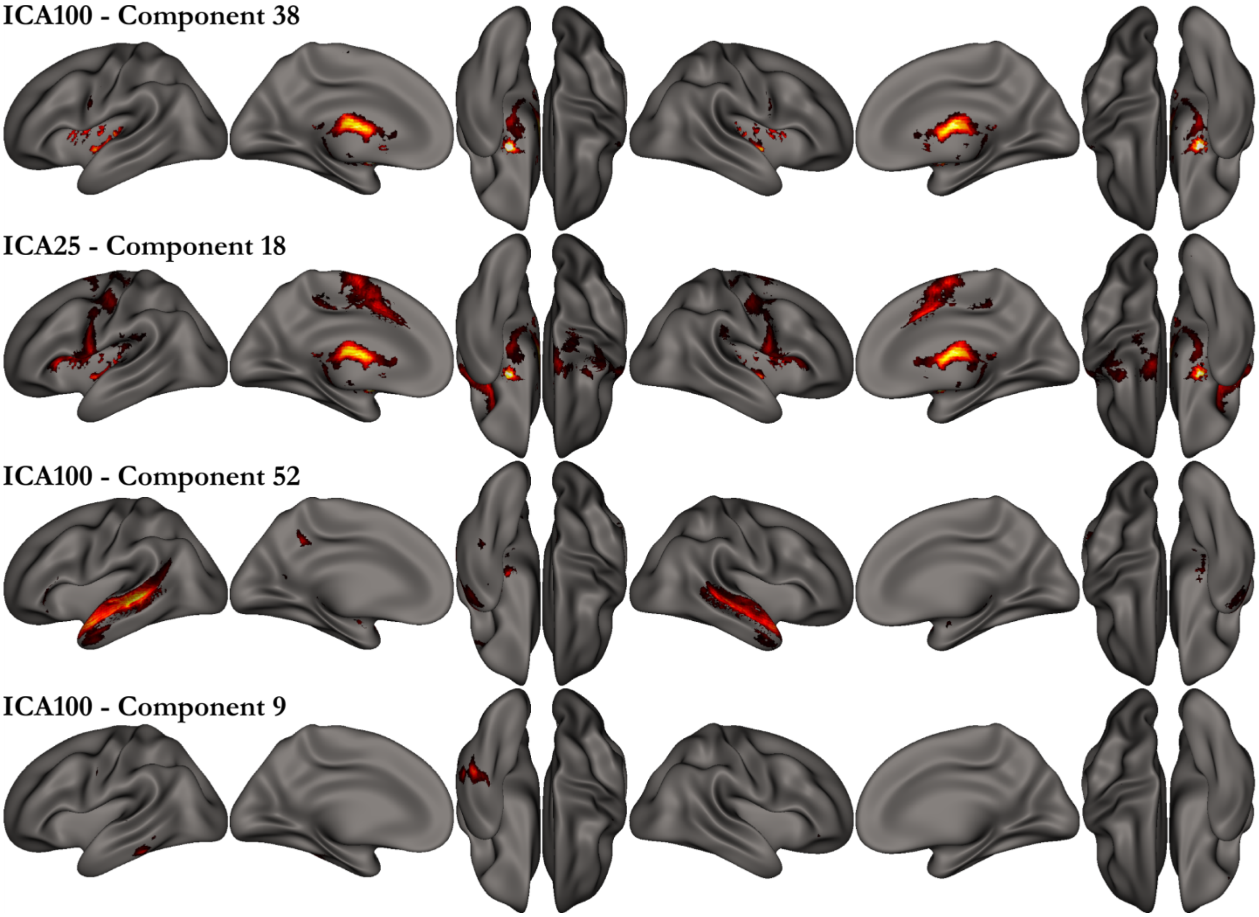


**Fig. S13. Resting-state ICAs with amplitudes most strongly associated with rs160459.** ICA100 – Component 38 (p-val = 0.0021), ICA25 – Component 18 (p-val = 0.0093), ICA100 – Component 52 (p-val = 0.0099), ICA100 – Component 9 (p-val = 0.0182).

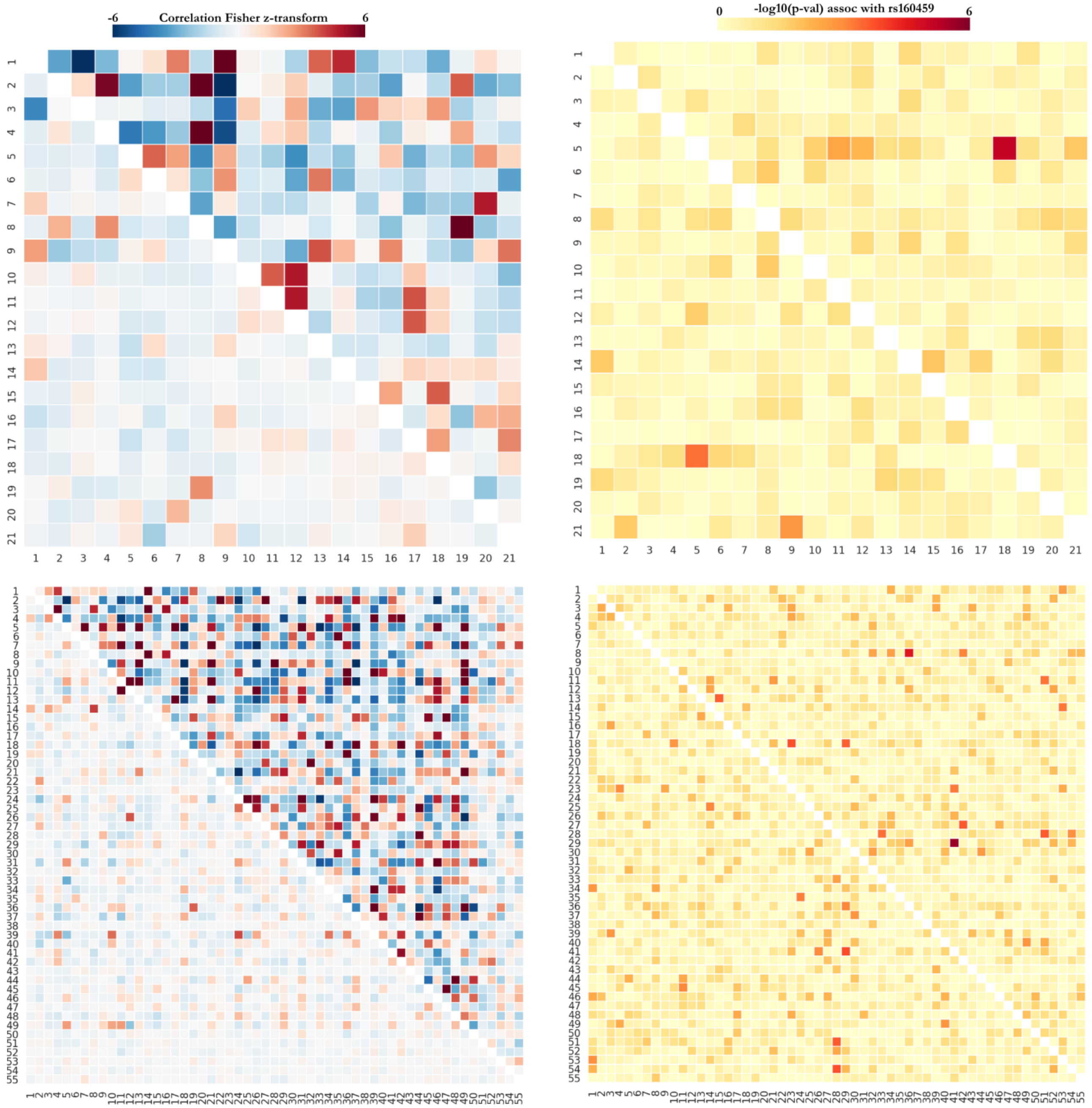


**Fig. S14. Left column: full correlation (above diagonal) and partial correlation (below diagonal) strength; right column: association p-values with rs160459 for ICA with 25 components (first line) and 100 components (second line).** A total of 21 and 55 ICA components that were labeled "good components" by the UK Biobank were retained in the analysis. The significant pairwise correlations are for ICA25 link 5-18 (pval = $7.6{\cdot}10^{-6}$) and for ICA100 link 29-41 (pval = $2.1{\cdot}10^{-6}$).

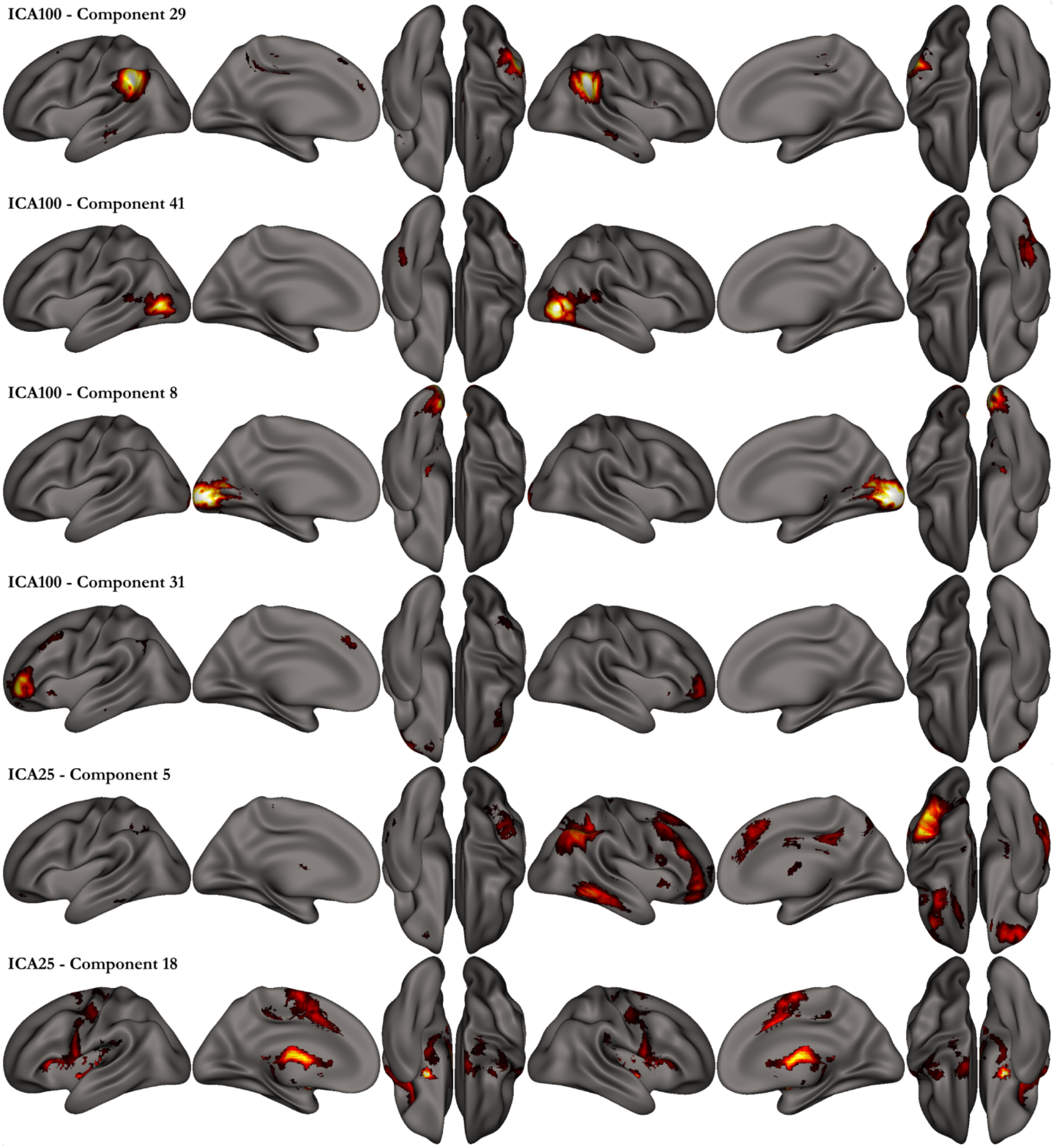


**Fig. S15. The 3 top pairs of ICA for the association of rs160459 with the full correlation among these pairs. First pair: for ICA100 - Good components 29-41 (edge 1146; p-val = $2.1 \cdot 10^{-6}$); Second pair: for ICA100 – Good components 8-31 (edge 385; p-val = $2.5 \cdot 10^{-5}$); Third pair: for ICA25 – Good components 5-18 (edge 87; p-val = $7.6 \cdot 10^{-6}$).**

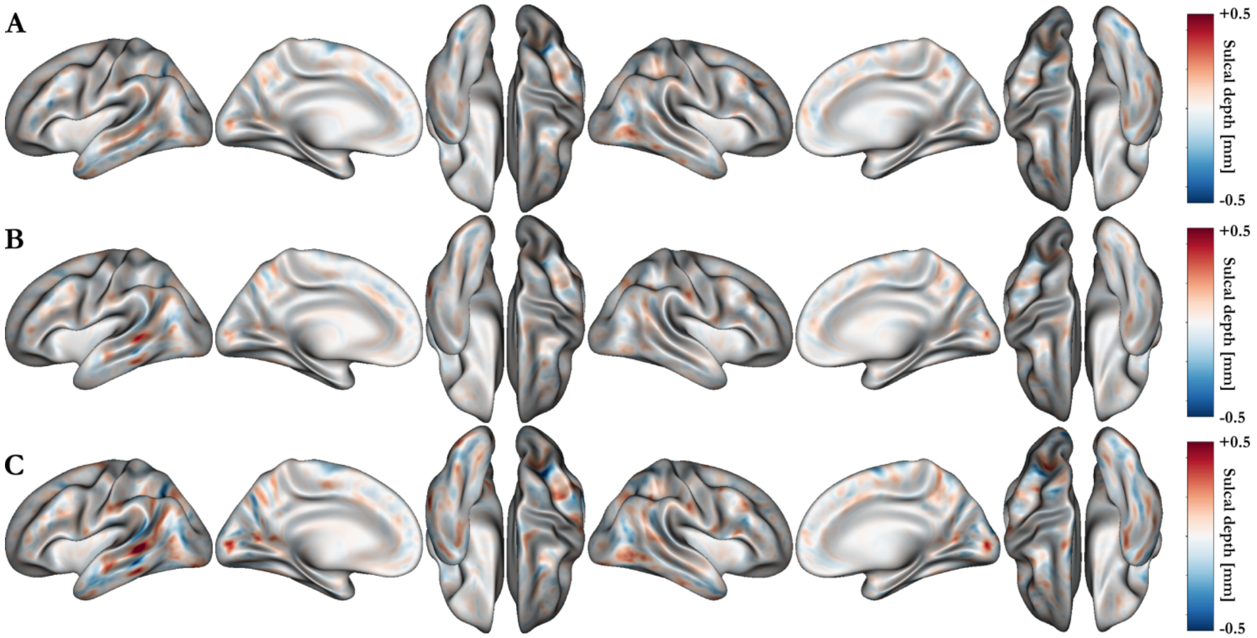


**Fig. S16. Mapping the sulcal depth vertexwise difference between the groups of subjects with different rs160459 allele: 'CC'-'CA' (A), 'CA'-'AA' (B), and 'CC'-AA" (C).**

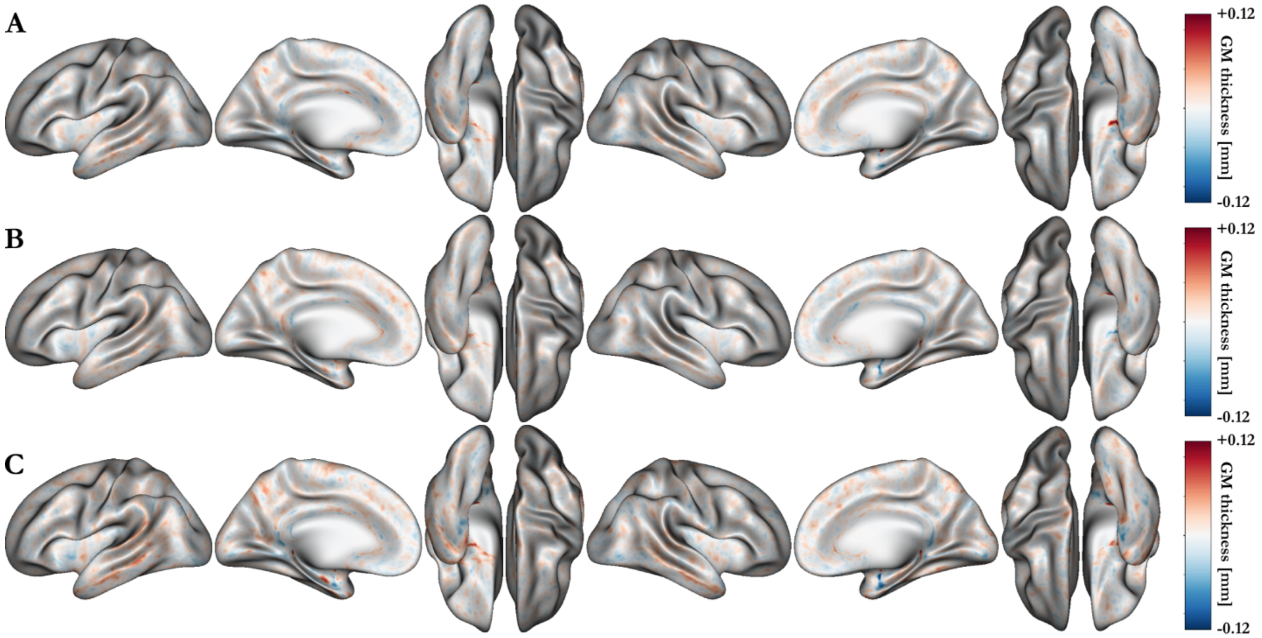


**Fig. S17. Mapping the gray matter thickness vertexwise difference between the groups of subjects with different rs160459 allele: 'CC'-'CA' (A), 'CA'-'AA' (B), and 'CC'-AA" (C).**

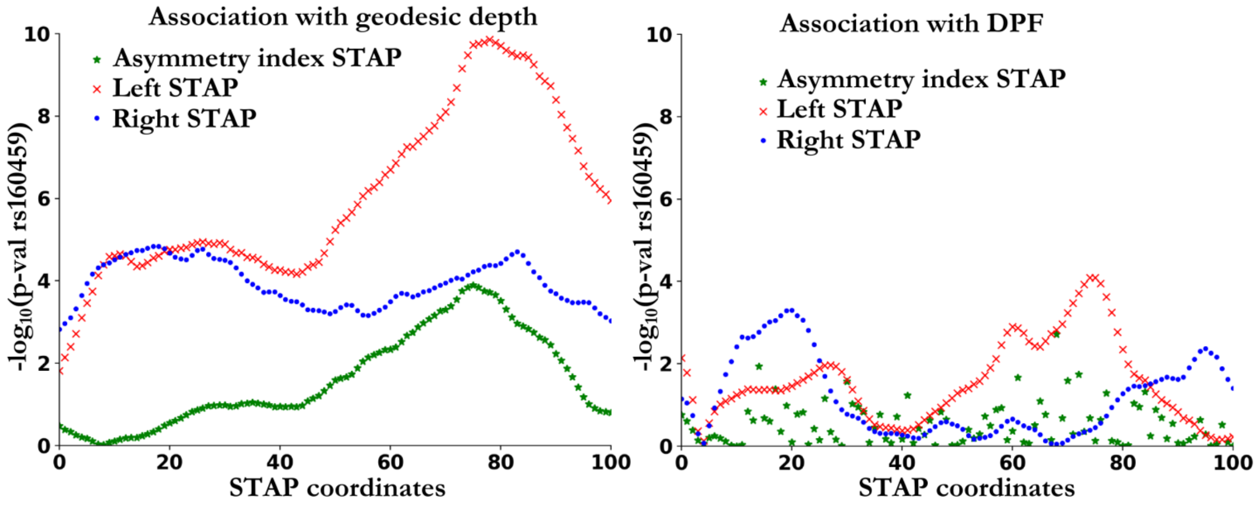


**Fig. S18. rs160459 $-\log_{10}$(p-val) along the STAP for the geodesic depth and DPF.**
