## Supplementary material for "Enhancer locus in ch14q23.1 modulates brain asymmetric temporal regions involved in language processing": Table S6

| SNP | Chr | BP | A1 | A2 | Freq | Probe | Gene | LNCipedia | Orientation | b | SE | p | p_GWAS |
|---|---|---|---|---|---|---|---|---|---|---|---|---|---|
| **rs170239** | 14 | 59064739 | C | T | 0,5468 | ENSG00000032219 | ARID4A | | N | -0,00640914 | 0,0448172 | 0,886285 | 3,55901E-11 |
| **rs170239** | 14 | 59064739 | C | T | 0,5468 | ENSG00000050130 | JKAMP | | N | 0,00812167 | 0,0359745 | 0,821387 | 3,55901E-11 |
| **rs170239** | 14 | 59064739 | C | T | 0,5468 | ENSG00000100567 | PSMA3 | | N | -0,0738344 | 0,0462408 | 0,110324 | 3,55901E-11 |
| **rs170239** | 14 | 59064739 | C | T | 0,5468 | ENSG00000100575 | TIMM9 | | N | -0,0398939 | 0,0428371 | 0,351701 | 3,55901E-11 |
| **rs170239** | 14 | 59064739 | C | T | 0,5468 | ENSG00000100578 | KIAA0586 | | N | 0,0461953 | 0,0440935 | 0,294792 | 3,55901E-11 |
| **rs170239** | 14 | 59064739 | C | T | 0,5468 | ENSG00000100592 | DAAM1 | | N | -0,0500776 | 0,0441854 | 0,257067 | 3,55901E-11 |
| **rs170239** | 14 | 59064739 | C | T | 0,5468 | ENSG00000126790 | L3HYPDH | | N | -0,0185379 | 0,055631 | 0,738961 | 3,55901E-11 |
| **rs170239** | 14 | 59064739 | C | T | 0,5468 | ENSG00000131966 | ACTR10 | | N | 0,153965 | 0,048344 | 0,00144868 | 3,55901E-11 |
| **rs170239** | 14 | 59064739 | C | T | 0,5468 | ENSG00000139971 | C14orf37 | | - | -0,0204219 | 0,0545153 | 0,707952 | 3,55901E-11 |
| **rs170239** | 14 | 59064739 | C | T | 0,5468 | ENSG00000151812 | SLC35F4 | | - | -0,0471178 | 0,053169 | 0,375516 | 3,55901E-11 |
| **rs170239** | 14 | 59064739 | C | T | 0,5468 | ENSG00000151838 | CCDC175 | | - | -0,0878043 | 0,0589691 | 0,13649 | 3,55901E-11 |
| **rs170239** | 14 | 59064739 | C | T | 0,5468 | ENSG00000165617 | DACT1 | | + | -0,105273 | 0,064279 | 0,101475 | 3,55901E-11 |
| **rs170239** | 14 | 59064739 | C | T | 0,5468 | ENSG00000180189 | HMGB1P14 | | + | 0,0394836 | 0,0403102 | 0,327336 | 3,55901E-11 |
| **rs170239** | 14 | 59064739 | C | T | 0,5468 | ENSG00000181619 | GPR135 | | - | -0,0131477 | 0,0591164 | 824 | 3,55901E-11 |
| **rs170239** | 14 | 59064739 | C | T | 0,5468 | ENSG00000196860 | TOMM20L | | + | -0,00592915 | 0,0923838 | 0,948827 | 3,55901E-11 |
| **rs170239** | 14 | 59064739 | C | T | 0,5468 | ENSG00000213598 | RP11-112J1.1 | LINC01500:1 | + | 0,0342914 | 0,0486128 | 0,480562 | 3,55901E-11 |
| **rs170239** | 14 | 59064739 | C | T | 0,5468 | ENSG00000239510 | RPL9P5 | | + | -0,18559 | 0,13876 | 0,181063 | 3,55901E-11 |
| **rs170239** | 14 | 59064739 | C | T | 0,5468 | ENSG00000257621 | RP11-349A22.5 | PSMA3-AS1 | N | 0,117659 | 0,0495156 | 0,0174922 | 3,55901E-11 |
| **rs170239** | 14 | 59064739 | C | T | 0,5468 | ENSG00000258378 | RP11-517O13.1 | lnc-TOMM20L-1:1 | + | -0,0665146 | 0,0493089 | 0,177357 | 3,55901E-11 |
| **rs170239** | 14 | 59064739 | C | T | 0,5468 | ENSG00000258682 | CTD-2002H8.2 | lnc-ARID4A-4:1 | + | 0,0209139 | 0,0436978 | 0,632221 | 3,55901E-11 |
| **rs170239** | 14 | 59064739 | C | T | 0,5468 | ENSG00000258734 | RP11-112J1.3 | | - | 0,0419163 | 0,0375819 | 0,264708 | 3,55901E-11 |
| **rs170239** | 14 | 59064739 | C | T | 0,5468 | ENSG00000258782 | RP11-701B16.2 | lnc-CCDC175-1:1 | - | -0,0810848 | 0,0545294 | 0,137016 | 3,55901E-11 |
| **rs170239** | 14 | 59064739 | C | T | 0,5468 | ENSG00000258856 | CTD-2325K12.1 | | + | -0,170087 | 0,099784 | 0,0882781 | 3,55901E-11 |
| **rs170239** | 14 | 59064739 | C | T | 0,5468 | ENSG00000258900 | HNRNPCP1 | | + | 0,154903 | 0,0478442 | 0,00120516 | 3,55901E-11 |
| **rs170239** | 14 | 59064739 | C | T | 0,5468 | ENSG00000259969 | RP11-999E24.3 | lnc-C14orf37-1:1 | N | -0,125629 | 0,050351 | 0,0125932 | 3,55901E-11 |
| **rs170239** | 14 | 59064739 | C | T | 0,5468 | ENSG00000268466 | AL132989.1 | | + | 0,0662969 | 0,0572221 | 0,246624 | 3,55901E-11 |
| **rs221326** | 14 | 59069053 | G | T | 0,4756 | ENSG00000032219 | ARID4A | | N | 0,0316852 | 0,0445757 | 0,477198 | 2,99726E-10 |
| **rs221326** | 14 | 59069053 | G | T | 0,4756 | ENSG00000050130 | JKAMP | | N | 0,034073 | 0,0358202 | 0,341491 | 2,99726E-10 |
| **rs221326** | 14 | 59069053 | G | T | 0,4756 | ENSG00000100567 | PSMA3 | | N | -0,0750873 | 0,0459716 | 0,102397 | 2,99726E-10 |
| **rs221326** | 14 | 59069053 | G | T | 0,4756 | ENSG00000100575 | TIMM9 | | N | -0,0522788 | 0,0427705 | 0,221591 | 2,99726E-10 |
| **rs221326** | 14 | 59069053 | G | T | 0,4756 | ENSG00000100578 | KIAA0586 | | N | 0,00642562 | 0,04389 | 0,883603 | 2,99726E-10 |
| **rs221326** | 14 | 59069053 | G | T | 0,4756 | ENSG00000100592 | DAAM1 | | N | -0,00705319 | 0,0439957 | 0,872633 | 2,99726E-10 |
| **rs221326** | 14 | 59069053 | G | T | 0,4756 | ENSG00000126790 | L3HYPDH | | N | 0,0189685 | 0,0552779 | 0,731487 | 2,99726E-10 |
| **rs221326** | 14 | 59069053 | G | T | 0,4756 | ENSG00000131966 | ACTR10 | | N | 0,133801 | 0,0481298 | 0,00543582 | 2,99726E-10 |
| **rs221326** | 14 | 59069053 | G | T | 0,4756 | ENSG00000139971 | C14orf37 | | - | 0,0178757 | 0,0544282 | 0,742589 | 2,99726E-10 |
| **rs221326** | 14 | 59069053 | G | T | 0,4756 | ENSG00000151812 | SLC35F4 | | - | -0,110823 | 0,0530676 | 0,0367672 | 2,99726E-10 |
| **rs221326** | 14 | 59069053 | G | T | 0,4756 | ENSG00000151838 | CCDC175 | | - | -0,0492683 | 0,0588965 | 0,402861 | 2,99726E-10 |
| **rs221326** | 14 | 59069053 | G | T | 0,4756 | ENSG00000165617 | DACT1 | | + | -0,164191 | 0,0638674 | 0,0101459 | 2,99726E-10 |
| **rs221326** | 14 | 59069053 | G | T | 0,4756 | ENSG00000180189 | HMGB1P14 | | + | -0,0159032 | 0,0401267 | 0,691866 | 2,99726E-10 |
| **rs221326** | 14 | 59069053 | G | T | 0,4756 | ENSG00000181619 | GPR135 | | - | 0,0137961 | 0,0588047 | 0,814512 | 2,99726E-10 |
| **rs221326** | 14 | 59069053 | G | T | 0,4756 | ENSG00000196860 | TOMM20L | | + | -0,0762862 | 0,0916931 | 0,405424 | 2,99726E-10 |
| **rs221326** | 14 | 59069053 | G | T | 0,4756 | ENSG00000213598 | RP11-112J1.1 | LINC01500:1 | + | 0,0380621 | 0,0483968 | 0,431599 | 2,99726E-10 |
| **rs221326** | 14 | 59069053 | G | T | 0,4756 | ENSG00000239510 | RPL9P5 | | + | -0,141395 | 0,138816 | 0,308403 | 2,99726E-10 |
| **rs221326** | 14 | 59069053 | G | T | 0,4756 | ENSG00000257621 | RP11-349A22.5 | PSMA3-AS1 | N | 0,16473 | 0,0491917 | 0,000811849 | 2,99726E-10 |
| **rs221326** | 14 | 59069053 | G | T | 0,4756 | ENSG00000258378 | RP11-517O13.1 | lnc-TOMM20L-1:1 | + | -0,121982 | 0,0490736 | 0,01293 | 2,99726E-10 |
| **rs221326** | 14 | 59069053 | G | T | 0,4756 | ENSG00000258682 | CTD-2002H8.2 | lnc-ARID4A-4:1 | + | 0,081875 | 0,0434696 | 0,0596325 | 2,99726E-10 |
| **rs221326** | 14 | 59069053 | G | T | 0,4756 | ENSG00000258734 | RP11-112J1.3 | | - | 0,0454626 | 0,0374901 | 0,225261 | 2,99726E-10 |
| **rs221326** | 14 | 59069053 | G | T | 0,4756 | ENSG00000258782 | RP11-701B16.2 | lnc-CCDC175-1:1 | - | -0,0784367 | 0,05418 | 0,1477 | 2,99726E-10 |
| **rs221326** | 14 | 59069053 | G | T | 0,4756 | ENSG00000258856 | CTD-2325K12.1 | | + | -0,113115 | 0,0995062 | 0,255637 | 2,99726E-10 |
| **rs221326** | 14 | 59069053 | G | T | 0,4756 | ENSG00000258900 | HNRNPCP1 | | + | 0,119001 | 0,0478404 | 0,0128658 | 2,99726E-10 |
| **rs221326** | 14 | 59069053 | G | T | 0,4756 | ENSG00000259969 | RP11-999E24.3 | lnc-C14orf37-1:1 | N | -0,136365 | 0,050003 | 0,00638856 | 2,99726E-10 |
| **rs221326** | 14 | 59069053 | G | T | 0,4756 | ENSG00000268466 | AL132989.1 | | + | 0,0349245 | 0,0569152 | 0,539464 | 2,99726E-10 |
| **rs468213** | 14 | 59071100 | C | G | 0,5909 | ENSG00000032219 | ARID4A | | N | 0,0256602 | 0,045435 | 0,572232 | 1,80897E-08 |
| **rs468213** | 14 | 59071100 | C | G | 0,5909 | ENSG00000050130 | JKAMP | | N | 0,0134998 | 0,0364578 | 0,71117 | 1,80897E-08 |
| **rs468213** | 14 | 59071100 | C | G | 0,5909 | ENSG00000100567 | PSMA3 | | N | -0,0694653 | 0,0469493 | 0,138985 | 1,80897E-08 |
| **rs468213** | 14 | 59071100 | C | G | 0,5909 | ENSG00000100575 | TIMM9 | | N | -0,0337824 | 0,0435875 | 0,438311 | 1,80897E-08 |
| **rs468213** | 14 | 59071100 | C | G | 0,5909 | ENSG00000100578 | KIAA0586 | | N | 0,0646989 | 0,0446879 | 0,147674 | 1,80897E-08 |
| **rs468213** | 14 | 59071100 | C | G | 0,5909 | ENSG00000100592 | DAAM1 | | N | -0,0250226 | 0,0448608 | 0,576993 | 1,80897E-08 |
| **rs468213** | 14 | 59071100 | C | G | 0,5909 | ENSG00000126790 | L3HYPDH | | N | -0,00873593 | 0,0564581 | 0,877032 | 1,80897E-08 |

| | | | | | | | | | | | | | |
|---|---|---|---|---|---|---|---|---|---|---|---|---|---|
| **rs468213** | 14 | 59071100 | C | G | 0,5909 | ENSG00000131966 | ACTR10 | | N | 0,176954 | 0,048979 | 0,000302848 | 1,80897E-08 |
| **rs468213** | 14 | 59071100 | C | G | 0,5909 | ENSG00000139970 | RTN1 | | N | 0,0564333 | 0,065639 | 0,389925 | 1,80897E-08 |
| **rs468213** | 14 | 59071100 | C | G | 0,5909 | ENSG00000139971 | C14orf37 | | - | -0,0435227 | 0,0551967 | 0,430403 | 1,80897E-08 |
| **rs468213** | 14 | 59071100 | C | G | 0,5909 | ENSG00000151812 | SLC35F4 | | - | -0,0882983 | 0,0539315 | 0,101582 | 1,80897E-08 |
| **rs468213** | 14 | 59071100 | C | G | 0,5909 | ENSG00000151838 | CCDC175 | | - | -0,117624 | 0,0596133 | 0,0484815 | 1,80897E-08 |
| **rs468213** | 14 | 59071100 | C | G | 0,5909 | ENSG00000165617 | DACT1 | | + | -0,145758 | 0,0650144 | 0,0249654 | 1,80897E-08 |
| **rs468213** | 14 | 59071100 | C | G | 0,5909 | ENSG00000180189 | HMGB1P14 | | + | 0,0219796 | 0,040745 | 0,589582 | 1,80897E-08 |
| **rs468213** | 14 | 59071100 | C | G | 0,5909 | ENSG00000181619 | GPR135 | | - | 0,0111654 | 0,0598369 | 0,851977 | 1,80897E-08 |
| **rs468213** | 14 | 59071100 | C | G | 0,5909 | ENSG00000196860 | TOMM20L | | + | -0,115302 | 0,0931377 | 0,215728 | 1,80897E-08 |
| **rs468213** | 14 | 59071100 | C | G | 0,5909 | ENSG00000213598 | RP11-112J1.1 | LINC01500:1 | + | 0,0367109 | 0,0491597 | 0,455204 | 1,80897E-08 |
| **rs468213** | 14 | 59071100 | C | G | 0,5909 | ENSG00000239510 | RPL9P5 | | + | -0,129408 | 0,141133 | 0,359186 | 1,80897E-08 |
| **rs468213** | 14 | 59071100 | C | G | 0,5909 | ENSG00000257621 | RP11-349A22.5 | PSMA3-AS1 | N | 0,120281 | 0,0503667 | 0,0169356 | 1,80897E-08 |
| **rs468213** | 14 | 59071100 | C | G | 0,5909 | ENSG00000258378 | RP11-517O13.1 | lnc-TOMM20L-1:1 | + | -0,0848888 | 0,0499689 | 0,0893508 | 1,80897E-08 |
| **rs468213** | 14 | 59071100 | C | G | 0,5909 | ENSG00000258682 | CTD-2002H8.2 | lnc-ARID4A-4:1 | + | 0,0607947 | 0,0441512 | 0,168522 | 1,80897E-08 |
| **rs468213** | 14 | 59071100 | C | G | 0,5909 | ENSG00000258734 | RP11-112J1.3 | | - | 0,0545393 | 0,0380244 | 0,15148 | 1,80897E-08 |
| **rs468213** | 14 | 59071100 | C | G | 0,5909 | ENSG00000258782 | RP11-701B16.2 | lnc-CCDC175-1:1 | - | -0,120136 | 0,0549832 | 0,0288923 | 1,80897E-08 |
| **rs468213** | 14 | 59071100 | C | G | 0,5909 | ENSG00000258856 | CTD-2325K12.1 | | + | -0,125081 | 0,101608 | 0,218317 | 1,80897E-08 |
| **rs468213** | 14 | 59071100 | C | G | 0,5909 | ENSG00000258900 | HNRNPCP1 | | + | 0,144358 | 0,0484686 | 0,00289778 | 1,80897E-08 |
| **rs468213** | 14 | 59071100 | C | G | 0,5909 | ENSG00000259969 | RP11-999E24.3 | lnc-C14orf37-1:1 | N | -0,137006 | 0,0511071 | 0,0073455 | 1,80897E-08 |
| **rs468213** | 14 | 59071100 | C | G | 0,5909 | ENSG00000268466 | AL132989.1 | | + | 0,0800713 | 0,0577113 | 0,165306 | 1,80897E-08 |
| **rs149142** | 14 | 59071725 | T | C | 0,58 | ENSG00000032219 | ARID4A | | N | 0,0250602 | 0,0453439 | 0,580489 | 1,17005E-08 |
| **rs149142** | 14 | 59071725 | T | C | 0,58 | ENSG00000050130 | JKAMP | | N | 0,0140222 | 0,0363615 | 0,699768 | 1,17005E-08 |
| **rs149142** | 14 | 59071725 | T | C | 0,58 | ENSG00000100567 | PSMA3 | | N | -0,0698402 | 0,0468682 | 0,136187 | 1,17005E-08 |
| **rs149142** | 14 | 59071725 | T | C | 0,58 | ENSG00000100575 | TIMM9 | | N | -0,0400889 | 0,0434851 | 0,356582 | 1,17005E-08 |
| **rs149142** | 14 | 59071725 | T | C | 0,58 | ENSG00000100578 | KIAA0586 | | N | 0,0581301 | 0,0445868 | 0,192319 | 1,17005E-08 |
| **rs149142** | 14 | 59071725 | T | C | 0,58 | ENSG00000100592 | DAAM1 | | N | -0,0260112 | 0,0447678 | 0,561224 | 1,17005E-08 |
| **rs149142** | 14 | 59071725 | T | C | 0,58 | ENSG00000126790 | L3HYPDH | | N | -0,00618979 | 0,0563951 | 0,912602 | 1,17005E-08 |
| **rs149142** | 14 | 59071725 | T | C | 0,58 | ENSG00000131966 | ACTR10 | | N | 0,178226 | 0,0488826 | 0,000266349 | 1,17005E-08 |
| **rs149142** | 14 | 59071725 | T | C | 0,58 | ENSG00000139970 | RTN1 | | N | 0,0564333 | 0,065639 | 0,389925 | 1,17005E-08 |
| **rs149142** | 14 | 59071725 | T | C | 0,58 | ENSG00000139971 | C14orf37 | | - | -0,0388911 | 0,0549842 | 0,479371 | 1,17005E-08 |
| **rs149142** | 14 | 59071725 | T | C | 0,58 | ENSG00000151812 | SLC35F4 | | - | -0,084385 | 0,0537311 | 0,116298 | 1,17005E-08 |
| **rs149142** | 14 | 59071725 | T | C | 0,58 | ENSG00000151838 | CCDC175 | | - | -0,11071 | 0,0593946 | 0,0623255 | 1,17005E-08 |
| **rs149142** | 14 | 59071725 | T | C | 0,58 | ENSG00000165617 | DACT1 | | + | -0,156257 | 0,0646756 | 0,0156914 | 1,17005E-08 |
| **rs149142** | 14 | 59071725 | T | C | 0,58 | ENSG00000180189 | HMGB1P14 | | + | 0,0228863 | 0,0405841 | 0,572806 | 1,17005E-08 |
| **rs149142** | 14 | 59071725 | T | C | 0,58 | ENSG00000181619 | GPR135 | | - | 0,011893 | 0,0596119 | 0,841866 | 1,17005E-08 |
| **rs149142** | 14 | 59071725 | T | C | 0,58 | ENSG00000196860 | TOMM20L | | + | -0,116035 | 0,0927816 | 0,211072 | 1,17005E-08 |
| **rs149142** | 14 | 59071725 | T | C | 0,58 | ENSG00000213598 | RP11-112J1.1 | LINC01500:1 | + | 0,0405008 | 0,0489532 | 0,408046 | 1,17005E-08 |
| **rs149142** | 14 | 59071725 | T | C | 0,58 | ENSG00000239510 | RPL9P5 | | + | -0,139679 | 0,140493 | 0,320122 | 1,17005E-08 |
| **rs149142** | 14 | 59071725 | T | C | 0,58 | ENSG00000257621 | RP11-349A22.5 | PSMA3-AS1 | N | 0,126228 | 0,0502773 | 0,0120514 | 1,17005E-08 |
| **rs149142** | 14 | 59071725 | T | C | 0,58 | ENSG00000258378 | RP11-517O13.1 | lnc-TOMM20L-1:1 | + | -0,0854445 | 0,0497821 | 0,0860943 | 1,17005E-08 |
| **rs149142** | 14 | 59071725 | T | C | 0,58 | ENSG00000258682 | CTD-2002H8.2 | lnc-ARID4A-4:1 | + | 0,0636563 | 0,0439839 | 0,147824 | 1,17005E-08 |
| **rs149142** | 14 | 59071725 | T | C | 0,58 | ENSG00000258734 | RP11-112J1.3 | | - | 0,0571624 | 0,037872 | 0,131207 | 1,17005E-08 |
| **rs149142** | 14 | 59071725 | T | C | 0,58 | ENSG00000258782 | RP11-701B16.2 | lnc-CCDC175-1:1 | - | -0,117069 | 0,054757 | 0,0325182 | 1,17005E-08 |
| **rs149142** | 14 | 59071725 | T | C | 0,58 | ENSG00000258856 | CTD-2325K12.1 | | + | -0,131179 | 0,101189 | 0,194846 | 1,17005E-08 |
| **rs149142** | 14 | 59071725 | T | C | 0,58 | ENSG00000258900 | HNRNPCP1 | | + | 0,145851 | 0,0482886 | 0,00252432 | 1,17005E-08 |
| **rs149142** | 14 | 59071725 | T | C | 0,58 | ENSG00000259969 | RP11-999E24.3 | lnc-C14orf37-1:1 | N | -0,137637 | 0,0510352 | 0,00699865 | 1,17005E-08 |
| **rs149142** | 14 | 59071725 | T | C | 0,58 | ENSG00000268466 | AL132989.1 | | + | 0,0766534 | 0,0574922 | 0,182438 | 1,17005E-08 |
| **rs1078243** | 14 | 59072144 | T | C | 0,5033 | ENSG00000032219 | ARID4A | | N | 0,0189992 | 0,0447124 | 0,670896 | 6,80381E-09 |
| **rs1078243** | 14 | 59072144 | T | C | 0,5033 | ENSG00000050130 | JKAMP | | N | 0,0183337 | 0,0358348 | 0,608918 | 6,80381E-09 |
| **rs1078243** | 14 | 59072144 | T | C | 0,5033 | ENSG00000100567 | PSMA3 | | N | -0,0863118 | 0,0461129 | 0,0612411 | 6,80381E-09 |
| **rs1078243** | 14 | 59072144 | T | C | 0,5033 | ENSG00000100575 | TIMM9 | | N | -0,00466435 | 0,0429112 | 0,913442 | 6,80381E-09 |
| **rs1078243** | 14 | 59072144 | T | C | 0,5033 | ENSG00000100578 | KIAA0586 | | N | 0,0536251 | 0,0439169 | 0,222064 | 6,80381E-09 |
| **rs1078243** | 14 | 59072144 | T | C | 0,5033 | ENSG00000100592 | DAAM1 | | N | -0,0124748 | 0,0441278 | 0,777408 | 6,80381E-09 |
| **rs1078243** | 14 | 59072144 | T | C | 0,5033 | ENSG00000126790 | L3HYPDH | | N | 0,00824834 | 0,0555306 | 0,881919 | 6,80381E-09 |
| **rs1078243** | 14 | 59072144 | T | C | 0,5033 | ENSG00000131966 | ACTR10 | | N | 0,161141 | 0,0481657 | 0,000821207 | 6,80381E-09 |
| **rs1078243** | 14 | 59072144 | T | C | 0,5033 | ENSG00000139970 | RTN1 | | N | 0,0167667 | 0,0646181 | 0,79527 | 6,80381E-09 |
| **rs1078243** | 14 | 59072144 | T | C | 0,5033 | ENSG00000139971 | C14orf37 | | - | -0,013633 | 0,0542721 | 0,801662 | 6,80381E-09 |
| **rs1078243** | 14 | 59072144 | T | C | 0,5033 | ENSG00000151812 | SLC35F4 | | - | -0,105284 | 0,052996 | 0,0469628 | 6,80381E-09 |
| **rs1078243** | 14 | 59072144 | T | C | 0,5033 | ENSG00000151838 | CCDC175 | | - | -0,0812281 | 0,058735 | 0,166678 | 6,80381E-09 |
| **rs1078243** | 14 | 59072144 | T | C | 0,5033 | ENSG00000165617 | DACT1 | | + | -0,11934 | 0,0638422 | 0,0615825 | 6,80381E-09 |
| **rs1078243** | 14 | 59072144 | T | C | 0,5033 | ENSG00000180189 | HMGB1P14 | | + | -0,00657084 | 0,0401264 | 0,869925 | 6,80381E-09 |
| **rs1078243** | 14 | 59072144 | T | C | 0,5033 | ENSG00000181619 | GPR135 | | - | -0,00106021 | 0,0587348 | 0,985598 | 6,80381E-09 |

| | | | | | | | | | | | | | |
|---|---|---|---|---|---|---|---|---|---|---|---|---|---|
| **rs1078243** | 14 | 59072144 | T | C | 0,5033 | ENSG00000196860 | TOMM20L | | + | -0,0999924 | 0,0916231 | 0,275121 | 6,80381E-09 |
| **rs1078243** | 14 | 59072144 | T | C | 0,5033 | ENSG00000213598 | RP11-112J1.1 | LINC01500:1 | + | 0,0315183 | 0,0483203 | 0,514222 | 6,80381E-09 |
| **rs1078243** | 14 | 59072144 | T | C | 0,5033 | ENSG00000239510 | RPL9P5 | | + | -0,0386924 | 0,139298 | 0,781191 | 6,80381E-09 |
| **rs1078243** | 14 | 59072144 | T | C | 0,5033 | ENSG00000257621 | RP11-349A22.5 | PSMA3-AS1 | N | 0,196067 | 0,0493354 | 7,06293E-05 | 6,80381E-09 |
| **rs1078243** | 14 | 59072144 | T | C | 0,5033 | ENSG00000258378 | RP11-517O13.1 | lnc-TOMM20L-1:1 | + | -0,0863101 | 0,049109 | 0,0788291 | 6,80381E-09 |
| **rs1078243** | 14 | 59072144 | T | C | 0,5033 | ENSG00000258682 | CTD-2002H8.2 | lnc-ARID4A-4:1 | + | 0,085379 | 0,0434348 | 0,0493352 | 6,80381E-09 |
| **rs1078243** | 14 | 59072144 | T | C | 0,5033 | ENSG00000258734 | RP11-112J1.3 | | - | 0,0260092 | 0,0374026 | 0,486814 | 6,80381E-09 |
| **rs1078243** | 14 | 59072144 | T | C | 0,5033 | ENSG00000258782 | RP11-701B16.2 | lnc-CCDC175-1:1 | - | -0,0931839 | 0,0540301 | 0,0845875 | 6,80381E-09 |
| **rs1078243** | 14 | 59072144 | T | C | 0,5033 | ENSG00000258856 | CTD-2325K12.1 | | + | -0,160328 | 0,0991433 | 0,10585 | 6,80381E-09 |
| **rs1078243** | 14 | 59072144 | T | C | 0,5033 | ENSG00000258900 | HNRNPCP1 | | + | 0,163583 | 0,0475827 | 0,000586297 | 6,80381E-09 |
| **rs1078243** | 14 | 59072144 | T | C | 0,5033 | ENSG00000259969 | RP11-999E24.3 | lnc-C14orf37-1:1 | N | -0,151169 | 0,0502032 | 0,00260264 | 6,80381E-09 |
| **rs1078243** | 14 | 59072144 | T | C | 0,5033 | ENSG00000268466 | AL132989.1 | | + | 0,0684695 | 0,0566243 | 0,22659 | 6,80381E-09 |
| **rs186347** | 14 | 59072226 | G | T | 0,4755 | ENSG00000032219 | ARID4A | | N | 0,00672344 | 0,0445338 | 0,879996 | 2,92564E-12 |
| **rs186347** | 14 | 59072226 | G | T | 0,4755 | ENSG00000050130 | JKAMP | | N | 0,0362228 | 0,0357736 | 0,311272 | 2,92564E-12 |
| **rs186347** | 14 | 59072226 | G | T | 0,4755 | ENSG00000100567 | PSMA3 | | N | -0,0816456 | 0,0459188 | 0,0753969 | 2,92564E-12 |
| **rs186347** | 14 | 59072226 | G | T | 0,4755 | ENSG00000100575 | TIMM9 | | N | -0,0217117 | 0,0427504 | 0,611543 | 2,92564E-12 |
| **rs186347** | 14 | 59072226 | G | T | 0,4755 | ENSG00000100578 | KIAA0586 | | N | 0,0452678 | 0,0437903 | 0,301258 | 2,92564E-12 |
| **rs186347** | 14 | 59072226 | G | T | 0,4755 | ENSG00000100592 | DAAM1 | | N | -0,0181543 | 0,0439497 | 0,679555 | 2,92564E-12 |
| **rs186347** | 14 | 59072226 | G | T | 0,4755 | ENSG00000126790 | L3HYPDH | | N | 0,0221811 | 0,0551688 | 0,68764 | 2,92564E-12 |
| **rs186347** | 14 | 59072226 | G | T | 0,4755 | ENSG00000131966 | ACTR10 | | N | 0,155862 | 0,0479149 | 0,00114236 | 2,92564E-12 |
| **rs186347** | 14 | 59072226 | G | T | 0,4755 | ENSG00000139970 | RTN1 | | N | 0,0266682 | 0,0639319 | 0,67658 | 2,92564E-12 |
| **rs186347** | 14 | 59072226 | G | T | 0,4755 | ENSG00000139971 | C14orf37 | | - | 0,00483894 | 0,0543919 | 0,92911 | 2,92564E-12 |
| **rs186347** | 14 | 59072226 | G | T | 0,4755 | ENSG00000151812 | SLC35F4 | | - | -0,0989095 | 0,0531275 | 0,0626399 | 2,92564E-12 |
| **rs186347** | 14 | 59072226 | G | T | 0,4755 | ENSG00000151838 | CCDC175 | | - | -0,097617 | 0,0587804 | 0,0967724 | 2,92564E-12 |
| **rs186347** | 14 | 59072226 | G | T | 0,4755 | ENSG00000165617 | DACT1 | | + | -0,128501 | 0,0639815 | 0,0445998 | 2,92564E-12 |
| **rs186347** | 14 | 59072226 | G | T | 0,4755 | ENSG00000180189 | HMGB1P14 | | + | -0,010101 | 0,0401667 | 0,801444 | 2,92564E-12 |
| **rs186347** | 14 | 59072226 | G | T | 0,4755 | ENSG00000181619 | GPR135 | | - | 0,0101412 | 0,0587799 | 0,863022 | 2,92564E-12 |
| **rs186347** | 14 | 59072226 | G | T | 0,4755 | ENSG00000196860 | TOMM20L | | + | -0,0383379 | 0,0920664 | 0,677106 | 2,92564E-12 |
| **rs186347** | 14 | 59072226 | G | T | 0,4755 | ENSG00000213598 | RP11-112J1.1 | LINC01500:1 | + | 0,0528952 | 0,0483769 | 0,274219 | 2,92564E-12 |
| **rs186347** | 14 | 59072226 | G | T | 0,4755 | ENSG00000239510 | RPL9P5 | | + | -0,110726 | 0,139087 | 0,425976 | 2,92564E-12 |
| **rs186347** | 14 | 59072226 | G | T | 0,4755 | ENSG00000257621 | RP11-349A22.5 | PSMA3-AS1 | N | 0,186884 | 0,0490713 | 0,000139854 | 2,92564E-12 |
| **rs186347** | 14 | 59072226 | G | T | 0,4755 | ENSG00000258378 | RP11-517O13.1 | lnc-TOMM20L-1:1 | + | -0,104808 | 0,0490867 | 0,0327484 | 2,92564E-12 |
| **rs186347** | 14 | 59072226 | G | T | 0,4755 | ENSG00000258682 | CTD-2002H8.2 | lnc-ARID4A-4:1 | + | 0,0639887 | 0,04356 | 0,141838 | 2,92564E-12 |
| **rs186347** | 14 | 59072226 | G | T | 0,4755 | ENSG00000258734 | RP11-112J1.3 | | - | 0,0280534 | 0,0374913 | 0,454301 | 2,92564E-12 |
| **rs186347** | 14 | 59072226 | G | T | 0,4755 | ENSG00000258782 | RP11-701B16.2 | lnc-CCDC175-1:1 | - | -0,115029 | 0,0541872 | 0,0337704 | 2,92564E-12 |
| **rs186347** | 14 | 59072226 | G | T | 0,4755 | ENSG00000258856 | CTD-2325K12.1 | | + | -0,169104 | 0,0992508 | 0,0884178 | 2,92564E-12 |
| **rs186347** | 14 | 59072226 | G | T | 0,4755 | ENSG00000258900 | HNRNPCP1 | | + | 0,139237 | 0,047723 | 0,00352722 | 2,92564E-12 |
| **rs186347** | 14 | 59072226 | G | T | 0,4755 | ENSG00000259969 | RP11-999E24.3 | lnc-C14orf37-1:1 | N | -0,138867 | 0,0499393 | 0,00542406 | 2,92564E-12 |
| **rs186347** | 14 | 59072226 | G | T | 0,4755 | ENSG00000268466 | AL132989.1 | | + | 0,0487258 | 0,056842 | 0,391326 | 2,92564E-12 |
| **rs160460** | 14 | 59072964 | A | G | 0,5248 | ENSG00000032219 | ARID4A | | N | 0,00212127 | 0,0447622 | 0,962203 | 1,20294E-11 |
| **rs160460** | 14 | 59072964 | A | G | 0,5248 | ENSG00000050130 | JKAMP | | N | 0,0351439 | 0,0359127 | 0,327782 | 1,20294E-11 |
| **rs160460** | 14 | 59072964 | A | G | 0,5248 | ENSG00000100567 | PSMA3 | | N | -0,0642888 | 0,0462041 | 0,164102 | 1,20294E-11 |
| **rs160460** | 14 | 59072964 | A | G | 0,5248 | ENSG00000100575 | TIMM9 | | N | -0,0441571 | 0,042888 | 0,303203 | 1,20294E-11 |
| **rs160460** | 14 | 59072964 | A | G | 0,5248 | ENSG00000100578 | KIAA0586 | | N | 0,0324594 | 0,0440184 | 0,460877 | 1,20294E-11 |
| **rs160460** | 14 | 59072964 | A | G | 0,5248 | ENSG00000100592 | DAAM1 | | N | -0,0356627 | 0,0441465 | 0,419191 | 1,20294E-11 |
| **rs160460** | 14 | 59072964 | A | G | 0,5248 | ENSG00000126790 | L3HYPDH | | N | 0,0110095 | 0,0556498 | 0,843174 | 1,20294E-11 |
| **rs160460** | 14 | 59072964 | A | G | 0,5248 | ENSG00000131966 | ACTR10 | | N | 0,1746 | 0,0481774 | 0,000289961 | 1,20294E-11 |
| **rs160460** | 14 | 59072964 | A | G | 0,5248 | ENSG00000139970 | RTN1 | | N | 0,0609594 | 0,0645366 | 0,344878 | 1,20294E-11 |
| **rs160460** | 14 | 59072964 | A | G | 0,5248 | ENSG00000139971 | C14orf37 | | - | -0,00984853 | 0,0543809 | 0,856287 | 1,20294E-11 |
| **rs160460** | 14 | 59072964 | A | G | 0,5248 | ENSG00000151812 | SLC35F4 | | - | -0,0838457 | 0,0531252 | 0,114504 | 1,20294E-11 |
| **rs160460** | 14 | 59072964 | A | G | 0,5248 | ENSG00000151838 | CCDC175 | | - | -0,111642 | 0,058594 | 0,0567337 | 1,20294E-11 |
| **rs160460** | 14 | 59072964 | A | G | 0,5248 | ENSG00000165617 | DACT1 | | + | -0,16295 | 0,0638918 | 0,0107597 | 1,20294E-11 |
| **rs160460** | 14 | 59072964 | A | G | 0,5248 | ENSG00000180189 | HMGB1P14 | | + | 0,0138066 | 0,0401503 | 0,730943 | 1,20294E-11 |
| **rs160460** | 14 | 59072964 | A | G | 0,5248 | ENSG00000181619 | GPR135 | | - | -0,0108634 | 0,0588976 | 0,853663 | 1,20294E-11 |
| **rs160460** | 14 | 59072964 | A | G | 0,5248 | ENSG00000196860 | TOMM20L | | + | -0,0289047 | 0,092147 | 0,753764 | 1,20294E-11 |
| **rs160460** | 14 | 59072964 | A | G | 0,5248 | ENSG00000213598 | RP11-112J1.1 | LINC01500:1 | + | 0,0487886 | 0,0484445 | 0,313885 | 1,20294E-11 |
| **rs160460** | 14 | 59072964 | A | G | 0,5248 | ENSG00000239510 | RPL9P5 | | + | -0,144003 | 0,138795 | 0,299491 | 1,20294E-11 |
| **rs160460** | 14 | 59072964 | A | G | 0,5248 | ENSG00000257621 | RP11-349A22.5 | PSMA3-AS1 | N | 0,117223 | 0,049571 | 0,0180425 | 1,20294E-11 |
| **rs160460** | 14 | 59072964 | A | G | 0,5248 | ENSG00000258378 | RP11-517O13.1 | lnc-TOMM20L-1:1 | + | -0,116444 | 0,0491008 | 0,0177145 | 1,20294E-11 |
| **rs160460** | 14 | 59072964 | A | G | 0,5248 | ENSG00000258682 | CTD-2002H8.2 | lnc-ARID4A-4:1 | + | 0,0468564 | 0,0435053 | 0,281467 | 1,20294E-11 |
| **rs160460** | 14 | 59072964 | A | G | 0,5248 | ENSG00000258734 | RP11-112J1.3 | | - | 0,0400078 | 0,0374749 | 0,285706 | 1,20294E-11 |
| **rs160460** | 14 | 59072964 | A | G | 0,5248 | ENSG00000258782 | RP11-701B16.2 | lnc-CCDC175-1:1 | - | -0,118061 | 0,0541727 | 0,0293059 | 1,20294E-11 |

| | | | | | | | | | | | | | |
|---|---|---|---|---|---|---|---|---|---|---|---|---|---|
| rs160460 | 14 | 59072964 | A | G | 0,5248 | ENSG00000258856 | CTD-2325K12.1 | | + | -0,150189 | 0,0997677 | 0,132225 | 1,20294E-11 |
| rs160460 | 14 | 59072964 | A | G | 0,5248 | ENSG00000258900 | HNRNPCP1 | | + | 0,116107 | 0,0478867 | 0,0153244 | 1,20294E-11 |
| rs160460 | 14 | 59072964 | A | G | 0,5248 | ENSG00000259969 | RP11-999E24.3 | lnc-C14orf37-1:1 | N | -0,12631 | 0,0502981 | 0,0120313 | 1,20294E-11 |
| rs160460 | 14 | 59072964 | A | G | 0,5248 | ENSG00000268466 | AL132989.1 | | + | 0,0893099 | 0,0568676 | 0,116301 | 1,20294E-11 |
| rs160459 | 14 | 59074136 | A | C | 0,4744 | ENSG00000032219 | ARID4A | | N | 0,0067203 | 0,0445352 | 0,880056 | 4,10338E-12 |
| rs160459 | 14 | 59074136 | A | C | 0,4744 | ENSG00000050130 | JKAMP | | N | 0,0362288 | 0,0357756 | 0,31122 | 4,10338E-12 |
| rs160459 | 14 | 59074136 | A | C | 0,4744 | ENSG00000100567 | PSMA3 | | N | -0,0816511 | 0,04592 | 0,0753847 | 4,10338E-12 |
| rs160459 | 14 | 59074136 | A | C | 0,4744 | ENSG00000100575 | TIMM9 | | N | -0,0217138 | 0,042752 | 0,611523 | 4,10338E-12 |
| rs160459 | 14 | 59074136 | A | C | 0,4744 | ENSG00000100578 | KIAA0586 | | N | 0,0452757 | 0,0437918 | 0,30119 | 4,10338E-12 |
| rs160459 | 14 | 59074136 | A | C | 0,4744 | ENSG00000100592 | DAAM1 | | N | -0,0181558 | 0,0439511 | 0,67954 | 4,10338E-12 |
| rs160459 | 14 | 59074136 | A | C | 0,4744 | ENSG00000126790 | L3HYPDH | | N | 0,022179 | 0,0551683 | 0,687666 | 4,10338E-12 |
| rs160459 | 14 | 59074136 | A | C | 0,4744 | ENSG00000131966 | ACTR10 | | N | 0,155857 | 0,0479158 | 0,00114303 | 4,10338E-12 |
| rs160459 | 14 | 59074136 | A | C | 0,4744 | ENSG00000139970 | RTN1 | | N | 0,026667 | 0,0639289 | 0,67658 | 4,10338E-12 |
| rs160459 | 14 | 59074136 | A | C | 0,4744 | ENSG00000139971 | C14orf37 | | - | 0,00483946 | 0,0543976 | 0,92911 | 4,10338E-12 |
| rs160459 | 14 | 59074136 | A | C | 0,4744 | ENSG00000151812 | SLC35F4 | | - | -0,09892 | 0,0531331 | 0,0626399 | 4,10338E-12 |
| rs160459 | 14 | 59074136 | A | C | 0,4744 | ENSG00000151838 | CCDC175 | | - | -0,0976274 | 0,0587867 | 0,0967724 | 4,10338E-12 |
| rs160459 | 14 | 59074136 | A | C | 0,4744 | ENSG00000165617 | DACT1 | | + | -0,128515 | 0,0639883 | 0,0445998 | 4,10338E-12 |
| rs160459 | 14 | 59074136 | A | C | 0,4744 | ENSG00000180189 | HMGB1P14 | | + | -0,0101021 | 0,0401709 | 0,801444 | 4,10338E-12 |
| rs160459 | 14 | 59074136 | A | C | 0,4744 | ENSG00000181619 | GPR135 | | - | 0,0101423 | 0,0587862 | 0,863022 | 4,10338E-12 |
| rs160459 | 14 | 59074136 | A | C | 0,4744 | ENSG00000196860 | TOMM20L | | + | -0,0383419 | 0,0920762 | 0,677106 | 4,10338E-12 |
| rs160459 | 14 | 59074136 | A | C | 0,4744 | ENSG00000213598 | RP11-112J1.1 | LINC01500:1 | + | 0,0529008 | 0,048382 | 0,274219 | 4,10338E-12 |
| rs160459 | 14 | 59074136 | A | C | 0,4744 | ENSG00000239510 | RPL9P5 | | + | -0,110738 | 0,139101 | 0,425976 | 4,10338E-12 |
| rs160459 | 14 | 59074136 | A | C | 0,4744 | ENSG00000257621 | RP11-349A22.5 | PSMA3-AS1 | N | 0,186884 | 0,049072 | 0,000139886 | 4,10338E-12 |
| rs160459 | 14 | 59074136 | A | C | 0,4744 | ENSG00000258378 | RP11-517O13.1 | lnc-TOMM20L-1:1 | + | -0,104819 | 0,0490919 | 0,0327484 | 4,10338E-12 |
| rs160459 | 14 | 59074136 | A | C | 0,4744 | ENSG00000258682 | CTD-2002H8.2 | lnc-ARID4A-4:1 | + | 0,0639954 | 0,0435646 | 0,141839 | 4,10338E-12 |
| rs160459 | 14 | 59074136 | A | C | 0,4744 | ENSG00000258734 | RP11-112J1.3 | | - | 0,0280564 | 0,0374953 | 0,454301 | 4,10338E-12 |
| rs160459 | 14 | 59074136 | A | C | 0,4744 | ENSG00000258782 | RP11-701B16.2 | lnc-CCDC175-1:1 | - | -0,115041 | 0,0541929 | 0,0337704 | 4,10338E-12 |
| rs160459 | 14 | 59074136 | A | C | 0,4744 | ENSG00000258856 | CTD-2325K12.1 | | + | -0,169122 | 0,0992613 | 0,0884178 | 4,10338E-12 |
| rs160459 | 14 | 59074136 | A | C | 0,4744 | ENSG00000258900 | HNRNPCP1 | | + | 0,139252 | 0,047728 | 0,00352722 | 4,10338E-12 |
| rs160459 | 14 | 59074136 | A | C | 0,4744 | ENSG00000259969 | RP11-999E24.3 | lnc-C14orf37-1:1 | N | -0,13887 | 0,0499399 | 0,00542345 | 4,10338E-12 |
| rs160459 | 14 | 59074136 | A | C | 0,4744 | ENSG00000268466 | AL132989.1 | | + | 0,048731 | 0,056848 | 0,391326 | 4,10338E-12 |
| rs160458 | 14 | 59074878 | T | C | 0,4533 | ENSG00000032219 | ARID4A | | N | -0,00355893 | 0,0445014 | 0,936258 | 2,03377E-12 |
| rs160458 | 14 | 59074878 | T | C | 0,4533 | ENSG00000050130 | JKAMP | | N | 0,0202841 | 0,0358413 | 0,571433 | 2,03377E-12 |
| rs160458 | 14 | 59074878 | T | C | 0,4533 | ENSG00000100567 | PSMA3 | | N | -0,0995349 | 0,0458999 | 0,0301189 | 2,03377E-12 |
| rs160458 | 14 | 59074878 | T | C | 0,4533 | ENSG00000100575 | TIMM9 | | N | -0,0256097 | 0,0427718 | 0,549338 | 2,03377E-12 |
| rs160458 | 14 | 59074878 | T | C | 0,4533 | ENSG00000100578 | KIAA0586 | | N | 0,0268637 | 0,0437783 | 0,539459 | 2,03377E-12 |
| rs160458 | 14 | 59074878 | T | C | 0,4533 | ENSG00000100592 | DAAM1 | | N | 0,00332144 | 0,0439624 | 0,939776 | 2,03377E-12 |
| rs160458 | 14 | 59074878 | T | C | 0,4533 | ENSG00000126790 | L3HYPDH | | N | 0,0231642 | 0,0551242 | 0,674326 | 2,03377E-12 |
| rs160458 | 14 | 59074878 | T | C | 0,4533 | ENSG00000131966 | ACTR10 | | N | 0,14927 | 0,0479568 | 0,00185449 | 2,03377E-12 |
| rs160458 | 14 | 59074878 | T | C | 0,4533 | ENSG00000139970 | RTN1 | | N | 0,0536853 | 0,0637323 | 0,399589 | 2,03377E-12 |
| rs160458 | 14 | 59074878 | T | C | 0,4533 | ENSG00000139971 | C14orf37 | | - | -0,0315459 | 0,0544486 | 0,562339 | 2,03377E-12 |
| rs160458 | 14 | 59074878 | T | C | 0,4533 | ENSG00000151812 | SLC35F4 | | - | -0,0924254 | 0,0533605 | 0,0832567 | 2,03377E-12 |
| rs160458 | 14 | 59074878 | T | C | 0,4533 | ENSG00000151838 | CCDC175 | | - | -0,16599 | 0,0586695 | 0,00466592 | 2,03377E-12 |
| rs160458 | 14 | 59074878 | T | C | 0,4533 | ENSG00000165617 | DACT1 | | + | -0,169142 | 0,0640199 | 0,00824133 | 2,03377E-12 |
| rs160458 | 14 | 59074878 | T | C | 0,4533 | ENSG00000180189 | HMGB1P14 | | + | 0,0285749 | 0,0402333 | 0,477561 | 2,03377E-12 |
| rs160458 | 14 | 59074878 | T | C | 0,4533 | ENSG00000181619 | GPR135 | | - | 0,0215898 | 0,0589679 | 0,71427 | 2,03377E-12 |
| rs160458 | 14 | 59074878 | T | C | 0,4533 | ENSG00000196860 | TOMM20L | | + | -0,0615064 | 0,0922625 | 0,504998 | 2,03377E-12 |
| rs160458 | 14 | 59074878 | T | C | 0,4533 | ENSG00000213598 | RP11-112J1.1 | LINC01500:1 | + | 0,0111486 | 0,0486047 | 0,818579 | 2,03377E-12 |
| rs160458 | 14 | 59074878 | T | C | 0,4533 | ENSG00000239510 | RPL9P5 | | + | -0,0815334 | 0,139726 | 0,559541 | 2,03377E-12 |
| rs160458 | 14 | 59074878 | T | C | 0,4533 | ENSG00000257621 | RP11-349A22.5 | PSMA3-AS1 | N | 0,166842 | 0,0491797 | 0,000692604 | 2,03377E-12 |
| rs160458 | 14 | 59074878 | T | C | 0,4533 | ENSG00000258378 | RP11-517O13.1 | lnc-TOMM20L-1:1 | + | -0,117087 | 0,0492789 | 0,0175015 | 2,03377E-12 |
| rs160458 | 14 | 59074878 | T | C | 0,4533 | ENSG00000258682 | CTD-2002H8.2 | lnc-ARID4A-4:1 | + | 0,0645894 | 0,0436172 | 0,138653 | 2,03377E-12 |
| rs160458 | 14 | 59074878 | T | C | 0,4533 | ENSG00000258734 | RP11-112J1.3 | | - | 0,0219317 | 0,0375943 | 0,559639 | 2,03377E-12 |
| rs160458 | 14 | 59074878 | T | C | 0,4533 | ENSG00000258782 | RP11-701B16.2 | lnc-CCDC175-1:1 | - | -0,167859 | 0,0541689 | 0,00194303 | 2,03377E-12 |
| rs160458 | 14 | 59074878 | T | C | 0,4533 | ENSG00000258856 | CTD-2325K12.1 | | + | -0,161375 | 0,0999138 | 0,10628 | 2,03377E-12 |
| rs160458 | 14 | 59074878 | T | C | 0,4533 | ENSG00000258900 | HNRNPCP1 | | + | 0,0772638 | 0,0480552 | 0,107876 | 2,03377E-12 |
| rs160458 | 14 | 59074878 | T | C | 0,4533 | ENSG00000259969 | RP11-999E24.3 | lnc-C14orf37-1:1 | N | -0,148256 | 0,0498954 | 0,00296504 | 2,03377E-12 |
| rs160458 | 14 | 59074878 | T | C | 0,4533 | ENSG00000268466 | AL132989.1 | | + | 0,0480896 | 0,057085 | 0,399552 | 2,03377E-12 |
| rs4898962 | 14 | 59079971 | T | C | 0,5111 | ENSG00000032219 | ARID4A | | N | -0,0167579 | 0,0444202 | 0,705981 | 4,64432E-08 |
| rs4898962 | 14 | 59079971 | T | C | 0,5111 | ENSG00000050130 | JKAMP | | N | -0,00822001 | 0,0357126 | 0,817959 | 4,64432E-08 |
| rs4898962 | 14 | 59079971 | T | C | 0,5111 | ENSG00000100567 | PSMA3 | | N | -0,0452952 | 0,0459204 | 0,323944 | 4,64432E-08 |
| rs4898962 | 14 | 59079971 | T | C | 0,5111 | ENSG00000100575 | TIMM9 | | N | -0,0256977 | 0,0427191 | 0,547473 | 4,64432E-08 |

| | | | | | | | | | | | | | |
|---|---|---|---|---|---|---|---|---|---|---|---|---|---|
| **rs4898962** | 14 | 59079971 | T | C | 0,5111 | ENSG00000100578 | KIAA0586 | | N | 0,0194346 | 0,0436632 | 0,656246 | 4,64432E-08 |
| **rs4898962** | 14 | 59079971 | T | C | 0,5111 | ENSG00000100592 | DAAM1 | | N | -0,0188459 | 0,0438635 | 0,667452 | 4,64432E-08 |
| **rs4898962** | 14 | 59079971 | T | C | 0,5111 | ENSG00000126790 | L3HYPDH | | N | 0,0177913 | 0,0550739 | 0,746661 | 4,64432E-08 |
| **rs4898962** | 14 | 59079971 | T | C | 0,5111 | ENSG00000131966 | ACTR10 | | N | 0,0965077 | 0,0481408 | 0,0449954 | 4,64432E-08 |
| **rs4898962** | 14 | 59079971 | T | C | 0,5111 | ENSG00000139970 | RTN1 | | N | 0,141272 | 0,0634802 | 0,0260515 | 4,64432E-08 |
| **rs4898962** | 14 | 59079971 | T | C | 0,5111 | ENSG00000139971 | C14orf37 | | - | 0,0165298 | 0,0542265 | 0,760497 | 4,64432E-08 |
| **rs4898962** | 14 | 59079971 | T | C | 0,5111 | ENSG00000151812 | SLC35F4 | | - | -0,0424772 | 0,0531769 | 0,424412 | 4,64432E-08 |
| **rs4898962** | 14 | 59079971 | T | C | 0,5111 | ENSG00000151838 | CCDC175 | | - | -0,130828 | 0,0584724 | 0,0252577 | 4,64432E-08 |
| **rs4898962** | 14 | 59079971 | T | C | 0,5111 | ENSG00000165617 | DACT1 | | + | -0,165769 | 0,0636448 | 0,00919848 | 4,64432E-08 |
| **rs4898962** | 14 | 59079971 | T | C | 0,5111 | ENSG00000180189 | HMGB1P14 | | + | 0,0916555 | 0,0401407 | 0,0224094 | 4,64432E-08 |
| **rs4898962** | 14 | 59079971 | T | C | 0,5111 | ENSG00000181619 | GPR135 | | - | 0,0376574 | 0,0586402 | 0,520758 | 4,64432E-08 |
| **rs4898962** | 14 | 59079971 | T | C | 0,5111 | ENSG00000196860 | TOMM20L | | + | 0,0201064 | 0,0917765 | 0,826588 | 4,64432E-08 |
| **rs4898962** | 14 | 59079971 | T | C | 0,5111 | ENSG00000213598 | RP11-112J1.1 | LINC01500:1 | + | -0,0242609 | 0,0483231 | 0,615628 | 4,64432E-08 |
| **rs4898962** | 14 | 59079971 | T | C | 0,5111 | ENSG00000239510 | RPL9P5 | | + | -0,110345 | 0,138956 | 0,427137 | 4,64432E-08 |
| **rs4898962** | 14 | 59079971 | T | C | 0,5111 | ENSG00000257621 | RP11-349A22.5 | PSMA3-AS1 | N | 0,108391 | 0,0492736 | 0,0278227 | 4,64432E-08 |
| **rs4898962** | 14 | 59079971 | T | C | 0,5111 | ENSG00000258378 | RP11-517O13.1 | lnc-TOMM20L-1:1 | + | -0,0999712 | 0,0491526 | 0,0419621 | 4,64432E-08 |
| **rs4898962** | 14 | 59079971 | T | C | 0,5111 | ENSG00000258682 | CTD-2002H8.2 | lnc-ARID4A-4:1 | + | 0,0030588 | 0,0434824 | 0,943919 | 4,64432E-08 |
| **rs4898962** | 14 | 59079971 | T | C | 0,5111 | ENSG00000258734 | RP11-112J1.3 | | - | 0,0261259 | 0,0372486 | 0,483059 | 4,64432E-08 |
| **rs4898962** | 14 | 59079971 | T | C | 0,5111 | ENSG00000258782 | RP11-701B16.2 | lnc-CCDC175-1:1 | - | -0,0903251 | 0,0541718 | 0,0954385 | 4,64432E-08 |
| **rs4898962** | 14 | 59079971 | T | C | 0,5111 | ENSG00000258856 | CTD-2325K12.1 | | + | -0,125264 | 0,0999804 | 0,210247 | 4,64432E-08 |
| **rs4898962** | 14 | 59079971 | T | C | 0,5111 | ENSG00000258900 | HNRNPCP1 | | + | 0,0558953 | 0,0479254 | 0,243493 | 4,64432E-08 |
| **rs4898962** | 14 | 59079971 | T | C | 0,5111 | ENSG00000259969 | RP11-999E24.3 | lnc-C14orf37-1:1 | N | -0,100365 | 0,0500097 | 0,0447595 | 4,64432E-08 |
| **rs4898962** | 14 | 59079971 | T | C | 0,5111 | ENSG00000268466 | AL132989.1 | | + | 0,0327778 | 0,0570388 | 0,565523 | 4,64432E-08 |
| **rs311814** | 14 | 59085726 | C | T | 0,5245 | ENSG00000032219 | ARID4A | | N | -0,0148015 | 0,0444112 | 0,738921 | 2,39181E-08 |
| **rs311814** | 14 | 59085726 | C | T | 0,5245 | ENSG00000050130 | JKAMP | | N | 0,00388336 | 0,0357464 | 0,913491 | 2,39181E-08 |
| **rs311814** | 14 | 59085726 | C | T | 0,5245 | ENSG00000100567 | PSMA3 | | N | -0,0436971 | 0,0459676 | 0,341805 | 2,39181E-08 |
| **rs311814** | 14 | 59085726 | C | T | 0,5245 | ENSG00000100575 | TIMM9 | | N | -0,0348595 | 0,042733 | 0,414641 | 2,39181E-08 |
| **rs311814** | 14 | 59085726 | C | T | 0,5245 | ENSG00000100578 | KIAA0586 | | N | 0,0333593 | 0,0437519 | 0,445782 | 2,39181E-08 |
| **rs311814** | 14 | 59085726 | C | T | 0,5245 | ENSG00000100592 | DAAM1 | | N | -0,0133633 | 0,0439168 | 0,76091 | 2,39181E-08 |
| **rs311814** | 14 | 59085726 | C | T | 0,5245 | ENSG00000126790 | L3HYPDH | | N | 0,00924386 | 0,0551129 | 0,866799 | 2,39181E-08 |
| **rs311814** | 14 | 59085726 | C | T | 0,5245 | ENSG00000131966 | ACTR10 | | N | 0,09965 | 0,0481571 | 0,038521 | 2,39181E-08 |
| **rs311814** | 14 | 59085726 | C | T | 0,5245 | ENSG00000139970 | RTN1 | | N | 0,141272 | 0,0634802 | 0,0260515 | 2,39181E-08 |
| **rs311814** | 14 | 59085726 | C | T | 0,5245 | ENSG00000139971 | C14orf37 | | - | -0,0259379 | 0,0543263 | 0,633044 | 2,39181E-08 |
| **rs311814** | 14 | 59085726 | C | T | 0,5245 | ENSG00000151812 | SLC35F4 | | - | -0,0561234 | 0,0532264 | 0,291687 | 2,39181E-08 |
| **rs311814** | 14 | 59085726 | C | T | 0,5245 | ENSG00000151838 | CCDC175 | | - | -0,145683 | 0,0585275 | 0,0128056 | 2,39181E-08 |
| **rs311814** | 14 | 59085726 | C | T | 0,5245 | ENSG00000165617 | DACT1 | | + | -0,166109 | 0,0637603 | 0,00918162 | 2,39181E-08 |
| **rs311814** | 14 | 59085726 | C | T | 0,5245 | ENSG00000180189 | HMGB1P14 | | + | 0,0672558 | 0,0402223 | 0,0945039 | 2,39181E-08 |
| **rs311814** | 14 | 59085726 | C | T | 0,5245 | ENSG00000181619 | GPR135 | | - | 0,0455509 | 0,0586995 | 0,437748 | 2,39181E-08 |
| **rs311814** | 14 | 59085726 | C | T | 0,5245 | ENSG00000196860 | TOMM20L | | + | 0,0264927 | 0,0921243 | 0,773671 | 2,39181E-08 |
| **rs311814** | 14 | 59085726 | C | T | 0,5245 | ENSG00000213598 | RP11-112J1.1 | LINC01500:1 | + | -0,0153018 | 0,0483793 | 0,751784 | 2,39181E-08 |
| **rs311814** | 14 | 59085726 | C | T | 0,5245 | ENSG00000239510 | RPL9P5 | | + | -0,128767 | 0,138936 | 0,354026 | 2,39181E-08 |
| **rs311814** | 14 | 59085726 | C | T | 0,5245 | ENSG00000257621 | RP11-349A22.5 | PSMA3-AS1 | N | 0,117998 | 0,0492467 | 0,0165724 | 2,39181E-08 |
| **rs311814** | 14 | 59085726 | C | T | 0,5245 | ENSG00000258378 | RP11-517O13.1 | lnc-TOMM20L-1:1 | + | -0,105946 | 0,0492194 | 0,031356 | 2,39181E-08 |
| **rs311814** | 14 | 59085726 | C | T | 0,5245 | ENSG00000258682 | CTD-2002H8.2 | lnc-ARID4A-4:1 | + | 0,00285351 | 0,0435271 | 0,94773 | 2,39181E-08 |
| **rs311814** | 14 | 59085726 | C | T | 0,5245 | ENSG00000258734 | RP11-112J1.3 | | - | 0,0212145 | 0,0372914 | 0,569435 | 2,39181E-08 |
| **rs311814** | 14 | 59085726 | C | T | 0,5245 | ENSG00000258782 | RP11-701B16.2 | lnc-CCDC175-1:1 | - | -0,0966005 | 0,0542222 | 0,0748201 | 2,39181E-08 |
| **rs311814** | 14 | 59085726 | C | T | 0,5245 | ENSG00000258856 | CTD-2325K12.1 | | + | -0,131664 | 0,100064 | 0,188239 | 2,39181E-08 |
| **rs311814** | 14 | 59085726 | C | T | 0,5245 | ENSG00000258900 | HNRNPCP1 | | + | 0,0386979 | 0,0479205 | 0,419353 | 2,39181E-08 |
| **rs311814** | 14 | 59085726 | C | T | 0,5245 | ENSG00000259969 | RP11-999E24.3 | lnc-C14orf37-1:1 | N | -0,112131 | 0,0500087 | 0,024946 | 2,39181E-08 |
| **rs311814** | 14 | 59085726 | C | T | 0,5245 | ENSG00000268466 | AL132989.1 | | + | 0,0175637 | 0,0570895 | 0,758347 | 2,39181E-08 |
| **rs311813** | 14 | 59086392 | G | A | 0,5256 | ENSG00000032219 | ARID4A | | N | -0,0148024 | 0,0444143 | 0,738922 | 0,000000038 |
| **rs311813** | 14 | 59086392 | G | A | 0,5256 | ENSG00000050130 | JKAMP | | N | 0,003885 | 0,0357494 | 0,913462 | 0,000000038 |
| **rs311813** | 14 | 59086392 | G | A | 0,5256 | ENSG00000100567 | PSMA3 | | N | -0,043701 | 0,0459708 | 0,341795 | 0,000000038 |
| **rs311813** | 14 | 59086392 | G | A | 0,5256 | ENSG00000100575 | TIMM9 | | N | -0,0348624 | 0,0427361 | 0,414638 | 0,000000038 |
| **rs311813** | 14 | 59086392 | G | A | 0,5256 | ENSG00000100578 | KIAA0586 | | N | 0,0333652 | 0,043755 | 0,445735 | 0,000000038 |
| **rs311813** | 14 | 59086392 | G | A | 0,5256 | ENSG00000100592 | DAAM1 | | N | -0,0133644 | 0,04392 | 0,760907 | 0,000000038 |
| **rs311813** | 14 | 59086392 | G | A | 0,5256 | ENSG00000126790 | L3HYPDH | | N | 0,00924325 | 0,0551158 | 0,866815 | 0,000000038 |
| **rs311813** | 14 | 59086392 | G | A | 0,5256 | ENSG00000131966 | ACTR10 | | N | 0,099656 | 0,0481603 | 0,0385218 | 0,000000038 |
| **rs311813** | 14 | 59086392 | G | A | 0,5256 | ENSG00000139970 | RTN1 | | N | 0,141277 | 0,0634824 | 0,0260515 | 0,000000038 |
| **rs311813** | 14 | 59086392 | G | A | 0,5256 | ENSG00000139971 | C14orf37 | | - | -0,0259407 | 0,0543321 | 0,633044 | 0,000000038 |
| **rs311813** | 14 | 59086392 | G | A | 0,5256 | ENSG00000151812 | SLC35F4 | | - | -0,0561293 | 0,053232 | 0,291687 | 0,000000038 |
| **rs311813** | 14 | 59086392 | G | A | 0,5256 | ENSG00000151838 | CCDC175 | | - | -0,145698 | 0,0585337 | 0,0128056 | 0,000000038 |

| | | | | | | | | | | | | | |
|---|---|---|---|---|---|---|---|---|---|---|---|---|---|
| **rs311813** | 14 | 59086392 | G | A | 0,5256 | ENSG00000165617 | DACT1 | | + | -0,166127 | 0,0637671 | 0,00918163 | 0,000000038 |
| **rs311813** | 14 | 59086392 | G | A | 0,5256 | ENSG00000180189 | HMGB1P14 | | + | 0,067263 | 0,0402266 | 0,0945038 | 0,000000038 |
| **rs311813** | 14 | 59086392 | G | A | 0,5256 | ENSG00000181619 | GPR135 | | - | 0,0455558 | 0,0587057 | 0,437748 | 0,000000038 |
| **rs311813** | 14 | 59086392 | G | A | 0,5256 | ENSG00000196860 | TOMM20L | | + | 0,0264955 | 0,092134 | 0,773671 | 0,000000038 |
| **rs311813** | 14 | 59086392 | G | A | 0,5256 | ENSG00000213598 | RP11-112J1.1 | LINC01500:1 | + | -0,0153034 | 0,0483844 | 0,751784 | 0,000000038 |
| **rs311813** | 14 | 59086392 | G | A | 0,5256 | ENSG00000239510 | RPL9P5 | | + | -0,12878 | 0,13895 | 0,354026 | 0,000000038 |
| **rs311813** | 14 | 59086392 | G | A | 0,5256 | ENSG00000257621 | RP11-349A22.5 | PSMA3-AS1 | N | 0,118004 | 0,0492498 | 0,0165742 | 0,000000038 |
| **rs311813** | 14 | 59086392 | G | A | 0,5256 | ENSG00000258378 | RP11-517O13.1 | lnc-TOMM20L-1:1 | + | -0,105957 | 0,0492246 | 0,031356 | 0,000000038 |
| **rs311813** | 14 | 59086392 | G | A | 0,5256 | ENSG00000258682 | CTD-2002H8.2 | lnc-ARID4A-4:1 | + | 0,00285381 | 0,0435317 | 0,947731 | 0,000000038 |
| **rs311813** | 14 | 59086392 | G | A | 0,5256 | ENSG00000258734 | RP11-112J1.3 | | - | 0,0212167 | 0,0372954 | 0,569435 | 0,000000038 |
| **rs311813** | 14 | 59086392 | G | A | 0,5256 | ENSG00000258782 | RP11-701B16.2 | lnc-CCDC175-1:1 | - | -0,0966107 | 0,054228 | 0,0748202 | 0,000000038 |
| **rs311813** | 14 | 59086392 | G | A | 0,5256 | ENSG00000258856 | CTD-2325K12.1 | | + | -0,131678 | 0,100074 | 0,188239 | 0,000000038 |
| **rs311813** | 14 | 59086392 | G | A | 0,5256 | ENSG00000258900 | HNRNPCP1 | | + | 0,0387021 | 0,0479256 | 0,419353 | 0,000000038 |
| **rs311813** | 14 | 59086392 | G | A | 0,5256 | ENSG00000259969 | RP11-999E24.3 | lnc-C14orf37-1:1 | N | -0,112139 | 0,0500118 | 0,0249449 | 0,000000038 |
| **rs311813** | 14 | 59086392 | G | A | 0,5256 | ENSG00000268466 | AL132989.1 | | + | 0,0175656 | 0,0570955 | 0,758347 | 0,000000038 |
| **rs1742882** | 14 | 59086975 | G | A | 0,4878 | ENSG00000032219 | ARID4A | | N | -0,00908481 | 0,0443972 | 0,837865 | 2,94919E-08 |
| **rs1742882** | 14 | 59086975 | G | A | 0,4878 | ENSG00000050130 | JKAMP | | N | 0,00919237 | 0,0357319 | 0,796978 | 2,94919E-08 |
| **rs1742882** | 14 | 59086975 | G | A | 0,4878 | ENSG00000100567 | PSMA3 | | N | -0,0526529 | 0,0459265 | 0,251604 | 2,94919E-08 |
| **rs1742882** | 14 | 59086975 | G | A | 0,4878 | ENSG00000100575 | TIMM9 | | N | -0,0372708 | 0,0427324 | 0,383105 | 2,94919E-08 |
| **rs1742882** | 14 | 59086975 | G | A | 0,4878 | ENSG00000100578 | KIAA0586 | | N | 0,0375523 | 0,0437223 | 0,390406 | 2,94919E-08 |
| **rs1742882** | 14 | 59086975 | G | A | 0,4878 | ENSG00000100592 | DAAM1 | | N | -0,0098114 | 0,0438988 | 0,823146 | 2,94919E-08 |
| **rs1742882** | 14 | 59086975 | G | A | 0,4878 | ENSG00000126790 | L3HYPDH | | N | 0,00965763 | 0,0551118 | 0,860893 | 2,94919E-08 |
| **rs1742882** | 14 | 59086975 | G | A | 0,4878 | ENSG00000131966 | ACTR10 | | N | 0,125135 | 0,048043 | 0,00919713 | 2,94919E-08 |
| **rs1742882** | 14 | 59086975 | G | A | 0,4878 | ENSG00000139970 | RTN1 | | N | 0,141272 | 0,0634802 | 0,0260515 | 2,94919E-08 |
| **rs1742882** | 14 | 59086975 | G | A | 0,4878 | ENSG00000139971 | C14orf37 | | - | -0,0514066 | 0,0542246 | 0,343114 | 2,94919E-08 |
| **rs1742882** | 14 | 59086975 | G | A | 0,4878 | ENSG00000151812 | SLC35F4 | | - | -0,0844773 | 0,0531469 | 0,111947 | 2,94919E-08 |
| **rs1742882** | 14 | 59086975 | G | A | 0,4878 | ENSG00000151838 | CCDC175 | | - | -0,183601 | 0,0582162 | 0,0016117 | 2,94919E-08 |
| **rs1742882** | 14 | 59086975 | G | A | 0,4878 | ENSG00000165617 | DACT1 | | + | -0,196722 | 0,0635076 | 0,00195084 | 2,94919E-08 |
| **rs1742882** | 14 | 59086975 | G | A | 0,4878 | ENSG00000180189 | HMGB1P14 | | + | 0,0448023 | 0,0401325 | 0,264268 | 2,94919E-08 |
| **rs1742882** | 14 | 59086975 | G | A | 0,4878 | ENSG00000181619 | GPR135 | | - | 0,0400901 | 0,0584713 | 0,492942 | 2,94919E-08 |
| **rs1742882** | 14 | 59086975 | G | A | 0,4878 | ENSG00000196860 | TOMM20L | | + | 0,0901601 | 0,0918053 | 0,32606 | 2,94919E-08 |
| **rs1742882** | 14 | 59086975 | G | A | 0,4878 | ENSG00000213598 | RP11-112J1.1 | LINC01500:1 | + | 0,00486901 | 0,0483925 | 0,919856 | 2,94919E-08 |
| **rs1742882** | 14 | 59086975 | G | A | 0,4878 | ENSG00000239510 | RPL9P5 | | + | -0,116061 | 0,138919 | 0,403459 | 2,94919E-08 |
| **rs1742882** | 14 | 59086975 | G | A | 0,4878 | ENSG00000257621 | RP11-349A22.5 | PSMA3-AS1 | N | 0,107803 | 0,0492919 | 0,0287399 | 2,94919E-08 |
| **rs1742882** | 14 | 59086975 | G | A | 0,4878 | ENSG00000258378 | RP11-517O13.1 | lnc-TOMM20L-1:1 | + | -0,118096 | 0,0491703 | 0,0163154 | 2,94919E-08 |
| **rs1742882** | 14 | 59086975 | G | A | 0,4878 | ENSG00000258682 | CTD-2002H8.2 | lnc-ARID4A-4:1 | + | 0,0190861 | 0,0434553 | 0,660508 | 2,94919E-08 |
| **rs1742882** | 14 | 59086975 | G | A | 0,4878 | ENSG00000258734 | RP11-112J1.3 | | - | 0,0358299 | 0,0374139 | 0,338232 | 2,94919E-08 |
| **rs1742882** | 14 | 59086975 | G | A | 0,4878 | ENSG00000258782 | RP11-701B16.2 | lnc-CCDC175-1:1 | - | -0,144522 | 0,0539596 | 0,00739886 | 2,94919E-08 |
| **rs1742882** | 14 | 59086975 | G | A | 0,4878 | ENSG00000258856 | CTD-2325K12.1 | | + | -0,144755 | 0,0998736 | 0,147231 | 2,94919E-08 |
| **rs1742882** | 14 | 59086975 | G | A | 0,4878 | ENSG00000258900 | HNRNPCP1 | | + | 0,0381158 | 0,0478581 | 0,425781 | 2,94919E-08 |
| **rs1742882** | 14 | 59086975 | G | A | 0,4878 | ENSG00000259969 | RP11-999E24.3 | lnc-C14orf37-1:1 | N | -0,116279 | 0,0499715 | 0,0199708 | 2,94919E-08 |
| **rs1742882** | 14 | 59086975 | G | A | 0,4878 | ENSG00000268466 | AL132989.1 | | + | 0,0501016 | 0,0569367 | 0,378885 | 2,94919E-08 |
| **rs191103** | 14 | 59092075 | T | C | 0,4552 | ENSG00000032219 | ARID4A | | N | -0,0167055 | 0,0444671 | 0,707153 | 1,5785E-08 |
| **rs191103** | 14 | 59092075 | T | C | 0,4552 | ENSG00000050130 | JKAMP | | N | 0,0146171 | 0,0357752 | 0,682847 | 1,5785E-08 |
| **rs191103** | 14 | 59092075 | T | C | 0,4552 | ENSG00000100567 | PSMA3 | | N | -0,0852262 | 0,0458617 | 0,0631224 | 1,5785E-08 |
| **rs191103** | 14 | 59092075 | T | C | 0,4552 | ENSG00000100575 | TIMM9 | | N | -0,0206619 | 0,0427847 | 0,629147 | 1,5785E-08 |
| **rs191103** | 14 | 59092075 | T | C | 0,4552 | ENSG00000100578 | KIAA0586 | | N | 0,0387784 | 0,043771 | 0,375651 | 1,5785E-08 |
| **rs191103** | 14 | 59092075 | T | C | 0,4552 | ENSG00000100592 | DAAM1 | | N | -0,00494618 | 0,0439282 | 0,91035 | 1,5785E-08 |
| **rs191103** | 14 | 59092075 | T | C | 0,4552 | ENSG00000126790 | L3HYPDH | | N | 0,0241467 | 0,0550481 | 0,660917 | 1,5785E-08 |
| **rs191103** | 14 | 59092075 | T | C | 0,4552 | ENSG00000131966 | ACTR10 | | N | 0,115724 | 0,0480397 | 0,0159993 | 1,5785E-08 |
| **rs191103** | 14 | 59092075 | T | C | 0,4552 | ENSG00000139970 | RTN1 | | N | 0,134847 | 0,0633393 | 0,033258 | 1,5785E-08 |
| **rs191103** | 14 | 59092075 | T | C | 0,4552 | ENSG00000139971 | C14orf37 | | - | -0,0312875 | 0,0544509 | 0,565561 | 1,5785E-08 |
| **rs191103** | 14 | 59092075 | T | C | 0,4552 | ENSG00000151812 | SLC35F4 | | - | -0,0788165 | 0,0533955 | 0,13992 | 1,5785E-08 |
| **rs191103** | 14 | 59092075 | T | C | 0,4552 | ENSG00000151838 | CCDC175 | | - | -0,185994 | 0,0585086 | 0,00147823 | 1,5785E-08 |
| **rs191103** | 14 | 59092075 | T | C | 0,4552 | ENSG00000165617 | DACT1 | | + | -0,186223 | 0,0638244 | 0,00352576 | 1,5785E-08 |
| **rs191103** | 14 | 59092075 | T | C | 0,4552 | ENSG00000180189 | HMGB1P14 | | + | 0,0305128 | 0,0403194 | 0,449183 | 1,5785E-08 |
| **rs191103** | 14 | 59092075 | T | C | 0,4552 | ENSG00000181619 | GPR135 | | - | 0,0504103 | 0,0587535 | 0,390894 | 1,5785E-08 |
| **rs191103** | 14 | 59092075 | T | C | 0,4552 | ENSG00000196860 | TOMM20L | | + | 0,0297015 | 0,0924391 | 0,747976 | 1,5785E-08 |
| **rs191103** | 14 | 59092075 | T | C | 0,4552 | ENSG00000213598 | RP11-112J1.1 | LINC01500:1 | + | 0,0279716 | 0,0484531 | 0,563741 | 1,5785E-08 |
| **rs191103** | 14 | 59092075 | T | C | 0,4552 | ENSG00000239510 | RPL9P5 | | + | -0,0757657 | 0,139711 | 0,58761 | 1,5785E-08 |
| **rs191103** | 14 | 59092075 | T | C | 0,4552 | ENSG00000257621 | RP11-349A22.5 | PSMA3-AS1 | N | 0,15712 | 0,0491789 | 0,001399 | 1,5785E-08 |
| **rs191103** | 14 | 59092075 | T | C | 0,4552 | ENSG00000258378 | RP11-517O13.1 | lnc-TOMM20L-1:1 | + | -0,104024 | 0,0493725 | 0,035125 | 1,5785E-08 |

| | | | | | | | | | | | | | |
|---|---|---|---|---|---|---|---|---|---|---|---|---|---|
| **rs191103** | 14 | 59092075 | T | C | 0,4552 | ENSG00000258682 | CTD-2002H8.2 | lnc-ARID4A-4:1 | + | 0,0377984 | 0,0436747 | 0,38679 | 1,5785E-08 |
| **rs191103** | 14 | 59092075 | T | C | 0,4552 | ENSG00000258734 | RP11-112J1.3 | | - | 0,0249938 | 0,0375257 | 0,505382 | 1,5785E-08 |
| **rs191103** | 14 | 59092075 | T | C | 0,4552 | ENSG00000258782 | RP11-701B16.2 | lnc-CCDC175-1:1 | - | -0,143015 | 0,054258 | 0,00839332 | 1,5785E-08 |
| **rs191103** | 14 | 59092075 | T | C | 0,4552 | ENSG00000258856 | CTD-2325K12.1 | | + | -0,16087 | 0,100129 | 0,108135 | 1,5785E-08 |
| **rs191103** | 14 | 59092075 | T | C | 0,4552 | ENSG00000258900 | HNRNPCP1 | | + | 0,0387627 | 0,0479183 | 0,418554 | 1,5785E-08 |
| **rs191103** | 14 | 59092075 | T | C | 0,4552 | ENSG00000259969 | RP11-999E24.3 | lnc-C14orf37-1:1 | N | -0,134993 | 0,0498953 | 0,00681964 | 1,5785E-08 |
| **rs191103** | 14 | 59092075 | T | C | 0,4552 | ENSG00000268466 | AL132989.1 | | + | 0,01796 | 0,0571478 | 0,753314 | 1,5785E-08 |
