## Supplementary material for "Enhancer locus in ch14q23.1 modulates brain asymmetric temporal regions involved in language processing": Table S12

| Phenotypes | Pvals | N Subjects |
|---|---|---|
| meandepth Morphologist STs_left | 1.4·10-13 | 18101 |
| SurfArea Destrieux Pole_temporal.lh | 5.8·10-12 | 17127 |
| GrayVol Desikan bankssts.lh | 7.6·10-8 | 17127 |
| SurfArea Destrieux G_cuneus.rh | 2.3·10-7 | 17127 |
| SurfArea Desikan bankssts.lh | 2.8·10-7 | 17127 |
| FA Tapetum_L | 5.7·10-7 | 16541 |
| SurfArea Desikan cuneus.rh | 6.6·10-7 | 17127 |
| ThickAvg Destrieux G_temporal_middle.rh | 6.9·10-7 | 17127 |
| ThickAvg Desikan bankssts.lh | 8.8·10-7 | 17127 |
| ThickAvg Desikan middletemporal.lh | 1.1·10-6 | 17127 |
| maxdepth Morphologist STiant_left | 1.5·10-6 | 18101 |
| ThickAvg Destrieux G_temporal_middle.lh | 1.9·10-6 | 17127 |
| rfMRI full corr ICA100 edge 1146 link 29-41 | 2.1·10-6 | 15859 |
| meandepth Morphologist STs_right | 2.3·10-6 | 18100 |
| SurfArea Destrieux Pole_temporal.rh | 2.4·10-6 | 17126 |
| ThickAvg Desikan bankssts.rh | 6.0·10-6 | 17127 |
| rfMRI full corr ICA25 edge 87 link 5-18 | 7.6·10-6 | 15864 |
| GrayVol Destrieux S_interm_prim-Jensen.lh | 7.6·10-6 | 17108 |
| GrayVol Desikan transversetemporal.lh | 7.9·10-6 | 17126 |
| surface Morphologist STiant_left | 8.6·10-6 | 18101 |
| hull_junction_length Morphologist STiant_left | 1.2·10-5 | 18101 |
| SurfArea Destrieux G_cuneus.lh | 1.4·10-5 | 17127 |
| GrayVol Destrieux S_circular_insula_inf.lh | 1.4·10-5 | 17126 |
| MO Cerebral_peduncle_L | 1.4·10-5 | 16541 |
| SurfArea Destrieux S_circular_insula_inf.lh | 1.6·10-5 | 17126 |
| ThickAvg Desikan middletemporal.rh | 2.3·10-5 | 17127 |
| ThickAvg Destrieux S_temporal_sup.lh | 2.5·10-5 | 17127 |
| rfMRI full corr ICA100 edge 385 link 8-36 | 2.5·10-5 | 15863 |
| SurfArea Desikan cuneus.lh | 2.6·10-5 | 17127 |
| maxdepth Morphologist STs_left | 2.8·10-5 | 18101 |
| GrayVol Desikan supramarginal.rh | 2.9·10-5 | 17127 |
| GrayVol Destrieux G_pariet_inf-Supramar.rh | 2.9·10-5 | 17127 |

| | | |
|---|---|---|
| **SurfArea Destrieux S_interm_prim-Jensen.lh** | 3.2·10-5 | 17108 |
| **opening Morphologist SOTlatint_left** | 3.5·10-5 | 16594 |
| **SurfArea Destrieux G_pariet_inf-Supramar.rh** | 3.6·10-5 | 17127 |
| **GrayVol Destrieux G_temp_sup-G_T_transv.lh** | 9.7·10-5 | 17126 |
| **hull_junction_length Morphologist SOTlatant_left** | 1.0·10-4 | 18076 |
| **GrayVol Desikan bankssts.rh** | 1.0·10-4 | 17127 |
| **GrayVol Destrieux G_cingul-Post-ventral.rh** | 1.1·10-4 | 17127 |
| **ThickAvg Destrieux S_circular_insula_inf.rh** | 1.2·10-4 | 17125 |
| **maxdepth Morphologist OCCIPITAL_left** | 1.6·10-4 | 18097 |
| **rfMRI part corr ICA100 edge 1146 link 29-41** | 1.6·10-4 | 15864 |
| **maxdepth Morphologist SPat_right** | 1.7·10-4 | 16760 |
| **rfMRI full corr ICA100 edge 787 link 18-23** | 1.8·10-4 | 15864 |
| **rfMRI part corr ICA100 edge 1133 link 28-54** | 1.8·10-4 | 15864 |
| **MO Anterior_limb_of_internal_capsule_R** | 1.9·10-4 | 16541 |
| **maxdepth Morphologist SPat_left** | 1.9·10-4 | 13931 |
| **MO Cerebral_peduncle_R** | 2.0·10-4 | 16541 |
| **SurfArea Desikan supramarginal.rh** | 2.2·10-4 | 17127 |
| **rfMRI full corr ICA100 edge 1094 link 27-42** | 2.3·10-4 | 15861 |
| **rfMRI part corr ICA100 edge 1130 link 28-51** | 2.4·10-4 | 15864 |
| **SurfArea Desikan transversetemporal.lh** | 2.5·10-4 | 17126 |
| **rfMRI full corr ICA100 edge 1130 link 28-51** | 2.5·10-4 | 15863 |
| **GrayVol Destrieux G_cingul-Post-ventral.lh** | 2.6·10-4 | 17127 |
| **rfMRI full corr ICA100 edge 584 link 13-15** | 2.9·10-4 | 15864 |
| **rfMRI full corr ICA100 edge 793 link 18-29** | 2.9·10-4 | 15864 |
| **FA Tapetum_R** | 3.0·10-4 | 16541 |
| **ThickAvg Destrieux G_temp_sup-Plan_tempo.lh** | 3.5·10-4 | 17127 |
| **GrayVol Destrieux Pole_temporal.lh** | 3.8·10-4 | 17127 |
| **ThickAvg Destrieux S_temporal_sup.rh** | 4.2·10-4 | 17127 |
| **rfMRI full corr ICA100 edge 1112 link 28-33** | 4.5·10-4 | 15864 |
| **GrayVol Destrieux S_intrapariet_and_P_trans.lh** | 4.6·10-4 | 17127 |
| **MO Tapetum_R** | 4.6·10-4 | 16541 |
| **rfMRI part corr ICA25 edge 87 link 5-18** | 4.9·10-4 | 15864 |
| **surface Morphologist STs_right** | 5.3·10-4 | 18100 |
| **rfMRI full corr ICA100 edge 535 link 11-51** | 5.3·10-4 | 15863 |

| | | |
|---|---|---|
| **rfMRI full corr ICA100 edge 663 link 14-53** | 5.7·10-4 | 15864 |
| **GrayVol Destrieux G_parietal_sup.lh** | 5.9·10-4 | 17127 |
| **rfMRI full corr ICA100 edge 975 link 23-41** | 5.9·10-4 | 15864 |
| **surface Morphologist FCalant-ScCal_left** | 5.9·10-4 | 18100 |
| **hull_junction_length Morphologist STs_left** | 6.0·10-4 | 18101 |
| **rfMRI part corr ICA100 edge 1000 link 24-35** | 6.9·10-4 | 15864 |
| **GrayVol Destrieux G_precuneus.lh** | 7.2·10-4 | 17127 |
| **hull_junction_length Morphologist SRh_left** | 7.8·10-4 | 17719 |
| **SurfArea Desikan caudalmiddlefrontal.rh** | 7.8·10-4 | 17127 |
| **GrayVol Destrieux G_pariet_inf-Angular.lh** | 7.9·10-4 | 17127 |
| **SurfArea Destrieux S_front_sup.rh** | 8.1·10-4 | 17127 |
| **opening Morphologist SRh_left** | 8.3·10-4 | 17719 |
| **rfMRI full corr ICA100 edge 391 link 8-42** | 8.4·10-4 | 15861 |
| **maxdepth Morphologist FCLrasc_left** | 8.5·10-4 | 17784 |
| **rfMRI part corr ICA100 edge 1141 link 29-36** | 8.9·10-4 | 15864 |
| **rfMRI part corr ICA100 edge 635 link 14-25** | 8.9·10-4 | 15864 |
| **surface Morphologist SOp_right** | 9.0·10-4 | 17447 |
| **rfMRI part corr ICA100 edge 529 link 11-45** | 9.1·10-4 | 15864 |
| **surface Morphologist FIPPoCinf_right** | 9.5·10-4 | 18073 |
| **rfMRI part corr ICA100 edge 1065 link 26-41** | 9.5·10-4 | 15864 |
| **SurfArea Destrieux S_parieto_occipital.lh** | 1.0·10-3 | 17127 |
| **rfMRI part corr ICA100 edge 178 link 4-23** | 1.0·10-3 | 15864 |
| **SurfArea Destrieux G_temp_sup-G_T_transv.lh** | 1.1·10-3 | 17126 |
| **SurfArea Desikan temporalpole.rh** | 1.1·10-3 | 17126 |
| **rfMRI full corr ICA100 edge 1133 link 28-54** | 1.1·10-3 | 15864 |
| **surface Morphologist SRh_left** | 1.1·10-3 | 17719 |
| **GrayVol Destrieux S_front_sup.rh** | 1.1·10-3 | 17127 |
| **rfMRI part corr ICA100 edge 52 link 1-53** | 1.2·10-3 | 15864 |
| **OD Superior_corona_radiata_R** | 1.3·10-3 | 16541 |
| **SurfArea Destrieux S_intrapariet_and_P_trans.lh** | 1.3·10-3 | 17127 |
| **ThickAvg Destrieux S_circular_insula_inf.lh** | 1.3·10-3 | 17126 |
| **rfMRI full corr ICA100 edge 1374 link 40-49** | 1.5·10-3 | 15864 |
| **rfMRI part corr ICA100 edge 1167 link 30-37** | 1.5·10-3 | 15864 |
| **rfMRI part corr ICA25 edge 144 link 9-21** | 1.5·10-3 | 15864 |

| | | |
|---|---|---|
| **rfMRI part corr ICA100 edge 1374 link 40-49** | 1.5·10-3 | 15864 |
| **rfMRI part corr ICA100 edge 458 link 10-18** | 1.5·10-3 | 15864 |
| **opening Morphologist FCLrant_right** | 1.6·10-3 | 16205 |
| **meandepth Morphologist SPat_left** | 1.6·10-3 | 13839 |
| **rfMRI part corr ICA100 edge 143 link 3-39** | 1.6·10-3 | 15864 |
| **SurfArea Desikan entorhinal.lh** | 1.7·10-3 | 17127 |
| **GrayVol Destrieux S_circular_insula_inf.rh** | 1.7·10-3 | 17125 |
| **rfMRI part corr ICA100 edge 33 link 1-34** | 1.7·10-3 | 15864 |
| **GrayVol Destrieux G_cuneus.rh** | 1.7·10-3 | 17127 |
| **rfMRI full corr ICA100 edge 1174 link 30-44** | 1.8·10-3 | 15864 |
| **SurfArea Destrieux G_oc-temp_med-Parahip.rh** | 1.8·10-3 | 17127 |
| **SurfArea Desikan temporalpole.lh** | 1.9·10-3 | 17127 |
| **rfMRI full corr ICA100 edge 505 link 11-21** | 1.9·10-3 | 15864 |
| **ThickAvg Destrieux Pole_temporal.lh** | 2.0·10-3 | 17127 |
| **opening Morphologist SOTlatant_left** | 2.1·10-3 | 18076 |
| **GrayVol Desikan caudalmiddlefrontal.rh** | 2.1·10-3 | 17127 |
| **rfMRI amplitude ICA100 component 38** | 2.1·10-3 | 15863 |
| **MO Anterior_limb_of_internal_capsule_L** | 2.2·10-3 | 16541 |
| **SurfArea Destrieux S_central.rh** | 2.3·10-3 | 17127 |
| **rfMRI full corr ICA100 edge 52 link 1-53** | 2.4·10-3 | 15864 |
| **rfMRI full corr ICA100 edge 383 link 8-34** | 2.4·10-3 | 15862 |
| **rfMRI full corr ICA100 edge 1141 link 29-36** | 2.5·10-3 | 15864 |
| **GrayVol Destrieux G_precuneus.rh** | 2.5·10-3 | 17127 |
| **ThickAvg Destrieux G_cingul-Post-ventral.rh** | 2.5·10-3 | 17127 |
| **rfMRI full corr ICA100 edge 1162 link 30-32** | 2.6·10-3 | 15863 |
| **rfMRI part corr ICA100 edge 1112 link 28-33** | 2.6·10-3 | 15864 |
| **OD Posterior_limb_of_internal_capsule_R** | 2.6·10-3 | 16541 |
| **GrayVol Destrieux G_cuneus.lh** | 2.7·10-3 | 17127 |
| **surface Morphologist SPat_right** | 2.7·10-3 | 16760 |
| **SurfArea Destrieux S_front_sup.lh** | 2.8·10-3 | 17127 |
| **ThickAvg Destrieux S_calcarine.rh** | 2.8·10-3 | 17127 |
| **maxdepth Morphologist FIP_right** | 2.9·10-3 | 18097 |
| **rfMRI full corr ICA25 edge 80 link 5-11** | 2.9·10-3 | 15864 |
| **rfMRI full corr ICA100 edge 541 link 12-14** | 3.0·10-3 | 15864 |

| | | |
|---|---|---|
| **rfMRI full corr ICA100 edge 579 link 12-52** | 3.0·10-3 | 15863 |
| **GrayVol Desikan cuneus.rh** | 3.1·10-3 | 17127 |
| **rfMRI full corr ICA100 edge 381 link 8-32** | 3.2·10-3 | 15864 |
| **opening Morphologist SFpolairetr_left** | 3.2·10-3 | 18062 |
| **MD Cerebral_peduncle_L** | 3.3·10-3 | 16541 |
| **rfMRI full corr ICA100 edge 143 link 3-39** | 3.4·10-3 | 15862 |
| **rfMRI part corr ICA100 edge 1131 link 28-52** | 3.4·10-3 | 15864 |
| **rfMRI full corr ICA100 edge 127 link 3-23** | 3.4·10-3 | 15864 |
| **rfMRI full corr ICA100 edge 178 link 4-23** | 3.5·10-3 | 15864 |
| **meandepth Morphologist STiant_left** | 3.5·10-3 | 18101 |
| **rfMRI part corr ICA100 edge 1025 link 25-30** | 3.6·10-3 | 15864 |
| **SurfArea Destrieux S_occipital_ant.rh** | 3.6·10-3 | 17127 |
| **rfMRI part corr ICA100 edge 584 link 13-15** | 3.6·10-3 | 15864 |
| **rfMRI part corr ICA100 edge 775 link 17-48** | 3.7·10-3 | 15864 |
| **rfMRI full corr ICA100 edge 498 link 11-14** | 3.7·10-3 | 15864 |
| **opening Morphologist SLiant_left** | 3.7·10-3 | 17423 |
| **surface Morphologist SPat_left** | 3.7·10-3 | 13931 |
| **rfMRI part corr ICA100 edge 45 link 1-46** | 3.8·10-3 | 15864 |
| **rfMRI part corr ICA100 edge 528 link 11-44** | 3.8·10-3 | 15864 |
| **rfMRI part corr ICA100 edge 120 link 3-16** | 3.8·10-3 | 15864 |
| **rfMRI part corr ICA100 edge 383 link 8-34** | 3.8·10-3 | 15864 |
| **rfMRI part corr ICA100 edge 418 link 9-23** | 3.8·10-3 | 15864 |
| **rfMRI full corr ICA100 edge 1376 link 40-51** | 3.9·10-3 | 15864 |
| **rfMRI part corr ICA100 edge 55 link 2-3** | 4.1·10-3 | 15864 |
| **rfMRI full corr ICA100 edge 987 link 23-53** | 4.2·10-3 | 15864 |
| **meandepth Morphologist SOp_right** | 4.2·10-3 | 17422 |
| **rfMRI part corr ICA100 edge 204 link 4-49** | 4.2·10-3 | 15864 |
| **OD Cerebral_peduncle_R** | 4.4·10-3 | 16541 |
| **maxdepth Morphologist OCCIPITAL_right** | 4.4·10-3 | 18096 |
| **GrayVol Destrieux G_temp_sup-Plan_polar.lh** | 4.5·10-3 | 17126 |
| **SurfArea Destrieux G_front_middle.lh** | 4.6·10-3 | 17127 |
| **rfMRI part corr ICA100 edge 982 link 23-48** | 4.8·10-3 | 15864 |
| **GM_thickness Morphologist STiant_left** | 4.8·10-3 | 18091 |
| **rfMRI full corr ICA100 edge 147 link 3-43** | 5.0·10-3 | 15856 |

| | | |
|---|---|---|
| **FA Superior_corona_radiata_R** | 5.0·10-3 | 16541 |
| **GrayVol Destrieux Pole_temporal.rh** | 5.1·10-3 | 17126 |
| **rfMRI part corr ICA100 edge 249 link 5-44** | 5.2·10-3 | 15864 |
| **MO Tapetum_L** | 5.5·10-3 | 16541 |
| **rfMRI full corr ICA100 edge 33 link 1-34** | 5.5·10-3 | 15864 |
| **GrayVol Destrieux G_insular_short.lh** | 5.5·10-3 | 17126 |
| **maxdepth Morphologist SOp_right** | 5.6·10-3 | 17447 |
| **GrayVol Destrieux S_occipital_ant.rh** | 5.7·10-3 | 17127 |
| **opening Morphologist SFpolairetr_right** | 5.7·10-3 | 18086 |
| **rfMRI part corr ICA100 edge 108 link 3-4** | 5.8·10-3 | 15864 |
| **rfMRI full corr ICA100 edge 1098 link 27-46** | 5.8·10-3 | 15864 |
| **meandepth Morphologist SOTlatint_left** | 5.8·10-3 | 16588 |
| **rfMRI part corr ICA100 edge 1098 link 27-46** | 5.8·10-3 | 15864 |
| **rfMRI part corr ICA100 edge 53 link 1-54** | 5.9·10-3 | 15864 |
| **GrayVol Desikan inferiorparietal.lh** | 5.9·10-3 | 17127 |
| **rfMRI full corr ICA100 edge 1065 link 26-41** | 6.0·10-3 | 15864 |
| **rfMRI part corr ICA100 edge 1371 link 40-46** | 6.0·10-3 | 15864 |
| **rfMRI full corr ICA100 edge 563 link 12-36** | 6.2·10-3 | 15864 |
| **rfMRI part corr ICA100 edge 1481 link 52-54** | 6.2·10-3 | 15864 |
| **rfMRI full corr ICA100 edge 91 link 2-39** | 6.3·10-3 | 15864 |
| **SurfArea Destrieux G_oc-temp_med-Parahip.lh** | 6.4·10-3 | 17127 |
| **rfMRI part corr ICA100 edge 1091 link 27-39** | 6.4·10-3 | 15864 |
| **SurfArea Destrieux G_precentral.rh** | 6.5·10-3 | 17127 |
| **GrayVol Destrieux G_oc-temp_med-Parahip.rh** | 6.6·10-3 | 17127 |
| **rfMRI part corr ICA100 edge 1092 link 27-40** | 6.6·10-3 | 15864 |
| **rfMRI part corr ICA100 edge 1353 link 39-43** | 6.6·10-3 | 15864 |
| **GrayVol Desikan parahippocampal.lh** | 6.7·10-3 | 17127 |
| **rfMRI full corr ICA100 edge 1159 link 29-54** | 6.7·10-3 | 15863 |
| **OD Cerebral_peduncle_L** | 6.8·10-3 | 16541 |
| **SurfArea Destrieux G_precuneus.lh** | 6.8·10-3 | 17127 |
| **rfMRI full corr ICA25 edge 81 link 5-12** | 6.9·10-3 | 15864 |
| **SurfArea Destrieux G_pariet_inf-Angular.lh** | 6.9·10-3 | 17127 |
| **rfMRI full corr ICA100 edge 120 link 3-16** | 6.9·10-3 | 15864 |
| **rfMRI full corr ICA100 edge 104 link 2-52** | 6.9·10-3 | 15864 |

| | | |
|---|---|---|
| **OD Tapetum_L** | 7.0·10-3 | 16541 |
| **GrayVol Desikan superiorparietal.lh** | 7.0·10-3 | 17127 |
| **rfMRI full corr ICA100 edge 1157 link 29-52** | 7.1·10-3 | 15864 |
| **rfMRI part corr ICA100 edge 237 link 5-32** | 7.1·10-3 | 15864 |
| **rfMRI part corr ICA100 edge 1088 link 27-36** | 7.1·10-3 | 15864 |
| **rfMRI full corr ICA100 edge 204 link 4-49** | 7.1·10-3 | 15864 |
| **SurfArea Destrieux G_insular_short.lh** | 7.1·10-3 | 17126 |
| **rfMRI part corr ICA100 edge 1159 link 29-54** | 7.2·10-3 | 15864 |
| **meandepth Morphologist OCCIPITAL_left** | 7.3·10-3 | 18097 |
| **opening Morphologist SOTlatant_right** | 7.5·10-3 | 18047 |
| **GrayVol Desikan lingual.lh** | 7.5·10-3 | 17127 |
| **rfMRI full corr ICA100 edge 1444 link 46-50** | 7.6·10-3 | 15864 |
| **SurfArea Desikan precentral.rh** | 7.7·10-3 | 17127 |
| **rfMRI full corr ICA100 edge 1058 link 26-34** | 7.7·10-3 | 15864 |
| **rfMRI full corr ICA100 edge 237 link 5-32** | 7.7·10-3 | 15864 |
| **OD Sagittal_stratum-inf_longitidinal_fasci_and_inf** | 7.7·10-3 | 16541 |
| **GrayVol Destrieux S_temporal_sup.rh** | 8.0·10-3 | 17127 |
| **SurfArea Destrieux S_collat_transv_post.rh** | 8.0·10-3 | 17126 |
| **FA Cingulum-hippocampus-L** | 8.1·10-3 | 16541 |
| **rfMRI full corr ICA100 edge 1357 link 39-47** | 8.2·10-3 | 15864 |
| **rfMRI part corr ICA100 edge 386 link 8-37** | 8.2·10-3 | 15864 |
| **hull_junction_length Morphologist FCLrdiag_right** | 8.4·10-3 | 11519 |
| **rfMRI full corr ICA100 edge 1170 link 30-40** | 8.4·10-3 | 15862 |
| **rfMRI full corr ICA100 edge 403 link 8-54** | 8.4·10-3 | 15864 |
| **SurfArea Destrieux G_front_inf-Triangul.rh** | 8.5·10-3 | 17127 |
| **rfMRI full corr ICA100 edge 1025 link 25-30** | 8.5·10-3 | 15864 |
| **rfMRI full corr ICA100 edge 404 link 8-55** | 8.6·10-3 | 15864 |
| **rfMRI part corr ICA100 edge 753 link 17-26** | 8.7·10-3 | 15864 |
| **rfMRI full corr ICA100 edge 185 link 4-30** | 8.9·10-3 | 15864 |
| **rfMRI full corr ICA100 edge 1086 link 27-34** | 9.0·10-3 | 15862 |
| **rfMRI full corr ICA100 edge 74 link 2-22** | 9.1·10-3 | 15864 |
| **rfMRI part corr ICA100 edge 1202 link 31-48** | 9.1·10-3 | 15864 |
| **rfMRI part corr ICA100 edge 975 link 23-41** | 9.1·10-3 | 15864 |
| **rfMRI full corr ICA100 edge 1353 link 39-43** | 9.2·10-3 | 15864 |

| | | |
|---|---|---|
| **hull_junction_length Morphologist INSULA_left** | 9.2·10-3 | 18101 |
| **rfMRI part corr ICA100 edge 329 link 7-27** | 9.2·10-3 | 15864 |
| **MD Cerebral_peduncle_R** | 9.2·10-3 | 16541 |
| **Left-Amygdala** | 9.3·10-3 | 17127 |
| **rfMRI amplitude ICA25 component 18** | 9.3·10-3 | 15863 |
| **GrayVol Desikan parstriangularis.rh** | 9.5·10-3 | 17127 |
| **maxdepth Morphologist SFinfant_left** | 9.6·10-3 | 17653 |
| **ThickAvg Desikan temporalpole.lh** | 9.6·10-3 | 17127 |
| **rfMRI full corr ICA100 edge 442 link 9-47** | 9.7·10-3 | 15862 |
| **OD Uncinate_fasciculus_L** | 9.8·10-3 | 16541 |
| **surface Morphologist FCalant-ScCal_right** | 9.8·10-3 | 18100 |
| **rfMRI full corr ICA100 edge 1085 link 27-33** | 9.8·10-3 | 15861 |
| **rfMRI amplitude ICA100 component 52** | 9.9·10-3 | 15863 |
| **FA Cingulum-hippocampus-R** | 1.0·10-2 | 16541 |
| **GrayVol Destrieux Lat_Fis-post.lh** | 0.01 | 17126 |
| **rfMRI part corr ICA100 edge 319 link 7-17** | 0.01 | 15864 |
| **rfMRI part corr ICA100 edge 1004 link 24-39** | 0.01 | 15864 |
| **rfMRI full corr ICA100 edge 785 link 18-21** | 0.01 | 15864 |
| **rfMRI part corr ICA100 edge 921 link 21-52** | 0.01 | 15864 |
| **GrayVol Destrieux G_temp_sup-Plan_tempo.lh** | 0.01 | 17127 |
| **opening Morphologist SLiant_right** | 0.01 | 17219 |
| **rfMRI full corr ICA100 edge 921 link 21-52** | 0.01 | 15864 |
| **GrayVol Destrieux S_central.rh** | 0.01 | 17127 |
| **rfMRI full corr ICA100 edge 1458 link 48-49** | 0.01 | 15864 |
| **rfMRI full corr ICA100 edge 1449 link 46-55** | 0.01 | 15864 |
| **meandepth Morphologist FCLrasc_left** | 0.01 | 17626 |
| **SurfArea Destrieux Lat_Fis-ant-Vertical.rh** | 0.01 | 17114 |
| **rfMRI full corr ICA100 edge 153 link 3-49** | 0.01 | 15864 |
| **rfMRI full corr ICA100 edge 628 link 14-18** | 0.01 | 15863 |
| **ThickAvg Desikan pericalcarine.rh** | 0.01 | 17127 |
| **GrayVol Destrieux G_oc-temp_med-Lingual.lh** | 0.01 | 17127 |
| **opening Morphologist STs_left** | 0.01 | 18101 |
| **rfMRI part corr ICA100 edge 1041 link 25-46** | 0.01 | 15864 |
| **rfMRI full corr ICA100 edge 66 link 2-14** | 0.01 | 15864 |

| | | |
|---|---|---|
| **SurfArea Destrieux G_temp_sup-Plan_polar.lh** | 0.01 | 17126 |
| **GrayVol Destrieux G_front_inf-Triangul.lh** | 0.01 | 17127 |
| **rfMRI part corr ICA100 edge 1170 link 30-40** | 0.01 | 15864 |
| **meandepth Morphologist SCu_right** | 0.01 | 18067 |
| **hull_junction_length Morphologist SFinter_left** | 0.01 | 18101 |
| **ThickAvg Destrieux S_oc-temp_med_and_Lingual.** | 0.01 | 17127 |
| **rfMRI part corr ICA100 edge 1139 link 29-34** | 0.01 | 15864 |
| **hull_junction_length Morphologist SFinfant_left** | 0.01 | 17653 |
| **rfMRI part corr ICA100 edge 637 link 14-27** | 0.01 | 15864 |
| **rfMRI part corr ICA100 edge 1174 link 30-44** | 0.01 | 15864 |
| **rfMRI part corr ICA100 edge 98 link 2-46** | 0.01 | 15864 |
| **rfMRI part corr ICA100 edge 338 link 7-36** | 0.01 | 15864 |
| **GrayVol Destrieux G_and_S_paracentral.rh** | 0.01 | 17127 |
| **rfMRI part corr ICA100 edge 551 link 12-24** | 0.01 | 15864 |
| **rfMRI part corr ICA100 edge 1094 link 27-42** | 0.01 | 15864 |
| **MO Posterior_thalamic_radiation-include_optic_ra** | 0.01 | 16541 |
| **GrayVol Destrieux S_orbital_med-olfact.rh** | 0.01 | 17125 |
| **rfMRI full corr ICA25 edge 183 link 14-15** | 0.01 | 15863 |
| **surface Morphologist SCu_right** | 0.01 | 18068 |
| **GrayVol Destrieux S_front_sup.lh** | 0.01 | 17127 |
| **meandepth Morphologist OCCIPITAL_right** | 0.01 | 18096 |
| **rfMRI part corr ICA100 edge 496 link 11-12** | 0.01 | 15864 |
| **rfMRI part corr ICA100 edge 1053 link 26-29** | 0.01 | 15864 |
| **OD Posterior_limb_of_internal_capsule_L** | 0.01 | 16541 |
| **SurfArea Destrieux S_collat_transv_post.lh** | 0.01 | 17125 |
| **rfMRI full corr ICA100 edge 1372 link 40-47** | 0.01 | 15864 |
| **meandepth Morphologist FIPPoCinf_left** | 0.01 | 18078 |
| **surface Morphologist FCLrscpost_right** | 0.01 | 16126 |
| **hull_junction_length Morphologist FCLrscant_left** | 0.01 | 4323 |
| **rfMRI part corr ICA100 edge 56 link 2-4** | 0.01 | 15864 |
| **GrayVol Destrieux G_parietal_sup.rh** | 0.01 | 17127 |
| **rfMRI part corr ICA100 edge 772 link 17-45** | 0.01 | 15864 |
| **rfMRI full corr ICA100 edge 478 link 10-38** | 0.01 | 15864 |
| **rfMRI part corr ICA100 edge 591 link 13-22** | 0.01 | 15864 |

| | | |
|---|---|---|
| **SurfArea Destrieux Lat_Fis-post.lh** | 0.01 | 17126 |
| **rfMRI full corr ICA100 edge 1251 link 33-52** | 0.01 | 15864 |
| **rfMRI full corr ICA100 edge 910 link 21-41** | 0.01 | 15861 |
| **rfMRI part corr ICA100 edge 1459 link 48-50** | 0.01 | 15864 |
| **GrayVol Destrieux G_front_inf-Triangul.rh** | 0.01 | 17127 |
| **GrayVol Destrieux S_collat_transv_post.rh** | 0.01 | 17126 |
| **rfMRI full corr ICA100 edge 1323 link 37-46** | 0.01 | 15864 |
| **GM_thickness Morphologist SOTlatant_right** | 0.01 | 18038 |
| **rfMRI full corr ICA100 edge 48 link 1-49** | 0.01 | 15864 |
| **SurfArea Desikan pericalcarine.rh** | 0.01 | 17127 |
| **rfMRI part corr ICA100 edge 1122 link 28-43** | 0.01 | 15864 |
| **rfMRI full corr ICA100 edge 694 link 15-44** | 0.01 | 15864 |
| **rfMRI part corr ICA100 edge 1215 link 32-38** | 0.01 | 15864 |
| **FA Posterior_limb_of_internal_capsule_R** | 0.02 | 16541 |
| **MD Fornix-cres-Stria_terminalis-not_resolved_wit** | 0.02 | 16541 |
| **rfMRI part corr ICA100 edge 78 link 2-26** | 0.02 | 15864 |
| **rfMRI full corr ICA25 edge 90 link 5-21** | 0.02 | 15864 |
| **rfMRI part corr ICA100 edge 777 link 17-50** | 0.02 | 15864 |
| **rfMRI part corr ICA100 edge 740 link 16-51** | 0.02 | 15864 |
| **rfMRI full corr ICA100 edge 843 link 19-43** | 0.02 | 15864 |
| **rfMRI part corr ICA100 edge 640 link 14-30** | 0.02 | 15864 |
| **rfMRI full corr ICA100 edge 1390 link 41-51** | 0.02 | 15863 |
| **SurfArea Desikan rostralmiddlefrontal.lh** | 0.02 | 17127 |
| **GrayVol Desikan cuneus.lh** | 0.02 | 17127 |
| **opening Morphologist STpol_right** | 0.02 | 18062 |
| **rfMRI full corr ICA100 edge 1140 link 29-35** | 0.02 | 15862 |
| **ISOVF Cerebral_peduncle_L** | 0.02 | 16527 |
| **rfMRI part corr ICA100 edge 1211 link 32-34** | 0.02 | 15864 |
| **rfMRI part corr ICA100 edge 721 link 16-32** | 0.02 | 15864 |
| **rfMRI full corr ICA100 edge 276 link 6-22** | 0.02 | 15863 |
| **GrayVol Destrieux S_parieto_occipital.lh** | 0.02 | 17127 |
| **rfMRI part corr ICA100 edge 605 link 13-36** | 0.02 | 15864 |
| **rfMRI full corr ICA100 edge 553 link 12-26** | 0.02 | 15864 |
| **rfMRI full corr ICA100 edge 932 link 22-30** | 0.02 | 15864 |

| | | |
|---|---|---|
| **rfMRI full corr ICA100 edge 519 link 11-35** | 0.02 | 15864 |
| **rfMRI part corr ICA25 edge 39 link 2-21** | 0.02 | 15864 |
| **rfMRI part corr ICA25 edge 121 link 8-10** | 0.02 | 15864 |
| **opening Morphologist SOTlatmed_right** | 0.02 | 17219 |
| **rfMRI part corr ICA25 edge 13 link 1-14** | 0.02 | 15864 |
| **opening Morphologist SCu_right** | 0.02 | 18068 |
| **rfMRI part corr ICA100 edge 353 link 7-51** | 0.02 | 15864 |
| **surface Morphologist SFinfant_left** | 0.02 | 17653 |
| **GM_thickness Morphologist SOlf_left** | 0.02 | 18084 |
| **rfMRI full corr ICA25 edge 92 link 6-8** | 0.02 | 15864 |
| **opening Morphologist STsterascant_left** | 0.02 | 17560 |
| **GrayVol Destrieux S_temporal_sup.lh** | 0.02 | 17127 |
| **rfMRI full corr ICA100 edge 88 link 2-36** | 0.02 | 15862 |
| **rfMRI part corr ICA100 edge 1086 link 27-34** | 0.02 | 15864 |
| **maxdepth Morphologist FIPrint2_left** | 0.02 | 16244 |
| **surface Morphologist FCLrasc_left** | 0.02 | 17784 |
| **rfMRI amplitude ICA100 component 9** | 0.02 | 15863 |
| **rfMRI full corr ICA100 edge 20 link 1-21** | 0.02 | 15864 |
| **rfMRI part corr ICA100 edge 600 link 13-31** | 0.02 | 15864 |
| **hull_junction_length Morphologist SOp_right** | 0.02 | 17447 |
| **rfMRI part corr ICA100 edge 404 link 8-55** | 0.02 | 15864 |
| **rfMRI part corr ICA100 edge 948 link 22-46** | 0.02 | 15864 |
| **rfMRI full corr ICA100 edge 179 link 4-24** | 0.02 | 15863 |
| **ThickAvg Destrieux S_calcarine.lh** | 0.02 | 17127 |
| **rfMRI part corr ICA100 edge 758 link 17-31** | 0.02 | 15864 |
| **surface Morphologist SOTlatant_left** | 0.02 | 18076 |
| **meandepth Morphologist FIPPoCinf_right** | 0.02 | 18073 |
| **rfMRI part corr ICA100 edge 1057 link 26-33** | 0.02 | 15864 |
| **rfMRI part corr ICA100 edge 74 link 2-22** | 0.02 | 15864 |
| **rfMRI full corr ICA100 edge 529 link 11-45** | 0.02 | 15864 |
| **rfMRI full corr ICA100 edge 1139 link 29-34** | 0.02 | 15864 |
| **rfMRI full corr ICA100 edge 573 link 12-46** | 0.02 | 15864 |
| **GrayVol Destrieux G_temp_sup-Plan_tempo.rh** | 0.02 | 17127 |
| **rfMRI part corr ICA25 edge 81 link 5-12** | 0.02 | 15864 |

| | | |
|---|---|---|
| **hull_junction_length Morphologist STpol_right** | 0.02 | 18062 |
| **GrayVol Destrieux S_subparietal.rh** | 0.02 | 17126 |
| **GrayVol Destrieux G_front_middle.lh** | 0.02 | 17127 |
| **rfMRI part corr ICA100 edge 741 link 16-52** | 0.02 | 15864 |
| **rfMRI full corr ICA100 edge 997 link 24-32** | 0.02 | 15864 |
| **GrayVol Desikan rostralmiddlefrontal.lh** | 0.02 | 17127 |
| **rfMRI full corr ICA100 edge 1154 link 29-49** | 0.02 | 15857 |
| **rfMRI part corr ICA100 edge 683 link 15-33** | 0.02 | 15864 |
| **rfMRI full corr ICA100 edge 1477 link 51-53** | 0.02 | 15864 |
| **rfMRI full corr ICA100 edge 451 link 10-11** | 0.02 | 15864 |
| **opening Morphologist STs_right** | 0.02 | 18100 |
| **rfMRI full corr ICA100 edge 108 link 3-4** | 0.02 | 15864 |
| **rfMRI full corr ICA25 edge 79 link 5-10** | 0.02 | 15864 |
| **rfMRI full corr ICA100 edge 1137 link 29-32** | 0.02 | 15864 |
| **rfMRI part corr ICA100 edge 1157 link 29-52** | 0.02 | 15864 |
| **rfMRI part corr ICA100 edge 879 link 20-44** | 0.02 | 15864 |
| **rfMRI part corr ICA100 edge 1357 link 39-47** | 0.02 | 15864 |
| **hull_junction_length Morphologist SC_left** | 0.02 | 18101 |
| **rfMRI part corr ICA100 edge 693 link 15-43** | 0.02 | 15864 |
| **GrayVol Destrieux G_oc-temp_med-Parahip.lh** | 0.02 | 17127 |
| **GrayVol Destrieux S_postcentral.rh** | 0.02 | 17127 |
| **rfMRI full corr ICA100 edge 1051 link 26-27** | 0.02 | 15864 |
| **rfMRI part corr ICA100 edge 663 link 14-53** | 0.02 | 15864 |
| **rfMRI full corr ICA100 edge 1092 link 27-40** | 0.02 | 15860 |
| **rfMRI part corr ICA100 edge 391 link 8-42** | 0.02 | 15864 |
| **meandepth Morphologist SC_left** | 0.02 | 18101 |
| **rfMRI full corr ICA100 edge 801 link 18-37** | 0.02 | 15863 |
| **meandepth Morphologist SPeCmedian_right** | 0.02 | 17071 |
| **rfMRI full corr ICA100 edge 538 link 11-54** | 0.02 | 15864 |
| **GrayVol Desikan insula.lh** | 0.02 | 17127 |
| **ThickAvg Desikan precuneus.lh** | 0.03 | 17127 |
| **SurfArea Destrieux S_cingul-Marginalis.lh** | 0.03 | 17127 |
| **rfMRI full corr ICA100 edge 791 link 18-27** | 0.03 | 15863 |
| **rfMRI part corr ICA100 edge 127 link 3-23** | 0.03 | 15864 |

| | | |
|---|---|---|
| **rfMRI full corr ICA100 edge 399 link 8-50** | 0.03 | 15864 |
| **rfMRI full corr ICA100 edge 1035 link 25-40** | 0.03 | 15864 |
| **rfMRI part corr ICA100 edge 1467 link 49-52** | 0.03 | 15864 |
| **SurfArea Desikan insula.lh** | 0.03 | 17127 |
| **FA Uncinate_fasciculus_L** | 0.03 | 16541 |
| **rfMRI full corr ICA100 edge 378 link 8-29** | 0.03 | 15864 |
| **rfMRI part corr ICA100 edge 153 link 3-49** | 0.03 | 15864 |
| **rfMRI part corr ICA100 edge 66 link 2-14** | 0.03 | 15864 |
| **ThickAvg Destrieux S_circular_insula_ant.lh** | 0.03 | 17126 |
| **FA Anterior_limb_of_internal_capsule_R** | 0.03 | 16541 |
| **rfMRI full corr ICA100 edge 637 link 14-27** | 0.03 | 15859 |
| **SurfArea Destrieux G_parietal_sup.lh** | 0.03 | 17127 |
| **rfMRI full corr ICA25 edge 185 link 14-17** | 0.03 | 15864 |
| **rfMRI part corr ICA100 edge 1002 link 24-37** | 0.03 | 15864 |
| **rfMRI full corr ICA100 edge 1155 link 29-50** | 0.03 | 15863 |
| **Right-Amygdala** | 0.03 | 17127 |
| **rfMRI full corr ICA100 edge 1003 link 24-38** | 0.03 | 15864 |
| **rfMRI full corr ICA100 edge 517 link 11-33** | 0.03 | 15863 |
| **rfMRI part corr ICA100 edge 75 link 2-23** | 0.03 | 15864 |
| **rfMRI full corr ICA100 edge 798 link 18-34** | 0.03 | 15864 |
| **rfMRI part corr ICA100 edge 276 link 6-22** | 0.03 | 15864 |
| **ISOVF Retrolenticular_part_of_internal_capsule_L** | 0.03 | 16527 |
| **meandepth Morphologist FColl_right** | 0.03 | 18100 |
| **rfMRI full corr ICA100 edge 830 link 19-30** | 0.03 | 15864 |
| **maxdepth Morphologist STs_right** | 0.03 | 18100 |
| **rfMRI part corr ICA100 edge 179 link 4-24** | 0.03 | 15864 |
| **opening Morphologist SCsylvian_right** | 0.03 | 14992 |
| **rfMRI part corr ICA100 edge 1366 link 40-41** | 0.03 | 15864 |
| **GrayVol Destrieux S_cingul-Marginalis.lh** | 0.03 | 17127 |
| **rfMRI full corr ICA100 edge 775 link 17-48** | 0.03 | 15864 |
| **rfMRI part corr ICA100 edge 498 link 11-14** | 0.03 | 15864 |
| **rfMRI part corr ICA100 edge 763 link 17-36** | 0.03 | 15864 |
| **rfMRI full corr ICA100 edge 1165 link 30-35** | 0.03 | 15863 |
| **rfMRI part corr ICA100 edge 253 link 5-48** | 0.03 | 15864 |

| | | |
|---|---|---|
| **SurfArea Desikan parahippocampal.rh** | 0.03 | 17127 |
| **rfMRI part corr ICA100 edge 1355 link 39-45** | 0.03 | 15864 |
| **SurfArea Destrieux G_precuneus.rh** | 0.03 | 17127 |
| **meandepth Morphologist FIPrint1_right** | 0.03 | 17025 |
| **rfMRI full corr ICA25 edge 137 link 9-14** | 0.03 | 15864 |
| **SurfArea Destrieux G_front_sup.rh** | 0.03 | 17127 |
| **rfMRI part corr ICA100 edge 1163 link 30-33** | 0.03 | 15864 |
| **rfMRI full corr ICA100 edge 332 link 7-30** | 0.03 | 15864 |
| **rfMRI full corr ICA100 edge 1083 link 27-31** | 0.03 | 15864 |
| **OD External_capsule_R** | 0.03 | 16541 |
| **rfMRI full corr ICA100 edge 1462 link 48-53** | 0.03 | 15864 |
| **SurfArea Desikan inferiorparietal.lh** | 0.03 | 17127 |
| **rfMRI full corr ICA100 edge 777 link 17-50** | 0.03 | 15864 |
| **hull_junction_length Morphologist SPaint_right** | 0.03 | 18007 |
| **rfMRI full corr ICA100 edge 1454 link 47-52** | 0.03 | 15864 |
| **rfMRI full corr ICA100 edge 1452 link 47-50** | 0.03 | 15864 |
| **rfMRI full corr ICA100 edge 1440 link 45-55** | 0.03 | 15864 |
| **rfMRI full corr ICA25 edge 131 link 8-20** | 0.03 | 15864 |
| **rfMRI amplitude ICA100 component 8** | 0.03 | 15863 |
| **rfMRI full corr ICA100 edge 814 link 18-50** | 0.03 | 15864 |
| **maxdepth Morphologist SLiant_left** | 0.03 | 17423 |
| **rfMRI amplitude ICA25 component 17** | 0.03 | 15863 |
| **meandepth Morphologist SpC_right** | 0.03 | 16302 |
| **SurfArea Desikan caudalmiddlefrontal.lh** | 0.03 | 17127 |
| **rfMRI part corr ICA100 edge 790 link 18-26** | 0.03 | 15864 |
| **SurfArea Desikan parstriangularis.rh** | 0.03 | 17127 |
| **rfMRI full corr ICA100 edge 338 link 7-36** | 0.03 | 15860 |
| **rfMRI full corr ICA100 edge 816 link 18-52** | 0.03 | 15864 |
| **rfMRI part corr ICA100 edge 4 link 1-5** | 0.03 | 15864 |
| **maxdepth Morphologist FIP_left** | 0.03 | 18098 |
| **GM_thickness Morphologist SPaint_left** | 0.03 | 17998 |
| **rfMRI full corr ICA100 edge 546 link 12-19** | 0.03 | 15864 |
| **rfMRI part corr ICA100 edge 579 link 12-52** | 0.03 | 15864 |
| **ThickAvg Destrieux S_oc_middle_and_Lunatus.rh** | 0.03 | 17127 |

| | | |
|---|---|---|
| **SurfArea Destrieux G_cingul-Post-ventral.rh** | 0.03 | 17127 |
| **meandepth Morphologist FIPrint2_left** | 0.03 | 16204 |
| **maxdepth Morphologist SPeCmedian_right** | 0.03 | 17074 |
| **rfMRI full corr ICA100 edge 1355 link 39-45** | 0.03 | 15864 |
| **rfMRI part corr ICA25 edge 102 link 6-18** | 0.03 | 15864 |
| **MD Tapetum_L** | 0.03 | 16541 |
| **rfMRI part corr ICA100 edge 1333 link 38-39** | 0.03 | 15864 |
| **ThickAvg Destrieux S_oc-temp_lat.rh** | 0.03 | 17127 |
| **rfMRI full corr ICA100 edge 606 link 13-37** | 0.03 | 15864 |
| **ThickAvg Destrieux G_temporal_inf.lh** | 0.03 | 17127 |
| **rfMRI part corr ICA100 edge 964 link 23-30** | 0.03 | 15864 |
| **SurfArea Destrieux G_and_S_occipital_inf.rh** | 0.03 | 17127 |
| **SurfArea Destrieux G_temp_sup-Lateral.rh** | 0.03 | 17127 |
| **rfMRI full corr ICA100 edge 880 link 20-45** | 0.03 | 15864 |
| **rfMRI part corr ICA100 edge 147 link 3-43** | 0.03 | 15864 |
| **MO Superior_corona_radiata_R** | 0.03 | 16541 |
| **rfMRI part corr ICA25 edge 180 link 13-19** | 0.03 | 15864 |
| **rfMRI full corr ICA100 edge 1000 link 24-35** | 0.03 | 15862 |
| **OD Anterior_limb_of_internal_capsule_L** | 0.03 | 16541 |
| **rfMRI part corr ICA100 edge 787 link 18-23** | 0.03 | 15864 |
| **rfMRI part corr ICA100 edge 1300 link 36-41** | 0.03 | 15864 |
| **rfMRI part corr ICA100 edge 381 link 8-32** | 0.03 | 15864 |
| **Right-Cerebellum-White-Matter** | 0.03 | 17127 |
| **maxdepth Morphologist SCu_right** | 0.03 | 18068 |
| **SurfArea Destrieux S_circular_insula_inf.rh** | 0.03 | 17125 |
| **GrayVol Destrieux G_oc-temp_med-Lingual.rh** | 0.03 | 17127 |
| **rfMRI full corr ICA100 edge 1172 link 30-42** | 0.03 | 15864 |
| **ThickAvg Desikan posteriorcingulate.lh** | 0.03 | 17126 |
| **GrayVol Desikan precuneus.rh** | 0.03 | 17127 |
| **rfMRI part corr ICA25 edge 92 link 6-8** | 0.03 | 15864 |
| **rfMRI part corr ICA100 edge 431 link 9-36** | 0.03 | 15864 |
| **rfMRI full corr ICA100 edge 595 link 13-26** | 0.03 | 15864 |
| **GrayVol Desikan parstriangularis.lh** | 0.03 | 17127 |
| **rfMRI full corr ICA100 edge 609 link 13-40** | 0.04 | 15863 |

| | | |
|---|---|---|
| **OD Anterior_limb_of_internal_capsule_R** | 0.04 | 16541 |
| **OD Tapetum_R** | 0.04 | 16541 |
| **GrayVol Desikan isthmuscingulate.lh** | 0.04 | 17126 |
| **OD Superior_fronto-occipital_fasciculus-part_of_a** | 0.04 | 16541 |
| **rfMRI part corr ICA100 edge 1417 link 43-53** | 0.04 | 15864 |
| **rfMRI full corr ICA100 edge 1037 link 25-42** | 0.04 | 15863 |
| **ThickAvg Destrieux G_and_S_cingul-Mid-Ant.lh** | 0.04 | 17127 |
| **SurfArea Destrieux S_parieto_occipital.rh** | 0.04 | 17127 |
| **meandepth Morphologist FCLrscpost_left** | 0.04 | 13744 |
| **ThickAvg Destrieux G_cingul-Post-ventral.lh** | 0.04 | 17127 |
| **rfMRI part corr ICA100 edge 1231 link 32-54** | 0.04 | 15864 |
| **SurfArea Destrieux G_front_middle.rh** | 0.04 | 17127 |
| **rfMRI part corr ICA100 edge 517 link 11-33** | 0.04 | 15864 |
| **rfMRI full corr ICA100 edge 591 link 13-22** | 0.04 | 15864 |
| **meandepth Morphologist SFpolairetr_left** | 0.04 | 18062 |
| **rfMRI part corr ICA100 edge 628 link 14-18** | 0.04 | 15864 |
| **rfMRI full corr ICA100 edge 806 link 18-42** | 0.04 | 15864 |
| **rfMRI full corr ICA100 edge 324 link 7-22** | 0.04 | 15864 |
| **rfMRI full corr ICA100 edge 1122 link 28-43** | 0.04 | 15863 |
| **rfMRI part corr ICA100 edge 1067 link 26-43** | 0.04 | 15864 |
| **rfMRI full corr ICA100 edge 1359 link 39-49** | 0.04 | 15864 |
| **rfMRI part corr ICA100 edge 535 link 11-51** | 0.04 | 15864 |
| **maxdepth Morphologist SPeCmedian_left** | 0.04 | 16802 |
| **rfMRI full corr ICA100 edge 1399 link 42-47** | 0.04 | 15864 |
| **rfMRI part corr ICA100 edge 1015 link 24-50** | 0.04 | 15864 |
| **rfMRI part corr ICA100 edge 530 link 11-46** | 0.04 | 15864 |
| **rfMRI part corr ICA100 edge 531 link 11-47** | 0.04 | 15864 |
| **ThickAvg Destrieux G_parietal_sup.lh** | 0.04 | 17127 |
| **surface Morphologist FColl_right** | 0.04 | 18100 |
| **SurfArea Destrieux G_parietal_sup.rh** | 0.04 | 17127 |
| **SurfArea Destrieux G_front_inf-Triangul.lh** | 0.04 | 17127 |
| **rfMRI part corr ICA100 edge 773 link 17-46** | 0.04 | 15864 |
| **rfMRI full corr ICA100 edge 1471 link 50-51** | 0.04 | 15864 |
| **rfMRI full corr ICA100 edge 1103 link 27-51** | 0.04 | 15864 |

| | | |
|---|---|---|
| **maxdepth Morphologist SpC_right** | 0.04 | 16309 |
| **rfMRI part corr ICA100 edge 1165 link 30-35** | 0.04 | 15864 |
| **rfMRI full corr ICA25 edge 135 link 9-12** | 0.04 | 15864 |
| **SurfArea Destrieux S_circular_insula_ant.rh** | 0.04 | 17125 |
| **rfMRI full corr ICA100 edge 39 link 1-40** | 0.04 | 15861 |
| **rfMRI part corr ICA25 edge 94 link 6-10** | 0.04 | 15864 |
| **SurfArea Desikan superiorfrontal.rh** | 0.04 | 17127 |
| **MO Posterior_limb_of_internal_capsule_R** | 0.04 | 16541 |
| **rfMRI full corr ICA25 edge 13 link 1-14** | 0.04 | 15864 |
| **GrayVol Desikan lingual.rh** | 0.04 | 17127 |
| **OD Fornix-cres-Stria_terminalis-not_resolved_wit** | 0.04 | 16541 |
| **hull_junction_length Morphologist FCalant-ScCal_** | 0.04 | 18100 |
| **rfMRI part corr ICA100 edge 1480 link 52-53** | 0.04 | 15864 |
| **rfMRI part corr ICA25 edge 18 link 1-19** | 0.04 | 15864 |
| **rfMRI full corr ICA100 edge 507 link 11-23** | 0.04 | 15864 |
| **rfMRI part corr ICA100 edge 932 link 22-30** | 0.04 | 15864 |
| **rfMRI full corr ICA100 edge 1131 link 28-52** | 0.04 | 15861 |
| **rfMRI full corr ICA100 edge 75 link 2-23** | 0.04 | 15864 |
| **rfMRI full corr ICA100 edge 886 link 20-51** | 0.04 | 15863 |
| **OD Superior_corona_radiata_L** | 0.04 | 16541 |
| **GrayVol Desikan paracentral.lh** | 0.04 | 17127 |
| **rfMRI full corr ICA25 edge 83 link 5-14** | 0.04 | 15864 |
| **rfMRI full corr ICA25 edge 120 link 8-9** | 0.04 | 15864 |
| **ThickAvg Destrieux Pole_temporal.rh** | 0.04 | 17126 |
| **rfMRI part corr ICA100 edge 393 link 8-44** | 0.04 | 15864 |
| **GrayVol Destrieux G_occipital_middle.rh** | 0.04 | 17127 |
| **rfMRI part corr ICA100 edge 1376 link 40-51** | 0.04 | 15864 |
| **rfMRI full corr ICA100 edge 592 link 13-23** | 0.04 | 15864 |
| **rfMRI full corr ICA100 edge 1281 link 35-41** | 0.04 | 15864 |
| **rfMRI full corr ICA100 edge 1101 link 27-49** | 0.04 | 15864 |
| **rfMRI part corr ICA100 edge 546 link 12-19** | 0.04 | 15864 |
| **MO External_capsule_L** | 0.04 | 16541 |
| **OD Uncinate_fasciculus_R** | 0.04 | 16541 |
| **maxdepth Morphologist SOTlatint_left** | 0.04 | 16594 |

| | | |
|---|---|---|
| **rfMRI full corr ICA25 edge 94 link 6-10** | 0.04 | 15864 |
| **rfMRI full corr ICA25 edge 50 link 3-14** | 0.04 | 15864 |
| **rfMRI full corr ICA100 edge 1273 link 34-53** | 0.04 | 15864 |
| **rfMRI full corr ICA100 edge 1305 link 36-46** | 0.04 | 15864 |
| **surface Morphologist SOp_left** | 0.04 | 15273 |
| **rfMRI part corr ICA100 edge 374 link 8-25** | 0.04 | 15864 |
| **SurfArea Desikan frontalpole.lh** | 0.04 | 17127 |
| **ThickAvg Destrieux S_interm_prim-Jensen.rh** | 0.04 | 17127 |
| **rfMRI full corr ICA100 edge 23 link 1-24** | 0.04 | 15864 |
| **ICVF Tapetum_L** | 0.04 | 16540 |
| **rfMRI full corr ICA100 edge 1433 link 45-48** | 0.04 | 15864 |
| **ISOVF Fornix-cres-Stria_terminalis-not_resolved_** | 0.04 | 16527 |
| **ICVF Superior_fronto-occipital_fasciculus-part_of** | 0.04 | 16540 |
| **rfMRI part corr ICA100 edge 200 link 4-45** | 0.04 | 15864 |
| **ThickAvg Destrieux G_and_S_paracentral.rh** | 0.04 | 17127 |
| **rfMRI part corr ICA100 edge 563 link 12-36** | 0.04 | 15864 |
| **rfMRI full corr ICA100 edge 860 link 20-25** | 0.04 | 15864 |
| **MD Retrolenticular_part_of_internal_capsule_L** | 0.04 | 16541 |
| **rfMRI full corr ICA100 edge 1246 link 33-47** | 0.04 | 15863 |
| **rfMRI full corr ICA100 edge 1033 link 25-38** | 0.04 | 15864 |
| **rfMRI part corr ICA100 edge 648 link 14-38** | 0.04 | 15864 |
| **rfMRI full corr ICA100 edge 1332 link 37-55** | 0.04 | 15864 |
| **rfMRI full corr ICA100 edge 78 link 2-26** | 0.04 | 15864 |
| **maxdepth Morphologist FCLrscpost_left** | 0.05 | 13973 |
| **maxdepth Morphologist FIPPoCinf_right** | 0.05 | 18073 |
| **rfMRI full corr ICA25 edge 181 link 13-20** | 0.05 | 15864 |
| **rfMRI full corr ICA100 edge 126 link 3-22** | 0.05 | 15864 |
| **rfMRI full corr ICA100 edge 1149 link 29-44** | 0.05 | 15864 |
| **GrayVol Desikan precuneus.lh** | 0.05 | 17127 |
| **SurfArea Desikan bankssts.rh** | 0.05 | 17127 |
| **GM_thickness Morphologist SPoCsup_right** | 0.05 | 17834 |
| **rfMRI part corr ICA100 edge 880 link 20-45** | 0.05 | 15864 |
| **rfMRI full corr ICA100 edge 256 link 5-51** | 0.05 | 15863 |
| **rfMRI part corr ICA100 edge 1389 link 41-50** | 0.05 | 15864 |

| | | |
|---|---|---|
| **ThickAvg Destrieux G_front_inf-Orbital.lh** | 0.05 | 17125 |
| **rfMRI amplitude ICA100 component 50** | 0.05 | 15863 |
| **MO Cingulum-hippocampus-R** | 0.05 | 16541 |
| **rfMRI full corr ICA100 edge 616 link 13-47** | 0.05 | 15864 |
| **rfMRI full corr ICA100 edge 389 link 8-40** | 0.05 | 15864 |
| **rfMRI full corr ICA100 edge 1316 link 37-39** | 0.05 | 15864 |
| **SurfArea Desikan superiortemporal.lh** | 0.05 | 17127 |
| **ISOVF Tapetum_L** | 0.05 | 16527 |
| **rfMRI part corr ICA100 edge 749 link 17-22** | 0.05 | 15864 |
| **SurfArea Destrieux S_precentral-inf-part.rh** | 0.05 | 17127 |
| **MD Posterior_limb_of_internal_capsule_L** | 0.05 | 16541 |
| **rfMRI part corr ICA25 edge 196 link 16-17** | 0.05 | 15864 |
| **rfMRI full corr ICA100 edge 53 link 1-54** | 0.05 | 15864 |
| **ThickAvg Desikan inferiortemporal.lh** | 0.05 | 17127 |
| **surface Morphologist SOTlatpost_left** | 0.05 | 18062 |
| **rfMRI full corr ICA25 edge 60 link 4-7** | 0.05 | 15864 |
| **maxdepth Morphologist SOTlatint_right** | 0.05 | 15621 |
| **rfMRI full corr ICA100 edge 203 link 4-48** | 0.05 | 15864 |
| **ThickAvg Desikan lingual.lh** | 0.05 | 17127 |
| **rfMRI part corr ICA100 edge 1115 link 28-36** | 0.05 | 15864 |
| **surface Morphologist SFinter_left** | 0.05 | 18101 |
| **maxdepth Morphologist SC_left** | 0.05 | 18101 |
| **rfMRI full corr ICA100 edge 87 link 2-35** | 0.05 | 15864 |
| **SurfArea Destrieux S_orbital_med-olfact.rh** | 0.05 | 17125 |
| **SurfArea Destrieux S_postcentral.rh** | 0.05 | 17127 |
| **rfMRI full corr ICA100 edge 1450 link 47-48** | 0.05 | 15864 |
| **rfMRI part corr ICA100 edge 987 link 23-53** | 0.05 | 15864 |
| **rfMRI part corr ICA100 edge 666 link 15-16** | 0.05 | 15864 |
| **rfMRI part corr ICA100 edge 616 link 13-47** | 0.05 | 15864 |
| **rfMRI full corr ICA100 edge 1091 link 27-39** | 0.05 | 15864 |
| **rfMRI part corr ICA100 edge 756 link 17-29** | 0.05 | 15864 |
| **rfMRI part corr ICA100 edge 694 link 15-44** | 0.05 | 15864 |
| **rfMRI full corr ICA100 edge 35 link 1-36** | 0.05 | 15864 |
| **GrayVol Desikan rostralanteriorcingulate.rh** | 0.05 | 17127 |

| | | |
|---|---|---|
| **rfMRI part corr ICA100 edge 289 link 6-35** | 0.05 | 15864 |
| **surface Morphologist STsterascpost_left** | 0.05 | 17904 |
| **rfMRI part corr ICA100 edge 538 link 11-54** | 0.05 | 15864 |
| **rfMRI full corr ICA100 edge 721 link 16-32** | 0.05 | 15864 |
| **rfMRI part corr ICA100 edge 478 link 10-38** | 0.05 | 15864 |
| **rfMRI full corr ICA25 edge 132 link 8-21** | 0.05 | 15864 |
| **ThickAvg Destrieux G_subcallosal.lh** | 0.05 | 17125 |
| **rfMRI full corr ICA100 edge 365 link 8-16** | 0.05 | 15864 |
| **rfMRI part corr ICA100 edge 1080 link 27-28** | 0.05 | 15864 |
| **rfMRI full corr ICA100 edge 635 link 14-25** | 0.05 | 15861 |
| **rfMRI part corr ICA100 edge 1303 link 36-44** | 0.05 | 15864 |
| **rfMRI full corr ICA100 edge 253 link 5-48** | 0.05 | 15864 |
| **hull_junction_length Morphologist FIPPoCinf_righ** | 0.05 | 18073 |
| **rfMRI full corr ICA100 edge 648 link 14-38** | 0.05 | 15864 |
| **GM_thickness Morphologist SLiant_right** | 0.05 | 17214 |
| **rfMRI part corr ICA100 edge 91 link 2-39** | 0.05 | 15864 |
| **rfMRI part corr ICA100 edge 606 link 13-37** | 0.05 | 15864 |
| **rfMRI full corr ICA100 edge 1232 link 32-55** | 0.05 | 15864 |
| **rfMRI full corr ICA100 edge 1460 link 48-51** | 0.05 | 15864 |
| **GM_thickness Morphologist SOlf_right** | 0.05 | 18083 |
| **rfMRI full corr ICA25 edge 144 link 9-21** | 0.05 | 15864 |
| **GM_thickness Morphologist SOp_right** | 0.05 | 17438 |
| **rfMRI part corr ICA100 edge 302 link 6-48** | 0.05 | 15864 |
| **rfMRI part corr ICA100 edge 231 link 5-26** | 0.05 | 15864 |
| **rfMRI part corr ICA100 edge 485 link 10-45** | 0.05 | 15864 |
| **GrayVol Destrieux S_interm_prim-Jensen.rh** | 0.05 | 17127 |
| **rfMRI part corr ICA25 edge 7 link 1-8** | 0.05 | 15864 |
| **rfMRI part corr ICA100 edge 1281 link 35-41** | 0.05 | 15864 |
| **rfMRI part corr ICA100 edge 793 link 18-29** | 0.05 | 15864 |
| **rfMRI full corr ICA100 edge 1183 link 30-53** | 0.05 | 15863 |
| **rfMRI full corr ICA100 edge 1194 link 31-40** | 0.05 | 15864 |
| **rfMRI full corr ICA100 edge 896 link 21-27** | 0.05 | 15864 |
| **rfMRI part corr ICA100 edge 63 link 2-11** | 0.05 | 15864 |
| **rfMRI part corr ICA100 edge 1116 link 28-37** | 0.05 | 15864 |

| | | |
|---|---|---|
| **GM_thickness Morphologist SRinf_left** | 0.05 | 16688 |
| **rfMRI full corr ICA100 edge 914 link 21-45** | 0.06 | 15862 |
| **rfMRI part corr ICA100 edge 47 link 1-48** | 0.06 | 15864 |
| **rfMRI part corr ICA100 edge 1233 link 33-34** | 0.06 | 15864 |
| **MD Tapetum_R** | 0.06 | 16541 |
| **rfMRI full corr ICA100 edge 1053 link 26-29** | 0.06 | 15863 |
| **rfMRI full corr ICA100 edge 1480 link 52-53** | 0.06 | 15864 |
| **rfMRI part corr ICA100 edge 650 link 14-40** | 0.06 | 15864 |
| **rfMRI amplitude ICA100 component 5** | 0.06 | 15863 |
| **rfMRI full corr ICA25 edge 188 link 14-20** | 0.06 | 15864 |
| **SurfArea Desikan isthmuscingulate.lh** | 0.06 | 17126 |
| **hull_junction_length Morphologist SFint_left** | 0.06 | 18101 |
| **ThickAvg Desikan inferiorparietal.lh** | 0.06 | 17127 |
| **ThickAvg Destrieux G_occipital_sup.rh** | 0.06 | 17127 |
| **rfMRI part corr ICA100 edge 615 link 13-46** | 0.06 | 15864 |
| **surface Morphologist SPeCmedian_left** | 0.06 | 16802 |
| **rfMRI part corr ICA25 edge 71 link 4-18** | 0.06 | 15864 |
| **GrayVol Desikan middletemporal.rh** | 0.06 | 17127 |
| **maxdepth Morphologist FIPPoCinf_left** | 0.06 | 18079 |
| **rfMRI part corr ICA100 edge 304 link 6-50** | 0.06 | 15864 |
| **rfMRI part corr ICA100 edge 306 link 6-52** | 0.06 | 15864 |
| **SurfArea Destrieux S_subparietal.rh** | 0.06 | 17126 |
| **GM_thickness Morphologist STiant_right** | 0.06 | 18092 |
| **rfMRI part corr ICA25 edge 156 link 11-12** | 0.06 | 15864 |
| **rfMRI part corr ICA100 edge 654 link 14-44** | 0.06 | 15864 |
| **rfMRI full corr ICA100 edge 438 link 9-43** | 0.06 | 15864 |
| **rfMRI full corr ICA100 edge 600 link 13-31** | 0.06 | 15864 |
| **GrayVol Desikan entorhinal.lh** | 0.06 | 17127 |
| **rfMRI full corr ICA100 edge 398 link 8-49** | 0.06 | 15864 |
| **rfMRI full corr ICA100 edge 63 link 2-11** | 0.06 | 15864 |
| **GrayVol Desikan supramarginal.lh** | 0.06 | 17127 |
| **maxdepth Morphologist FIPrint1_right** | 0.06 | 17034 |
| **SurfArea Desikan superiorparietal.lh** | 0.06 | 17127 |
| **rfMRI part corr ICA25 edge 116 link 7-18** | 0.06 | 15864 |

| | | |
|---|---|---|
| **OD External_capsule_L** | 0.06 | 16541 |
| **hull_junction_length Morphologist FColl_left** | 0.06 | 18101 |
| **rfMRI full corr ICA25 edge 77 link 5-8** | 0.06 | 15864 |
| **rfMRI full corr ICA100 edge 36 link 1-37** | 0.06 | 15864 |
| **maxdepth Morphologist SPeCinf_right** | 0.06 | 16608 |
| **rfMRI part corr ICA100 edge 126 link 3-22** | 0.06 | 15864 |
| **GrayVol Destrieux G_and_S_paracentral.lh** | 0.06 | 17127 |
| **rfMRI full corr ICA100 edge 171 link 4-16** | 0.06 | 15864 |
| **GrayVol Destrieux G_front_middle.rh** | 0.06 | 17127 |
| **rfMRI full corr ICA100 edge 184 link 4-29** | 0.06 | 15864 |
| **GrayVol Destrieux G_front_inf-Opercular.lh** | 0.06 | 17127 |
| **maxdepth Morphologist SPeCmarginal_left** | 0.06 | 15613 |
| **rfMRI full corr ICA100 edge 861 link 20-26** | 0.06 | 15864 |
| **rfMRI full corr ICA100 edge 1102 link 27-50** | 0.06 | 15864 |
| **rfMRI full corr ICA100 edge 1467 link 49-52** | 0.06 | 15864 |
| **MO Cingulum-hippocampus-L** | 0.06 | 16541 |
| **rfMRI full corr ICA100 edge 859 link 20-24** | 0.06 | 15864 |
| **rfMRI part corr ICA100 edge 1172 link 30-42** | 0.06 | 15864 |
| **ThickAvg Destrieux G_temp_sup-Plan_tempo.rh** | 0.06 | 17127 |
| **rfMRI part corr ICA100 edge 385 link 8-36** | 0.06 | 15864 |
| **rfMRI part corr ICA100 edge 732 link 16-43** | 0.06 | 15864 |
| **rfMRI full corr ICA100 edge 1175 link 30-45** | 0.06 | 15864 |
| **rfMRI full corr ICA100 edge 807 link 18-43** | 0.06 | 15864 |
| **surface Morphologist FIPrint1_right** | 0.06 | 17034 |
| **SurfArea Desikan inferiortemporal.lh** | 0.06 | 17127 |
| **rfMRI full corr ICA100 edge 622 link 13-53** | 0.06 | 15864 |
| **rfMRI full corr ICA100 edge 1387 link 41-48** | 0.06 | 15864 |
| **ThickAvg Destrieux G_oc-temp_med-Lingual.lh** | 0.06 | 17127 |
| **rfMRI part corr ICA100 edge 505 link 11-21** | 0.06 | 15864 |
| **rfMRI part corr ICA25 edge 127 link 8-16** | 0.06 | 15864 |
| **rfMRI full corr ICA100 edge 45 link 1-46** | 0.06 | 15862 |
| **meandepth Morphologist SPat_right** | 0.06 | 16616 |
| **Left-Accumbens-area** | 0.06 | 17127 |
| **surface Morphologist SFpolairetr_right** | 0.06 | 18086 |

| | | |
|---|---|---|
| **MO Fornix-cres-Stria_terminalis-not_resolved_wit** | 0.06 | 16541 |
| **rfMRI part corr ICA100 edge 81 link 2-29** | 0.06 | 15864 |
| **rfMRI full corr ICA25 edge 30 link 2-12** | 0.06 | 15856 |
| **rfMRI part corr ICA100 edge 1310 link 36-51** | 0.06 | 15864 |
| **rfMRI full corr ICA100 edge 396 link 8-47** | 0.06 | 15864 |
| **rfMRI full corr ICA100 edge 198 link 4-43** | 0.06 | 15864 |
| **rfMRI full corr ICA100 edge 757 link 17-30** | 0.06 | 15864 |
| **ThickAvg Destrieux S_oc_sup_and_transversal.rh** | 0.06 | 17127 |
| **rfMRI full corr ICA25 edge 82 link 5-13** | 0.06 | 15864 |
| **rfMRI amplitude ICA100 component 27** | 0.06 | 15863 |
| **rfMRI full corr ICA100 edge 252 link 5-47** | 0.06 | 15864 |
| **rfMRI part corr ICA100 edge 51 link 1-52** | 0.06 | 15864 |
| **rfMRI full corr ICA100 edge 1202 link 31-48** | 0.06 | 15864 |
| **rfMRI part corr ICA100 edge 676 link 15-26** | 0.06 | 15864 |
| **ThickAvg Desikan transversetemporal.lh** | 0.06 | 17126 |
| **rfMRI full corr ICA100 edge 576 link 12-49** | 0.06 | 15864 |
| **Left-Hippocampus** | 0.06 | 17127 |
| **rfMRI amplitude ICA100 component 41** | 0.06 | 15863 |
| **rfMRI part corr ICA25 edge 169 link 12-16** | 0.06 | 15864 |
| **rfMRI full corr ICA100 edge 483 link 10-43** | 0.06 | 15864 |
| **MO Medial_lemniscus_R** | 0.06 | 16541 |
| **rfMRI full corr ICA100 edge 117 link 3-13** | 0.06 | 15863 |
| **rfMRI full corr ICA100 edge 1075 link 26-51** | 0.06 | 15864 |
| **opening Morphologist SFinfant_left** | 0.07 | 17653 |
| **rfMRI part corr ICA100 edge 1313 link 36-54** | 0.07 | 15864 |
| **GrayVol Destrieux S_collat_transv_ant.lh** | 0.07 | 17127 |
| **rfMRI full corr ICA25 edge 180 link 13-19** | 0.07 | 15864 |
| **rfMRI full corr ICA100 edge 1307 link 36-48** | 0.07 | 15864 |
| **rfMRI part corr ICA100 edge 1103 link 27-51** | 0.07 | 15864 |
| **SurfArea Desikan superiorfrontal.lh** | 0.07 | 17127 |
| **rfMRI part corr ICA100 edge 1061 link 26-37** | 0.07 | 15864 |
| **MD External_capsule_L** | 0.07 | 16541 |
| **meandepth Morphologist FColl_left** | 0.07 | 18101 |
| **SurfArea Destrieux G_temporal_middle.lh** | 0.07 | 17127 |

| | | |
|---|---|---|
| **meandepth Morphologist SFinfant_left** | 0.07 | 17647 |
| **FA Uncinate_fasciculus_R** | 0.07 | 16541 |
| **rfMRI part corr ICA25 edge 120 link 8-9** | 0.07 | 15864 |
| **rfMRI part corr ICA25 edge 187 link 14-19** | 0.07 | 15864 |
| **SurfArea Destrieux S_oc-temp_med_and_Lingual.** | 0.07 | 17127 |
| **opening Morphologist FColl_right** | 0.07 | 18100 |
| **maxdepth Morphologist STsterascant_right** | 0.07 | 17642 |
| **ISOVF Cerebral_peduncle_R** | 0.07 | 16527 |
| **ThickAvg Desikan entorhinal.lh** | 0.07 | 17127 |
| **rfMRI part corr ICA100 edge 1394 link 41-55** | 0.07 | 15864 |
| **rfMRI full corr ICA25 edge 88 link 5-19** | 0.07 | 15864 |
| **rfMRI full corr ICA100 edge 448 link 9-53** | 0.07 | 15864 |
| **GrayVol Desikan insula.rh** | 0.07 | 17127 |
| **rfMRI part corr ICA100 edge 788 link 18-24** | 0.07 | 15864 |
| **ThickAvg Desikan pericalcarine.lh** | 0.07 | 17127 |
| **rfMRI full corr ICA100 edge 189 link 4-34** | 0.07 | 15864 |
| **meandepth Morphologist SPaint_right** | 0.07 | 18007 |
| **rfMRI full corr ICA100 edge 1304 link 36-45** | 0.07 | 15864 |
| **SurfArea Destrieux Lat_Fis-post.rh** | 0.07 | 17127 |
| **OD Corticospinal_tract_L** | 0.07 | 16541 |
| **SurfArea Destrieux G_occipital_middle.rh** | 0.07 | 17127 |
| **MD Cingulum-hippocampus-L** | 0.07 | 16541 |
| **rfMRI full corr ICA100 edge 1015 link 24-50** | 0.07 | 15864 |
| **rfMRI part corr ICA100 edge 50 link 1-51** | 0.07 | 15864 |
| **rfMRI part corr ICA100 edge 1235 link 33-36** | 0.07 | 15864 |
| **rfMRI part corr ICA100 edge 324 link 7-22** | 0.07 | 15864 |
| **rfMRI full corr ICA100 edge 1406 link 42-54** | 0.07 | 15864 |
| **rfMRI full corr ICA100 edge 56 link 2-4** | 0.07 | 15864 |
| **rfMRI full corr ICA100 edge 44 link 1-45** | 0.07 | 15862 |
| **rfMRI part corr ICA100 edge 218 link 5-13** | 0.07 | 15864 |
| **rfMRI full corr ICA100 edge 1328 link 37-51** | 0.07 | 15864 |
| **rfMRI part corr ICA100 edge 1406 link 42-54** | 0.07 | 15864 |
| **rfMRI part corr ICA25 edge 185 link 14-17** | 0.07 | 15864 |
| **SurfArea Destrieux Pole_occipital.rh** | 0.07 | 17127 |

| | | |
|---|---|---|
| **SurfArea Destrieux G_oc-temp_med-Lingual.lh** | 0.07 | 17127 |
| **rfMRI part corr ICA100 edge 17 link 1-18** | 0.07 | 15864 |
| **rfMRI full corr ICA25 edge 102 link 6-18** | 0.07 | 15862 |
| **SurfArea Destrieux S_precentral-sup-part.rh** | 0.07 | 17127 |
| **rfMRI part corr ICA25 edge 77 link 5-8** | 0.07 | 15864 |
| **rfMRI full corr ICA100 edge 81 link 2-29** | 0.07 | 15864 |
| **rfMRI part corr ICA100 edge 411 link 9-16** | 0.07 | 15864 |
| **ThickAvg Destrieux S_subparietal.lh** | 0.07 | 17125 |
| **rfMRI part corr ICA100 edge 688 link 15-38** | 0.07 | 15864 |
| **hull_junction_length Morphologist SPeCmedian_le** | 0.07 | 16802 |
| **rfMRI part corr ICA100 edge 1323 link 37-46** | 0.07 | 15864 |
| **rfMRI full corr ICA100 edge 1145 link 29-40** | 0.07 | 15864 |
| **surface Morphologist SPaint_left** | 0.07 | 18007 |
| **GrayVol Destrieux S_circular_insula_ant.rh** | 0.07 | 17125 |
| **rfMRI part corr ICA100 edge 1117 link 28-38** | 0.07 | 15864 |
| **rfMRI full corr ICA100 edge 1163 link 30-33** | 0.07 | 15864 |
| **hull_junction_length Morphologist SFinter_right** | 0.07 | 18099 |
| **opening Morphologist SFinfant_right** | 0.07 | 18042 |
| **rfMRI part corr ICA100 edge 491 link 10-51** | 0.07 | 15864 |
| **rfMRI full corr ICA100 edge 1115 link 28-36** | 0.07 | 15864 |
| **rfMRI part corr ICA100 edge 595 link 13-26** | 0.07 | 15864 |
| **SurfArea Desikan lingual.lh** | 0.07 | 17127 |
| **opening Morphologist SForbitaire_right** | 0.07 | 17279 |
| **rfMRI part corr ICA25 edge 135 link 9-12** | 0.07 | 15864 |
| **surface Morphologist SPaint_right** | 0.08 | 18007 |
| **rfMRI full corr ICA100 edge 162 link 4-7** | 0.08 | 15864 |
| **rfMRI full corr ICA100 edge 502 link 11-18** | 0.08 | 15863 |
| **SurfArea Destrieux G_Ins_lg_and_S_cent_ins.lh** | 0.08 | 17126 |
| **ICVF Cingulum-hippocampus-L** | 0.08 | 16540 |
| **rfMRI part corr ICA100 edge 499 link 11-15** | 0.08 | 15864 |
| **ThickAvg Destrieux G_temp_sup-G_T_transv.lh** | 0.08 | 17126 |
| **rfMRI full corr ICA100 edge 246 link 5-41** | 0.08 | 15864 |
| **rfMRI full corr ICA100 edge 215 link 5-10** | 0.08 | 15862 |
| **rfMRI part corr ICA100 edge 382 link 8-33** | 0.08 | 15864 |

| | | |
|---|---|---|
| **rfMRI part corr ICA100 edge 1398 link 42-46** | 0.08 | 15864 |
| **SurfArea Destrieux G_occipital_sup.rh** | 0.08 | 17127 |
| **surface Morphologist SLiant_right** | 0.08 | 17219 |
| **Left-Putamen** | 0.08 | 17127 |
| **rfMRI part corr ICA100 edge 445 link 9-50** | 0.08 | 15864 |
| **rfMRI part corr ICA100 edge 193 link 4-38** | 0.08 | 15864 |
| **rfMRI part corr ICA100 edge 1014 link 24-49** | 0.08 | 15864 |
| **rfMRI full corr ICA100 edge 654 link 14-44** | 0.08 | 15864 |
| **rfMRI full corr ICA100 edge 286 link 6-32** | 0.08 | 15864 |
| **rfMRI full corr ICA25 edge 139 link 9-16** | 0.08 | 15864 |
| **MD External_capsule_R** | 0.08 | 16541 |
| **rfMRI part corr ICA100 edge 398 link 8-49** | 0.08 | 15864 |
| **rfMRI part corr ICA100 edge 965 link 23-31** | 0.08 | 15864 |
| **hull_junction_length Morphologist FCLp_right** | 0.08 | 18100 |
| **rfMRI amplitude ICA25 component 9** | 0.08 | 15863 |
| **rfMRI full corr ICA100 edge 172 link 4-17** | 0.08 | 15864 |
| **ThickAvg Destrieux G_precuneus.lh** | 0.08 | 17127 |
| **rfMRI full corr ICA100 edge 1005 link 24-40** | 0.08 | 15864 |
| **hull_junction_length Morphologist STsterascpost_** | 0.08 | 17904 |
| **rfMRI full corr ICA100 edge 835 link 19-35** | 0.08 | 15864 |
| **rfMRI full corr ICA100 edge 9 link 1-10** | 0.08 | 15864 |
| **rfMRI full corr ICA100 edge 640 link 14-30** | 0.08 | 15864 |
| **rfMRI full corr ICA100 edge 1394 link 41-55** | 0.08 | 15864 |
| **rfMRI full corr ICA100 edge 42 link 1-43** | 0.08 | 15864 |
| **rfMRI full corr ICA100 edge 255 link 5-50** | 0.08 | 15860 |
| **GrayVol Desikan parahippocampal.rh** | 0.08 | 17127 |
| **rfMRI full corr ICA100 edge 1303 link 36-44** | 0.08 | 15863 |
| **ICVF Tapetum_R** | 0.08 | 16540 |
| **FA Medial_lemniscus_R** | 0.08 | 16541 |
| **rfMRI full corr ICA100 edge 1333 link 38-39** | 0.08 | 15864 |
| **rfMRI part corr ICA100 edge 1435 link 45-50** | 0.08 | 15864 |
| **hull_junction_length Morphologist SOTlatpost_left** | 0.08 | 18062 |
| **rfMRI part corr ICA100 edge 155 link 3-51** | 0.08 | 15864 |
| **rfMRI part corr ICA100 edge 752 link 17-25** | 0.08 | 15864 |

| | | |
|---|---|---|
| **SurfArea Destrieux S_temporal_transverse.lh** | 0.08 | 17126 |
| **rfMRI part corr ICA100 edge 68 link 2-16** | 0.08 | 15864 |
| **maxdepth Morphologist FCMpost_right** | 0.08 | 18100 |
| **rfMRI part corr ICA100 edge 1026 link 25-31** | 0.08 | 15864 |
| **GM_thickness Morphologist FCLrant_right** | 0.08 | 16190 |
| **rfMRI part corr ICA100 edge 284 link 6-30** | 0.08 | 15864 |
| **rfMRI full corr ICA100 edge 350 link 7-48** | 0.08 | 15864 |
| **ISOVF Sagittal_stratum-inf_longitidinal_fasci_and_** | 0.08 | 16527 |
| **rfMRI part corr ICA100 edge 352 link 7-50** | 0.08 | 15864 |
| **ThickAvg Destrieux G_pariet_inf-Supramar.lh** | 0.08 | 17127 |
| **meandepth Morphologist SPaint_left** | 0.08 | 18005 |
| **GM_thickness Morphologist SPasup_right** | 0.08 | 17395 |
| **rfMRI part corr ICA100 edge 130 link 3-26** | 0.08 | 15864 |
| **rfMRI part corr ICA25 edge 139 link 9-16** | 0.08 | 15864 |
| **rfMRI full corr ICA100 edge 470 link 10-30** | 0.08 | 15864 |
| **rfMRI part corr ICA100 edge 1083 link 27-31** | 0.08 | 15864 |
| **hull_junction_length Morphologist SCsylvian_left** | 0.08 | 13814 |
| **rfMRI part corr ICA100 edge 432 link 9-37** | 0.08 | 15864 |
| **rfMRI part corr ICA100 edge 885 link 20-50** | 0.08 | 15864 |
| **SurfArea Destrieux G_and_S_transv_frontopol.lh** | 0.08 | 17127 |
| **SurfArea Desikan paracentral.lh** | 0.08 | 17127 |
| **GrayVol Destrieux S_precentral-inf-part.rh** | 0.08 | 17127 |
| **rfMRI part corr ICA100 edge 241 link 5-36** | 0.08 | 15864 |
| **rfMRI full corr ICA100 edge 605 link 13-36** | 0.09 | 15864 |
| **rfMRI full corr ICA25 edge 130 link 8-19** | 0.09 | 15864 |
| **rfMRI part corr ICA100 edge 750 link 17-23** | 0.09 | 15864 |
| **rfMRI full corr ICA25 edge 18 link 1-19** | 0.09 | 15864 |
| **meandepth Morphologist STiant_right** | 0.09 | 18100 |
| **rfMRI part corr ICA100 edge 626 link 14-16** | 0.09 | 15864 |
| **rfMRI part corr ICA100 edge 1463 link 48-54** | 0.09 | 15864 |
| **rfMRI part corr ICA100 edge 375 link 8-26** | 0.09 | 15864 |
| **rfMRI part corr ICA100 edge 603 link 13-34** | 0.09 | 15864 |
| **rfMRI full corr ICA100 edge 551 link 12-24** | 0.09 | 15864 |
| **rfMRI part corr ICA100 edge 154 link 3-50** | 0.09 | 15864 |

| | | |
|---|---|---|
| **rfMRI part corr ICA100 edge 940 link 22-38** | 0.09 | 15864 |
| **rfMRI part corr ICA25 edge 193 link 15-19** | 0.09 | 15864 |
| **rfMRI full corr ICA100 edge 361 link 8-12** | 0.09 | 15863 |
| **GrayVol Destrieux G_front_sup.rh** | 0.09 | 17127 |
| **maxdepth Morphologist SFpolairetr_left** | 0.09 | 18062 |
| **rfMRI full corr ICA100 edge 320 link 7-18** | 0.09 | 15864 |
| **rfMRI full corr ICA100 edge 216 link 5-11** | 0.09 | 15864 |
| **rfMRI full corr ICA100 edge 797 link 18-33** | 0.09 | 15864 |
| **GrayVol Destrieux Lat_Fis-ant-Vertical.rh** | 0.09 | 17114 |
| **ThickAvg Destrieux S_parieto_occipital.lh** | 0.09 | 17127 |
| **rfMRI part corr ICA100 edge 905 link 21-36** | 0.09 | 15864 |
| **rfMRI part corr ICA100 edge 111 link 3-7** | 0.09 | 15864 |
| **rfMRI part corr ICA100 edge 1074 link 26-50** | 0.09 | 15864 |
| **rfMRI part corr ICA100 edge 20 link 1-21** | 0.09 | 15864 |
| **rfMRI part corr ICA100 edge 19 link 1-20** | 0.09 | 15864 |
| **rfMRI full corr ICA100 edge 1100 link 27-48** | 0.09 | 15864 |
| **MO Uncinate_fasciculus_L** | 0.09 | 16541 |
| **opening Morphologist SPeCmarginal_left** | 0.09 | 15613 |
| **rfMRI full corr ICA25 edge 177 link 13-16** | 0.09 | 15864 |
| **rfMRI full corr ICA100 edge 1167 link 30-37** | 0.09 | 15863 |
| **rfMRI full corr ICA100 edge 572 link 12-45** | 0.09 | 15864 |
| **rfMRI part corr ICA100 edge 1033 link 25-38** | 0.09 | 15864 |
| **rfMRI part corr ICA100 edge 576 link 12-49** | 0.09 | 15864 |
| **GM_thickness Morphologist SCsylvian_right** | 0.09 | 14984 |
| **rfMRI part corr ICA100 edge 1403 link 42-51** | 0.09 | 15864 |
| **rfMRI full corr ICA25 edge 41 link 3-5** | 0.09 | 15864 |
| **maxdepth Morphologist SRh_left** | 0.09 | 17719 |
| **rfMRI full corr ICA100 edge 109 link 3-5** | 0.09 | 15863 |
| **MD Superior_corona_radiata_L** | 0.09 | 16541 |
| **opening Morphologist SPaint_left** | 0.09 | 18007 |
| **rfMRI full corr ICA100 edge 822 link 19-22** | 0.09 | 15864 |
| **rfMRI part corr ICA100 edge 574 link 12-47** | 0.09 | 15864 |
| **surface Morphologist OCCIPITAL_left** | 0.09 | 18097 |
| **rfMRI part corr ICA25 edge 179 link 13-18** | 0.09 | 15864 |

| | | |
|---|---|---|
| **rfMRI part corr ICA100 edge 305 link 6-51** | 0.09 | 15864 |
| **rfMRI part corr ICA100 edge 1150 link 29-45** | 0.09 | 15864 |
| **meandepth Morphologist SLiant_left** | 0.09 | 17411 |
| **rfMRI full corr ICA100 edge 641 link 14-31** | 0.09 | 15860 |
| **ThickAvg Desikan caudalanteriorcingulate.lh** | 0.09 | 17126 |
| **rfMRI full corr ICA100 edge 650 link 14-40** | 0.09 | 15864 |
| **GrayVol Desikan caudalmiddlefrontal.lh** | 0.09 | 17127 |
| **OD Inferior_cerebellar_peduncle_R** | 0.09 | 16541 |
| **hull_junction_length Morphologist FCLrant_right** | 0.09 | 16205 |
| **rfMRI part corr ICA100 edge 294 link 6-40** | 0.09 | 15864 |
| **rfMRI part corr ICA100 edge 224 link 5-19** | 0.1 | 15864 |
| **SurfArea Destrieux S_central.lh** | 0.1 | 17127 |
| **SurfArea Desikan parstriangularis.lh** | 0.1 | 17127 |
| **GM_thickness Morphologist INSULA_left** | 0.1 | 18092 |
| **hull_junction_length Morphologist SFpolairetr_rigl** | 0.1 | 18086 |
| **rfMRI part corr ICA100 edge 855 link 19-55** | 0.1 | 15864 |
| **rfMRI full corr ICA100 edge 1134 link 28-55** | 0.1 | 15864 |
| **MD Posterior_limb_of_internal_capsule_R** | 0.1 | 16541 |
| **rfMRI full corr ICA100 edge 1114 link 28-35** | 0.1 | 15864 |
| **rfMRI full corr ICA100 edge 1231 link 32-54** | 0.1 | 15864 |
| **rfMRI full corr ICA100 edge 187 link 4-32** | 0.1 | 15864 |
| **rfMRI part corr ICA100 edge 1391 link 41-52** | 0.1 | 15864 |
| **rfMRI part corr ICA100 edge 714 link 16-25** | 0.1 | 15864 |
| **rfMRI part corr ICA100 edge 1102 link 27-50** | 0.1 | 15864 |
| **surface Morphologist SFinter_right** | 0.1 | 18099 |
| **rfMRI part corr ICA100 edge 744 link 16-55** | 0.1 | 15864 |
| **rfMRI part corr ICA100 edge 1343 link 38-49** | 0.1 | 15864 |
| **rfMRI part corr ICA100 edge 361 link 8-12** | 0.1 | 15864 |
| **rfMRI full corr ICA25 edge 7 link 1-8** | 0.1 | 15864 |
| **rfMRI part corr ICA100 edge 784 link 18-20** | 0.1 | 15864 |
| **surface Morphologist OCCIPITAL_right** | 0.1 | 18096 |
| **GrayVol Desikan superiorparietal.rh** | 0.1 | 17127 |
| **ThickAvg Desikan parahippocampal.lh** | 0.1 | 17127 |
| **rfMRI part corr ICA100 edge 240 link 5-35** | 0.1 | 15864 |

| | | |
|---|---|---|
| **rfMRI full corr ICA100 edge 514 link 11-30** | 0.1 | 15864 |
| **rfMRI full corr ICA100 edge 1389 link 41-50** | 0.1 | 15864 |
| **rfMRI part corr ICA100 edge 133 link 3-29** | 0.1 | 15864 |
| **rfMRI full corr ICA100 edge 269 link 6-15** | 0.1 | 15864 |
| **GrayVol Desikan inferiortemporal.rh** | 0.1 | 17127 |
| **rfMRI full corr ICA100 edge 1472 link 50-52** | 0.1 | 15864 |
| **hull_junction_length Morphologist FCLrretroCtr_le** | 0.1 | 16831 |
| **rfMRI part corr ICA100 edge 1381 link 41-42** | 0.1 | 15864 |
| **rfMRI full corr ICA100 edge 773 link 17-46** | 0.1 | 15864 |
| **rfMRI part corr ICA100 edge 946 link 22-44** | 0.1 | 15864 |
| **MD Medial_lemniscus_R** | 0.1 | 16541 |
| **rfMRI full corr ICA100 edge 1189 link 31-35** | 0.1 | 15864 |
| **rfMRI part corr ICA100 edge 58 link 2-6** | 0.1 | 15864 |
| **maxdepth Morphologist SCall_left** | 0.1 | 18064 |
| **CC_Anterior** | 0.1 | 17127 |
| **rfMRI full corr ICA100 edge 683 link 15-33** | 0.1 | 15864 |
| **rfMRI part corr ICA25 edge 170 link 12-17** | 0.1 | 15864 |
| **ThickAvg Destrieux G_Ins_lg_and_S_cent_ins.rh** | 0.1 | 17125 |
| **surface Morphologist SpC_left** | 0.1 | 17043 |
| **rfMRI full corr ICA100 edge 456 link 10-16** | 0.1 | 15864 |
| **SurfArea Destrieux G_front_sup.lh** | 0.1 | 17127 |
| **ISOVF Superior_corona_radiata_L** | 0.1 | 16527 |
| **SurfArea Destrieux S_orbital-H_Shaped.rh** | 0.1 | 17125 |
| **rfMRI full corr ICA100 edge 744 link 16-55** | 0.1 | 15864 |
| **opening Morphologist STsterascpost_left** | 0.1 | 17904 |
| **rfMRI full corr ICA100 edge 666 link 15-16** | 0.1 | 15861 |
| **rfMRI full corr ICA100 edge 515 link 11-31** | 0.1 | 15864 |
| **SurfArea Desikan frontalpole.rh** | 0.1 | 17126 |
| **MO Retrolenticular_part_of_internal_capsule_L** | 0.1 | 16541 |
| **GM_thickness Morphologist STpol_left** | 0.1 | 18067 |
| **rfMRI full corr ICA100 edge 908 link 21-39** | 0.1 | 15862 |
| **rfMRI amplitude ICA100 component 43** | 0.1 | 15863 |
| **rfMRI full corr ICA100 edge 1181 link 30-51** | 0.1 | 15864 |
| **rfMRI full corr ICA100 edge 352 link 7-50** | 0.1 | 15864 |

| | | |
|---|---|---|
| **rfMRI part corr ICA100 edge 84 link 2-32** | 0.1 | 15864 |
| **rfMRI full corr ICA100 edge 55 link 2-3** | 0.1 | 15864 |
| **rfMRI part corr ICA100 edge 233 link 5-28** | 0.1 | 15864 |
| **rfMRI part corr ICA25 edge 44 link 3-8** | 0.1 | 15864 |
| **rfMRI full corr ICA25 edge 104 link 6-20** | 0.1 | 15864 |
| **SurfArea Destrieux G_oc-temp_med-Lingual.rh** | 0.1 | 17127 |
| **rfMRI full corr ICA100 edge 917 link 21-48** | 0.1 | 15864 |
| **surface Morphologist STsterascant_right** | 0.1 | 17642 |
| **rfMRI full corr ICA100 edge 1404 link 42-52** | 0.1 | 15864 |
| **rfMRI amplitude ICA25 component 16** | 0.1 | 15863 |
| **opening Morphologist SFint_left** | 0.11 | 18101 |
| **GrayVol Destrieux G_and_S_cingul-Mid-Post.lh** | 0.11 | 17127 |
| **rfMRI part corr ICA100 edge 390 link 8-41** | 0.11 | 15864 |
| **rfMRI full corr ICA100 edge 1448 link 46-54** | 0.11 | 15864 |
| **rfMRI part corr ICA100 edge 860 link 20-25** | 0.11 | 15864 |
| **rfMRI part corr ICA100 edge 1137 link 29-32** | 0.11 | 15864 |
| **rfMRI part corr ICA100 edge 974 link 23-40** | 0.11 | 15864 |
| **rfMRI full corr ICA100 edge 608 link 13-39** | 0.11 | 15864 |
| **rfMRI part corr ICA100 edge 910 link 21-41** | 0.11 | 15864 |
| **ISOVF Tapetum_R** | 0.11 | 16527 |
| **rfMRI full corr ICA100 edge 642 link 14-32** | 0.11 | 15864 |
| **ThickAvg Destrieux G_temp_sup-Lateral.rh** | 0.11 | 17127 |
| **rfMRI part corr ICA100 edge 490 link 10-50** | 0.11 | 15864 |
| **rfMRI full corr ICA100 edge 461 link 10-21** | 0.11 | 15864 |
| **rfMRI amplitude ICA25 component 10** | 0.11 | 15863 |
| **rfMRI part corr ICA100 edge 710 link 16-21** | 0.11 | 15864 |
| **GrayVol Destrieux S_parieto_occipital.rh** | 0.11 | 17127 |
| **rfMRI part corr ICA100 edge 18 link 1-19** | 0.11 | 15864 |
| **rfMRI full corr ICA25 edge 11 link 1-12** | 0.11 | 15864 |
| **rfMRI part corr ICA100 edge 923 link 21-54** | 0.11 | 15864 |
| **rfMRI full corr ICA100 edge 249 link 5-44** | 0.11 | 15864 |
| **rfMRI full corr ICA100 edge 69 link 2-17** | 0.11 | 15864 |
| **rfMRI full corr ICA25 edge 21 link 2-3** | 0.11 | 15864 |
| **rfMRI part corr ICA100 edge 1275 link 34-55** | 0.11 | 15864 |

| | | |
|---|---|---|
| **meandepth Morphologist SPeCmarginal_left** | 0.11 | 15566 |
| **rfMRI part corr ICA100 edge 3 link 1-4** | 0.11 | 15864 |
| **rfMRI full corr ICA100 edge 1186 link 31-32** | 0.11 | 15861 |
| **rfMRI part corr ICA100 edge 1162 link 30-32** | 0.11 | 15864 |
| **rfMRI part corr ICA100 edge 256 link 5-51** | 0.11 | 15864 |
| **SurfArea Desikan supramarginal.lh** | 0.11 | 17127 |
| **rfMRI full corr ICA100 edge 15 link 1-16** | 0.11 | 15864 |
| **rfMRI part corr ICA100 edge 199 link 4-44** | 0.11 | 15864 |
| **surface Morphologist FIPrint1_left** | 0.11 | 17282 |
| **rfMRI full corr ICA100 edge 923 link 21-54** | 0.11 | 15862 |
| **meandepth Morphologist STsterascant_right** | 0.11 | 17627 |
| **rfMRI part corr ICA100 edge 1166 link 30-36** | 0.11 | 15864 |
| **rfMRI part corr ICA100 edge 896 link 21-27** | 0.11 | 15864 |
| **rfMRI part corr ICA100 edge 1244 link 33-45** | 0.11 | 15864 |
| **OD Body_of_corpus_callosum** | 0.11 | 16541 |
| **rfMRI full corr ICA100 edge 289 link 6-35** | 0.11 | 15864 |
| **rfMRI amplitude ICA25 component 14** | 0.11 | 15863 |
| **rfMRI part corr ICA100 edge 408 link 9-13** | 0.11 | 15864 |
| **rfMRI full corr ICA25 edge 33 link 2-15** | 0.11 | 15864 |
| **rfMRI full corr ICA100 edge 1215 link 32-38** | 0.11 | 15864 |
| **opening Morphologist SOIf_right** | 0.11 | 18091 |
| **rfMRI part corr ICA100 edge 1085 link 27-33** | 0.11 | 15864 |
| **SurfArea Destrieux S_front_middle.lh** | 0.11 | 17127 |
| **GrayVol Desikan paracentral.rh** | 0.11 | 17127 |
| **rfMRI part corr ICA100 edge 1055 link 26-31** | 0.12 | 15864 |
| **ThickAvg Desikan precuneus.rh** | 0.12 | 17127 |
| **maxdepth Morphologist FIPrint2_right** | 0.12 | 13064 |
| **rfMRI part corr ICA25 edge 159 link 11-15** | 0.12 | 15864 |
| **rfMRI part corr ICA100 edge 396 link 8-47** | 0.12 | 15864 |
| **meandepth Morphologist FPO_right** | 0.12 | 18097 |
| **SurfArea Destrieux G_front_inf-Opercular.lh** | 0.12 | 17127 |
| **SurfArea Destrieux S_oc_middle_and_Lunatus.rh** | 0.12 | 17127 |
| **rfMRI full corr ICA100 edge 262 link 6-8** | 0.12 | 15863 |
| **rfMRI part corr ICA100 edge 87 link 2-35** | 0.12 | 15864 |

| | | |
|---|---|---|
| **GrayVol Destrieux S_oc_sup_and_transversal.lh** | 0.12 | 17126 |
| **rfMRI part corr ICA100 edge 347 link 7-45** | 0.12 | 15864 |
| **ICVF Cingulum-hippocampus-R** | 0.12 | 16540 |
| **opening Morphologist SGSM_left** | 0.12 | 12712 |
| **surface Morphologist FCLrdiag_right** | 0.12 | 11519 |
| **GrayVol Desikan superiorfrontal.rh** | 0.12 | 17127 |
| **rfMRI full corr ICA100 edge 1041 link 25-46** | 0.12 | 15864 |
| **opening Morphologist FColl_left** | 0.12 | 18101 |
| **rfMRI part corr ICA100 edge 809 link 18-45** | 0.12 | 15864 |
| **rfMRI full corr ICA100 edge 959 link 23-25** | 0.12 | 15864 |
| **rfMRI part corr ICA100 edge 31 link 1-32** | 0.12 | 15864 |
| **rfMRI part corr ICA100 edge 1011 link 24-46** | 0.12 | 15864 |
| **ThickAvg Destrieux S_temporal_inf.lh** | 0.12 | 17127 |
| **rfMRI full corr ICA100 edge 1021 link 25-26** | 0.12 | 15864 |
| **rfMRI full corr ICA100 edge 1067 link 26-43** | 0.12 | 15864 |
| **rfMRI full corr ICA100 edge 1364 link 39-54** | 0.12 | 15864 |
| **rfMRI full corr ICA25 edge 91 link 6-7** | 0.12 | 15863 |
| **GrayVol Destrieux S_circular_insula_ant.lh** | 0.12 | 17126 |
| **rfMRI full corr ICA100 edge 706 link 16-17** | 0.12 | 15861 |
| **rfMRI part corr ICA100 edge 1164 link 30-34** | 0.12 | 15864 |
| **rfMRI part corr ICA100 edge 1444 link 46-50** | 0.12 | 15864 |
| **rfMRI part corr ICA100 edge 897 link 21-28** | 0.12 | 15864 |
| **rfMRI full corr ICA100 edge 285 link 6-31** | 0.12 | 15864 |
| **rfMRI full corr ICA100 edge 857 link 20-22** | 0.12 | 15863 |
| **rfMRI part corr ICA100 edge 1224 link 32-47** | 0.12 | 15864 |
| **hull_junction_length Morphologist FCalant-ScCal_** | 0.12 | 18100 |
| **GrayVol Destrieux S_precentral-sup-part.rh** | 0.12 | 17127 |
| **rfMRI full corr ICA100 edge 745 link 17-18** | 0.12 | 15864 |
| **meandepth Morphologist SLiant_right** | 0.12 | 17201 |
| **MO External_capsule_R** | 0.12 | 16541 |
| **rfMRI full corr ICA100 edge 963 link 23-29** | 0.12 | 15864 |
| **rfMRI full corr ICA100 edge 449 link 9-54** | 0.12 | 15864 |
| **rfMRI part corr ICA100 edge 379 link 8-30** | 0.12 | 15864 |
| **maxdepth Morphologist SPeCmarginal_right** | 0.12 | 17010 |

| | | |
|---|---|---|
| **rfMRI full corr ICA100 edge 778 link 17-51** | 0.12 | 15864 |
| **rfMRI full corr ICA100 edge 1319 link 37-42** | 0.12 | 15864 |
| **rfMRI part corr ICA100 edge 474 link 10-34** | 0.12 | 15864 |
| **rfMRI full corr ICA100 edge 756 link 17-29** | 0.12 | 15864 |
| **rfMRI part corr ICA100 edge 1335 link 38-41** | 0.12 | 15864 |
| **rfMRI part corr ICA100 edge 830 link 19-30** | 0.12 | 15864 |
| **surface Morphologist SFinfant_right** | 0.12 | 18042 |
| **rfMRI part corr ICA100 edge 1175 link 30-45** | 0.12 | 15864 |
| **GrayVol Destrieux Lat_Fis-post.rh** | 0.12 | 17127 |
| **rfMRI full corr ICA100 edge 1002 link 24-37** | 0.12 | 15864 |
| **rfMRI full corr ICA100 edge 149 link 3-45** | 0.13 | 15859 |
| **ThickAvg Destrieux Pole_occipital.rh** | 0.13 | 17127 |
| **rfMRI full corr ICA100 edge 297 link 6-43** | 0.13 | 15864 |
| **rfMRI part corr ICA100 edge 185 link 4-30** | 0.13 | 15864 |
| **rfMRI part corr ICA100 edge 901 link 21-32** | 0.13 | 15864 |
| **rfMRI part corr ICA100 edge 388 link 8-39** | 0.13 | 15864 |
| **maxdepth Morphologist FColl_right** | 0.13 | 18100 |
| **rfMRI full corr ICA100 edge 624 link 13-55** | 0.13 | 15864 |
| **opening Morphologist FCLp_left** | 0.13 | 18101 |
| **rfMRI part corr ICA100 edge 100 link 2-48** | 0.13 | 15864 |
| **rfMRI part corr ICA100 edge 608 link 13-39** | 0.13 | 15864 |
| **rfMRI part corr ICA100 edge 412 link 9-17** | 0.13 | 15864 |
| **ThickAvg Desikan isthmuscingulate.rh** | 0.13 | 17127 |
| **rfMRI full corr ICA25 edge 125 link 8-14** | 0.13 | 15862 |
| **OD Superior_fronto-occipital_fasciculus-part_of_a** | 0.13 | 16541 |
| **rfMRI full corr ICA100 edge 73 link 2-21** | 0.13 | 15864 |
| **rfMRI part corr ICA100 edge 1456 link 47-54** | 0.13 | 15864 |
| **GrayVol Destrieux S_front_middle.lh** | 0.13 | 17127 |
| **rfMRI part corr ICA100 edge 573 link 12-46** | 0.13 | 15864 |
| **rfMRI part corr ICA100 edge 593 link 13-24** | 0.13 | 15864 |
| **rfMRI part corr ICA100 edge 270 link 6-16** | 0.13 | 15864 |
| **rfMRI part corr ICA100 edge 188 link 4-33** | 0.13 | 15864 |
| **maxdepth Morphologist FCLrretroCtr_left** | 0.13 | 16831 |
| **rfMRI amplitude ICA100 component 51** | 0.13 | 15863 |

| | | |
|---|---|---|
| **rfMRI part corr ICA100 edge 1307 link 36-48** | 0.13 | 15864 |
| **SurfArea Destrieux S_oc-temp_med_and_Lingual.** | 0.13 | 17127 |
| **rfMRI full corr ICA100 edge 374 link 8-25** | 0.13 | 15864 |
| **opening Morphologist SRh_right** | 0.13 | 17623 |
| **SurfArea Destrieux G_cingul-Post-ventral.lh** | 0.13 | 17127 |
| **rfMRI full corr ICA100 edge 303 link 6-49** | 0.13 | 15863 |
| **ThickAvg Destrieux G_temp_sup-Lateral.lh** | 0.13 | 17127 |
| **maxdepth Morphologist FCLrasc_right** | 0.13 | 17670 |
| **rfMRI full corr ICA100 edge 266 link 6-12** | 0.13 | 15864 |
| **rfMRI full corr ICA100 edge 524 link 11-40** | 0.13 | 15864 |
| **rfMRI full corr ICA100 edge 499 link 11-15** | 0.13 | 15864 |
| **rfMRI part corr ICA100 edge 585 link 13-16** | 0.13 | 15864 |
| **rfMRI full corr ICA100 edge 1011 link 24-46** | 0.13 | 15864 |
| **FA Inferior_cerebellar_peduncle_L** | 0.13 | 16541 |
| **rfMRI full corr ICA100 edge 598 link 13-29** | 0.13 | 15859 |
| **rfMRI part corr ICA25 edge 54 link 3-18** | 0.13 | 15864 |
| **rfMRI part corr ICA100 edge 1228 link 32-51** | 0.13 | 15864 |
| **rfMRI part corr ICA100 edge 482 link 10-42** | 0.13 | 15864 |
| **rfMRI part corr ICA100 edge 886 link 20-51** | 0.13 | 15864 |
| **rfMRI part corr ICA25 edge 105 link 6-21** | 0.13 | 15864 |
| **GrayVol Desikan middletemporal.lh** | 0.13 | 17127 |
| **rfMRI full corr ICA100 edge 523 link 11-39** | 0.13 | 15864 |
| **rfMRI full corr ICA100 edge 207 link 4-52** | 0.13 | 15864 |
| **rfMRI part corr ICA100 edge 1173 link 30-43** | 0.14 | 15864 |
| **rfMRI part corr ICA100 edge 455 link 10-15** | 0.14 | 15864 |
| **rfMRI full corr ICA100 edge 1335 link 38-41** | 0.14 | 15864 |
| **rfMRI full corr ICA100 edge 763 link 17-36** | 0.14 | 15863 |
| **rfMRI full corr ICA100 edge 703 link 15-53** | 0.14 | 15864 |
| **rfMRI part corr ICA100 edge 378 link 8-29** | 0.14 | 15864 |
| **rfMRI full corr ICA100 edge 68 link 2-16** | 0.14 | 15861 |
| **rfMRI part corr ICA100 edge 1436 link 45-51** | 0.14 | 15864 |
| **rfMRI full corr ICA100 edge 590 link 13-21** | 0.14 | 15864 |
| **rfMRI part corr ICA100 edge 274 link 6-20** | 0.14 | 15864 |
| **rfMRI full corr ICA25 edge 193 link 15-19** | 0.14 | 15864 |

| | | |
|---|---|---|
| **ThickAvg Destrieux G_precentral.rh** | 0.14 | 17127 |
| **opening Morphologist SRinf_left** | 0.14 | 16697 |
| **rfMRI full corr ICA100 edge 70 link 2-18** | 0.14 | 15861 |
| **rfMRI part corr ICA100 edge 852 link 19-52** | 0.14 | 15864 |
| **rfMRI full corr ICA100 edge 1398 link 42-46** | 0.14 | 15864 |
| **rfMRI full corr ICA100 edge 779 link 17-52** | 0.14 | 15864 |
| **rfMRI full corr ICA100 edge 1064 link 26-40** | 0.14 | 15864 |
| **rfMRI part corr ICA100 edge 1477 link 51-53** | 0.14 | 15864 |
| **rfMRI full corr ICA100 edge 497 link 11-13** | 0.14 | 15864 |
| **rfMRI full corr ICA25 edge 169 link 12-16** | 0.14 | 15864 |
| **OD Medial_lemniscus_R** | 0.14 | 16541 |
| **ThickAvg Destrieux G_precuneus.rh** | 0.14 | 17127 |
| **rfMRI part corr ICA100 edge 776 link 17-49** | 0.14 | 15864 |
| **rfMRI part corr ICA100 edge 532 link 11-48** | 0.14 | 15864 |
| **rfMRI part corr ICA25 edge 126 link 8-15** | 0.14 | 15864 |
| **rfMRI full corr ICA100 edge 58 link 2-6** | 0.14 | 15864 |
| **rfMRI part corr ICA100 edge 947 link 22-45** | 0.14 | 15864 |
| **rfMRI full corr ICA100 edge 1026 link 25-31** | 0.14 | 15864 |
| **rfMRI full corr ICA25 edge 110 link 7-12** | 0.14 | 15864 |
| **rfMRI part corr ICA100 edge 624 link 13-55** | 0.14 | 15864 |
| **rfMRI part corr ICA100 edge 801 link 18-37** | 0.14 | 15864 |
| **rfMRI part corr ICA100 edge 1126 link 28-47** | 0.14 | 15864 |
| **rfMRI part corr ICA100 edge 30 link 1-31** | 0.14 | 15864 |
| **rfMRI full corr ICA100 edge 195 link 4-40** | 0.14 | 15864 |
| **rfMRI part corr ICA100 edge 1390 link 41-51** | 0.14 | 15864 |
| **GrayVol Destrieux G_oc-temp_lat-fusifor.lh** | 0.14 | 17127 |
| **Right-Putamen** | 0.14 | 17127 |
| **surface Morphologist SC_right** | 0.14 | 18100 |
| **rfMRI part corr ICA100 edge 1349 link 38-55** | 0.14 | 15864 |
| **rfMRI full corr ICA100 edge 174 link 4-19** | 0.14 | 15864 |
| **hull_junction_length Morphologist FIPrint1_left** | 0.14 | 17282 |
| **rfMRI full corr ICA100 edge 808 link 18-44** | 0.14 | 15864 |
| **rfMRI full corr ICA100 edge 473 link 10-33** | 0.14 | 15864 |
| **ThickAvg Destrieux G_and_S_transv_frontopol.rh** | 0.14 | 17127 |

| | | |
|---|---|---|
| **rfMRI full corr ICA100 edge 1057 link 26-33** | 0.14 | 15864 |
| **rfMRI part corr ICA100 edge 246 link 5-41** | 0.14 | 15864 |
| **rfMRI full corr ICA100 edge 1072 link 26-48** | 0.14 | 15864 |
| **rfMRI full corr ICA100 edge 135 link 3-31** | 0.14 | 15863 |
| **rfMRI full corr ICA100 edge 379 link 8-30** | 0.14 | 15864 |
| **rfMRI part corr ICA25 edge 174 link 12-21** | 0.15 | 15864 |
| **meandepth Morphologist SPeCsup_right** | 0.15 | 17267 |
| **rfMRI full corr ICA100 edge 1296 link 36-37** | 0.15 | 15864 |
| **rfMRI full corr ICA100 edge 1313 link 36-54** | 0.15 | 15864 |
| **rfMRI full corr ICA100 edge 444 link 9-49** | 0.15 | 15863 |
| **rfMRI amplitude ICA25 component 15** | 0.15 | 15863 |
| **rfMRI part corr ICA100 edge 824 link 19-24** | 0.15 | 15864 |
| **rfMRI part corr ICA100 edge 117 link 3-13** | 0.15 | 15864 |
| **rfMRI part corr ICA100 edge 1051 link 26-27** | 0.15 | 15864 |
| **rfMRI part corr ICA25 edge 37 link 2-19** | 0.15 | 15864 |
| **rfMRI full corr ICA100 edge 1031 link 25-36** | 0.15 | 15862 |
| **maxdepth Morphologist SPeCsup_right** | 0.15 | 17272 |
| **rfMRI part corr ICA100 edge 702 link 15-52** | 0.15 | 15864 |
| **rfMRI part corr ICA100 edge 465 link 10-25** | 0.15 | 15864 |
| **hull_junction_length Morphologist SFinfant_right** | 0.15 | 18042 |
| **rfMRI part corr ICA100 edge 470 link 10-30** | 0.15 | 15864 |
| **rfMRI amplitude ICA25 component 20** | 0.15 | 15863 |
| **rfMRI part corr ICA100 edge 449 link 9-54** | 0.15 | 15864 |
| **Brain-Stem** | 0.15 | 17127 |
| **GrayVol Desikan pericalcarine.rh** | 0.15 | 17127 |
| **rfMRI amplitude ICA100 component 23** | 0.15 | 15863 |
| **rfMRI full corr ICA100 edge 197 link 4-42** | 0.15 | 15864 |
| **rfMRI full corr ICA100 edge 813 link 18-49** | 0.15 | 15864 |
| **rfMRI part corr ICA100 edge 1009 link 24-44** | 0.15 | 15864 |
| **rfMRI full corr ICA100 edge 965 link 23-31** | 0.15 | 15864 |
| **rfMRI part corr ICA25 edge 200 link 16-21** | 0.15 | 15864 |
| **meandepth Morphologist FCLrretroCtr_left** | 0.15 | 16743 |
| **Right-Caudate** | 0.15 | 17127 |
| **GrayVol Desikan postcentral.lh** | 0.15 | 17127 |

| | | |
|---|---|---|
| **rfMRI full corr ICA100 edge 790 link 18-26** | 0.15 | 15864 |
| **rfMRI full corr ICA100 edge 967 link 23-33** | 0.15 | 15864 |
| **rfMRI part corr ICA100 edge 236 link 5-31** | 0.15 | 15864 |
| **GrayVol Destrieux G_temporal_middle.rh** | 0.15 | 17127 |
| **rfMRI amplitude ICA100 component 53** | 0.15 | 15863 |
| **rfMRI part corr ICA100 edge 1154 link 29-49** | 0.15 | 15864 |
| **rfMRI full corr ICA100 edge 1441 link 46-47** | 0.15 | 15864 |
| **rfMRI full corr ICA100 edge 1008 link 24-43** | 0.15 | 15861 |
| **rfMRI part corr ICA100 edge 88 link 2-36** | 0.15 | 15864 |
| **meandepth Morphologist FIPrint2_right** | 0.15 | 12965 |
| **surface Morphologist FCMpost_left** | 0.15 | 18101 |
| **rfMRI full corr ICA100 edge 1061 link 26-37** | 0.15 | 15864 |
| **SurfArea Destrieux G_cingul-Post-dorsal.lh** | 0.15 | 17125 |
| **GrayVol Destrieux G_insular_short.rh** | 0.15 | 17125 |
| **rfMRI full corr ICA100 edge 1381 link 41-42** | 0.15 | 15864 |
| **rfMRI full corr ICA100 edge 510 link 11-26** | 0.15 | 15864 |
| **rfMRI part corr ICA100 edge 403 link 8-54** | 0.16 | 15864 |
| **rfMRI part corr ICA100 edge 1345 link 38-51** | 0.16 | 15864 |
| **rfMRI part corr ICA100 edge 1161 link 30-31** | 0.16 | 15864 |
| **opening Morphologist SCu_left** | 0.16 | 18058 |
| **rfMRI part corr ICA100 edge 914 link 21-45** | 0.16 | 15864 |
| **rfMRI full corr ICA100 edge 668 link 15-18** | 0.16 | 15864 |
| **SurfArea Destrieux S_oc_sup_and_transversal.lh** | 0.16 | 17126 |
| **rfMRI part corr ICA100 edge 227 link 5-22** | 0.16 | 15864 |
| **opening Morphologist SFmarginal_right** | 0.16 | 18006 |
| **rfMRI full corr ICA100 edge 205 link 4-50** | 0.16 | 15864 |
| **rfMRI part corr ICA100 edge 629 link 14-19** | 0.16 | 15864 |
| **rfMRI full corr ICA100 edge 239 link 5-34** | 0.16 | 15864 |
| **rfMRI part corr ICA100 edge 299 link 6-45** | 0.16 | 15864 |
| **GrayVol Destrieux G_temp_sup-Plan_polar.rh** | 0.16 | 17126 |
| **hull_junction_length Morphologist SpC_left** | 0.16 | 17043 |
| **meandepth Morphologist SForbitaire_left** | 0.16 | 17519 |
| **rfMRI part corr ICA100 edge 1104 link 27-52** | 0.16 | 15864 |
| **GrayVol Desikan frontalpole.rh** | 0.16 | 17126 |

| | | |
|---|---|---|
| **rfMRI part corr ICA100 edge 928 link 22-26** | 0.16 | 15864 |
| **GM_thickness Morphologist STipost_left** | 0.16 | 18087 |
| **rfMRI part corr ICA100 edge 835 link 19-35** | 0.16 | 15864 |
| **rfMRI full corr ICA100 edge 646 link 14-36** | 0.16 | 15864 |
| **rfMRI part corr ICA100 edge 173 link 4-18** | 0.16 | 15864 |
| **MD Superior_fronto-occipital_fasciculus-part_of_a** | 0.16 | 16541 |
| **rfMRI full corr ICA100 edge 732 link 16-43** | 0.16 | 15863 |
| **rfMRI part corr ICA100 edge 859 link 20-24** | 0.16 | 15864 |
| **GrayVol Destrieux G_pariet_inf-Supramar.lh** | 0.16 | 17127 |
| **rfMRI amplitude ICA100 component 15** | 0.16 | 15863 |
| **rfMRI full corr ICA100 edge 429 link 9-34** | 0.16 | 15864 |
| **rfMRI full corr ICA100 edge 941 link 22-39** | 0.16 | 15864 |
| **GrayVol Desikan caudalanteriorcingulate.lh** | 0.16 | 17126 |
| **rfMRI full corr ICA100 edge 298 link 6-44** | 0.16 | 15864 |
| **rfMRI full corr ICA100 edge 264 link 6-10** | 0.16 | 15864 |
| **rfMRI part corr ICA100 edge 645 link 14-35** | 0.16 | 15864 |
| **OD Posterior_thalamic_radiation-include_optic_ra** | 0.16 | 16541 |
| **rfMRI full corr ICA100 edge 155 link 3-51** | 0.16 | 15864 |
| **maxdepth Morphologist SOTlatant_right** | 0.16 | 18047 |
| **rfMRI full corr ICA100 edge 653 link 14-43** | 0.16 | 15864 |
| **GrayVol Desikan isthmuscingulate.rh** | 0.16 | 17127 |
| **rfMRI full corr ICA100 edge 390 link 8-41** | 0.16 | 15864 |
| **rfMRI full corr ICA100 edge 353 link 7-51** | 0.16 | 15864 |
| **rfMRI amplitude ICA25 component 2** | 0.16 | 15863 |
| **rfMRI part corr ICA100 edge 592 link 13-23** | 0.16 | 15864 |
| **rfMRI part corr ICA100 edge 524 link 11-40** | 0.16 | 15864 |
| **rfMRI full corr ICA100 edge 522 link 11-38** | 0.16 | 15864 |
| **rfMRI part corr ICA100 edge 286 link 6-32** | 0.16 | 15864 |
| **SurfArea Destrieux G_precentral.lh** | 0.16 | 17127 |
| **rfMRI full corr ICA100 edge 982 link 23-48** | 0.16 | 15864 |
| **rfMRI part corr ICA100 edge 679 link 15-29** | 0.16 | 15864 |
| **rfMRI part corr ICA100 edge 598 link 13-29** | 0.16 | 15864 |
| **surface Morphologist FIPPoCinf_left** | 0.16 | 18079 |
| **rfMRI part corr ICA100 edge 994 link 24-29** | 0.16 | 15864 |

| | | |
|---|---|---|
| **rfMRI part corr ICA100 edge 1099 link 27-47** | 0.16 | 15864 |
| **hull_junction_length Morphologist SOTlatant_right** | 0.16 | 18047 |
| **opening Morphologist SFinter_right** | 0.16 | 18099 |
| **rfMRI part corr ICA100 edge 1010 link 24-45** | 0.17 | 15864 |
| **meandepth Morphologist FCalant-ScCal_left** | 0.17 | 18100 |
| **rfMRI full corr ICA100 edge 1309 link 36-50** | 0.17 | 15864 |
| **rfMRI full corr ICA100 edge 51 link 1-52** | 0.17 | 15864 |
| **rfMRI full corr ICA100 edge 907 link 21-38** | 0.17 | 15864 |
| **rfMRI full corr ICA100 edge 940 link 22-38** | 0.17 | 15864 |
| **FA Body_of_corpus_callosum** | 0.17 | 16541 |
| **rfMRI full corr ICA100 edge 710 link 16-21** | 0.17 | 15864 |
| **surface Morphologist SFint_left** | 0.17 | 18101 |
| **rfMRI part corr ICA100 edge 785 link 18-21** | 0.17 | 15864 |
| **hull_junction_length Morphologist SLiant_left** | 0.17 | 17423 |
| **SurfArea Destrieux G_temporal_middle.rh** | 0.17 | 17127 |
| **rfMRI part corr ICA100 edge 497 link 11-13** | 0.17 | 15864 |
| **ThickAvg Desikan temporalpole.rh** | 0.17 | 17126 |
| **rfMRI part corr ICA25 edge 145 link 10-11** | 0.17 | 15864 |
| **ThickAvg Destrieux G_front_inf-Opercular.lh** | 0.17 | 17127 |
| **rfMRI part corr ICA100 edge 1457 link 47-55** | 0.17 | 15864 |
| **rfMRI amplitude ICA100 component 49** | 0.17 | 15863 |
| **rfMRI part corr ICA100 edge 1433 link 45-48** | 0.17 | 15864 |
| **rfMRI full corr ICA100 edge 1150 link 29-45** | 0.17 | 15864 |
| **rfMRI full corr ICA100 edge 431 link 9-36** | 0.17 | 15863 |
| **rfMRI part corr ICA100 edge 65 link 2-13** | 0.17 | 15864 |
| **rfMRI part corr ICA25 edge 188 link 14-20** | 0.17 | 15864 |
| **ThickAvg Destrieux S_orbital_lateral.lh** | 0.17 | 17126 |
| **surface Morphologist FCLrasc_right** | 0.17 | 17670 |
| **rfMRI full corr ICA100 edge 971 link 23-37** | 0.17 | 15863 |
| **ISOVF External_capsule_R** | 0.17 | 16527 |
| **MO Superior_longitudinal_fasciculus_R** | 0.17 | 16541 |
| **rfMRI part corr ICA100 edge 122 link 3-18** | 0.17 | 15864 |
| **SurfArea Desikan middletemporal.lh** | 0.17 | 17127 |
| **rfMRI full corr ICA100 edge 776 link 17-49** | 0.17 | 15864 |

| | | |
|---|---|---|
| **rfMRI full corr ICA100 edge 1345 link 38-51** | 0.17 | 15864 |
| **GrayVol Destrieux S_orbital_lateral.lh** | 0.17 | 17126 |
| **meandepth Morphologist SC_right** | 0.17 | 18100 |
| **opening Morphologist SOTlatpost_right** | 0.17 | 18075 |
| **rfMRI full corr ICA100 edge 1158 link 29-53** | 0.17 | 15864 |
| **GM_thickness Morphologist FCLrscpost_right** | 0.17 | 16115 |
| **rfMRI part corr ICA100 edge 1158 link 29-53** | 0.17 | 15864 |
| **rfMRI full corr ICA100 edge 1300 link 36-41** | 0.17 | 15864 |
| **maxdepth Morphologist STsterascant_left** | 0.17 | 17560 |
| **SurfArea Destrieux S_intrapariet_and_P_trans.rh** | 0.17 | 17127 |
| **SurfArea Desikan parahippocampal.lh** | 0.17 | 17127 |
| **surface Morphologist SPeCmarginal_left** | 0.17 | 15613 |
| **rfMRI part corr ICA100 edge 843 link 19-43** | 0.17 | 15864 |
| **rfMRI part corr ICA100 edge 280 link 6-26** | 0.17 | 15864 |
| **rfMRI full corr ICA100 edge 1459 link 48-50** | 0.17 | 15863 |
| **rfMRI part corr ICA100 edge 660 link 14-50** | 0.17 | 15864 |
| **rfMRI part corr ICA100 edge 446 link 9-51** | 0.17 | 15864 |
| **ThickAvg Desikan cuneus.lh** | 0.17 | 17127 |
| **rfMRI part corr ICA25 edge 138 link 9-15** | 0.17 | 15864 |
| **ThickAvg Destrieux S_interm_prim-Jensen.lh** | 0.17 | 17108 |
| **SurfArea Destrieux G_oc-temp_lat-fusifor.lh** | 0.17 | 17127 |
| **ThickAvg Destrieux G_and_S_cingul-Mid-Post.rh** | 0.17 | 17127 |
| **SurfArea Desikan rostralanteriorcingulate.rh** | 0.17 | 17127 |
| **GM_thickness Morphologist SPeCinter_left** | 0.17 | 18018 |
| **rfMRI part corr ICA100 edge 1273 link 34-53** | 0.17 | 15864 |
| **rfMRI amplitude ICA100 component 55** | 0.17 | 15863 |
| **rfMRI full corr ICA100 edge 1228 link 32-51** | 0.17 | 15864 |
| **rfMRI full corr ICA25 edge 64 link 4-11** | 0.17 | 15864 |
| **OD Cingulum-cingulate_gyrus-L** | 0.17 | 16541 |
| **GM_thickness Morphologist INSULA_right** | 0.18 | 18093 |
| **rfMRI part corr ICA100 edge 646 link 14-36** | 0.18 | 15864 |
| **Left-Cerebellum-White-Matter** | 0.18 | 17127 |
| **OD Inferior_cerebellar_peduncle_L** | 0.18 | 16541 |
| **rfMRI full corr ICA100 edge 764 link 17-37** | 0.18 | 15864 |

| | | |
|---|---|---|
| **rfMRI full corr ICA100 edge 1123 link 28-44** | 0.18 | 15864 |
| **rfMRI full corr ICA100 edge 1192 link 31-38** | 0.18 | 15864 |
| **rfMRI part corr ICA100 edge 720 link 16-31** | 0.18 | 15864 |
| **rfMRI amplitude ICA25 component 12** | 0.18 | 15863 |
| **rfMRI part corr ICA100 edge 1287 link 35-47** | 0.18 | 15864 |
| **rfMRI full corr ICA100 edge 852 link 19-52** | 0.18 | 15864 |
| **rfMRI part corr ICA100 edge 933 link 22-31** | 0.18 | 15864 |
| **rfMRI part corr ICA100 edge 820 link 19-20** | 0.18 | 15864 |
| **maxdepth Morphologist SOTlatant_left** | 0.18 | 18076 |
| **GrayVol Destrieux G_orbital.lh** | 0.18 | 17127 |
| **rfMRI part corr ICA100 edge 28 link 1-29** | 0.18 | 15864 |
| **ThickAvg Destrieux G_Ins_lg_and_S_cent_ins.lh** | 0.18 | 17126 |
| **rfMRI full corr ICA100 edge 475 link 10-35** | 0.18 | 15864 |
| **GrayVol Destrieux S_postcentral.lh** | 0.18 | 17127 |
| **opening Morphologist SPeCinf_right** | 0.18 | 16608 |
| **rfMRI full corr ICA25 edge 86 link 5-17** | 0.18 | 15864 |
| **MO Corticospinal_tract_L** | 0.18 | 16541 |
| **hull_junction_length Morphologist SFmarginal_rig** | 0.18 | 18006 |
| **rfMRI full corr ICA100 edge 1244 link 33-45** | 0.18 | 15864 |
| **rfMRI part corr ICA100 edge 1412 link 43-48** | 0.18 | 15864 |
| **hull_junction_length Morphologist STsterascant_ri** | 0.18 | 17642 |
| **rfMRI part corr ICA100 edge 1018 link 24-53** | 0.18 | 15864 |
| **rfMRI part corr ICA100 edge 1005 link 24-40** | 0.18 | 15864 |
| **rfMRI full corr ICA100 edge 1260 link 34-40** | 0.18 | 15864 |
| **SurfArea Destrieux S_temporal_sup.rh** | 0.18 | 17127 |
| **rfMRI part corr ICA100 edge 1156 link 29-51** | 0.18 | 15864 |
| **rfMRI part corr ICA100 edge 459 link 10-19** | 0.18 | 15864 |
| **rfMRI full corr ICA100 edge 457 link 10-17** | 0.18 | 15864 |
| **rfMRI part corr ICA100 edge 301 link 6-47** | 0.18 | 15864 |
| **rfMRI full corr ICA25 edge 66 link 4-13** | 0.18 | 15864 |
| **rfMRI full corr ICA100 edge 1275 link 34-55** | 0.18 | 15864 |
| **rfMRI full corr ICA100 edge 676 link 15-26** | 0.18 | 15864 |
| **SurfArea Desikan lingual.rh** | 0.18 | 17127 |
| **rfMRI part corr ICA100 edge 873 link 20-38** | 0.18 | 15864 |

| | | |
|---|---|---|
| **ThickAvg Desikan superiorparietal.lh** | 0.18 | 17127 |
| **rfMRI full corr ICA100 edge 393 link 8-44** | 0.18 | 15864 |
| **SurfArea Destrieux G_temporal_inf.rh** | 0.18 | 17127 |
| **rfMRI part corr ICA100 edge 42 link 1-43** | 0.18 | 15864 |
| **rfMRI full corr ICA100 edge 1023 link 25-28** | 0.18 | 15863 |
| **rfMRI part corr ICA100 edge 541 link 12-14** | 0.18 | 15864 |
| **rfMRI full corr ICA100 edge 1216 link 32-39** | 0.18 | 15864 |
| **MD Posterior_thalamic_radiation-include_optic_ra** | 0.18 | 16541 |
| **GM_thickness Morphologist SC_right** | 0.18 | 18093 |
| **maxdepth Morphologist SOp_left** | 0.18 | 15273 |
| **ThickAvg Desikan superiortemporal.rh** | 0.18 | 17127 |
| **rfMRI part corr ICA100 edge 373 link 8-24** | 0.18 | 15864 |
| **GM_thickness Morphologist SLiant_left** | 0.18 | 17414 |
| **rfMRI full corr ICA25 edge 121 link 8-10** | 0.18 | 15864 |
| **rfMRI part corr ICA25 edge 132 link 8-21** | 0.18 | 15864 |
| **meandepth Morphologist SOp_left** | 0.18 | 15232 |
| **maxdepth Morphologist SFpolairetr_right** | 0.18 | 18086 |
| **rfMRI full corr ICA100 edge 443 link 9-48** | 0.18 | 15864 |
| **hull_junction_length Morphologist SGSM_left** | 0.18 | 12712 |
| **rfMRI part corr ICA100 edge 832 link 19-32** | 0.18 | 15864 |
| **rfMRI full corr ICA100 edge 1482 link 52-55** | 0.18 | 15864 |
| **rfMRI part corr ICA100 edge 1013 link 24-48** | 0.18 | 15864 |
| **rfMRI full corr ICA100 edge 101 link 2-49** | 0.18 | 15864 |
| **GM_thickness Morphologist SPeCmedian_right** | 0.19 | 17067 |
| **rfMRI part corr ICA100 edge 238 link 5-33** | 0.19 | 15864 |
| **rfMRI full corr ICA25 edge 150 link 10-16** | 0.19 | 15864 |
| **rfMRI full corr ICA100 edge 747 link 17-20** | 0.19 | 15864 |
| **GrayVol Destrieux G_temporal_inf.rh** | 0.19 | 17127 |
| **rfMRI full corr ICA100 edge 800 link 18-36** | 0.19 | 15864 |
| **hull_junction_length Morphologist SCsylvian_right** | 0.19 | 14992 |
| **rfMRI part corr ICA25 edge 41 link 3-5** | 0.19 | 15864 |
| **rfMRI full corr ICA100 edge 1465 link 49-50** | 0.19 | 15864 |
| **rfMRI full corr ICA100 edge 1210 link 32-33** | 0.19 | 15864 |
| **rfMRI part corr ICA100 edge 39 link 1-40** | 0.19 | 15864 |

| | | |
|---|---|---|
| **rfMRI part corr ICA100 edge 731 link 16-42** | 0.19 | 15864 |
| **GrayVol Destrieux S_central.lh** | 0.19 | 17127 |
| **rfMRI part corr ICA100 edge 680 link 15-30** | 0.19 | 15864 |
| **rfMRI full corr ICA100 edge 479 link 10-39** | 0.19 | 15864 |
| **rfMRI full corr ICA100 edge 1010 link 24-45** | 0.19 | 15864 |
| **rfMRI full corr ICA100 edge 1310 link 36-51** | 0.19 | 15864 |
| **rfMRI part corr ICA100 edge 1387 link 41-48** | 0.19 | 15864 |
| **rfMRI part corr ICA100 edge 856 link 20-21** | 0.19 | 15864 |
| **OD Superior_longitudinal_fasciculus_L** | 0.19 | 16541 |
| **rfMRI part corr ICA100 edge 639 link 14-29** | 0.19 | 15864 |
| **rfMRI part corr ICA100 edge 1195 link 31-41** | 0.19 | 15864 |
| **rfMRI part corr ICA100 edge 952 link 22-50** | 0.19 | 15864 |
| **rfMRI full corr ICA100 edge 146 link 3-42** | 0.19 | 15864 |
| **rfMRI full corr ICA100 edge 83 link 2-31** | 0.19 | 15863 |
| **OD Cingulum-cingulate_gyrus-R** | 0.19 | 16541 |
| **OD Posterior_corona_radiata_L** | 0.19 | 16541 |
| **SurfArea Desikan inferiortemporal.rh** | 0.19 | 17127 |
| **rfMRI full corr ICA100 edge 1334 link 38-40** | 0.19 | 15864 |
| **FA Superior_cerebellar_peduncle_R** | 0.19 | 16541 |
| **rfMRI part corr ICA100 edge 198 link 4-43** | 0.19 | 15864 |
| **rfMRI part corr ICA100 edge 255 link 5-50** | 0.19 | 15864 |
| **meandepth Morphologist SRinf_left** | 0.19 | 16696 |
| **rfMRI full corr ICA100 edge 294 link 6-40** | 0.19 | 15864 |
| **rfMRI part corr ICA25 edge 162 link 11-18** | 0.19 | 15864 |
| **rfMRI part corr ICA100 edge 1372 link 40-47** | 0.19 | 15864 |
| **surface Morphologist SOr_right** | 0.19 | 18098 |
| **rfMRI part corr ICA100 edge 275 link 6-21** | 0.19 | 15864 |
| **hull_junction_length Morphologist SPaint_left** | 0.19 | 18007 |
| **GrayVol Desikan lateralorbitofrontal.lh** | 0.19 | 17127 |
| **rfMRI full corr ICA25 edge 52 link 3-16** | 0.19 | 15864 |
| **hull_junction_length Morphologist SCu_right** | 0.19 | 18068 |
| **surface Morphologist SPeCmarginal_right** | 0.19 | 17010 |
| **rfMRI part corr ICA100 edge 257 link 5-52** | 0.19 | 15864 |
| **rfMRI full corr ICA25 edge 84 link 5-15** | 0.19 | 15864 |

| | | |
|---|---|---|
| GM_thickness Morphologist FCLrasc_right | 0.19 | 17657 |
| hull_junction_length Morphologist SRinf_right | 0.19 | 17661 |
| FA Fornix-cres-Stria_terminalis-not_resolved_with | 0.19 | 16541 |
| GM_thickness Morphologist SOTlatint_right | 0.19 | 15615 |
| rfMRI full corr ICA100 edge 758 link 17-31 | 0.19 | 15864 |
| rfMRI amplitude ICA100 component 54 | 0.19 | 15863 |
| surface Morphologist SGSM_left | 0.19 | 12712 |
| ISOVF External_capsule_L | 0.2 | 16527 |
| rfMRI full corr ICA100 edge 220 link 5-15 | 0.2 | 15863 |
| rfMRI part corr ICA100 edge 1251 link 33-52 | 0.2 | 15864 |
| rfMRI part corr ICA100 edge 1471 link 50-51 | 0.2 | 15864 |
| OD Superior_longitudinal_fasciculus_R | 0.2 | 16541 |
| SurfArea Destrieux G_temp_sup-Plan_tempo.rh | 0.2 | 17127 |
| rfMRI full corr ICA100 edge 968 link 23-34 | 0.2 | 15864 |
| maxdepth Morphologist SOTlatmed_right | 0.2 | 17219 |
| rfMRI full corr ICA100 edge 173 link 4-18 | 0.2 | 15863 |
| rfMRI full corr ICA100 edge 1096 link 27-44 | 0.2 | 15864 |
| rfMRI amplitude ICA100 component 11 | 0.2 | 15863 |
| rfMRI part corr ICA100 edge 1243 link 33-44 | 0.2 | 15864 |
| rfMRI full corr ICA100 edge 1238 link 33-39 | 0.2 | 15864 |
| hull_junction_length Morphologist SPeCmarginal_ | 0.2 | 17010 |
| rfMRI full corr ICA100 edge 1324 link 37-47 | 0.2 | 15864 |
| rfMRI part corr ICA25 edge 66 link 4-13 | 0.2 | 15864 |
| rfMRI full corr ICA100 edge 437 link 9-42 | 0.2 | 15859 |
| hull_junction_length Morphologist FCLrscant_righ | 0.2 | 5353 |
| SurfArea Destrieux G_and_S_cingul-Mid-Post.lh | 0.2 | 17127 |
| SurfArea Destrieux S_temporal_sup.lh | 0.2 | 17127 |
| rfMRI part corr ICA100 edge 166 link 4-11 | 0.2 | 15864 |
| SurfArea Destrieux G_orbital.lh | 0.2 | 17127 |
| GrayVol Destrieux G_temporal_middle.lh | 0.2 | 17127 |
| ThickAvg Desikan postcentral.lh | 0.2 | 17127 |
| maxdepth Morphologist SOTlatpost_left | 0.2 | 18062 |
| rfMRI part corr ICA100 edge 857 link 20-22 | 0.2 | 15864 |
| surface Morphologist SCu_left | 0.2 | 18058 |

| | | |
|---|---|---|
| **rfMRI part corr ICA100 edge 607 link 13-38** | 0.2 | 15864 |
| **MO Sagittal_stratum-inf_longitidinal_fasci_and_inf** | 0.2 | 16541 |
| **rfMRI part corr ICA100 edge 719 link 16-30** | 0.2 | 15864 |
| **rfMRI part corr ICA25 edge 125 link 8-14** | 0.2 | 15864 |
| **rfMRI part corr ICA100 edge 764 link 17-37** | 0.2 | 15864 |
| **rfMRI full corr ICA100 edge 228 link 5-23** | 0.2 | 15864 |
| **rfMRI part corr ICA100 edge 674 link 15-24** | 0.2 | 15864 |
| **rfMRI full corr ICA100 edge 460 link 10-20** | 0.2 | 15864 |
| **rfMRI full corr ICA100 edge 1250 link 33-51** | 0.2 | 15864 |
| **meandepth Morphologist SForbitaire_right** | 0.2 | 17278 |
| **hull_junction_length Morphologist SCLPC_right** | 0.2 | 5279 |
| **rfMRI part corr ICA100 edge 1316 link 37-39** | 0.2 | 15864 |
| **ISOVF Cingulum-hippocampus-L** | 0.2 | 16527 |
| **rfMRI full corr ICA100 edge 340 link 7-38** | 0.2 | 15864 |
| **opening Morphologist SPat_right** | 0.2 | 16760 |
| **surface Morphologist SPeCsup_right** | 0.2 | 17272 |
| **rfMRI amplitude ICA100 component 22** | 0.2 | 15863 |
| **rfMRI part corr ICA100 edge 703 link 15-53** | 0.2 | 15864 |
| **rfMRI part corr ICA100 edge 709 link 16-20** | 0.2 | 15864 |
| **ThickAvg Destrieux S_cingul-Marginalis.rh** | 0.2 | 17127 |
| **rfMRI full corr ICA100 edge 47 link 1-48** | 0.2 | 15862 |
| **opening Morphologist SOTlatmed_left** | 0.2 | 17430 |
| **rfMRI full corr ICA100 edge 1211 link 32-34** | 0.2 | 15864 |
| **maxdepth Morphologist STsterascpost_left** | 0.21 | 17904 |
| **rfMRI amplitude ICA25 component 4** | 0.21 | 15863 |
| **FA Genu_of_corpus_callosum** | 0.21 | 16541 |
| **rfMRI part corr ICA25 edge 131 link 8-20** | 0.21 | 15864 |
| **rfMRI full corr ICA100 edge 772 link 17-45** | 0.21 | 15864 |
| **rfMRI full corr ICA100 edge 275 link 6-21** | 0.21 | 15864 |
| **rfMRI part corr ICA100 edge 687 link 15-37** | 0.21 | 15864 |
| **opening Morphologist FCMant_right** | 0.21 | 17828 |
| **opening Morphologist SPeCinter_right** | 0.21 | 18084 |
| **rfMRI part corr ICA100 edge 451 link 10-11** | 0.21 | 15864 |
| **rfMRI part corr ICA100 edge 894 link 21-25** | 0.21 | 15864 |

| | | |
|---|---|---|
| **rfMRI part corr ICA25 edge 183 link 14-15** | 0.21 | 15864 |
| **GrayVol Desikan precentral.rh** | 0.21 | 17127 |
| **rfMRI full corr ICA25 edge 36 link 2-18** | 0.21 | 15863 |
| **rfMRI full corr ICA100 edge 1164 link 30-34** | 0.21 | 15864 |
| **rfMRI part corr ICA25 edge 199 link 16-20** | 0.21 | 15864 |
| **rfMRI full corr ICA100 edge 1306 link 36-47** | 0.21 | 15864 |
| **ThickAvg Destrieux G_cuneus.rh** | 0.21 | 17127 |
| **rfMRI full corr ICA100 edge 496 link 11-12** | 0.21 | 15864 |
| **rfMRI full corr ICA100 edge 1124 link 28-45** | 0.21 | 15864 |
| **rfMRI part corr ICA100 edge 704 link 15-54** | 0.21 | 15864 |
| **meandepth Morphologist SCall_left** | 0.21 | 18064 |
| **ThickAvg Desikan insula.rh** | 0.21 | 17127 |
| **rfMRI full corr ICA100 edge 1161 link 30-31** | 0.21 | 15864 |
| **rfMRI part corr ICA100 edge 642 link 14-32** | 0.21 | 15864 |
| **rfMRI part corr ICA25 edge 40 link 3-4** | 0.21 | 15864 |
| **rfMRI full corr ICA100 edge 879 link 20-44** | 0.21 | 15864 |
| **rfMRI full corr ICA100 edge 554 link 12-27** | 0.21 | 15864 |
| **rfMRI part corr ICA100 edge 1089 link 27-37** | 0.21 | 15864 |
| **maxdepth Morphologist FColl_left** | 0.21 | 18101 |
| **GM_thickness Morphologist SC_left** | 0.21 | 18092 |
| **rfMRI part corr ICA100 edge 917 link 21-48** | 0.21 | 15864 |
| **rfMRI part corr ICA100 edge 609 link 13-40** | 0.21 | 15864 |
| **GM_thickness Morphologist SPeCinter_right** | 0.21 | 18076 |
| **rfMRI part corr ICA100 edge 1252 link 33-53** | 0.21 | 15864 |
| **opening Morphologist FIPrint2_right** | 0.21 | 13064 |
| **Right-Accumbens-area** | 0.21 | 17127 |
| **hull_junction_length Morphologist SCLPC_left** | 0.21 | 13421 |
| **rfMRI amplitude ICA100 component 48** | 0.21 | 15863 |
| **rfMRI part corr ICA100 edge 662 link 14-52** | 0.21 | 15864 |
| **rfMRI full corr ICA100 edge 330 link 7-28** | 0.21 | 15860 |
| **GrayVol Destrieux S_orbital-H_Shaped.rh** | 0.21 | 17125 |
| **rfMRI full corr ICA100 edge 937 link 22-35** | 0.21 | 15864 |
| **ISOVF Medial_lemniscus_R** | 0.21 | 16527 |
| **rfMRI part corr ICA100 edge 1388 link 41-49** | 0.21 | 15864 |

| | | |
|---|---|---|
| **FA Medial_lemniscus_L** | 0.21 | 16541 |
| **surface Morphologist SLiant_left** | 0.21 | 17423 |
| **rfMRI full corr ICA100 edge 799 link 18-35** | 0.21 | 15864 |
| **GrayVol Destrieux S_temporal_transverse.rh** | 0.21 | 17124 |
| **CC_Mid_Anterior** | 0.21 | 17127 |
| **GM_thickness Morphologist FIPrint2_left** | 0.21 | 16234 |
| **opening Morphologist FIPrint2_left** | 0.22 | 16244 |
| **maxdepth Morphologist SPaint_right** | 0.22 | 18007 |
| **rfMRI full corr ICA100 edge 753 link 17-26** | 0.22 | 15864 |
| **rfMRI full corr ICA100 edge 1197 link 31-43** | 0.22 | 15864 |
| **rfMRI part corr ICA100 edge 828 link 19-28** | 0.22 | 15864 |
| **rfMRI full corr ICA100 edge 408 link 9-13** | 0.22 | 15864 |
| **ICVF Anterior_limb_of_internal_capsule_L** | 0.22 | 16540 |
| **rfMRI part corr ICA100 edge 202 link 4-47** | 0.22 | 15864 |
| **rfMRI full corr ICA25 edge 56 link 3-20** | 0.22 | 15864 |
| **rfMRI part corr ICA100 edge 297 link 6-43** | 0.22 | 15864 |
| **rfMRI full corr ICA100 edge 920 link 21-51** | 0.22 | 15864 |
| **rfMRI full corr ICA100 edge 1179 link 30-49** | 0.22 | 15864 |
| **FA Middle_cerebellar_peduncle** | 0.22 | 16541 |
| **SurfArea Destrieux S_interm_prim-Jensen.rh** | 0.22 | 17127 |
| **rfMRI part corr ICA100 edge 868 link 20-33** | 0.22 | 15864 |
| **rfMRI part corr ICA25 edge 43 link 3-7** | 0.22 | 15864 |
| **rfMRI amplitude ICA100 component 4** | 0.22 | 15863 |
| **rfMRI part corr ICA100 edge 566 link 12-39** | 0.22 | 15864 |
| **SurfArea Desikan superiorparietal.rh** | 0.22 | 17127 |
| **rfMRI full corr ICA100 edge 111 link 3-7** | 0.22 | 15864 |
| **rfMRI part corr ICA100 edge 421 link 9-26** | 0.22 | 15864 |
| **rfMRI full corr ICA100 edge 291 link 6-37** | 0.22 | 15864 |
| **opening Morphologist STsterascant_right** | 0.22 | 17642 |
| **rfMRI part corr ICA100 edge 920 link 21-51** | 0.22 | 15864 |
| **GM_thickness Morphologist SPeCsup_right** | 0.22 | 17265 |
| **rfMRI full corr ICA100 edge 137 link 3-33** | 0.22 | 15864 |
| **rfMRI part corr ICA25 edge 98 link 6-14** | 0.22 | 15864 |
| **meandepth Morphologist STipost_left** | 0.22 | 18096 |

| | | |
|---|---|---|
| **rfMRI part corr ICA100 edge 443 link 9-48** | 0.22 | 15864 |
| **rfMRI part corr ICA100 edge 973 link 23-39** | 0.22 | 15864 |
| **SurfArea Destrieux S_collat_transv_ant.lh** | 0.22 | 17127 |
| **surface Morphologist FPO_right** | 0.22 | 18097 |
| **rfMRI part corr ICA100 edge 285 link 6-31** | 0.22 | 15864 |
| **MD Superior_fronto-occipital_fasciculus-part_of_a** | 0.22 | 16541 |
| **rfMRI full corr ICA100 edge 1426 link 44-51** | 0.22 | 15864 |
| **GrayVol Destrieux S_oc-temp_med_and_Lingual.lh** | 0.22 | 17127 |
| **opening Morphologist STsterascpost_right** | 0.22 | 18048 |
| **rfMRI full corr ICA100 edge 6 link 1-7** | 0.22 | 15864 |
| **rfMRI part corr ICA100 edge 1177 link 30-47** | 0.22 | 15864 |
| **GrayVol Destrieux G_and_S_subcentral.rh** | 0.22 | 17127 |
| **ThickAvg Destrieux S_pericallosal.lh** | 0.22 | 17127 |
| **rfMRI full corr ICA100 edge 679 link 15-29** | 0.22 | 15864 |
| **GM_thickness Morphologist STsterascant_right** | 0.22 | 17630 |
| **ISOVF Cingulum-cingulate_gyrus-R** | 0.22 | 16527 |
| **ThickAvg Desikan posteriorcingulate.rh** | 0.22 | 17127 |
| **ThickAvg Desikan precentral.rh** | 0.22 | 17127 |
| **surface Morphologist SPeCinf_left** | 0.22 | 17300 |
| **OD Anterior_corona_radiata_R** | 0.23 | 16541 |
| **SurfArea Destrieux G_and_S_subcentral.rh** | 0.23 | 17127 |
| **rfMRI part corr ICA100 edge 442 link 9-47** | 0.23 | 15864 |
| **rfMRI part corr ICA100 edge 1134 link 28-55** | 0.23 | 15864 |
| **rfMRI full corr ICA100 edge 359 link 8-10** | 0.23 | 15864 |
| **opening Morphologist SPeCinf_left** | 0.23 | 17300 |
| **GrayVol Destrieux S_temporal_transverse.lh** | 0.23 | 17126 |
| **SurfArea Destrieux S_postcentral.lh** | 0.23 | 17127 |
| **rfMRI part corr ICA25 edge 56 link 3-20** | 0.23 | 15864 |
| **rfMRI full corr ICA100 edge 4 link 1-5** | 0.23 | 15864 |
| **GrayVol Destrieux G_precentral.rh** | 0.23 | 17127 |
| **rfMRI full corr ICA100 edge 386 link 8-37** | 0.23 | 15864 |
| **rfMRI full corr ICA25 edge 61 link 4-8** | 0.23 | 15864 |
| **hull_junction_length Morphologist FIP_left** | 0.23 | 18098 |
| **rfMRI part corr ICA100 edge 1309 link 36-50** | 0.23 | 15864 |

| | | |
|---|---|---|
| **rfMRI full corr ICA100 edge 1099 link 27-47** | 0.23 | 15863 |
| **rfMRI part corr ICA100 edge 850 link 19-50** | 0.23 | 15864 |
| **rfMRI part corr ICA100 edge 1332 link 37-55** | 0.23 | 15864 |
| **SurfArea Destrieux G_temp_sup-Lateral.lh** | 0.23 | 17127 |
| **rfMRI full corr ICA100 edge 7 link 1-8** | 0.23 | 15864 |
| **rfMRI full corr ICA100 edge 845 link 19-45** | 0.23 | 15864 |
| **rfMRI part corr ICA100 edge 583 link 13-14** | 0.23 | 15864 |
| **maxdepth Morphologist SPaint_left** | 0.23 | 18007 |
| **rfMRI full corr ICA100 edge 1135 link 29-30** | 0.23 | 15864 |
| **GrayVol Destrieux S_oc-temp_lat.rh** | 0.23 | 17127 |
| **surface Morphologist FCLp_left** | 0.23 | 18101 |
| **SurfArea Desikan precuneus.rh** | 0.23 | 17127 |
| **rfMRI part corr ICA100 edge 57 link 2-5** | 0.23 | 15864 |
| **rfMRI full corr ICA100 edge 302 link 6-48** | 0.23 | 15864 |
| **rfMRI full corr ICA100 edge 804 link 18-40** | 0.23 | 15864 |
| **rfMRI full corr ICA100 edge 872 link 20-37** | 0.23 | 15864 |
| **ThickAvg Destrieux S_intrapariet_and_P_trans.lh** | 0.23 | 17127 |
| **maxdepth Morphologist SForbitaire_left** | 0.23 | 17520 |
| **rfMRI full corr ICA100 edge 432 link 9-37** | 0.23 | 15864 |
| **ThickAvg Destrieux G_cuneus.lh** | 0.23 | 17127 |
| **rfMRI full corr ICA100 edge 334 link 7-32** | 0.23 | 15864 |
| **rfMRI full corr ICA100 edge 1469 link 49-54** | 0.23 | 15864 |
| **ThickAvg Destrieux G_oc-temp_lat-fusifor.rh** | 0.23 | 17127 |
| **ThickAvg Destrieux G_and_S_cingul-Mid-Post.lh** | 0.23 | 17127 |
| **rfMRI part corr ICA25 edge 130 link 8-19** | 0.23 | 15864 |
| **rfMRI full corr ICA100 edge 1411 link 43-47** | 0.23 | 15864 |
| **rfMRI part corr ICA100 edge 1185 link 30-55** | 0.23 | 15864 |
| **rfMRI full corr ICA100 edge 512 link 11-28** | 0.23 | 15864 |
| **rfMRI part corr ICA100 edge 816 link 18-52** | 0.23 | 15864 |
| **rfMRI full corr ICA100 edge 643 link 14-33** | 0.23 | 15862 |
| **meandepth Morphologist STsterascant_left** | 0.23 | 17507 |
| **rfMRI full corr ICA100 edge 402 link 8-53** | 0.23 | 15864 |
| **rfMRI part corr ICA100 edge 869 link 20-34** | 0.23 | 15864 |
| **rfMRI part corr ICA100 edge 705 link 15-55** | 0.23 | 15864 |

| | | |
|---|---|---|
| **rfMRI part corr ICA100 edge 653 link 14-43** | 0.23 | 15864 |
| **rfMRI full corr ICA25 edge 44 link 3-8** | 0.23 | 15864 |
| **rfMRI full corr ICA100 edge 5 link 1-6** | 0.23 | 15864 |
| **meandepth Morphologist SOTlatint_right** | 0.23 | 15609 |
| **surface Morphologist FCLrscant_left** | 0.24 | 4323 |
| **rfMRI part corr ICA100 edge 781 link 17-54** | 0.24 | 15864 |
| **meandepth Morphologist SPeCmarginal_right** | 0.24 | 17001 |
| **meandepth Morphologist FIP_left** | 0.24 | 18098 |
| **rfMRI part corr ICA100 edge 668 link 15-18** | 0.24 | 15864 |
| **rfMRI full corr ICA100 edge 329 link 7-27** | 0.24 | 15864 |
| **GrayVol Destrieux G_oc-temp_lat-fusifor.rh** | 0.24 | 17127 |
| **rfMRI part corr ICA100 edge 486 link 10-46** | 0.24 | 15864 |
| **rfMRI part corr ICA25 edge 166 link 12-13** | 0.24 | 15864 |
| **maxdepth Morphologist FCalant-ScCal_left** | 0.24 | 18100 |
| **MD Sagittal_stratum-inf_longitidinal_fasci_and_inf** | 0.24 | 16541 |
| **rfMRI part corr ICA100 edge 283 link 6-29** | 0.24 | 15864 |
| **rfMRI part corr ICA100 edge 1052 link 26-28** | 0.24 | 15864 |
| **rfMRI part corr ICA100 edge 1121 link 28-42** | 0.24 | 15864 |
| **ThickAvg Desikan parahippocampal.rh** | 0.24 | 17127 |
| **surface Morphologist FCLrant_left** | 0.24 | 17708 |
| **rfMRI part corr ICA100 edge 1035 link 25-40** | 0.24 | 15864 |
| **maxdepth Morphologist SOlf_left** | 0.24 | 18094 |
| **SurfArea Destrieux S_circular_insula_ant.lh** | 0.24 | 17126 |
| **rfMRI part corr ICA100 edge 632 link 14-22** | 0.24 | 15864 |
| **rfMRI full corr ICA100 edge 945 link 22-43** | 0.24 | 15864 |
| **rfMRI full corr ICA100 edge 1205 link 31-51** | 0.24 | 15864 |
| **rfMRI part corr ICA100 edge 348 link 7-46** | 0.24 | 15864 |
| **rfMRI full corr ICA100 edge 1121 link 28-42** | 0.24 | 15864 |
| **rfMRI part corr ICA100 edge 590 link 13-21** | 0.24 | 15864 |
| **opening Morphologist SFmedian_left** | 0.24 | 18078 |
| **rfMRI part corr ICA100 edge 712 link 16-23** | 0.24 | 15864 |
| **rfMRI part corr ICA100 edge 287 link 6-33** | 0.24 | 15864 |
| **rfMRI amplitude ICA100 component 1** | 0.24 | 15863 |
| **rfMRI part corr ICA100 edge 1438 link 45-53** | 0.24 | 15864 |

| | | |
|---|---|---|
| **maxdepth Morphologist SFint_right** | 0.24 | 18100 |
| **rfMRI part corr ICA100 edge 1179 link 30-49** | 0.24 | 15864 |
| **MO Posterior_limb_of_internal_capsule_L** | 0.24 | 16541 |
| **rfMRI full corr ICA100 edge 915 link 21-46** | 0.24 | 15864 |
| **rfMRI part corr ICA100 edge 1081 link 27-29** | 0.24 | 15864 |
| **rfMRI full corr ICA100 edge 699 link 15-49** | 0.24 | 15864 |
| **FA Splenium_of_corpus_callosum** | 0.24 | 16541 |
| **rfMRI full corr ICA100 edge 1354 link 39-44** | 0.24 | 15863 |
| **rfMRI part corr ICA100 edge 943 link 22-41** | 0.24 | 15864 |
| **rfMRI part corr ICA100 edge 171 link 4-16** | 0.24 | 15864 |
| **hull_junction_length Morphologist FCMpost_left** | 0.24 | 18101 |
| **rfMRI part corr ICA25 edge 79 link 5-10** | 0.24 | 15864 |
| **ICVF Superior_fronto-occipital_fasciculus-part_of** | 0.24 | 16540 |
| **rfMRI full corr ICA100 edge 122 link 3-18** | 0.24 | 15863 |
| **rfMRI part corr ICA100 edge 205 link 4-50** | 0.24 | 15864 |
| **rfMRI part corr ICA100 edge 476 link 10-36** | 0.24 | 15864 |
| **rfMRI full corr ICA100 edge 819 link 18-55** | 0.24 | 15864 |
| **rfMRI full corr ICA100 edge 1350 link 39-40** | 0.24 | 15864 |
| **rfMRI full corr ICA100 edge 467 link 10-27** | 0.24 | 15864 |
| **rfMRI full corr ICA100 edge 1283 link 35-43** | 0.24 | 15864 |
| **rfMRI full corr ICA100 edge 1132 link 28-53** | 0.24 | 15863 |
| **rfMRI part corr ICA100 edge 604 link 13-35** | 0.24 | 15864 |
| **rfMRI part corr ICA100 edge 636 link 14-26** | 0.24 | 15864 |
| **rfMRI full corr ICA100 edge 973 link 23-39** | 0.25 | 15864 |
| **MD Fornix-cres-Stria_terminalis-not_resolved_wit** | 0.25 | 16541 |
| **rfMRI full corr ICA100 edge 1127 link 28-48** | 0.25 | 15864 |
| **rfMRI full corr ICA100 edge 40 link 1-41** | 0.25 | 15864 |
| **rfMRI full corr ICA100 edge 418 link 9-23** | 0.25 | 15864 |
| **GM_thickness Morphologist FCLrscant_right** | 0.25 | 5349 |
| **rfMRI part corr ICA100 edge 1109 link 28-30** | 0.25 | 15864 |
| **rfMRI full corr ICA100 edge 673 link 15-23** | 0.25 | 15864 |
| **rfMRI part corr ICA100 edge 596 link 13-27** | 0.25 | 15864 |
| **rfMRI part corr ICA100 edge 456 link 10-16** | 0.25 | 15864 |
| **rfMRI full corr ICA100 edge 1302 link 36-43** | 0.25 | 15864 |

| | | |
|---|---|---|
| **rfMRI part corr ICA100 edge 415 link 9-20** | 0.25 | 15864 |
| **rfMRI full corr ICA100 edge 866 link 20-31** | 0.25 | 15864 |
| **rfMRI full corr ICA100 edge 680 link 15-30** | 0.25 | 15864 |
| **rfMRI full corr ICA100 edge 233 link 5-28** | 0.25 | 15864 |
| **meandepth Morphologist SCsylvian_left** | 0.25 | 13657 |
| **rfMRI full corr ICA100 edge 425 link 9-30** | 0.25 | 15864 |
| **rfMRI part corr ICA100 edge 477 link 10-37** | 0.25 | 15864 |
| **ICVF Posterior_limb_of_internal_capsule_L** | 0.25 | 16540 |
| **surface Morphologist SForbitaire_left** | 0.25 | 17520 |
| **FA Pontine_crossing_tract-a_part_of_MCP** | 0.25 | 16541 |
| **rfMRI part corr ICA100 edge 334 link 7-32** | 0.25 | 15864 |
| **rfMRI part corr ICA100 edge 1265 link 34-45** | 0.25 | 15864 |
| **ThickAvg Destrieux G_cingul-Post-dorsal.rh** | 0.25 | 17127 |
| **rfMRI full corr ICA25 edge 126 link 8-15** | 0.25 | 15864 |
| **GrayVol Destrieux G_and_S_transv_frontopol.rh** | 0.25 | 17127 |
| **ThickAvg Destrieux S_front_sup.lh** | 0.25 | 17127 |
| **rfMRI full corr ICA100 edge 1016 link 24-51** | 0.25 | 15864 |
| **rfMRI full corr ICA25 edge 10 link 1-11** | 0.25 | 15864 |
| **rfMRI part corr ICA100 edge 804 link 18-40** | 0.25 | 15864 |
| **rfMRI part corr ICA100 edge 958 link 23-24** | 0.25 | 15864 |
| **rfMRI full corr ICA100 edge 1403 link 42-51** | 0.25 | 15864 |
| **GrayVol Destrieux S_cingul-Marginalis.rh** | 0.25 | 17127 |
| **rfMRI part corr ICA100 edge 1084 link 27-32** | 0.25 | 15864 |
| **rfMRI part corr ICA100 edge 354 link 7-52** | 0.25 | 15864 |
| **rfMRI full corr ICA100 edge 299 link 6-45** | 0.25 | 15864 |
| **rfMRI part corr ICA100 edge 798 link 18-34** | 0.25 | 15864 |
| **rfMRI part corr ICA100 edge 335 link 7-33** | 0.25 | 15864 |
| **Left-Caudate** | 0.25 | 17127 |
| **GrayVol Destrieux G_front_sup.lh** | 0.25 | 17127 |
| **rfMRI part corr ICA100 edge 32 link 1-33** | 0.25 | 15864 |
| **GrayVol Destrieux G_postcentral.lh** | 0.25 | 17127 |
| **rfMRI full corr ICA25 edge 62 link 4-9** | 0.25 | 15864 |
| **rfMRI full corr ICA100 edge 1299 link 36-40** | 0.25 | 15864 |
| **rfMRI part corr ICA100 edge 341 link 7-39** | 0.25 | 15864 |

| | | |
|---|---|---|
| **GrayVol Destrieux G_temp_sup-Lateral.rh** | 0.25 | 17127 |
| **rfMRI full corr ICA100 edge 741 link 16-52** | 0.25 | 15864 |
| **rfMRI part corr ICA100 edge 1100 link 27-48** | 0.25 | 15864 |
| **rfMRI part corr ICA100 edge 195 link 4-40** | 0.25 | 15864 |
| **hull_junction_length Morphologist SPasup_left** | 0.25 | 16537 |
| **rfMRI full corr ICA100 edge 1391 link 41-52** | 0.25 | 15864 |
| **rfMRI full corr ICA100 edge 1089 link 27-37** | 0.25 | 15862 |
| **rfMRI full corr ICA100 edge 169 link 4-14** | 0.25 | 15864 |
| **rfMRI full corr ICA100 edge 530 link 11-46** | 0.26 | 15864 |
| **rfMRI part corr ICA25 edge 45 link 3-9** | 0.26 | 15864 |
| **GrayVol Desikan temporalpole.rh** | 0.26 | 17126 |
| **opening Morphologist FPO_left** | 0.26 | 18097 |
| **rfMRI full corr ICA25 edge 208 link 19-20** | 0.26 | 15864 |
| **meandepth Morphologist SPeCmedian_left** | 0.26 | 16792 |
| **rfMRI full corr ICA100 edge 357 link 7-55** | 0.26 | 15864 |
| **rfMRI part corr ICA100 edge 765 link 17-38** | 0.26 | 15864 |
| **FA Inferior_cerebellar_peduncle_R** | 0.26 | 16541 |
| **FA Superior_longitudinal_fasciculus_R** | 0.26 | 16541 |
| **rfMRI part corr ICA100 edge 727 link 16-38** | 0.26 | 15864 |
| **rfMRI part corr ICA25 edge 11 link 1-12** | 0.26 | 15864 |
| **rfMRI part corr ICA100 edge 215 link 5-10** | 0.26 | 15864 |
| **rfMRI full corr ICA100 edge 165 link 4-10** | 0.26 | 15864 |
| **rfMRI part corr ICA100 edge 1409 link 43-45** | 0.26 | 15864 |
| **GrayVol Desikan postcentral.rh** | 0.26 | 17127 |
| **hull_junction_length Morphologist FIPrint1_right** | 0.26 | 17034 |
| **maxdepth Morphologist FCLp_left** | 0.26 | 18101 |
| **rfMRI full corr ICA100 edge 306 link 6-52** | 0.26 | 15863 |
| **opening Morphologist FCMant_left** | 0.26 | 17879 |
| **SurfArea Desikan insula.rh** | 0.26 | 17127 |
| **rfMRI full corr ICA100 edge 221 link 5-16** | 0.26 | 15864 |
| **rfMRI part corr ICA100 edge 1306 link 36-47** | 0.26 | 15864 |
| **rfMRI part corr ICA100 edge 1038 link 25-43** | 0.26 | 15864 |
| **rfMRI full corr ICA100 edge 1436 link 45-51** | 0.26 | 15864 |
| **GrayVol Desikan superiorfrontal.lh** | 0.26 | 17127 |

| | | |
|---|---|---|
| **rfMRI full corr ICA100 edge 62 link 2-10** | 0.26 | 15864 |
| **rfMRI full corr ICA100 edge 67 link 2-15** | 0.26 | 15864 |
| **rfMRI part corr ICA25 edge 34 link 2-16** | 0.26 | 15864 |
| **rfMRI full corr ICA100 edge 1311 link 36-52** | 0.26 | 15864 |
| **rfMRI full corr ICA100 edge 210 link 4-55** | 0.26 | 15864 |
| **rfMRI part corr ICA100 edge 206 link 4-51** | 0.26 | 15864 |
| **rfMRI full corr ICA100 edge 1351 link 39-41** | 0.26 | 15863 |
| **meandepth Morphologist SFmedian_right** | 0.26 | 18080 |
| **rfMRI full corr ICA100 edge 980 link 23-46** | 0.26 | 15864 |
| **ICVF External_capsule_L** | 0.26 | 16540 |
| **surface Morphologist STpol_right** | 0.26 | 18062 |
| **rfMRI part corr ICA100 edge 722 link 16-33** | 0.26 | 15864 |
| **ThickAvg Desikan parsopercularis.rh** | 0.26 | 17127 |
| **opening Morphologist FCLp_right** | 0.26 | 18100 |
| **rfMRI part corr ICA100 edge 922 link 21-53** | 0.26 | 15864 |
| **rfMRI full corr ICA100 edge 1446 link 46-52** | 0.26 | 15864 |
| **rfMRI full corr ICA100 edge 176 link 4-21** | 0.26 | 15864 |
| **rfMRI full corr ICA100 edge 1451 link 47-49** | 0.26 | 15864 |
| **maxdepth Morphologist SPeCinter_left** | 0.26 | 18028 |
| **rfMRI full corr ICA100 edge 411 link 9-16** | 0.26 | 15864 |
| **rfMRI part corr ICA100 edge 59 link 2-7** | 0.26 | 15864 |
| **rfMRI full corr ICA100 edge 714 link 16-25** | 0.26 | 15863 |
| **rfMRI full corr ICA25 edge 151 link 10-17** | 0.26 | 15864 |
| **rfMRI full corr ICA100 edge 525 link 11-41** | 0.26 | 15864 |
| **rfMRI full corr ICA100 edge 658 link 14-48** | 0.26 | 15863 |
| **rfMRI full corr ICA25 edge 73 link 4-20** | 0.26 | 15864 |
| **rfMRI part corr ICA100 edge 252 link 5-47** | 0.26 | 15864 |
| **rfMRI part corr ICA100 edge 971 link 23-37** | 0.26 | 15864 |
| **GrayVol Desikan lateraloccipital.rh** | 0.26 | 17127 |
| **rfMRI part corr ICA100 edge 1242 link 33-43** | 0.26 | 15864 |
| **rfMRI full corr ICA100 edge 936 link 22-34** | 0.26 | 15864 |
| **FA Posterior_limb_of_internal_capsule_L** | 0.26 | 16541 |
| **hull_junction_length Morphologist SPeCinf_left** | 0.26 | 17300 |
| **MD Anterior_limb_of_internal_capsule_L** | 0.26 | 16541 |

| | | |
|---|---|---|
| **rfMRI part corr ICA100 edge 892 link 21-23** | 0.26 | 15864 |
| **rfMRI full corr ICA100 edge 1377 link 40-52** | 0.26 | 15864 |
| **SurfArea Destrieux Lat_Fis-ant-Vertical.lh** | 0.26 | 17122 |
| **rfMRI full corr ICA100 edge 1407 link 42-55** | 0.26 | 15864 |
| **SurfArea Destrieux S_orbital_lateral.lh** | 0.27 | 17126 |
| **rfMRI full corr ICA25 edge 155 link 10-21** | 0.27 | 15864 |
| **rfMRI part corr ICA100 edge 165 link 4-10** | 0.27 | 15864 |
| **rfMRI amplitude ICA100 component 21** | 0.27 | 15863 |
| **rfMRI part corr ICA100 edge 1246 link 33-47** | 0.27 | 15864 |
| **rfMRI full corr ICA100 edge 1024 link 25-29** | 0.27 | 15858 |
| **rfMRI full corr ICA100 edge 97 link 2-45** | 0.27 | 15864 |
| **rfMRI part corr ICA100 edge 1239 link 33-40** | 0.27 | 15864 |
| **rfMRI full corr ICA100 edge 416 link 9-21** | 0.27 | 15863 |
| **rfMRI part corr ICA100 edge 196 link 4-41** | 0.27 | 15864 |
| **rfMRI part corr ICA25 edge 203 link 17-20** | 0.27 | 15864 |
| **rfMRI full corr ICA100 edge 805 link 18-41** | 0.27 | 15864 |
| **maxdepth Morphologist SCsylvian_left** | 0.27 | 13814 |
| **ThickAvg Destrieux G_pariet_inf-Angular.lh** | 0.27 | 17127 |
| **rfMRI part corr ICA100 edge 981 link 23-47** | 0.27 | 15864 |
| **meandepth Morphologist SPeCinf_right** | 0.27 | 16568 |
| **rfMRI part corr ICA100 edge 502 link 11-18** | 0.27 | 15864 |
| **rfMRI full corr ICA25 edge 167 link 12-14** | 0.27 | 15864 |
| **rfMRI part corr ICA100 edge 700 link 15-50** | 0.27 | 15864 |
| **rfMRI part corr ICA100 edge 216 link 5-11** | 0.27 | 15864 |
| **rfMRI part corr ICA100 edge 1362 link 39-52** | 0.27 | 15864 |
| **rfMRI part corr ICA100 edge 469 link 10-29** | 0.27 | 15864 |
| **rfMRI full corr ICA100 edge 167 link 4-12** | 0.27 | 15864 |
| **rfMRI full corr ICA100 edge 842 link 19-42** | 0.27 | 15864 |
| **rfMRI full corr ICA100 edge 718 link 16-29** | 0.27 | 15862 |
| **rfMRI full corr ICA100 edge 1074 link 26-50** | 0.27 | 15863 |
| **rfMRI full corr ICA25 edge 182 link 13-21** | 0.27 | 15864 |
| **rfMRI part corr ICA100 edge 1296 link 36-37** | 0.27 | 15864 |
| **SurfArea Destrieux G_temp_sup-G_T_transv.rh** | 0.27 | 17125 |
| **rfMRI full corr ICA100 edge 224 link 5-19** | 0.27 | 15863 |

| | | |
|---|---|---|
| **rfMRI part corr ICA100 edge 164 link 4-9** | 0.27 | 15864 |
| **rfMRI full corr ICA100 edge 827 link 19-27** | 0.27 | 15864 |
| **rfMRI part corr ICA100 edge 951 link 22-49** | 0.27 | 15864 |
| **rfMRI part corr ICA100 edge 1363 link 39-53** | 0.27 | 15864 |
| **rfMRI full corr ICA100 edge 531 link 11-47** | 0.27 | 15864 |
| **meandepth Morphologist SOTlatmed_right** | 0.27 | 17207 |
| **rfMRI part corr ICA100 edge 1022 link 25-27** | 0.27 | 15864 |
| **rfMRI full corr ICA100 edge 43 link 1-44** | 0.27 | 15864 |
| **ThickAvg Destrieux S_circular_insula_sup.lh** | 0.27 | 17126 |
| **GrayVol Destrieux S_calcarine.lh** | 0.27 | 17127 |
| **rfMRI part corr ICA100 edge 907 link 21-38** | 0.27 | 15864 |
| **rfMRI part corr ICA100 edge 841 link 19-41** | 0.27 | 15864 |
| **rfMRI full corr ICA100 edge 1314 link 36-55** | 0.27 | 15864 |
| **rfMRI part corr ICA100 edge 291 link 6-37** | 0.27 | 15864 |
| **rfMRI full corr ICA100 edge 1456 link 47-54** | 0.27 | 15864 |
| **rfMRI part corr ICA100 edge 986 link 23-52** | 0.27 | 15864 |
| **rfMRI part corr ICA100 edge 350 link 7-48** | 0.27 | 15864 |
| **hull_junction_length Morphologist FCLrasc_left** | 0.27 | 17784 |
| **rfMRI full corr ICA100 edge 50 link 1-51** | 0.27 | 15864 |
| **rfMRI full corr ICA100 edge 180 link 4-25** | 0.27 | 15864 |
| **rfMRI full corr ICA100 edge 277 link 6-23** | 0.27 | 15862 |
| **rfMRI part corr ICA100 edge 79 link 2-27** | 0.27 | 15864 |
| **rfMRI part corr ICA100 edge 310 link 7-8** | 0.27 | 15864 |
| **rfMRI part corr ICA100 edge 1064 link 26-40** | 0.27 | 15864 |
| **rfMRI part corr ICA100 edge 808 link 18-44** | 0.27 | 15864 |
| **rfMRI full corr ICA25 edge 95 link 6-11** | 0.27 | 15864 |
| **meandepth Morphologist STsterascpost_left** | 0.27 | 17892 |
| **GrayVol Destrieux S_pericallosal.rh** | 0.28 | 17127 |
| **rfMRI part corr ICA100 edge 356 link 7-54** | 0.28 | 15864 |
| **rfMRI full corr ICA100 edge 1461 link 48-52** | 0.28 | 15864 |
| **rfMRI full corr ICA100 edge 267 link 6-13** | 0.28 | 15864 |
| **rfMRI full corr ICA100 edge 991 link 24-26** | 0.28 | 15864 |
| **rfMRI full corr ICA100 edge 796 link 18-32** | 0.28 | 15864 |
| **rfMRI part corr ICA100 edge 267 link 6-13** | 0.28 | 15864 |

| | | |
|---|---|---|
| **rfMRI part corr ICA100 edge 109 link 3-5** | 0.28 | 15864 |
| **GrayVol Destrieux S_subparietal.lh** | 0.28 | 17125 |
| **rfMRI full corr ICA100 edge 728 link 16-39** | 0.28 | 15863 |
| **rfMRI part corr ICA100 edge 1336 link 38-42** | 0.28 | 15864 |
| **rfMRI full corr ICA25 edge 205 link 18-19** | 0.28 | 15864 |
| **ThickAvg Desikan superiorfrontal.lh** | 0.28 | 17127 |
| **rfMRI full corr ICA100 edge 783 link 18-19** | 0.28 | 15864 |
| **rfMRI full corr ICA100 edge 626 link 14-16** | 0.28 | 15863 |
| **rfMRI part corr ICA100 edge 846 link 19-46** | 0.28 | 15864 |
| **rfMRI part corr ICA100 edge 522 link 11-38** | 0.28 | 15864 |
| **rfMRI full corr ICA100 edge 874 link 20-39** | 0.28 | 15864 |
| **hull_junction_length Morphologist FCLrdiag_left** | 0.28 | 8551 |
| **rfMRI part corr ICA100 edge 410 link 9-15** | 0.28 | 15864 |
| **SurfArea Destrieux G_insular_short.rh** | 0.28 | 17125 |
| **rfMRI full corr ICA100 edge 490 link 10-50** | 0.28 | 15864 |
| **ThickAvg Destrieux S_cingul-Marginalis.lh** | 0.28 | 17127 |
| **rfMRI part corr ICA100 edge 1454 link 47-52** | 0.28 | 15864 |
| **rfMRI full corr ICA100 edge 1173 link 30-43** | 0.28 | 15864 |
| **rfMRI part corr ICA100 edge 46 link 1-47** | 0.28 | 15864 |
| **meandepth Morphologist SFinfant_right** | 0.28 | 18042 |
| **meandepth Morphologist FCLrscpost_right** | 0.28 | 16029 |
| **rfMRI full corr ICA25 edge 149 link 10-15** | 0.28 | 15864 |
| **GM_thickness Morphologist STs_right** | 0.28 | 18092 |
| **ThickAvg Desikan inferiorparietal.rh** | 0.28 | 17127 |
| **rfMRI full corr ICA100 edge 565 link 12-38** | 0.28 | 15864 |
| **FA Retrolenticular_part_of_internal_capsule_L** | 0.28 | 16541 |
| **SurfArea Desikan rostralmiddlefrontal.rh** | 0.28 | 17127 |
| **meandepth Morphologist FIP_right** | 0.28 | 18097 |
| **rfMRI part corr ICA100 edge 1101 link 27-49** | 0.28 | 15864 |
| **rfMRI full corr ICA25 edge 15 link 1-16** | 0.28 | 15864 |
| **MO Corticospinal_tract_R** | 0.28 | 16541 |
| **rfMRI part corr ICA100 edge 810 link 18-46** | 0.28 | 15864 |
| **rfMRI part corr ICA100 edge 1453 link 47-51** | 0.28 | 15864 |
| **rfMRI full corr ICA100 edge 229 link 5-24** | 0.28 | 15864 |

| | | |
|---|---|---|
| **rfMRI full corr ICA100 edge 200 link 4-45** | 0.28 | 15860 |
| **rfMRI part corr ICA100 edge 718 link 16-29** | 0.28 | 15864 |
| **rfMRI part corr ICA100 edge 968 link 23-34** | 0.28 | 15864 |
| **hull_junction_length Morphologist FCLa_right** | 0.28 | 18097 |
| **rfMRI part corr ICA100 edge 861 link 20-26** | 0.28 | 15864 |
| **rfMRI part corr ICA100 edge 1192 link 31-38** | 0.28 | 15864 |
| **GM_thickness Morphologist SPeCmarginal_right** | 0.28 | 16999 |
| **rfMRI full corr ICA100 edge 305 link 6-51** | 0.28 | 15861 |
| **rfMRI part corr ICA25 edge 176 link 13-15** | 0.28 | 15864 |
| **rfMRI full corr ICA100 edge 850 link 19-50** | 0.28 | 15864 |
| **meandepth Morphologist FIPrint1_left** | 0.28 | 17276 |
| **rfMRI part corr ICA100 edge 1046 link 25-51** | 0.28 | 15864 |
| **rfMRI full corr ICA100 edge 1367 link 40-42** | 0.28 | 15864 |
| **rfMRI full corr ICA100 edge 401 link 8-52** | 0.28 | 15864 |
| **GM_thickness Morphologist SOTlatant_left** | 0.28 | 18066 |
| **rfMRI full corr ICA100 edge 952 link 22-50** | 0.28 | 15864 |
| **CSF** | 0.28 | 17127 |
| **rfMRI full corr ICA25 edge 184 link 14-16** | 0.28 | 15864 |
| **GrayVol Destrieux G_subcallosal.lh** | 0.28 | 17125 |
| **rfMRI part corr ICA100 edge 796 link 18-32** | 0.28 | 15864 |
| **rfMRI full corr ICA25 edge 127 link 8-16** | 0.28 | 15864 |
| **rfMRI part corr ICA100 edge 614 link 13-45** | 0.28 | 15864 |
| **CC_Posterior** | 0.29 | 17127 |
| **GrayVol Desikan frontalpole.lh** | 0.29 | 17127 |
| **rfMRI part corr ICA100 edge 771 link 17-44** | 0.29 | 15864 |
| **rfMRI part corr ICA25 edge 64 link 4-11** | 0.29 | 15864 |
| **rfMRI full corr ICA100 edge 1369 link 40-44** | 0.29 | 15864 |
| **rfMRI part corr ICA25 edge 114 link 7-16** | 0.29 | 15864 |
| **SurfArea Destrieux G_temp_sup-Plan_polar.rh** | 0.29 | 17126 |
| **rfMRI full corr ICA100 edge 528 link 11-44** | 0.29 | 15863 |
| **ThickAvg Destrieux S_orbital_lateral.rh** | 0.29 | 17126 |
| **rfMRI part corr ICA100 edge 678 link 15-28** | 0.29 | 15864 |
| **rfMRI full corr ICA100 edge 321 link 7-19** | 0.29 | 15860 |
| **rfMRI part corr ICA100 edge 641 link 14-31** | 0.29 | 15864 |

| | | |
|---|---|---|
| **GM_thickness Morphologist FCLp_left** | 0.29 | 18092 |
| **rfMRI full corr ICA25 edge 114 link 7-16** | 0.29 | 15864 |
| **rfMRI part corr ICA25 edge 36 link 2-18** | 0.29 | 15864 |
| **maxdepth Morphologist SCu_left** | 0.29 | 18058 |
| **rfMRI full corr ICA100 edge 639 link 14-29** | 0.29 | 15860 |
| **MO Body_of_corpus_callosum** | 0.29 | 16541 |
| **rfMRI full corr ICA100 edge 1388 link 41-49** | 0.29 | 15864 |
| **rfMRI part corr ICA100 edge 691 link 15-41** | 0.29 | 15864 |
| **GrayVol Destrieux S_temporal_inf.rh** | 0.29 | 17127 |
| **rfMRI part corr ICA100 edge 1027 link 25-32** | 0.29 | 15864 |
| **rfMRI full corr ICA25 edge 123 link 8-12** | 0.29 | 15864 |
| **rfMRI part corr ICA100 edge 597 link 13-28** | 0.29 | 15864 |
| **rfMRI part corr ICA100 edge 2 link 1-3** | 0.29 | 15864 |
| **meandepth Morphologist FCLrscant_right** | 0.29 | 5320 |
| **rfMRI part corr ICA100 edge 457 link 10-17** | 0.29 | 15864 |
| **rfMRI full corr ICA100 edge 585 link 13-16** | 0.29 | 15863 |
| **rfMRI part corr ICA100 edge 959 link 23-25** | 0.29 | 15864 |
| **rfMRI part corr ICA100 edge 313 link 7-11** | 0.29 | 15864 |
| **rfMRI part corr ICA100 edge 254 link 5-49** | 0.29 | 15864 |
| **rfMRI full corr ICA100 edge 28 link 1-29** | 0.29 | 15864 |
| **GM_thickness Morphologist FCMpost_right** | 0.29 | 18093 |
| **hull_junction_length Morphologist SPat_left** | 0.29 | 13931 |
| **rfMRI part corr ICA100 edge 40 link 1-41** | 0.29 | 15864 |
| **rfMRI part corr ICA100 edge 991 link 24-26** | 0.29 | 15864 |
| **OD Retrolenticular_part_of_internal_capsule_L** | 0.29 | 16541 |
| **rfMRI part corr ICA100 edge 1464 link 48-55** | 0.29 | 15864 |
| **rfMRI full corr ICA100 edge 1055 link 26-31** | 0.29 | 15864 |
| **rfMRI part corr ICA25 edge 210 link 20-21** | 0.29 | 15864 |
| **rfMRI full corr ICA100 edge 560 link 12-33** | 0.29 | 15864 |
| **SurfArea Destrieux S_circular_insula_sup.rh** | 0.29 | 17126 |
| **rfMRI part corr ICA100 edge 1248 link 33-49** | 0.29 | 15864 |
| **rfMRI full corr ICA100 edge 1233 link 33-34** | 0.29 | 15864 |
| **GrayVol Destrieux G_rectus.rh** | 0.29 | 17126 |
| **rfMRI full corr ICA100 edge 1475 link 50-55** | 0.29 | 15864 |

| | | |
|---|---|---|
| **rfMRI part corr ICA100 edge 1209 link 31-55** | 0.29 | 15864 |
| **rfMRI full corr ICA25 edge 39 link 2-21** | 0.29 | 15864 |
| **rfMRI full corr ICA100 edge 817 link 18-53** | 0.29 | 15864 |
| **hull_junction_length Morphologist FIPrint2_left** | 0.29 | 16244 |
| **rfMRI full corr ICA100 edge 536 link 11-52** | 0.29 | 15863 |
| **rfMRI part corr ICA100 edge 359 link 8-10** | 0.29 | 15864 |
| **rfMRI part corr ICA100 edge 441 link 9-46** | 0.29 | 15864 |
| **rfMRI full corr ICA100 edge 950 link 22-48** | 0.29 | 15864 |
| **rfMRI full corr ICA100 edge 1443 link 46-49** | 0.3 | 15864 |
| **rfMRI part corr ICA100 edge 552 link 12-25** | 0.3 | 15864 |
| **rfMRI part corr ICA25 edge 14 link 1-15** | 0.3 | 15864 |
| **rfMRI full corr ICA100 edge 1427 link 44-52** | 0.3 | 15864 |
| **rfMRI part corr ICA100 edge 1379 link 40-54** | 0.3 | 15864 |
| **rfMRI full corr ICA100 edge 1453 link 47-51** | 0.3 | 15864 |
| **GrayVol Desikan rostralmiddlefrontal.rh** | 0.3 | 17127 |
| **rfMRI full corr ICA100 edge 1093 link 27-41** | 0.3 | 15864 |
| **rfMRI part corr ICA100 edge 331 link 7-29** | 0.3 | 15864 |
| **rfMRI part corr ICA100 edge 872 link 20-37** | 0.3 | 15864 |
| **rfMRI full corr ICA100 edge 79 link 2-27** | 0.3 | 15864 |
| **hull_junction_length Morphologist STipost_left** | 0.3 | 18096 |
| **rfMRI full corr ICA100 edge 884 link 20-49** | 0.3 | 15864 |
| **rfMRI part corr ICA25 edge 190 link 15-16** | 0.3 | 15864 |
| **rfMRI full corr ICA100 edge 1366 link 40-41** | 0.3 | 15864 |
| **rfMRI full corr ICA25 edge 200 link 16-21** | 0.3 | 15863 |
| **rfMRI full corr ICA100 edge 435 link 9-40** | 0.3 | 15863 |
| **opening Morphologist SpC_left** | 0.3 | 17043 |
| **meandepth Morphologist SOr_right** | 0.3 | 18097 |
| **rfMRI part corr ICA100 edge 1108 link 28-29** | 0.3 | 15864 |
| **rfMRI part corr ICA100 edge 564 link 12-37** | 0.3 | 15864 |
| **rfMRI part corr ICA100 edge 77 link 2-25** | 0.3 | 15864 |
| **rfMRI part corr ICA100 edge 298 link 6-44** | 0.3 | 15864 |
| **ISOVF Retrolenticular_part_of_internal_capsule_R** | 0.3 | 16527 |
| **rfMRI full corr ICA100 edge 630 link 14-20** | 0.3 | 15863 |
| **rfMRI part corr ICA100 edge 217 link 5-12** | 0.3 | 15864 |

| | | |
|---|---|---|
| **ThickAvg Destrieux G_precentral.lh** | 0.3 | 17127 |
| **ICVF External_capsule_R** | 0.3 | 16540 |
| **rfMRI part corr ICA25 edge 177 link 13-16** | 0.3 | 15864 |
| **rfMRI full corr ICA100 edge 1176 link 30-46** | 0.3 | 15864 |
| **maxdepth Morphologist STiant_right** | 0.3 | 18100 |
| **SurfArea Desikan entorhinal.rh** | 0.3 | 17125 |
| **rfMRI part corr ICA100 edge 90 link 2-38** | 0.3 | 15864 |
| **opening Morphologist STipost_left** | 0.3 | 18096 |
| **rfMRI part corr ICA100 edge 183 link 4-28** | 0.3 | 15864 |
| **rfMRI part corr ICA100 edge 919 link 21-50** | 0.3 | 15864 |
| **rfMRI full corr ICA100 edge 96 link 2-44** | 0.3 | 15862 |
| **SurfArea Destrieux G_and_S_occipital_inf.lh** | 0.3 | 17127 |
| **ThickAvg Destrieux Lat_Fis-ant-Vertical.rh** | 0.3 | 17114 |
| **ThickAvg Desikan parsopercularis.lh** | 0.3 | 17127 |
| **rfMRI part corr ICA100 edge 136 link 3-32** | 0.3 | 15864 |
| **GrayVol Destrieux G_temp_sup-G_T_transv.rh** | 0.3 | 17125 |
| **rfMRI full corr ICA100 edge 193 link 4-38** | 0.3 | 15864 |
| **opening Morphologist SFmedian_right** | 0.3 | 18080 |
| **rfMRI part corr ICA100 edge 1399 link 42-47** | 0.3 | 15864 |
| **rfMRI part corr ICA100 edge 311 link 7-9** | 0.3 | 15864 |
| **rfMRI full corr ICA100 edge 335 link 7-33** | 0.3 | 15863 |
| **GrayVol Destrieux S_orbital-H_Shaped.lh** | 0.3 | 17127 |
| **opening Morphologist SC_left** | 0.3 | 18101 |
| **SurfArea Destrieux G_and_S_paracentral.lh** | 0.3 | 17127 |
| **rfMRI full corr ICA100 edge 382 link 8-33** | 0.3 | 15859 |
| **rfMRI part corr ICA100 edge 395 link 8-46** | 0.31 | 15864 |
| **rfMRI full corr ICA100 edge 211 link 5-6** | 0.31 | 15864 |
| **rfMRI part corr ICA100 edge 247 link 5-42** | 0.31 | 15864 |
| **rfMRI part corr ICA100 edge 1393 link 41-54** | 0.31 | 15864 |
| **GrayVol Desikan posteriorcingulate.lh** | 0.31 | 17126 |
| **ThickAvg Desikan superiortemporal.lh** | 0.31 | 17127 |
| **rfMRI full corr ICA100 edge 501 link 11-17** | 0.31 | 15860 |
| **rfMRI full corr ICA100 edge 388 link 8-39** | 0.31 | 15858 |
| **rfMRI full corr ICA100 edge 492 link 10-52** | 0.31 | 15864 |

| | | |
|---|---|---|
| **rfMRI full corr ICA100 edge 72 link 2-20** | 0.31 | 15864 |
| **GrayVol Destrieux G_and_S_subcentral.lh** | 0.31 | 17127 |
| **GM_thickness Morphologist SPat_right** | 0.31 | 16748 |
| **rfMRI part corr ICA100 edge 571 link 12-44** | 0.31 | 15864 |
| **rfMRI part corr ICA100 edge 1474 link 50-54** | 0.31 | 15864 |
| **rfMRI part corr ICA100 edge 132 link 3-28** | 0.31 | 15864 |
| **rfMRI part corr ICA100 edge 1218 link 32-41** | 0.31 | 15864 |
| **rfMRI full corr ICA100 edge 949 link 22-47** | 0.31 | 15864 |
| **ISOVF Posterior_limb_of_internal_capsule_L** | 0.31 | 16527 |
| **opening Morphologist SFinter_left** | 0.31 | 18101 |
| **ThickAvg Desikan parsorbitalis.lh** | 0.31 | 17126 |
| **rfMRI part corr ICA25 edge 175 link 13-14** | 0.31 | 15864 |
| **rfMRI part corr ICA100 edge 690 link 15-40** | 0.31 | 15864 |
| **rfMRI full corr ICA25 edge 156 link 11-12** | 0.31 | 15864 |
| **rfMRI part corr ICA100 edge 656 link 14-46** | 0.31 | 15864 |
| **rfMRI part corr ICA100 edge 340 link 7-38** | 0.31 | 15864 |
| **rfMRI full corr ICA100 edge 415 link 9-20** | 0.31 | 15864 |
| **rfMRI full corr ICA100 edge 223 link 5-18** | 0.31 | 15863 |
| **hull_junction_length Morphologist SRh_right** | 0.31 | 17623 |
| **rfMRI full corr ICA25 edge 29 link 2-11** | 0.31 | 15864 |
| **rfMRI part corr ICA100 edge 16 link 1-17** | 0.31 | 15864 |
| **opening Morphologist SFint_right** | 0.31 | 18100 |
| **rfMRI full corr ICA100 edge 644 link 14-34** | 0.31 | 15864 |
| **rfMRI part corr ICA100 edge 487 link 10-47** | 0.31 | 15864 |
| **rfMRI part corr ICA100 edge 1078 link 26-54** | 0.31 | 15864 |
| **rfMRI full corr ICA25 edge 204 link 17-21** | 0.31 | 15864 |
| **rfMRI full corr ICA100 edge 322 link 7-20** | 0.31 | 15864 |
| **rfMRI full corr ICA100 edge 876 link 20-41** | 0.31 | 15864 |
| **rfMRI part corr ICA100 edge 1369 link 40-44** | 0.31 | 15864 |
| **ISOVF Sagittal_stratum-inf_longitidinal_fasci_and_** | 0.31 | 16527 |
| **rfMRI full corr ICA100 edge 218 link 5-13** | 0.31 | 15864 |
| **rfMRI full corr ICA100 edge 240 link 5-35** | 0.31 | 15863 |
| **ThickAvg Destrieux Lat_Fis-ant-Horizont.lh** | 0.31 | 17126 |
| **OD Cingulum-hippocampus-L** | 0.31 | 16541 |

| | | |
|---|---|---|
| **ThickAvg Destrieux G_and_S_transv_frontopol.lh** | 0.31 | 17127 |
| **rfMRI part corr ICA100 edge 730 link 16-41** | 0.31 | 15864 |
| **rfMRI full corr ICA100 edge 986 link 23-52** | 0.31 | 15864 |
| **rfMRI part corr ICA100 edge 822 link 19-22** | 0.31 | 15864 |
| **maxdepth Morphologist STpol_right** | 0.31 | 18062 |
| **meandepth Morphologist STpol_left** | 0.31 | 18076 |
| **rfMRI part corr ICA100 edge 716 link 16-27** | 0.31 | 15864 |
| **GM_thickness Morphologist SOTlatmed_right** | 0.31 | 17211 |
| **GM_thickness Morphologist SCLPC_left** | 0.31 | 13412 |
| **rfMRI full corr ICA100 edge 1084 link 27-32** | 0.31 | 15864 |
| **rfMRI part corr ICA100 edge 9 link 1-10** | 0.32 | 15864 |
| **rfMRI full corr ICA25 edge 124 link 8-13** | 0.32 | 15864 |
| **rfMRI part corr ICA100 edge 602 link 13-33** | 0.32 | 15864 |
| **maxdepth Morphologist SFinfant_right** | 0.32 | 18042 |
| **maxdepth Morphologist SCall_right** | 0.32 | 18086 |
| **rfMRI part corr ICA100 edge 762 link 17-35** | 0.32 | 15864 |
| **rfMRI full corr ICA100 edge 1227 link 32-50** | 0.32 | 15864 |
| **rfMRI part corr ICA25 edge 96 link 6-12** | 0.32 | 15864 |
| **rfMRI part corr ICA25 edge 184 link 14-16** | 0.32 | 15864 |
| **meandepth Morphologist SpC_left** | 0.32 | 17039 |
| **GrayVol Destrieux G_and_S_frontomargin.rh** | 0.32 | 17126 |
| **rfMRI amplitude ICA100 component 30** | 0.32 | 15863 |
| **rfMRI full corr ICA100 edge 520 link 11-36** | 0.32 | 15858 |
| **rfMRI amplitude ICA25 component 19** | 0.32 | 15863 |
| **rfMRI amplitude ICA100 component 18** | 0.32 | 15863 |
| **rfMRI full corr ICA100 edge 1113 link 28-34** | 0.32 | 15864 |
| **opening Morphologist SPeCmedian_left** | 0.32 | 16802 |
| **ThickAvg Desikan supramarginal.lh** | 0.32 | 17127 |
| **rfMRI part corr ICA100 edge 394 link 8-45** | 0.32 | 15864 |
| **rfMRI part corr ICA100 edge 554 link 12-27** | 0.32 | 15864 |
| **rfMRI part corr ICA100 edge 1257 link 34-37** | 0.32 | 15864 |
| **meandepth Morphologist FCLrdiag_left** | 0.32 | 8388 |
| **rfMRI full corr ICA100 edge 281 link 6-27** | 0.32 | 15864 |
| **rfMRI full corr ICA100 edge 725 link 16-36** | 0.32 | 15862 |

| | | |
|---|---|---|
| **rfMRI part corr ICA100 edge 507 link 11-23** | 0.32 | 15864 |
| **rfMRI part corr ICA100 edge 1142 link 29-37** | 0.32 | 15864 |
| **rfMRI part corr ICA100 edge 220 link 5-15** | 0.32 | 15864 |
| **rfMRI full corr ICA100 edge 667 link 15-17** | 0.32 | 15864 |
| **rfMRI part corr ICA100 edge 1230 link 32-53** | 0.32 | 15864 |
| **GM_thickness Morphologist FPO_right** | 0.32 | 18090 |
| **rfMRI part corr ICA25 edge 129 link 8-18** | 0.32 | 15864 |
| **GrayVol Destrieux S_orbital_med-olfact.lh** | 0.32 | 17126 |
| **rfMRI part corr ICA100 edge 1418 link 43-54** | 0.32 | 15864 |
| **OD Sagittal_stratum-inf_longitidinal_fasci_and_inf** | 0.32 | 16541 |
| **rfMRI full corr ICA100 edge 889 link 20-54** | 0.32 | 15864 |
| **rfMRI full corr ICA25 edge 109 link 7-11** | 0.32 | 15864 |
| **rfMRI part corr ICA25 edge 194 link 15-20** | 0.32 | 15864 |
| **rfMRI part corr ICA100 edge 234 link 5-29** | 0.32 | 15864 |
| **GrayVol Desikan fusiform.rh** | 0.32 | 17127 |
| **rfMRI amplitude ICA100 component 31** | 0.32 | 15863 |
| **rfMRI part corr ICA100 edge 471 link 10-31** | 0.32 | 15864 |
| **rfMRI full corr ICA100 edge 485 link 10-45** | 0.32 | 15864 |
| **ThickAvg Desikan paracentral.rh** | 0.32 | 17127 |
| **rfMRI full corr ICA25 edge 48 link 3-12** | 0.32 | 15864 |
| **ISOVF Superior_longitudinal_fasciculus_R** | 0.32 | 16527 |
| **rfMRI part corr ICA100 edge 376 link 8-27** | 0.32 | 15864 |
| **rfMRI part corr ICA25 edge 55 link 3-19** | 0.32 | 15864 |
| **rfMRI full corr ICA100 edge 319 link 7-17** | 0.32 | 15863 |
| **MD Cingulum-hippocampus-R** | 0.32 | 16541 |
| **rfMRI full corr ICA100 edge 160 link 4-5** | 0.32 | 15864 |
| **rfMRI part corr ICA100 edge 1305 link 36-46** | 0.33 | 15864 |
| **MD Retrolenticular_part_of_internal_capsule_R** | 0.33 | 16541 |
| **maxdepth Morphologist FCMant_right** | 0.33 | 17828 |
| **rfMRI part corr ICA100 edge 104 link 2-52** | 0.33 | 15864 |
| **rfMRI full corr ICA100 edge 638 link 14-28** | 0.33 | 15864 |
| **rfMRI full corr ICA100 edge 1071 link 26-47** | 0.33 | 15864 |
| **rfMRI full corr ICA100 edge 1184 link 30-54** | 0.33 | 15864 |
| **rfMRI full corr ICA100 edge 194 link 4-39** | 0.33 | 15864 |

| | | |
|---|---|---|
| **rfMRI part corr ICA100 edge 399 link 8-50** | 0.33 | 15864 |
| **GrayVol Destrieux Pole_occipital.rh** | 0.33 | 17127 |
| **rfMRI part corr ICA100 edge 803 link 18-39** | 0.33 | 15864 |
| **rfMRI part corr ICA25 edge 50 link 3-14** | 0.33 | 15864 |
| **rfMRI part corr ICA100 edge 239 link 5-34** | 0.33 | 15864 |
| **rfMRI full corr ICA100 edge 698 link 15-48** | 0.33 | 15863 |
| **rfMRI full corr ICA100 edge 225 link 5-20** | 0.33 | 15864 |
| **rfMRI full corr ICA100 edge 1282 link 35-42** | 0.33 | 15864 |
| **rfMRI part corr ICA100 edge 475 link 10-35** | 0.33 | 15864 |
| **rfMRI part corr ICA100 edge 125 link 3-21** | 0.33 | 15864 |
| **rfMRI part corr ICA100 edge 925 link 22-23** | 0.33 | 15864 |
| **rfMRI full corr ICA100 edge 1373 link 40-48** | 0.33 | 15864 |
| **ThickAvg Destrieux G_front_inf-Opercular.rh** | 0.33 | 17127 |
| **rfMRI full corr ICA100 edge 279 link 6-25** | 0.33 | 15864 |
| **rfMRI full corr ICA100 edge 466 link 10-26** | 0.33 | 15864 |
| **rfMRI full corr ICA100 edge 494 link 10-54** | 0.33 | 15863 |
| **rfMRI full corr ICA100 edge 549 link 12-22** | 0.33 | 15862 |
| **rfMRI full corr ICA100 edge 645 link 14-35** | 0.33 | 15864 |
| **rfMRI part corr ICA100 edge 811 link 18-47** | 0.33 | 15864 |
| **rfMRI part corr ICA25 edge 164 link 11-20** | 0.33 | 15864 |
| **ICVF Posterior_limb_of_internal_capsule_R** | 0.33 | 16540 |
| **rfMRI part corr ICA100 edge 1160 link 29-55** | 0.33 | 15864 |
| **rfMRI part corr ICA25 edge 2 link 1-3** | 0.33 | 15864 |
| **rfMRI full corr ICA100 edge 107 link 2-55** | 0.33 | 15864 |
| **OD Superior_cerebellar_peduncle_R** | 0.33 | 16541 |
| **rfMRI full corr ICA100 edge 142 link 3-38** | 0.33 | 15864 |
| **rfMRI full corr ICA100 edge 1371 link 40-46** | 0.33 | 15864 |
| **rfMRI part corr ICA100 edge 1186 link 31-32** | 0.33 | 15864 |
| **GM_thickness Morphologist FCLrdiag_right** | 0.33 | 11510 |
| **rfMRI amplitude ICA100 component 3** | 0.33 | 15863 |
| **rfMRI part corr ICA100 edge 516 link 11-32** | 0.33 | 15864 |
| **MO Inferior_cerebellar_peduncle_R** | 0.33 | 16541 |
| **rfMRI part corr ICA100 edge 1314 link 36-55** | 0.33 | 15864 |
| **rfMRI full corr ICA100 edge 1294 link 35-54** | 0.33 | 15864 |

| | | |
|---|---|---|
| **ICVF Corticospinal_tract_R** | 0.33 | 16540 |
| **rfMRI part corr ICA25 edge 60 link 4-7** | 0.33 | 15864 |
| **rfMRI part corr ICA100 edge 747 link 17-20** | 0.33 | 15864 |
| **rfMRI full corr ICA100 edge 98 link 2-46** | 0.33 | 15864 |
| **rfMRI full corr ICA100 edge 17 link 1-18** | 0.33 | 15864 |
| **rfMRI part corr ICA25 edge 91 link 6-7** | 0.33 | 15864 |
| **rfMRI full corr ICA25 edge 37 link 2-19** | 0.33 | 15864 |
| **ICVF Medial_lemniscus_R** | 0.33 | 16540 |
| **rfMRI part corr ICA100 edge 1437 link 45-52** | 0.33 | 15864 |
| **rfMRI part corr ICA100 edge 462 link 10-22** | 0.33 | 15864 |
| **rfMRI part corr ICA100 edge 874 link 20-39** | 0.34 | 15864 |
| **rfMRI part corr ICA100 edge 1262 link 34-42** | 0.34 | 15864 |
| **rfMRI part corr ICA100 edge 759 link 17-32** | 0.34 | 15864 |
| **rfMRI part corr ICA100 edge 1395 link 42-43** | 0.34 | 15864 |
| **rfMRI part corr ICA100 edge 961 link 23-27** | 0.34 | 15864 |
| **rfMRI part corr ICA100 edge 36 link 1-37** | 0.34 | 15864 |
| **rfMRI full corr ICA100 edge 1125 link 28-46** | 0.34 | 15864 |
| **rfMRI amplitude ICA100 component 42** | 0.34 | 15863 |
| **MO Superior_corona_radiata_L** | 0.34 | 16541 |
| **rfMRI full corr ICA100 edge 1009 link 24-44** | 0.34 | 15864 |
| **rfMRI part corr ICA100 edge 578 link 12-51** | 0.34 | 15864 |
| **rfMRI full corr ICA100 edge 455 link 10-15** | 0.34 | 15864 |
| **rfMRI part corr ICA25 edge 155 link 10-21** | 0.34 | 15864 |
| **rfMRI part corr ICA25 edge 195 link 15-21** | 0.34 | 15864 |
| **rfMRI part corr ICA100 edge 1392 link 41-53** | 0.34 | 15864 |
| **OD Middle_cerebellar_peduncle** | 0.34 | 16541 |
| **rfMRI full corr ICA100 edge 1224 link 32-47** | 0.34 | 15864 |
| **rfMRI part corr ICA100 edge 1145 link 29-40** | 0.34 | 15864 |
| **rfMRI full corr ICA100 edge 1301 link 36-42** | 0.34 | 15864 |
| **rfMRI full corr ICA100 edge 1144 link 29-39** | 0.34 | 15863 |
| **rfMRI part corr ICA100 edge 488 link 10-48** | 0.34 | 15864 |
| **hull_junction_length Morphologist SRinf_left** | 0.34 | 16697 |
| **rfMRI part corr ICA100 edge 728 link 16-39** | 0.34 | 15864 |
| **GM_thickness Morphologist FCLrscant_left** | 0.34 | 4322 |

| | | |
|---|---|---|
| **ICVF Inferior_cerebellar_peduncle_L** | 0.34 | 16540 |
| **rfMRI part corr ICA100 edge 461 link 10-21** | 0.34 | 15864 |
| **meandepth Morphologist SFpolairetr_right** | 0.34 | 18086 |
| **rfMRI full corr ICA100 edge 1234 link 33-35** | 0.34 | 15864 |
| **GM_thickness Morphologist SGSM_left** | 0.34 | 12705 |
| **rfMRI full corr ICA100 edge 1081 link 27-29** | 0.34 | 15864 |
| **rfMRI full corr ICA100 edge 897 link 21-28** | 0.34 | 15864 |
| **rfMRI part corr ICA100 edge 736 link 16-47** | 0.34 | 15864 |
| **rfMRI part corr ICA25 edge 48 link 3-12** | 0.34 | 15864 |
| **rfMRI full corr ICA100 edge 989 link 23-55** | 0.34 | 15864 |
| **ThickAvg Destrieux G_front_middle.rh** | 0.34 | 17127 |
| **GM_thickness Morphologist SFsup_right** | 0.34 | 18093 |
| **rfMRI part corr ICA100 edge 1225 link 32-48** | 0.34 | 15864 |
| **rfMRI full corr ICA100 edge 177 link 4-22** | 0.34 | 15864 |
| **OD Cingulum-hippocampus-R** | 0.34 | 16541 |
| **rfMRI full corr ICA100 edge 331 link 7-29** | 0.34 | 15864 |
| **rfMRI part corr ICA25 edge 23 link 2-5** | 0.34 | 15864 |
| **rfMRI part corr ICA100 edge 1295 link 35-55** | 0.34 | 15864 |
| **rfMRI part corr ICA100 edge 229 link 5-24** | 0.34 | 15864 |
| **rfMRI part corr ICA100 edge 1285 link 35-45** | 0.34 | 15864 |
| **rfMRI part corr ICA25 edge 122 link 8-11** | 0.34 | 15864 |
| **ThickAvg Desikan supramarginal.rh** | 0.34 | 17127 |
| **meandepth Morphologist SPasup_left** | 0.34 | 16533 |
| **rfMRI full corr ICA100 edge 468 link 10-28** | 0.34 | 15864 |
| **rfMRI part corr ICA100 edge 903 link 21-34** | 0.34 | 15864 |
| **GrayVol Destrieux S_calcarine.rh** | 0.34 | 17127 |
| **hull_junction_length Morphologist SPeCsup_left** | 0.34 | 17143 |
| **rfMRI part corr ICA100 edge 464 link 10-24** | 0.34 | 15864 |
| **rfMRI full corr ICA100 edge 1199 link 31-45** | 0.34 | 15864 |
| **opening Morphologist FCMpost_right** | 0.35 | 18100 |
| **ThickAvg Desikan caudalmiddlefrontal.lh** | 0.35 | 17127 |
| **rfMRI full corr ICA100 edge 477 link 10-37** | 0.35 | 15864 |
| **rfMRI full corr ICA100 edge 296 link 6-42** | 0.35 | 15864 |
| **rfMRI part corr ICA100 edge 427 link 9-32** | 0.35 | 15864 |

| | | |
|---|---|---|
| **rfMRI part corr ICA100 edge 1236 link 33-37** | 0.35 | 15864 |
| **OD Medial_lemniscus_L** | 0.35 | 16541 |
| **rfMRI full corr ICA100 edge 717 link 16-28** | 0.35 | 15863 |
| **SurfArea Destrieux G_Ins_lg_and_S_cent_ins.rh** | 0.35 | 17125 |
| **rfMRI full corr ICA100 edge 333 link 7-31** | 0.35 | 15864 |
| **rfMRI part corr ICA100 edge 295 link 6-41** | 0.35 | 15864 |
| **rfMRI part corr ICA100 edge 766 link 17-39** | 0.35 | 15864 |
| **ThickAvg Destrieux G_rectus.lh** | 0.35 | 17127 |
| **rfMRI part corr ICA25 edge 136 link 9-13** | 0.35 | 15864 |
| **ThickAvg Desikan postcentral.rh** | 0.35 | 17127 |
| **rfMRI full corr ICA100 edge 1229 link 32-52** | 0.35 | 15864 |
| **rfMRI full corr ICA100 edge 996 link 24-31** | 0.35 | 15864 |
| **rfMRI part corr ICA100 edge 213 link 5-8** | 0.35 | 15864 |
| **ICVF Posterior_thalamic_radiation-include_optic_r** | 0.35 | 16540 |
| **meandepth Morphologist FCLrasc_right** | 0.35 | 17490 |
| **rfMRI full corr ICA100 edge 1019 link 24-54** | 0.35 | 15864 |
| **rfMRI full corr ICA100 edge 1007 link 24-42** | 0.35 | 15864 |
| **rfMRI part corr ICA100 edge 782 link 17-55** | 0.35 | 15864 |
| **rfMRI full corr ICA100 edge 771 link 17-44** | 0.35 | 15864 |
| **rfMRI full corr ICA100 edge 762 link 17-35** | 0.35 | 15864 |
| **rfMRI full corr ICA100 edge 1435 link 45-50** | 0.35 | 15864 |
| **SurfArea Destrieux S_orbital_med-olfact.lh** | 0.35 | 17126 |
| **rfMRI part corr ICA100 edge 707 link 16-18** | 0.35 | 15864 |
| **rfMRI full corr ICA100 edge 1429 link 44-54** | 0.35 | 15864 |
| **rfMRI full corr ICA100 edge 508 link 11-24** | 0.35 | 15864 |
| **rfMRI part corr ICA100 edge 405 link 9-10** | 0.35 | 15864 |
| **rfMRI full corr ICA100 edge 230 link 5-25** | 0.35 | 15864 |
| **rfMRI part corr ICA100 edge 1128 link 28-49** | 0.35 | 15864 |
| **rfMRI part corr ICA25 edge 85 link 5-16** | 0.35 | 15864 |
| **surface Morphologist FCLp_right** | 0.35 | 18100 |
| **rfMRI full corr ICA100 edge 962 link 23-28** | 0.35 | 15864 |
| **rfMRI full corr ICA100 edge 376 link 8-27** | 0.35 | 15864 |
| **ThickAvg Destrieux G_pariet_inf-Supramar.rh** | 0.35 | 17127 |
| **rfMRI full corr ICA100 edge 885 link 20-50** | 0.35 | 15864 |

| | | |
|---|---|---|
| **rfMRI part corr ICA25 edge 161 link 11-17** | 0.35 | 15864 |
| **rfMRI full corr ICA100 edge 526 link 11-42** | 0.35 | 15864 |
| **rfMRI full corr ICA100 edge 837 link 19-37** | 0.35 | 15864 |
| **rfMRI full corr ICA100 edge 1255 link 34-35** | 0.35 | 15864 |
| **rfMRI full corr ICA100 edge 134 link 3-30** | 0.35 | 15864 |
| **rfMRI part corr ICA100 edge 162 link 4-7** | 0.35 | 15864 |
| **rfMRI full corr ICA100 edge 440 link 9-45** | 0.35 | 15864 |
| **rfMRI full corr ICA25 edge 67 link 4-14** | 0.35 | 15864 |
| **rfMRI full corr ICA100 edge 128 link 3-24** | 0.36 | 15864 |
| **rfMRI full corr ICA100 edge 1109 link 28-30** | 0.36 | 15864 |
| **rfMRI part corr ICA100 edge 618 link 13-49** | 0.36 | 15864 |
| **maxdepth Morphologist INSULA_right** | 0.36 | 18100 |
| **rfMRI full corr ICA100 edge 1277 link 35-37** | 0.36 | 15864 |
| **rfMRI full corr ICA100 edge 151 link 3-47** | 0.36 | 15863 |
| **MD Corticospinal_tract_R** | 0.36 | 16541 |
| **hull_junction_length Morphologist SPat_right** | 0.36 | 16760 |
| **SurfArea Destrieux G_and_S_frontomargin.lh** | 0.36 | 17127 |
| **rfMRI part corr ICA100 edge 1268 link 34-48** | 0.36 | 15864 |
| **OD Posterior_corona_radiata_R** | 0.36 | 16541 |
| **rfMRI part corr ICA25 edge 110 link 7-12** | 0.36 | 15864 |
| **rfMRI full corr ICA100 edge 59 link 2-7** | 0.36 | 15863 |
| **hull_junction_length Morphologist SCu_left** | 0.36 | 18058 |
| **rfMRI full corr ICA100 edge 8 link 1-9** | 0.36 | 15855 |
| **rfMRI part corr ICA100 edge 1194 link 31-40** | 0.36 | 15864 |
| **OD Retrolenticular_part_of_internal_capsule_R** | 0.36 | 16541 |
| **rfMRI part corr ICA100 edge 434 link 9-39** | 0.36 | 15864 |
| **rfMRI part corr ICA100 edge 815 link 18-51** | 0.36 | 15864 |
| **rfMRI full corr ICA100 edge 235 link 5-30** | 0.36 | 15864 |
| **OD Pontine_crossing_tract-a_part_of_MCP** | 0.36 | 16541 |
| **rfMRI part corr ICA100 edge 148 link 3-44** | 0.36 | 15864 |
| **rfMRI amplitude ICA25 component 8** | 0.36 | 15863 |
| **rfMRI full corr ICA100 edge 759 link 17-32** | 0.36 | 15864 |
| **rfMRI full corr ICA25 edge 129 link 8-18** | 0.36 | 15864 |
| **rfMRI amplitude ICA100 component 45** | 0.36 | 15863 |

| | | |
|---|---|---|
| **MO Sagittal_stratum-inf_longitidinal_fasci_and_inf** | 0.36 | 16541 |
| **rfMRI part corr ICA100 edge 993 link 24-28** | 0.36 | 15864 |
| **rfMRI part corr ICA25 edge 189 link 14-21** | 0.36 | 15864 |
| **rfMRI full corr ICA100 edge 1049 link 25-54** | 0.36 | 15864 |
| **rfMRI part corr ICA100 edge 467 link 10-27** | 0.36 | 15864 |
| **rfMRI full corr ICA100 edge 238 link 5-33** | 0.36 | 15864 |
| **rfMRI part corr ICA100 edge 553 link 12-26** | 0.36 | 15864 |
| **ThickAvg Destrieux S_temporal_transverse.lh** | 0.36 | 17126 |
| **rfMRI full corr ICA100 edge 1343 link 38-49** | 0.36 | 15864 |
| **rfMRI part corr ICA100 edge 622 link 13-53** | 0.36 | 15864 |
| **GrayVol Destrieux G_and_S_cingul-Mid-Post.rh** | 0.36 | 17127 |
| **ISOVF Superior_longitudinal_fasciculus_L** | 0.36 | 16527 |
| **rfMRI part corr ICA100 edge 454 link 10-14** | 0.36 | 15864 |
| **meandepth Morphologist SOTlatant_right** | 0.36 | 18047 |
| **rfMRI full corr ICA25 edge 161 link 11-17** | 0.36 | 15864 |
| **rfMRI full corr ICA100 edge 716 link 16-27** | 0.36 | 15860 |
| **rfMRI part corr ICA100 edge 1260 link 34-40** | 0.36 | 15864 |
| **rfMRI full corr ICA100 edge 395 link 8-46** | 0.37 | 15864 |
| **rfMRI part corr ICA100 edge 570 link 12-43** | 0.37 | 15864 |
| **ICVF Superior_corona_radiata_L** | 0.37 | 16540 |
| **GM_thickness Morphologist FCLa_left** | 0.37 | 18048 |
| **GrayVol Destrieux S_intrapariet_and_P_trans.rh** | 0.37 | 17127 |
| **rfMRI part corr ICA100 edge 160 link 4-5** | 0.37 | 15864 |
| **rfMRI part corr ICA100 edge 644 link 14-34** | 0.37 | 15864 |
| **FA Anterior_limb_of_internal_capsule_L** | 0.37 | 16541 |
| **rfMRI part corr ICA100 edge 881 link 20-46** | 0.37 | 15864 |
| **rfMRI part corr ICA100 edge 303 link 6-49** | 0.37 | 15864 |
| **ThickAvg Destrieux G_parietal_sup.rh** | 0.37 | 17127 |
| **rfMRI part corr ICA100 edge 1063 link 26-39** | 0.37 | 15864 |
| **rfMRI part corr ICA100 edge 317 link 7-15** | 0.37 | 15864 |
| **surface Morphologist INSULA_right** | 0.37 | 18100 |
| **ThickAvg Desikan superiorparietal.rh** | 0.37 | 17127 |
| **rfMRI full corr ICA100 edge 571 link 12-44** | 0.37 | 15864 |
| **rfMRI part corr ICA100 edge 621 link 13-52** | 0.37 | 15864 |

| | | |
|---|---|---|
| **rfMRI full corr ICA100 edge 854 link 19-54** | 0.37 | 15864 |
| **rfMRI part corr ICA100 edge 208 link 4-53** | 0.37 | 15864 |
| **rfMRI part corr ICA100 edge 362 link 8-13** | 0.37 | 15864 |
| **rfMRI full corr ICA100 edge 532 link 11-48** | 0.37 | 15864 |
| **rfMRI full corr ICA100 edge 1434 link 45-49** | 0.37 | 15864 |
| **rfMRI part corr ICA100 edge 698 link 15-48** | 0.37 | 15864 |
| **rfMRI full corr ICA25 edge 89 link 5-20** | 0.37 | 15864 |
| **rfMRI full corr ICA25 edge 189 link 14-21** | 0.37 | 15864 |
| **hull_junction_length Morphologist STpol_left** | 0.37 | 18076 |
| **rfMRI part corr ICA100 edge 1113 link 28-34** | 0.37 | 15864 |
| **MD Anterior_corona_radiata_R** | 0.37 | 16541 |
| **hull_junction_length Morphologist FColl_right** | 0.37 | 18100 |
| **ThickAvg Destrieux S_occipital_ant.rh** | 0.37 | 17127 |
| **rfMRI part corr ICA100 edge 1411 link 43-47** | 0.37 | 15864 |
| **Right-Cerebellum-Cortex** | 0.37 | 17127 |
| **rfMRI full corr ICA100 edge 254 link 5-49** | 0.37 | 15864 |
| **hull_junction_length Morphologist SForbitaire_left** | 0.37 | 17520 |
| **GrayVol Destrieux G_and_S_cingul-Mid-Ant.rh** | 0.37 | 17127 |
| **rfMRI part corr ICA100 edge 544 link 12-17** | 0.37 | 15864 |
| **MD Superior_corona_radiata_R** | 0.37 | 16541 |
| **ThickAvg Desikan cuneus.rh** | 0.37 | 17127 |
| **rfMRI part corr ICA100 edge 320 link 7-18** | 0.37 | 15864 |
| **rfMRI full corr ICA100 edge 828 link 19-28** | 0.37 | 15864 |
| **hull_junction_length Morphologist SOp_left** | 0.37 | 15273 |
| **rfMRI full corr ICA100 edge 1063 link 26-39** | 0.37 | 15864 |
| **maxdepth Morphologist SFmedian_right** | 0.37 | 18080 |
| **rfMRI part corr ICA100 edge 1351 link 39-41** | 0.37 | 15864 |
| **rfMRI full corr ICA25 edge 70 link 4-17** | 0.37 | 15864 |
| **rfMRI full corr ICA25 edge 9 link 1-10** | 0.38 | 15864 |
| **rfMRI part corr ICA100 edge 152 link 3-48** | 0.38 | 15864 |
| **rfMRI full corr ICA100 edge 1258 link 34-38** | 0.38 | 15864 |
| **rfMRI full corr ICA100 edge 65 link 2-13** | 0.38 | 15864 |
| **rfMRI full corr ICA100 edge 761 link 17-34** | 0.38 | 15864 |
| **rfMRI full corr ICA100 edge 1405 link 42-53** | 0.38 | 15864 |

| | | |
|---|---|---|
| **opening Morphologist SPat_left** | 0.38 | 13931 |
| **rfMRI full corr ICA100 edge 1375 link 40-50** | 0.38 | 15864 |
| **rfMRI full corr ICA100 edge 878 link 20-43** | 0.38 | 15864 |
| **surface Morphologist SCall_left** | 0.38 | 18064 |
| **FA Superior_corona_radiata_L** | 0.38 | 16541 |
| **rfMRI full corr ICA100 edge 1256 link 34-36** | 0.38 | 15864 |
| **surface Morphologist FColl_left** | 0.38 | 18101 |
| **surface Morphologist SCLPC_left** | 0.38 | 13421 |
| **meandepth Morphologist SCLPC_right** | 0.38 | 5250 |
| **rfMRI part corr ICA100 edge 21 link 1-22** | 0.38 | 15864 |
| **rfMRI full corr ICA100 edge 1143 link 29-38** | 0.38 | 15864 |
| **ThickAvg Destrieux Lat_Fis-ant-Horizont.rh** | 0.38 | 17125 |
| **rfMRI part corr ICA100 edge 369 link 8-20** | 0.38 | 15864 |
| **rfMRI full corr ICA25 edge 187 link 14-19** | 0.38 | 15864 |
| **rfMRI full corr ICA100 edge 282 link 6-28** | 0.38 | 15864 |
| **rfMRI full corr ICA100 edge 459 link 10-19** | 0.38 | 15864 |
| **rfMRI part corr ICA25 edge 208 link 19-20** | 0.38 | 15864 |
| **opening Morphologist SRinf_right** | 0.38 | 17661 |
| **rfMRI full corr ICA100 edge 969 link 23-35** | 0.38 | 15864 |
| **ISOVF Inferior_cerebellar_peduncle_L** | 0.38 | 16527 |
| **rfMRI full corr ICA25 edge 71 link 4-18** | 0.38 | 15864 |
| **rfMRI part corr ICA100 edge 757 link 17-30** | 0.38 | 15864 |
| **GM_thickness Morphologist FIP_right** | 0.38 | 18090 |
| **opening Morphologist SpC_right** | 0.38 | 16309 |
| **rfMRI full corr ICA100 edge 603 link 13-34** | 0.38 | 15864 |
| **rfMRI full corr ICA100 edge 1193 link 31-39** | 0.38 | 15864 |
| **rfMRI full corr ICA25 edge 147 link 10-13** | 0.38 | 15864 |
| **surface Morphologist FIP_right** | 0.38 | 18097 |
| **rfMRI full corr ICA100 edge 14 link 1-15** | 0.38 | 15864 |
| **FA Cingulum-cingulate_gyrus-R** | 0.38 | 16541 |
| **rfMRI full corr ICA100 edge 739 link 16-50** | 0.38 | 15864 |
| **hull_junction_length Morphologist SLipost_right** | 0.38 | 17978 |
| **rfMRI full corr ICA100 edge 647 link 14-37** | 0.38 | 15864 |
| **rfMRI full corr ICA100 edge 384 link 8-35** | 0.38 | 15862 |

| | | |
|---|---|---|
| **rfMRI full corr ICA100 edge 829 link 19-29** | 0.38 | 15861 |
| **SurfArea Desikan parsopercularis.rh** | 0.38 | 17127 |
| **rfMRI full corr ICA100 edge 30 link 1-31** | 0.38 | 15862 |
| **ThickAvg Destrieux G_and_S_subcentral.lh** | 0.38 | 17127 |
| **rfMRI part corr ICA100 edge 222 link 5-17** | 0.38 | 15864 |
| **rfMRI amplitude ICA25 component 7** | 0.38 | 15863 |
| **meandepth Morphologist FCLrscant_left** | 0.38 | 4279 |
| **rfMRI part corr ICA100 edge 936 link 22-34** | 0.38 | 15864 |
| **rfMRI part corr ICA100 edge 967 link 23-33** | 0.38 | 15864 |
| **rfMRI part corr ICA25 edge 28 link 2-10** | 0.38 | 15864 |
| **GrayVol Destrieux G_front_inf-Opercular.rh** | 0.38 | 17127 |
| **surface Morphologist SRinf_right** | 0.38 | 17661 |
| **rfMRI full corr ICA100 edge 1022 link 25-27** | 0.38 | 15864 |
| **rfMRI full corr ICA100 edge 1378 link 40-53** | 0.38 | 15864 |
| **rfMRI amplitude ICA100 component 7** | 0.38 | 15863 |
| **rfMRI full corr ICA100 edge 293 link 6-39** | 0.39 | 15864 |
| **rfMRI part corr ICA100 edge 116 link 3-12** | 0.39 | 15864 |
| **rfMRI part corr ICA25 edge 21 link 2-3** | 0.39 | 15864 |
| **GM_thickness Morphologist SFpolairetr_right** | 0.39 | 18079 |
| **rfMRI part corr ICA25 edge 158 link 11-14** | 0.39 | 15864 |
| **rfMRI full corr ICA100 edge 674 link 15-24** | 0.39 | 15864 |
| **surface Morphologist SPeCsup_left** | 0.39 | 17143 |
| **rfMRI amplitude ICA100 component 2** | 0.39 | 15863 |
| **rfMRI full corr ICA25 edge 65 link 4-12** | 0.39 | 15864 |
| **surface Morphologist FCMpost_right** | 0.39 | 18100 |
| **rfMRI full corr ICA100 edge 1042 link 25-47** | 0.39 | 15864 |
| **rfMRI full corr ICA100 edge 567 link 12-40** | 0.39 | 15864 |
| **rfMRI full corr ICA100 edge 688 link 15-38** | 0.39 | 15864 |
| **rfMRI part corr ICA100 edge 1118 link 28-39** | 0.39 | 15864 |
| **ThickAvg Destrieux S_oc_sup_and_transversal.lh** | 0.39 | 17126 |
| **rfMRI full corr ICA100 edge 926 link 22-24** | 0.39 | 15864 |
| **rfMRI part corr ICA100 edge 978 link 23-44** | 0.39 | 15864 |
| **rfMRI full corr ICA25 edge 191 link 15-17** | 0.39 | 15864 |
| **GrayVol Destrieux G_cingul-Post-dorsal.lh** | 0.39 | 17125 |

| | | |
|---|---|---|
| **rfMRI full corr ICA25 edge 1 link 1-2** | 0.39 | 15863 |
| **rfMRI part corr ICA100 edge 557 link 12-30** | 0.39 | 15864 |
| **hull_junction_length Morphologist OCCIPITAL_left** | 0.39 | 18097 |
| **ICVF Anterior_limb_of_internal_capsule_R** | 0.39 | 16540 |
| **meandepth Morphologist FCMpost_right** | 0.39 | 18100 |
| **rfMRI full corr ICA100 edge 849 link 19-49** | 0.39 | 15862 |
| **hull_junction_length Morphologist FPO_right** | 0.39 | 18097 |
| **rfMRI full corr ICA100 edge 833 link 19-33** | 0.39 | 15864 |
| **rfMRI full corr ICA100 edge 594 link 13-25** | 0.39 | 15864 |
| **rfMRI part corr ICA100 edge 884 link 20-49** | 0.39 | 15864 |
| **meandepth Morphologist SRh_left** | 0.39 | 17717 |
| **rfMRI part corr ICA25 edge 112 link 7-14** | 0.39 | 15864 |
| **rfMRI part corr ICA100 edge 833 link 19-33** | 0.39 | 15864 |
| **GM_thickness Morphologist SpC_right** | 0.39 | 16301 |
| **meandepth Morphologist SCsylvian_right** | 0.39 | 14952 |
| **rfMRI full corr ICA25 edge 136 link 9-13** | 0.39 | 15864 |
| **rfMRI full corr ICA100 edge 610 link 13-41** | 0.4 | 15864 |
| **rfMRI part corr ICA100 edge 23 link 1-24** | 0.4 | 15864 |
| **rfMRI part corr ICA100 edge 380 link 8-31** | 0.4 | 15864 |
| **GrayVol Destrieux S_circular_insula_sup.lh** | 0.4 | 17126 |
| **rfMRI part corr ICA100 edge 314 link 7-12** | 0.4 | 15864 |
| **rfMRI part corr ICA100 edge 558 link 12-31** | 0.4 | 15864 |
| **ThickAvg Destrieux G_and_S_paracentral.lh** | 0.4 | 17127 |
| **rfMRI part corr ICA100 edge 1473 link 50-53** | 0.4 | 15864 |
| **rfMRI full corr ICA100 edge 342 link 7-40** | 0.4 | 15864 |
| **maxdepth Morphologist SFint_left** | 0.4 | 18101 |
| **rfMRI full corr ICA100 edge 1298 link 36-39** | 0.4 | 15864 |
| **rfMRI part corr ICA100 edge 1404 link 42-52** | 0.4 | 15864 |
| **rfMRI full corr ICA100 edge 1442 link 46-48** | 0.4 | 15864 |
| **rfMRI part corr ICA100 edge 611 link 13-42** | 0.4 | 15864 |
| **rfMRI part corr ICA100 edge 1434 link 45-49** | 0.4 | 15864 |
| **rfMRI full corr ICA100 edge 651 link 14-41** | 0.4 | 15863 |
| **rfMRI full corr ICA100 edge 315 link 7-13** | 0.4 | 15864 |
| **rfMRI full corr ICA100 edge 191 link 4-36** | 0.4 | 15864 |

| | | |
|---|---|---|
| **rfMRI part corr ICA100 edge 279 link 6-25** | 0.4 | 15864 |
| **rfMRI full corr ICA100 edge 979 link 23-45** | 0.4 | 15864 |
| **ThickAvg Destrieux G_postcentral.lh** | 0.4 | 17127 |
| **rfMRI full corr ICA100 edge 273 link 6-19** | 0.4 | 15863 |
| **rfMRI full corr ICA100 edge 1235 link 33-36** | 0.4 | 15864 |
| **surface Morphologist FCLrdiag_left** | 0.4 | 8551 |
| **rfMRI part corr ICA100 edge 420 link 9-25** | 0.4 | 15864 |
| **SurfArea Destrieux G_front_inf-Orbital.lh** | 0.4 | 17125 |
| **rfMRI full corr ICA100 edge 974 link 23-40** | 0.4 | 15863 |
| **rfMRI full corr ICA100 edge 367 link 8-18** | 0.4 | 15863 |
| **rfMRI full corr ICA100 edge 870 link 20-35** | 0.4 | 15864 |
| **maxdepth Morphologist FPO_right** | 0.4 | 18097 |
| **rfMRI full corr ICA100 edge 1046 link 25-51** | 0.4 | 15864 |
| **rfMRI full corr ICA100 edge 46 link 1-47** | 0.4 | 15864 |
| **GrayVol Destrieux G_and_S_occipital_inf.lh** | 0.4 | 17127 |
| **GrayVol Desikan inferiortemporal.lh** | 0.4 | 17127 |
| **rfMRI part corr ICA100 edge 1232 link 32-55** | 0.4 | 15864 |
| **rfMRI full corr ICA100 edge 1349 link 38-55** | 0.4 | 15864 |
| **SurfArea Desikan lateraloccipital.rh** | 0.4 | 17127 |
| **rfMRI full corr ICA100 edge 1339 link 38-45** | 0.4 | 15864 |
| **rfMRI amplitude ICA100 component 40** | 0.4 | 15863 |
| **hull_junction_length Morphologist FCLrscpost_lef** | 0.4 | 13973 |
| **rfMRI full corr ICA100 edge 113 link 3-9** | 0.4 | 15863 |
| **rfMRI part corr ICA100 edge 264 link 6-10** | 0.4 | 15864 |
| **rfMRI full corr ICA100 edge 781 link 17-54** | 0.41 | 15864 |
| **rfMRI full corr ICA100 edge 611 link 13-42** | 0.41 | 15864 |
| **rfMRI full corr ICA100 edge 736 link 16-47** | 0.41 | 15863 |
| **rfMRI full corr ICA100 edge 1059 link 26-35** | 0.41 | 15862 |
| **rfMRI part corr ICA100 edge 344 link 7-42** | 0.41 | 15864 |
| **rfMRI full corr ICA100 edge 583 link 13-14** | 0.41 | 15864 |
| **rfMRI full corr ICA25 edge 14 link 1-15** | 0.41 | 15864 |
| **GM_thickness Morphologist STpol_right** | 0.41 | 18054 |
| **SurfArea Destrieux G_postcentral.rh** | 0.41 | 17127 |
| **rfMRI full corr ICA100 edge 972 link 23-38** | 0.41 | 15863 |

| | | |
|---|---|---|
| **rfMRI full corr ICA100 edge 894 link 21-25** | 0.41 | 15864 |
| **rfMRI part corr ICA100 edge 839 link 19-39** | 0.41 | 15864 |
| **rfMRI part corr ICA100 edge 1320 link 37-43** | 0.41 | 15864 |
| **maxdepth Morphologist FCLrretroCtr_right** | 0.41 | 16212 |
| **surface Morphologist INSULA_left** | 0.41 | 18101 |
| **GM_thickness Morphologist SRh_left** | 0.41 | 17708 |
| **maxdepth Morphologist SFmarginal_left** | 0.41 | 18044 |
| **rfMRI part corr ICA100 edge 115 link 3-11** | 0.41 | 15864 |
| **rfMRI part corr ICA100 edge 613 link 13-44** | 0.41 | 15864 |
| **rfMRI amplitude ICA100 component 35** | 0.41 | 15863 |
| **rfMRI full corr ICA100 edge 1271 link 34-51** | 0.41 | 15864 |
| **rfMRI part corr ICA100 edge 260 link 5-55** | 0.41 | 15864 |
| **SurfArea Destrieux S_front_inf.lh** | 0.41 | 17127 |
| **rfMRI full corr ICA100 edge 1401 link 42-49** | 0.41 | 15864 |
| **rfMRI full corr ICA100 edge 947 link 22-45** | 0.41 | 15864 |
| **Optic-Chiasm** | 0.41 | 17127 |
| **rfMRI part corr ICA100 edge 1229 link 32-52** | 0.41 | 15864 |
| **rfMRI part corr ICA100 edge 212 link 5-7** | 0.41 | 15864 |
| **rfMRI full corr ICA25 edge 196 link 16-17** | 0.41 | 15864 |
| **rfMRI part corr ICA100 edge 778 link 17-51** | 0.41 | 15864 |
| **rfMRI part corr ICA100 edge 483 link 10-43** | 0.41 | 15864 |
| **OD Genu_of_corpus_callosum** | 0.41 | 16541 |
| **rfMRI part corr ICA25 edge 147 link 10-13** | 0.41 | 15864 |
| **hull_junction_length Morphologist FCMpost_right** | 0.41 | 18100 |
| **opening Morphologist STpol_left** | 0.41 | 18076 |
| **rfMRI part corr ICA100 edge 1178 link 30-48** | 0.41 | 15864 |
| **rfMRI full corr ICA100 edge 770 link 17-43** | 0.41 | 15864 |
| **GM_thickness Morphologist SOTlatint_left** | 0.41 | 16585 |
| **rfMRI part corr ICA100 edge 182 link 4-27** | 0.41 | 15864 |
| **rfMRI full corr ICA25 edge 96 link 6-12** | 0.41 | 15864 |
| **ThickAvg Destrieux G_occipital_sup.lh** | 0.41 | 17127 |
| **rfMRI part corr ICA100 edge 1077 link 26-53** | 0.41 | 15864 |
| **rfMRI part corr ICA100 edge 1144 link 29-39** | 0.41 | 15864 |
| **rfMRI part corr ICA100 edge 737 link 16-48** | 0.42 | 15864 |

| | | |
|---|---|---|
| opening Morphologist SPoCsup_right | 0.42 | 17842 |
| Left-VentralDC | 0.42 | 17127 |
| GrayVol Destrieux G_occipital_middle.lh | 0.42 | 17127 |
| surface Morphologist SOlf_right | 0.42 | 18091 |
| rfMRI part corr ICA100 edge 1378 link 40-53 | 0.42 | 15864 |
| SurfArea Desikan precentral.lh | 0.42 | 17127 |
| rfMRI part corr ICA100 edge 1322 link 37-45 | 0.42 | 15864 |
| GrayVol Destrieux G_and_S_occipital_inf.rh | 0.42 | 17127 |
| rfMRI part corr ICA100 edge 172 link 4-17 | 0.42 | 15864 |
| rfMRI part corr ICA100 edge 278 link 6-24 | 0.42 | 15864 |
| rfMRI full corr ICA100 edge 392 link 8-43 | 0.42 | 15864 |
| rfMRI part corr ICA100 edge 733 link 16-44 | 0.42 | 15864 |
| MD Uncinate_fasciculus_L | 0.42 | 16541 |
| ThickAvg Destrieux G_oc-temp_lat-fusifor.lh | 0.42 | 17127 |
| ICVF Fornix-column_and_body_of_fornix | 0.42 | 16540 |
| rfMRI part corr ICA100 edge 262 link 6-8 | 0.42 | 15864 |
| opening Morphologist SPeCmarginal_right | 0.42 | 17010 |
| MD Posterior_thalamic_radiation-include_optic_ra | 0.42 | 16541 |
| rfMRI part corr ICA100 edge 67 link 2-15 | 0.42 | 15864 |
| GM_thickness Morphologist FIP_left | 0.42 | 18089 |
| rfMRI part corr ICA100 edge 1282 link 35-42 | 0.42 | 15864 |
| rfMRI full corr ICA100 edge 258 link 5-53 | 0.42 | 15864 |
| rfMRI part corr ICA100 edge 1199 link 31-45 | 0.42 | 15864 |
| SurfArea Desikan rostralanteriorcingulate.lh | 0.42 | 17127 |
| rfMRI full corr ICA100 edge 18 link 1-19 | 0.42 | 15863 |
| rfMRI full corr ICA100 edge 602 link 13-33 | 0.42 | 15864 |
| Right-Thalamus-Proper | 0.42 | 17127 |
| rfMRI part corr ICA100 edge 813 link 18-49 | 0.42 | 15864 |
| rfMRI full corr ICA100 edge 115 link 3-11 | 0.42 | 15864 |
| rfMRI part corr ICA100 edge 1062 link 26-38 | 0.42 | 15864 |
| rfMRI full corr ICA100 edge 832 link 19-32 | 0.42 | 15864 |
| SurfArea Destrieux G_oc-temp_lat-fusifor.rh | 0.42 | 17127 |
| rfMRI part corr ICA100 edge 13 link 1-14 | 0.42 | 15864 |
| rfMRI full corr ICA100 edge 812 link 18-48 | 0.42 | 15864 |

| | | |
|---|---|---|
| **rfMRI full corr ICA100 edge 1437 link 45-52** | 0.42 | 15864 |
| **rfMRI full corr ICA25 edge 176 link 13-15** | 0.42 | 15864 |
| **rfMRI full corr ICA100 edge 410 link 9-15** | 0.42 | 15864 |
| **rfMRI part corr ICA100 edge 1132 link 28-53** | 0.42 | 15864 |
| **rfMRI amplitude ICA100 component 47** | 0.42 | 15863 |
| **rfMRI full corr ICA100 edge 92 link 2-40** | 0.42 | 15864 |
| **hull_junction_length Morphologist SFmedian_righ** | 0.42 | 18080 |
| **GM_thickness Morphologist SPasup_left** | 0.42 | 16529 |
| **rfMRI part corr ICA100 edge 569 link 12-42** | 0.42 | 15864 |
| **hull_junction_length Morphologist SLipost_left** | 0.42 | 17919 |
| **rfMRI part corr ICA100 edge 1196 link 31-42** | 0.42 | 15864 |
| **GM_thickness Morphologist SPat_left** | 0.42 | 13916 |
| **MO Retrolenticular_part_of_internal_capsule_R** | 0.42 | 16541 |
| **rfMRI part corr ICA100 edge 1143 link 29-38** | 0.42 | 15864 |
| **rfMRI part corr ICA25 edge 137 link 9-14** | 0.42 | 15864 |
| **rfMRI full corr ICA25 edge 32 link 2-14** | 0.43 | 15863 |
| **rfMRI full corr ICA100 edge 154 link 3-50** | 0.43 | 15864 |
| **rfMRI part corr ICA100 edge 1183 link 30-53** | 0.43 | 15864 |
| **rfMRI full corr ICA100 edge 464 link 10-24** | 0.43 | 15864 |
| **rfMRI part corr ICA100 edge 102 link 2-50** | 0.43 | 15864 |
| **rfMRI part corr ICA100 edge 367 link 8-18** | 0.43 | 15864 |
| **rfMRI part corr ICA100 edge 1127 link 28-48** | 0.43 | 15864 |
| **rfMRI full corr ICA100 edge 132 link 3-28** | 0.43 | 15864 |
| **rfMRI full corr ICA100 edge 1270 link 34-50** | 0.43 | 15864 |
| **ThickAvg Desikan rostralanteriorcingulate.lh** | 0.43 | 17127 |
| **rfMRI amplitude ICA100 component 14** | 0.43 | 15863 |
| **rfMRI part corr ICA100 edge 847 link 19-47** | 0.43 | 15864 |
| **rfMRI full corr ICA25 edge 98 link 6-14** | 0.43 | 15864 |
| **rfMRI part corr ICA100 edge 1037 link 25-42** | 0.43 | 15864 |
| **rfMRI full corr ICA25 edge 153 link 10-19** | 0.43 | 15864 |
| **rfMRI full corr ICA100 edge 599 link 13-30** | 0.43 | 15862 |
| **rfMRI part corr ICA25 edge 141 link 9-18** | 0.43 | 15864 |
| **rfMRI full corr ICA100 edge 1152 link 29-47** | 0.43 | 15864 |
| **rfMRI part corr ICA100 edge 514 link 11-30** | 0.43 | 15864 |

| | | |
|---|---|---|
| **rfMRI full corr ICA100 edge 1213 link 32-36** | 0.43 | 15864 |
| **rfMRI full corr ICA100 edge 503 link 11-19** | 0.43 | 15862 |
| **rfMRI full corr ICA100 edge 1151 link 29-46** | 0.43 | 15864 |
| **rfMRI part corr ICA100 edge 927 link 22-25** | 0.43 | 15864 |
| **rfMRI full corr ICA100 edge 1018 link 24-53** | 0.43 | 15863 |
| **rfMRI full corr ICA100 edge 157 link 3-53** | 0.43 | 15864 |
| **meandepth Morphologist STipost_right** | 0.43 | 18093 |
| **rfMRI part corr ICA100 edge 221 link 5-16** | 0.43 | 15864 |
| **rfMRI full corr ICA100 edge 1040 link 25-45** | 0.43 | 15863 |
| **rfMRI part corr ICA100 edge 1455 link 47-53** | 0.43 | 15864 |
| **rfMRI part corr ICA100 edge 540 link 12-13** | 0.43 | 15864 |
| **rfMRI full corr ICA100 edge 809 link 18-45** | 0.43 | 15863 |
| **rfMRI part corr ICA100 edge 425 link 9-30** | 0.43 | 15864 |
| **rfMRI full corr ICA100 edge 1168 link 30-38** | 0.43 | 15864 |
| **rfMRI part corr ICA25 edge 204 link 17-21** | 0.43 | 15864 |
| **rfMRI part corr ICA100 edge 675 link 15-25** | 0.43 | 15864 |
| **rfMRI part corr ICA25 edge 191 link 15-17** | 0.43 | 15864 |
| **SurfArea Destrieux G_rectus.rh** | 0.43 | 17126 |
| **SurfArea Desikan postcentral.rh** | 0.43 | 17127 |
| **rfMRI full corr ICA100 edge 537 link 11-53** | 0.43 | 15863 |
| **rfMRI full corr ICA100 edge 356 link 7-54** | 0.43 | 15864 |
| **ThickAvg Destrieux G_occipital_middle.rh** | 0.43 | 17127 |
| **rfMRI full corr ICA100 edge 248 link 5-43** | 0.43 | 15864 |
| **rfMRI part corr ICA100 edge 417 link 9-22** | 0.43 | 15864 |
| **opening Morphologist OCCIPITAL_right** | 0.43 | 18096 |
| **rfMRI part corr ICA100 edge 105 link 2-53** | 0.44 | 15864 |
| **GM_thickness Morphologist FIPPoCinf_right** | 0.44 | 18065 |
| **rfMRI part corr ICA100 edge 481 link 10-41** | 0.44 | 15864 |
| **rfMRI part corr ICA100 edge 526 link 11-42** | 0.44 | 15864 |
| **opening Morphologist SsP_left** | 0.44 | 18098 |
| **GrayVol Destrieux Lat_Fis-ant-Vertical.lh** | 0.44 | 17122 |
| **rfMRI full corr ICA100 edge 1220 link 32-43** | 0.44 | 15864 |
| **SurfArea Destrieux Lat_Fis-ant-Horizont.rh** | 0.44 | 17125 |
| **rfMRI part corr ICA100 edge 1003 link 24-38** | 0.44 | 15864 |

| | | |
|---|---|---|
| **ThickAvg Destrieux G_front_sup.lh** | 0.44 | 17127 |
| **rfMRI full corr ICA100 edge 25 link 1-26** | 0.44 | 15864 |
| **rfMRI full corr ICA100 edge 671 link 15-21** | 0.44 | 15864 |
| **rfMRI part corr ICA100 edge 1341 link 38-47** | 0.44 | 15864 |
| **rfMRI part corr ICA100 edge 950 link 22-48** | 0.44 | 15864 |
| **rfMRI part corr ICA100 edge 37 link 1-38** | 0.44 | 15864 |
| **rfMRI part corr ICA100 edge 601 link 13-32** | 0.44 | 15864 |
| **rfMRI part corr ICA25 edge 82 link 5-13** | 0.44 | 15864 |
| **rfMRI full corr ICA100 edge 274 link 6-20** | 0.44 | 15858 |
| **rfMRI full corr ICA100 edge 1208 link 31-54** | 0.44 | 15864 |
| **rfMRI full corr ICA100 edge 80 link 2-28** | 0.44 | 15864 |
| **GM_thickness Morphologist FCLrretroCtr_right** | 0.44 | 16206 |
| **rfMRI part corr ICA100 edge 1450 link 47-48** | 0.44 | 15864 |
| **rfMRI full corr ICA100 edge 488 link 10-48** | 0.44 | 15864 |
| **MD Sagittal_stratum-inf_longitidinal_fasci_and_inf** | 0.44 | 16541 |
| **ICVF Fornix-cres-Stria_terminalis-not_resolved_w** | 0.44 | 16540 |
| **ThickAvg Destrieux S_parieto_occipital.rh** | 0.44 | 17127 |
| **rfMRI full corr ICA100 edge 1468 link 49-53** | 0.44 | 15864 |
| **hull_junction_length Morphologist FCLa_left** | 0.44 | 18059 |
| **ISOVF Superior_cerebellar_peduncle_R** | 0.44 | 16527 |
| **rfMRI part corr ICA100 edge 638 link 14-28** | 0.44 | 15864 |
| **rfMRI full corr ICA100 edge 1117 link 28-38** | 0.44 | 15863 |
| **rfMRI part corr ICA100 edge 423 link 9-28** | 0.44 | 15864 |
| **ThickAvg Desikan paracentral.lh** | 0.44 | 17127 |
| **SurfArea Destrieux S_suborbital.rh** | 0.44 | 17125 |
| **rfMRI part corr ICA100 edge 416 link 9-21** | 0.44 | 15864 |
| **rfMRI part corr ICA100 edge 149 link 3-45** | 0.44 | 15864 |
| **rfMRI full corr ICA100 edge 1095 link 27-43** | 0.44 | 15864 |
| **rfMRI part corr ICA100 edge 865 link 20-30** | 0.44 | 15864 |
| **rfMRI part corr ICA100 edge 156 link 3-52** | 0.44 | 15864 |
| **rfMRI full corr ICA100 edge 556 link 12-29** | 0.45 | 15863 |
| **rfMRI full corr ICA100 edge 543 link 12-16** | 0.45 | 15863 |
| **rfMRI full corr ICA100 edge 1483 link 53-54** | 0.45 | 15864 |
| **opening Morphologist FIPrint1_right** | 0.45 | 17034 |

| | | |
|---|---|---|
| **rfMRI full corr ICA100 edge 1484 link 53-55** | 0.45 | 15864 |
| **Right-choroid-plexus** | 0.45 | 17127 |
| **Right-Inf-Lat-Vent** | 0.45 | 17127 |
| **maxdepth Morphologist FCLa_left** | 0.45 | 18059 |
| **rfMRI full corr ICA100 edge 789 link 18-25** | 0.45 | 15863 |
| **surface Morphologist SLipost_right** | 0.45 | 17978 |
| **ICVF Superior_corona_radiata_R** | 0.45 | 16540 |
| **rfMRI full corr ICA100 edge 578 link 12-51** | 0.45 | 15864 |
| **rfMRI part corr ICA100 edge 223 link 5-18** | 0.45 | 15864 |
| **rfMRI full corr ICA25 edge 75 link 5-6** | 0.45 | 15864 |
| **rfMRI full corr ICA100 edge 1060 link 26-36** | 0.45 | 15863 |
| **ThickAvg Destrieux G_subcallosal.rh** | 0.45 | 17126 |
| **rfMRI full corr ICA25 edge 57 link 3-21** | 0.45 | 15864 |
| **rfMRI part corr ICA100 edge 1279 link 35-39** | 0.45 | 15864 |
| **SurfArea Destrieux S_pericallosal.rh** | 0.45 | 17127 |
| **rfMRI full corr ICA100 edge 1076 link 26-52** | 0.45 | 15863 |
| **rfMRI full corr ICA100 edge 370 link 8-21** | 0.45 | 15864 |
| **rfMRI full corr ICA100 edge 1346 link 38-52** | 0.45 | 15864 |
| **rfMRI full corr ICA100 edge 780 link 17-53** | 0.45 | 15864 |
| **rfMRI full corr ICA100 edge 695 link 15-45** | 0.45 | 15864 |
| **rfMRI full corr ICA100 edge 1147 link 29-42** | 0.45 | 15864 |
| **rfMRI part corr ICA100 edge 581 link 12-54** | 0.45 | 15864 |
| **rfMRI full corr ICA100 edge 1090 link 27-38** | 0.45 | 15864 |
| **GrayVol Destrieux G_precentral.lh** | 0.45 | 17127 |
| **GM_thickness Morphologist FCMpost_left** | 0.45 | 18092 |
| **rfMRI full corr ICA100 edge 675 link 15-25** | 0.45 | 15864 |
| **rfMRI part corr ICA100 edge 1283 link 35-43** | 0.45 | 15864 |
| **opening Morphologist SOp_left** | 0.45 | 15273 |
| **rfMRI part corr ICA100 edge 210 link 4-55** | 0.45 | 15864 |
| **rfMRI full corr ICA100 edge 1240 link 33-41** | 0.45 | 15864 |
| **rfMRI full corr ICA100 edge 148 link 3-44** | 0.45 | 15858 |
| **ThickAvg Destrieux G_and_S_cingul-Mid-Ant.rh** | 0.45 | 17127 |
| **SurfArea Desikan precuneus.lh** | 0.45 | 17127 |
| **rfMRI part corr ICA100 edge 1420 link 44-45** | 0.45 | 15864 |

| | | |
|---|---|---|
| **rfMRI full corr ICA100 edge 106 link 2-54** | 0.45 | 15860 |
| **rfMRI part corr ICA100 edge 930 link 22-28** | 0.45 | 15864 |
| **rfMRI full corr ICA100 edge 550 link 12-23** | 0.46 | 15863 |
| **rfMRI part corr ICA100 edge 734 link 16-45** | 0.46 | 15864 |
| **opening Morphologist SC_right** | 0.46 | 18100 |
| **rfMRI full corr ICA100 edge 1177 link 30-47** | 0.46 | 15864 |
| **rfMRI full corr ICA100 edge 916 link 21-47** | 0.46 | 15864 |
| **rfMRI part corr ICA100 edge 1060 link 26-36** | 0.46 | 15864 |
| **rfMRI full corr ICA100 edge 558 link 12-31** | 0.46 | 15864 |
| **rfMRI part corr ICA100 edge 1465 link 49-50** | 0.46 | 15864 |
| **rfMRI full corr ICA100 edge 351 link 7-49** | 0.46 | 15864 |
| **rfMRI full corr ICA100 edge 844 link 19-44** | 0.46 | 15864 |
| **surface Morphologist SCLPC_right** | 0.46 | 5279 |
| **rfMRI full corr ICA100 edge 1272 link 34-52** | 0.46 | 15863 |
| **rfMRI part corr ICA100 edge 327 link 7-25** | 0.46 | 15864 |
| **MO Inferior_cerebellar_peduncle_L** | 0.46 | 16541 |
| **rfMRI part corr ICA100 edge 1276 link 35-36** | 0.46 | 15864 |
| **rfMRI full corr ICA25 edge 162 link 11-18** | 0.46 | 15864 |
| **rfMRI part corr ICA100 edge 1105 link 27-53** | 0.46 | 15864 |
| **maxdepth Morphologist SLiant_right** | 0.46 | 17219 |
| **meandepth Morphologist SPeCinf_left** | 0.46 | 17263 |
| **meandepth Morphologist FCLp_right** | 0.46 | 18100 |
| **rfMRI full corr ICA100 edge 41 link 1-42** | 0.46 | 15864 |
| **meandepth Morphologist SFint_left** | 0.46 | 18101 |
| **rfMRI full corr ICA100 edge 873 link 20-38** | 0.46 | 15864 |
| **rfMRI part corr ICA100 edge 1049 link 25-54** | 0.46 | 15864 |
| **rfMRI full corr ICA100 edge 893 link 21-24** | 0.46 | 15864 |
| **rfMRI full corr ICA100 edge 250 link 5-45** | 0.46 | 15864 |
| **rfMRI full corr ICA25 edge 63 link 4-10** | 0.46 | 15864 |
| **rfMRI full corr ICA100 edge 3 link 1-4** | 0.46 | 15864 |
| **rfMRI part corr ICA100 edge 1430 link 44-55** | 0.46 | 15864 |
| **rfMRI full corr ICA100 edge 85 link 2-33** | 0.46 | 15864 |
| **rfMRI full corr ICA100 edge 1119 link 28-40** | 0.46 | 15864 |
| **rfMRI part corr ICA25 edge 52 link 3-16** | 0.46 | 15864 |

| | | |
|---|---|---|
| **opening Morphologist SFmarginal_left** | 0.46 | 18044 |
| **rfMRI full corr ICA100 edge 713 link 16-24** | 0.46 | 15864 |
| **ThickAvg Destrieux S_temporal_inf.rh** | 0.46 | 17127 |
| **OD Superior_cerebellar_peduncle_L** | 0.46 | 16541 |
| **rfMRI full corr ICA100 edge 1430 link 44-55** | 0.46 | 15864 |
| **rfMRI part corr ICA25 edge 101 link 6-17** | 0.46 | 15864 |
| **ThickAvg Desikan entorhinal.rh** | 0.46 | 17125 |
| **rfMRI part corr ICA100 edge 867 link 20-32** | 0.46 | 15864 |
| **ThickAvg Destrieux S_postcentral.rh** | 0.46 | 17127 |
| **hull_junction_length Morphologist SFint_right** | 0.46 | 18100 |
| **rfMRI full corr ICA100 edge 1105 link 27-53** | 0.46 | 15864 |
| **SurfArea Desikan postcentral.lh** | 0.46 | 17127 |
| **rfMRI part corr ICA100 edge 889 link 20-54** | 0.46 | 15864 |
| **rfMRI part corr ICA100 edge 1198 link 31-44** | 0.46 | 15864 |
| **rfMRI full corr ICA100 edge 569 link 12-42** | 0.46 | 15864 |
| **rfMRI full corr ICA100 edge 1073 link 26-49** | 0.47 | 15864 |
| **rfMRI full corr ICA100 edge 905 link 21-36** | 0.47 | 15864 |
| **opening Morphologist FCMpost_left** | 0.47 | 18101 |
| **rfMRI part corr ICA100 edge 1476 link 51-52** | 0.47 | 15864 |
| **GM_thickness Morphologist FIPrint1_right** | 0.47 | 17027 |
| **rfMRI part corr ICA25 edge 33 link 2-15** | 0.47 | 15864 |
| **rfMRI full corr ICA25 edge 154 link 10-20** | 0.47 | 15864 |
| **SurfArea Destrieux G_subcallosal.rh** | 0.47 | 17126 |
| **rfMRI part corr ICA100 edge 1461 link 48-52** | 0.47 | 15864 |
| **rfMRI full corr ICA100 edge 434 link 9-39** | 0.47 | 15864 |
| **ISOVF Anterior_corona_radiata_R** | 0.47 | 16527 |
| **rfMRI full corr ICA25 edge 159 link 11-15** | 0.47 | 15864 |
| **rfMRI part corr ICA100 edge 1331 link 37-54** | 0.47 | 15864 |
| **rfMRI full corr ICA25 edge 122 link 8-11** | 0.47 | 15864 |
| **rfMRI part corr ICA100 edge 780 link 17-53** | 0.47 | 15864 |
| **ICVF Cerebral_peduncle_R** | 0.47 | 16540 |
| **rfMRI part corr ICA100 edge 35 link 1-36** | 0.47 | 15864 |
| **GM_thickness Morphologist SFsup_left** | 0.47 | 18092 |
| **rfMRI full corr ICA100 edge 1120 link 28-41** | 0.47 | 15864 |

| | | |
|---|---|---|
| **rfMRI full corr ICA100 edge 348 link 7-46** | 0.47 | 15864 |
| **rfMRI part corr ICA100 edge 1460 link 48-51** | 0.47 | 15864 |
| **rfMRI full corr ICA100 edge 344 link 7-42** | 0.47 | 15864 |
| **rfMRI part corr ICA25 edge 26 link 2-8** | 0.47 | 15864 |
| **rfMRI part corr ICA100 edge 893 link 21-24** | 0.47 | 15864 |
| **rfMRI full corr ICA100 edge 815 link 18-51** | 0.47 | 15864 |
| **rfMRI part corr ICA100 edge 912 link 21-43** | 0.47 | 15864 |
| **opening Morphologist FCLrant_left** | 0.47 | 17708 |
| **rfMRI part corr ICA100 edge 1019 link 24-54** | 0.47 | 15864 |
| **SurfArea Destrieux G_temporal_inf.lh** | 0.47 | 17127 |
| **rfMRI full corr ICA100 edge 580 link 12-53** | 0.47 | 15864 |
| **rfMRI part corr ICA100 edge 1066 link 26-42** | 0.47 | 15864 |
| **rfMRI part corr ICA100 edge 1440 link 45-55** | 0.47 | 15864 |
| **rfMRI full corr ICA100 edge 582 link 12-55** | 0.47 | 15863 |
| **rfMRI part corr ICA25 edge 148 link 10-14** | 0.47 | 15864 |
| **GrayVol Desikan parsorbitalis.lh** | 0.47 | 17126 |
| **hull_junction_length Morphologist SLiant_right** | 0.47 | 17219 |
| **rfMRI full corr ICA100 edge 621 link 13-52** | 0.47 | 15862 |
| **ICVF Anterior_corona_radiata_R** | 0.47 | 16540 |
| **ThickAvg Desikan medialorbitofrontal.rh** | 0.47 | 17126 |
| **rfMRI full corr ICA100 edge 123 link 3-19** | 0.47 | 15864 |
| **ICVF Genu_of_corpus_callosum** | 0.48 | 16540 |
| **ThickAvg Desikan frontalpole.rh** | 0.48 | 17126 |
| **MD Medial_lemniscus_L** | 0.48 | 16541 |
| **ThickAvg Destrieux S_suborbital.lh** | 0.48 | 17126 |
| **rfMRI part corr ICA100 edge 207 link 4-52** | 0.48 | 15864 |
| **GrayVol Destrieux S_circular_insula_sup.rh** | 0.48 | 17126 |
| **rfMRI full corr ICA100 edge 413 link 9-18** | 0.48 | 15864 |
| **GrayVol Destrieux G_and_S_cingul-Mid-Ant.lh** | 0.48 | 17127 |
| **rfMRI full corr ICA100 edge 636 link 14-26** | 0.48 | 15864 |
| **rfMRI part corr ICA25 edge 209 link 19-21** | 0.48 | 15864 |
| **surface Morphologist SPasup_left** | 0.48 | 16537 |
| **rfMRI full corr ICA100 edge 509 link 11-25** | 0.48 | 15864 |
| **rfMRI part corr ICA100 edge 426 link 9-31** | 0.48 | 15864 |

| | | |
|---|---|---|
| **rfMRI part corr ICA100 edge 5 link 1-6** | 0.48 | 15864 |
| **rfMRI part corr ICA100 edge 878 link 20-43** | 0.48 | 15864 |
| **ThickAvg Destrieux G_front_inf-Triangul.rh** | 0.48 | 17127 |
| **rfMRI part corr ICA100 edge 1054 link 26-30** | 0.48 | 15864 |
| **rfMRI full corr ICA100 edge 545 link 12-18** | 0.48 | 15863 |
| **surface Morphologist SPasup_right** | 0.48 | 17400 |
| **rfMRI part corr ICA100 edge 1044 link 25-49** | 0.48 | 15864 |
| **rfMRI part corr ICA100 edge 389 link 8-40** | 0.48 | 15864 |
| **rfMRI part corr ICA100 edge 24 link 1-25** | 0.48 | 15864 |
| **rfMRI full corr ICA100 edge 564 link 12-37** | 0.48 | 15864 |
| **rfMRI amplitude ICA100 component 39** | 0.48 | 15863 |
| **GM_thickness Morphologist STipost_right** | 0.48 | 18085 |
| **SurfArea Destrieux G_postcentral.lh** | 0.48 | 17127 |
| **rfMRI part corr ICA100 edge 1168 link 30-38** | 0.48 | 15864 |
| **rfMRI part corr ICA100 edge 1267 link 34-47** | 0.48 | 15864 |
| **SurfArea Destrieux G_occipital_sup.lh** | 0.48 | 17127 |
| **rfMRI full corr ICA100 edge 548 link 12-21** | 0.48 | 15864 |
| **rfMRI amplitude ICA25 component 11** | 0.48 | 15863 |
| **GM_thickness Morphologist SRh_right** | 0.48 | 17611 |
| **rfMRI full corr ICA100 edge 898 link 21-29** | 0.48 | 15864 |
| **GM_thickness Morphologist SPoCsup_left** | 0.48 | 17966 |
| **opening Morphologist SFinf_right** | 0.48 | 18080 |
| **rfMRI full corr ICA100 edge 751 link 17-24** | 0.48 | 15864 |
| **rfMRI full corr ICA100 edge 1054 link 26-30** | 0.48 | 15864 |
| **rfMRI part corr ICA25 edge 95 link 6-11** | 0.48 | 15864 |
| **rfMRI full corr ICA100 edge 998 link 24-33** | 0.48 | 15864 |
| **maxdepth Morphologist SCLPC_left** | 0.48 | 13421 |
| **meandepth Morphologist FCMant_left** | 0.48 | 17879 |
| **rfMRI full corr ICA100 edge 450 link 9-55** | 0.48 | 15864 |
| **rfMRI part corr ICA100 edge 85 link 2-33** | 0.48 | 15864 |
| **rfMRI full corr ICA100 edge 597 link 13-28** | 0.48 | 15862 |
| **rfMRI part corr ICA100 edge 142 link 3-38** | 0.48 | 15864 |
| **rfMRI full corr ICA100 edge 263 link 6-9** | 0.48 | 15864 |
| **ISOVF Medial_lemniscus_L** | 0.48 | 16527 |

| | | |
|---|---|---|
| **rfMRI full corr ICA25 edge 173 link 12-20** | 0.48 | 15864 |
| **rfMRI part corr ICA100 edge 1359 link 39-49** | 0.48 | 15864 |
| **ThickAvg Destrieux G_and_S_occipital_inf.rh** | 0.48 | 17127 |
| **rfMRI full corr ICA100 edge 992 link 24-27** | 0.48 | 15858 |
| **rfMRI part corr ICA25 edge 119 link 7-21** | 0.48 | 15864 |
| **rfMRI part corr ICA100 edge 368 link 8-19** | 0.48 | 15864 |
| **rfMRI full corr ICA100 edge 295 link 6-41** | 0.48 | 15864 |
| **rfMRI part corr ICA100 edge 12 link 1-13** | 0.48 | 15864 |
| **GrayVol Destrieux G_and_S_transv_frontopol.lh** | 0.48 | 17127 |
| **rfMRI full corr ICA25 edge 199 link 16-20** | 0.48 | 15864 |
| **surface Morphologist SOTlatint_right** | 0.48 | 15621 |
| **ICVF Superior_cerebellar_peduncle_L** | 0.48 | 16540 |
| **GrayVol Destrieux S_oc-temp_med_and_Lingual.rl** | 0.49 | 17127 |
| **maxdepth Morphologist FCLp_right** | 0.49 | 18100 |
| **SurfArea Desikan pericalcarine.lh** | 0.49 | 17127 |
| **rfMRI part corr ICA100 edge 1293 link 35-53** | 0.49 | 15864 |
| **rfMRI full corr ICA100 edge 301 link 6-47** | 0.49 | 15863 |
| **opening Morphologist SOTlatpost_left** | 0.49 | 18062 |
| **rfMRI full corr ICA100 edge 131 link 3-27** | 0.49 | 15864 |
| **rfMRI full corr ICA100 edge 284 link 6-30** | 0.49 | 15864 |
| **hull_junction_length Morphologist SOTlatpost_rigl** | 0.49 | 18075 |
| **rfMRI part corr ICA100 edge 789 link 18-25** | 0.49 | 15864 |
| **rfMRI full corr ICA100 edge 484 link 10-44** | 0.49 | 15864 |
| **maxdepth Morphologist FCLrant_left** | 0.49 | 17708 |
| **rfMRI full corr ICA25 edge 78 link 5-9** | 0.49 | 15864 |
| **hull_junction_length Morphologist SPeCsup_right** | 0.49 | 17272 |
| **rfMRI full corr ICA100 edge 1078 link 26-54** | 0.49 | 15864 |
| **rfMRI full corr ICA100 edge 615 link 13-46** | 0.49 | 15864 |
| **rfMRI full corr ICA100 edge 891 link 21-22** | 0.49 | 15864 |
| **rfMRI full corr ICA100 edge 1396 link 42-44** | 0.49 | 15864 |
| **rfMRI full corr ICA100 edge 347 link 7-45** | 0.49 | 15864 |
| **rfMRI full corr ICA25 edge 158 link 11-14** | 0.49 | 15864 |
| **rfMRI full corr ICA100 edge 1410 link 43-46** | 0.49 | 15864 |
| **rfMRI part corr ICA100 edge 515 link 11-31** | 0.49 | 15864 |

| | | |
|---|---|---|
| **rfMRI part corr ICA100 edge 800 link 18-36** | 0.49 | 15864 |
| **rfMRI part corr ICA100 edge 293 link 6-39** | 0.49 | 15864 |
| **SurfArea Desikan lateralorbitofrontal.lh** | 0.49 | 17127 |
| **rfMRI full corr ICA100 edge 84 link 2-32** | 0.49 | 15864 |
| **rfMRI part corr ICA100 edge 768 link 17-41** | 0.49 | 15864 |
| **rfMRI full corr ICA100 edge 629 link 14-19** | 0.49 | 15863 |
| **meandepth Morphologist SPoCsup_right** | 0.49 | 17839 |
| **rfMRI part corr ICA100 edge 1441 link 46-47** | 0.49 | 15864 |
| **meandepth Morphologist SCLPC_left** | 0.49 | 13404 |
| **rfMRI part corr ICA100 edge 177 link 4-22** | 0.49 | 15864 |
| **rfMRI full corr ICA25 edge 207 link 18-21** | 0.49 | 15864 |
| **MD Genu_of_corpus_callosum** | 0.49 | 16541 |
| **GM_thickness Morphologist SFinfant_left** | 0.49 | 17642 |
| **opening Morphologist SPeCmedian_right** | 0.49 | 17074 |
| **SurfArea Desikan fusiform.rh** | 0.49 | 17127 |
| **GrayVol Destrieux G_front_inf-Orbital.lh** | 0.49 | 17125 |
| **rfMRI part corr ICA100 edge 141 link 3-37** | 0.49 | 15864 |
| **maxdepth Morphologist SsP_right** | 0.49 | 18094 |
| **SurfArea Destrieux G_front_inf-Orbital.rh** | 0.49 | 17127 |
| **rfMRI full corr ICA100 edge 620 link 13-51** | 0.49 | 15862 |
| **SurfArea Desikan superiortemporal.rh** | 0.49 | 17127 |
| **surface Morphologist FCLrant_right** | 0.49 | 16205 |
| **rfMRI full corr ICA100 edge 1382 link 41-43** | 0.49 | 15864 |
| **rfMRI part corr ICA100 edge 26 link 1-27** | 0.5 | 15864 |
| **ICVF Middle_cerebellar_peduncle** | 0.5 | 16540 |
| **meandepth Morphologist SsP_right** | 0.5 | 18094 |
| **rfMRI full corr ICA100 edge 930 link 22-28** | 0.5 | 15863 |
| **rfMRI part corr ICA100 edge 842 link 19-42** | 0.5 | 15864 |
| **GrayVol Desikan inferiorparietal.rh** | 0.5 | 17127 |
| **MO Pontine_crossing_tract-a_part_of_MCP** | 0.5 | 16541 |
| **rfMRI full corr ICA100 edge 977 link 23-43** | 0.5 | 15864 |
| **rfMRI part corr ICA100 edge 357 link 7-55** | 0.5 | 15864 |
| **meandepth Morphologist SFsup_right** | 0.5 | 18100 |
| **rfMRI part corr ICA100 edge 225 link 5-20** | 0.5 | 15864 |

| | | |
|---|---|---|
| **rfMRI full corr ICA25 edge 145 link 10-11** | 0.5 | 15864 |
| **rfMRI full corr ICA25 edge 43 link 3-7** | 0.5 | 15864 |
| **rfMRI part corr ICA25 edge 51 link 3-15** | 0.5 | 15864 |
| **rfMRI full corr ICA100 edge 280 link 6-26** | 0.5 | 15864 |
| **rfMRI part corr ICA25 edge 173 link 12-20** | 0.5 | 15864 |
| **GM_thickness Morphologist SPaint_right** | 0.5 | 18000 |
| **ThickAvg Desikan caudalmiddlefrontal.rh** | 0.5 | 17127 |
| **SurfArea Destrieux G_and_S_subcentral.lh** | 0.5 | 17127 |
| **5th-Ventricle** | 0.5 | 17127 |
| **rfMRI part corr ICA100 edge 1337 link 38-43** | 0.5 | 15864 |
| **rfMRI full corr ICA100 edge 1166 link 30-36** | 0.5 | 15864 |
| **SurfArea Destrieux G_and_S_cingul-Mid-Ant.rh** | 0.5 | 17127 |
| **rfMRI part corr ICA100 edge 941 link 22-39** | 0.5 | 15864 |
| **ISOVF Posterior_thalamic_radiation-include_optic** | 0.5 | 16527 |
| **rfMRI full corr ICA100 edge 265 link 6-11** | 0.5 | 15863 |
| **rfMRI full corr ICA100 edge 76 link 2-24** | 0.5 | 15863 |
| **meandepth Morphologist SsP_left** | 0.5 | 18098 |
| **rfMRI part corr ICA100 edge 838 link 19-38** | 0.5 | 15864 |
| **rfMRI part corr ICA100 edge 1097 link 27-45** | 0.5 | 15864 |
| **SurfArea Destrieux G_and_S_paracentral.rh** | 0.5 | 17127 |
| **ISOVF Superior_fronto-occipital_fasciculus-part_c** | 0.5 | 16527 |
| **rfMRI full corr ICA100 edge 1107 link 27-55** | 0.5 | 15864 |
| **rfMRI full corr ICA100 edge 1052 link 26-28** | 0.5 | 15864 |
| **rfMRI part corr ICA100 edge 1370 link 40-45** | 0.5 | 15864 |
| **rfMRI full corr ICA100 edge 1182 link 30-52** | 0.51 | 15864 |
| **rfMRI part corr ICA100 edge 1292 link 35-52** | 0.51 | 15864 |
| **rfMRI full corr ICA100 edge 1014 link 24-49** | 0.51 | 15864 |
| **rfMRI full corr ICA100 edge 938 link 22-36** | 0.51 | 15864 |
| **rfMRI full corr ICA100 edge 903 link 21-34** | 0.51 | 15864 |
| **GrayVol Destrieux S_temporal_inf.lh** | 0.51 | 17127 |
| **rfMRI full corr ICA100 edge 358 link 8-9** | 0.51 | 15863 |
| **rfMRI full corr ICA100 edge 1201 link 31-47** | 0.51 | 15864 |
| **rfMRI part corr ICA100 edge 888 link 20-53** | 0.51 | 15864 |
| **rfMRI part corr ICA100 edge 151 link 3-47** | 0.51 | 15864 |

| | | |
|---|---|---|
| **Left-Cerebellum-Cortex** | 0.51 | 17127 |
| **rfMRI part corr ICA100 edge 1021 link 25-26** | 0.51 | 15864 |
| **rfMRI part corr ICA25 edge 67 link 4-14** | 0.51 | 15864 |
| **rfMRI full corr ICA100 edge 678 link 15-28** | 0.51 | 15864 |
| **GM_thickness Morphologist FColl_left** | 0.51 | 18091 |
| **rfMRI part corr ICA100 edge 711 link 16-22** | 0.51 | 15864 |
| **rfMRI full corr ICA100 edge 129 link 3-25** | 0.51 | 15864 |
| **rfMRI full corr ICA100 edge 686 link 15-36** | 0.51 | 15864 |
| **rfMRI full corr ICA100 edge 1285 link 35-45** | 0.51 | 15864 |
| **rfMRI full corr ICA100 edge 593 link 13-24** | 0.51 | 15864 |
| **rfMRI full corr ICA100 edge 612 link 13-43** | 0.51 | 15864 |
| **rfMRI part corr ICA100 edge 851 link 19-51** | 0.51 | 15864 |
| **rfMRI full corr ICA100 edge 1239 link 33-40** | 0.51 | 15864 |
| **opening Morphologist SOr_left** | 0.51 | 18100 |
| **rfMRI part corr ICA100 edge 1040 link 25-45** | 0.51 | 15864 |
| **hull_junction_length Morphologist SPasup_right** | 0.51 | 17400 |
| **rfMRI full corr ICA25 edge 35 link 2-17** | 0.51 | 15864 |
| **rfMRI full corr ICA100 edge 259 link 5-54** | 0.51 | 15864 |
| **SurfArea Destrieux S_front_inf.rh** | 0.51 | 17127 |
| **rfMRI part corr ICA100 edge 619 link 13-50** | 0.51 | 15864 |
| **maxdepth Morphologist SOlf_right** | 0.51 | 18091 |
| **rfMRI full corr ICA100 edge 182 link 4-27** | 0.51 | 15864 |
| **rfMRI part corr ICA100 edge 337 link 7-35** | 0.51 | 15864 |
| **rfMRI full corr ICA100 edge 649 link 14-39** | 0.51 | 15863 |
| **rfMRI full corr ICA100 edge 504 link 11-20** | 0.51 | 15864 |
| **rfMRI part corr ICA100 edge 493 link 10-53** | 0.51 | 15864 |
| **rfMRI full corr ICA100 edge 575 link 12-48** | 0.51 | 15864 |
| **rfMRI part corr ICA100 edge 29 link 1-30** | 0.51 | 15864 |
| **MD Superior_longitudinal_fasciculus_L** | 0.51 | 16541 |
| **hull_junction_length Morphologist STs_right** | 0.51 | 18100 |
| **rfMRI full corr ICA25 edge 143 link 9-20** | 0.52 | 15864 |
| **surface Morphologist SFmarginal_right** | 0.52 | 18006 |
| **FA Corticospinal_tract_L** | 0.52 | 16541 |
| **rfMRI full corr ICA100 edge 1253 link 33-54** | 0.52 | 15864 |

| | | |
|---|---|---|
| **rfMRI full corr ICA25 edge 69 link 4-16** | 0.52 | 15864 |
| **rfMRI full corr ICA100 edge 1066 link 26-42** | 0.52 | 15863 |
| **rfMRI full corr ICA100 edge 412 link 9-17** | 0.52 | 15864 |
| **rfMRI full corr ICA100 edge 999 link 24-34** | 0.52 | 15864 |
| **rfMRI part corr ICA100 edge 779 link 17-52** | 0.52 | 15864 |
| **rfMRI full corr ICA100 edge 268 link 6-14** | 0.52 | 15863 |
| **rfMRI part corr ICA100 edge 587 link 13-18** | 0.52 | 15864 |
| **GrayVol Destrieux G_postcentral.rh** | 0.52 | 17127 |
| **rfMRI full corr ICA100 edge 834 link 19-34** | 0.52 | 15863 |
| **rfMRI part corr ICA100 edge 6 link 1-7** | 0.52 | 15864 |
| **rfMRI full corr ICA25 edge 118 link 7-20** | 0.52 | 15864 |
| **GrayVol Destrieux S_oc_middle_and_Lunatus.rh** | 0.52 | 17127 |
| **rfMRI full corr ICA100 edge 409 link 9-14** | 0.52 | 15864 |
| **rfMRI part corr ICA100 edge 1182 link 30-52** | 0.52 | 15864 |
| **rfMRI full corr ICA100 edge 1266 link 34-46** | 0.52 | 15864 |
| **rfMRI full corr ICA100 edge 453 link 10-13** | 0.52 | 15864 |
| **rfMRI full corr ICA100 edge 964 link 23-30** | 0.52 | 15863 |
| **rfMRI part corr ICA100 edge 101 link 2-49** | 0.52 | 15864 |
| **rfMRI full corr ICA25 edge 133 link 9-10** | 0.52 | 15864 |
| **rfMRI part corr ICA100 edge 1462 link 48-53** | 0.52 | 15864 |
| **MD Superior_cerebellar_peduncle_R** | 0.52 | 16541 |
| **rfMRI amplitude ICA100 component 25** | 0.52 | 15863 |
| **rfMRI full corr ICA100 edge 1034 link 25-39** | 0.52 | 15864 |
| **rfMRI full corr ICA100 edge 209 link 4-54** | 0.52 | 15864 |
| **rfMRI part corr ICA100 edge 1188 link 31-34** | 0.52 | 15864 |
| **ThickAvg Destrieux G_oc-temp_med-Lingual.rh** | 0.52 | 17127 |
| **ThickAvg Desikan lingual.rh** | 0.52 | 17127 |
| **rfMRI part corr ICA100 edge 316 link 7-14** | 0.52 | 15864 |
| **rfMRI full corr ICA100 edge 1087 link 27-35** | 0.52 | 15864 |
| **rfMRI full corr ICA100 edge 1082 link 27-30** | 0.52 | 15864 |
| **rfMRI part corr ICA25 edge 27 link 2-9** | 0.52 | 15864 |
| **rfMRI full corr ICA100 edge 581 link 12-54** | 0.52 | 15864 |
| **rfMRI part corr ICA100 edge 419 link 9-24** | 0.52 | 15864 |
| **rfMRI part corr ICA100 edge 1152 link 29-47** | 0.52 | 15864 |

| | | |
|---|---|---|
| **rfMRI full corr ICA100 edge 1408 link 43-44** | 0.52 | 15864 |
| **ThickAvg Destrieux S_subparietal.rh** | 0.52 | 17126 |
| **rfMRI part corr ICA100 edge 754 link 17-27** | 0.52 | 15864 |
| **SurfArea Desikan paracentral.rh** | 0.52 | 17127 |
| **Right-vessel** | 0.52 | 17127 |
| **rfMRI full corr ICA100 edge 826 link 19-26** | 0.52 | 15864 |
| **rfMRI part corr ICA100 edge 92 link 2-40** | 0.53 | 15864 |
| **rfMRI part corr ICA100 edge 397 link 8-48** | 0.53 | 15864 |
| **rfMRI part corr ICA100 edge 447 link 9-52** | 0.53 | 15864 |
| **rfMRI part corr ICA100 edge 695 link 15-45** | 0.53 | 15864 |
| **3rd-Ventricle** | 0.53 | 17127 |
| **rfMRI part corr ICA25 edge 68 link 4-15** | 0.53 | 15864 |
| **rfMRI full corr ICA100 edge 919 link 21-50** | 0.53 | 15864 |
| **rfMRI part corr ICA100 edge 902 link 21-33** | 0.53 | 15864 |
| **rfMRI full corr ICA100 edge 1320 link 37-43** | 0.53 | 15864 |
| **rfMRI full corr ICA100 edge 925 link 22-23** | 0.53 | 15864 |
| **Right-Pallidum** | 0.53 | 17127 |
| **rfMRI part corr ICA100 edge 184 link 4-29** | 0.53 | 15864 |
| **rfMRI part corr ICA100 edge 508 link 11-24** | 0.53 | 15864 |
| **rfMRI full corr ICA100 edge 11 link 1-12** | 0.53 | 15864 |
| **rfMRI part corr ICA25 edge 104 link 6-20** | 0.53 | 15864 |
| **rfMRI part corr ICA100 edge 1124 link 28-45** | 0.53 | 15864 |
| **rfMRI full corr ICA100 edge 186 link 4-31** | 0.53 | 15864 |
| **rfMRI part corr ICA100 edge 883 link 20-48** | 0.53 | 15864 |
| **rfMRI full corr ICA100 edge 869 link 20-34** | 0.53 | 15863 |
| **rfMRI full corr ICA100 edge 604 link 13-35** | 0.53 | 15864 |
| **rfMRI part corr ICA100 edge 72 link 2-20** | 0.53 | 15864 |
| **rfMRI part corr ICA100 edge 1446 link 46-52** | 0.53 | 15864 |
| **rfMRI full corr ICA25 edge 55 link 3-19** | 0.53 | 15864 |
| **GrayVol Destrieux G_temporal_inf.lh** | 0.53 | 17127 |
| **rfMRI part corr ICA100 edge 468 link 10-28** | 0.53 | 15864 |
| **rfMRI part corr ICA100 edge 1226 link 32-49** | 0.53 | 15864 |
| **SurfArea Destrieux S_pericallosal.lh** | 0.53 | 17127 |
| **rfMRI part corr ICA100 edge 1449 link 46-55** | 0.53 | 15864 |

| | | |
|---|---|---|
| **rfMRI part corr ICA100 edge 979 link 23-45** | 0.53 | 15864 |
| **rfMRI full corr ICA100 edge 1195 link 31-41** | 0.53 | 15862 |
| **rfMRI part corr ICA100 edge 377 link 8-28** | 0.53 | 15864 |
| **rfMRI full corr ICA100 edge 272 link 6-18** | 0.53 | 15864 |
| **rfMRI full corr ICA100 edge 1097 link 27-45** | 0.53 | 15864 |
| **rfMRI full corr ICA100 edge 287 link 6-33** | 0.53 | 15862 |
| **rfMRI full corr ICA100 edge 447 link 9-52** | 0.53 | 15864 |
| **rfMRI full corr ICA100 edge 811 link 18-47** | 0.53 | 15863 |
| **GM_thickness Morphologist FCLrant_left** | 0.53 | 17692 |
| **rfMRI full corr ICA100 edge 1222 link 32-45** | 0.53 | 15864 |
| **maxdepth Morphologist SOTlatpost_right** | 0.53 | 18075 |
| **rfMRI full corr ICA100 edge 700 link 15-50** | 0.53 | 15864 |
| **rfMRI part corr ICA100 edge 44 link 1-45** | 0.53 | 15864 |
| **opening Morphologist SPeCinter_left** | 0.53 | 18028 |
| **rfMRI full corr ICA25 edge 166 link 12-13** | 0.53 | 15864 |
| **MO Uncinate_fasciculus_R** | 0.53 | 16541 |
| **SurfArea Destrieux G_and_S_frontomargin.rh** | 0.53 | 17126 |
| **rfMRI part corr ICA100 edge 226 link 5-21** | 0.54 | 15864 |
| **rfMRI full corr ICA100 edge 697 link 15-47** | 0.54 | 15864 |
| **rfMRI full corr ICA100 edge 1027 link 25-32** | 0.54 | 15864 |
| **GM_thickness Morphologist FCLrasc_left** | 0.54 | 17772 |
| **rfMRI part corr ICA100 edge 103 link 2-51** | 0.54 | 15864 |
| **rfMRI full corr ICA100 edge 1290 link 35-50** | 0.54 | 15864 |
| **meandepth Morphologist SFmarginal_right** | 0.54 | 18006 |
| **rfMRI full corr ICA100 edge 1080 link 27-28** | 0.54 | 15864 |
| **rfMRI part corr ICA100 edge 575 link 12-48** | 0.54 | 15864 |
| **ISOVF Cingulum-cingulate_gyrus-L** | 0.54 | 16527 |
| **rfMRI part corr ICA100 edge 746 link 17-19** | 0.54 | 15864 |
| **rfMRI full corr ICA100 edge 166 link 4-11** | 0.54 | 15862 |
| **ThickAvg Destrieux Lat_Fis-post.rh** | 0.54 | 17127 |
| **rfMRI part corr ICA100 edge 911 link 21-42** | 0.54 | 15864 |
| **rfMRI full corr ICA100 edge 939 link 22-37** | 0.54 | 15858 |
| **rfMRI part corr ICA100 edge 1377 link 40-52** | 0.54 | 15864 |
| **rfMRI full corr ICA100 edge 1386 link 41-47** | 0.54 | 15864 |

| | | |
|---|---|---|
| **rfMRI part corr ICA100 edge 1206 link 31-52** | 0.54 | 15864 |
| **ThickAvg Destrieux G_front_middle.lh** | 0.54 | 17127 |
| **SurfArea Destrieux G_occipital_middle.lh** | 0.54 | 17127 |
| **rfMRI part corr ICA100 edge 163 link 4-8** | 0.54 | 15864 |
| **rfMRI part corr ICA100 edge 899 link 21-30** | 0.54 | 15864 |
| **rfMRI part corr ICA100 edge 1237 link 33-38** | 0.54 | 15864 |
| **rfMRI part corr ICA100 edge 742 link 16-53** | 0.54 | 15864 |
| **ICVF Cingulum-cingulate_gyrus-R** | 0.54 | 16540 |
| **rfMRI full corr ICA100 edge 704 link 15-54** | 0.54 | 15864 |
| **meandepth Morphologist FCMpost_left** | 0.54 | 18101 |
| **rfMRI full corr ICA100 edge 948 link 22-46** | 0.54 | 15864 |
| **MO Medial_lemniscus_L** | 0.54 | 16541 |
| **surface Morphologist SpC_right** | 0.54 | 16309 |
| **maxdepth Morphologist SLipost_left** | 0.54 | 17919 |
| **rfMRI part corr ICA25 edge 107 link 7-9** | 0.54 | 15864 |
| **meandepth Morphologist SGSM_left** | 0.54 | 12701 |
| **rfMRI full corr ICA100 edge 1470 link 49-55** | 0.54 | 15864 |
| **rfMRI part corr ICA25 edge 205 link 18-19** | 0.54 | 15864 |
| **rfMRI part corr ICA100 edge 1354 link 39-44** | 0.54 | 15864 |
| **rfMRI part corr ICA100 edge 1429 link 44-54** | 0.54 | 15864 |
| **ThickAvg Desikan precentral.lh** | 0.54 | 17127 |
| **rfMRI part corr ICA25 edge 73 link 4-20** | 0.54 | 15864 |
| **rfMRI part corr ICA100 edge 643 link 14-33** | 0.54 | 15864 |
| **rfMRI amplitude ICA100 component 26** | 0.54 | 15863 |
| **rfMRI part corr ICA100 edge 684 link 15-34** | 0.54 | 15864 |
| **ThickAvg Desikan lateralorbitofrontal.rh** | 0.54 | 17127 |
| **ISOVF Posterior_thalamic_radiation-include_optic** | 0.54 | 16527 |
| **rfMRI part corr ICA100 edge 977 link 23-43** | 0.54 | 15864 |
| **rfMRI full corr ICA100 edge 130 link 3-26** | 0.55 | 15864 |
| **rfMRI full corr ICA100 edge 882 link 20-47** | 0.55 | 15864 |
| **rfMRI full corr ICA25 edge 40 link 3-4** | 0.55 | 15864 |
| **rfMRI full corr ICA100 edge 336 link 7-34** | 0.55 | 15862 |
| **rfMRI full corr ICA100 edge 90 link 2-38** | 0.55 | 15864 |
| **meandepth Morphologist SCall_right** | 0.55 | 18085 |

| | | |
|---|---|---|
| **rfMRI full corr ICA100 edge 858 link 20-23** | 0.55 | 15864 |
| **rfMRI full corr ICA100 edge 766 link 17-39** | 0.55 | 15864 |
| **rfMRI full corr ICA100 edge 539 link 11-55** | 0.55 | 15860 |
| **rfMRI full corr ICA100 edge 588 link 13-19** | 0.55 | 15864 |
| **rfMRI part corr ICA100 edge 692 link 15-42** | 0.55 | 15864 |
| **rfMRI full corr ICA100 edge 946 link 22-44** | 0.55 | 15864 |
| **rfMRI part corr ICA100 edge 99 link 2-47** | 0.55 | 15864 |
| **rfMRI part corr ICA100 edge 49 link 1-50** | 0.55 | 15864 |
| **GM_thickness Morphologist STs_left** | 0.55 | 18092 |
| **ICVF Corticospinal_tract_L** | 0.55 | 16540 |
| **rfMRI part corr ICA100 edge 1269 link 34-49** | 0.55 | 15864 |
| **ISOVF Inferior_cerebellar_peduncle_R** | 0.55 | 16527 |
| **rfMRI part corr ICA100 edge 1017 link 24-52** | 0.55 | 15864 |
| **surface Morphologist SRh_right** | 0.55 | 17623 |
| **rfMRI full corr ICA100 edge 685 link 15-35** | 0.55 | 15864 |
| **rfMRI part corr ICA100 edge 1250 link 33-51** | 0.55 | 15864 |
| **meandepth Morphologist SFmedian_left** | 0.55 | 18078 |
| **rfMRI part corr ICA100 edge 258 link 5-53** | 0.55 | 15864 |
| **opening Morphologist FCLrscpost_right** | 0.55 | 16126 |
| **opening Morphologist SPeCsup_right** | 0.55 | 17272 |
| **rfMRI part corr ICA25 edge 202 link 17-19** | 0.55 | 15864 |
| **rfMRI full corr ICA100 edge 1279 link 35-39** | 0.55 | 15864 |
| **GM_thickness Morphologist SOTlatpost_left** | 0.55 | 18053 |
| **rfMRI full corr ICA25 edge 157 link 11-13** | 0.55 | 15864 |
| **rfMRI full corr ICA100 edge 226 link 5-21** | 0.55 | 15864 |
| **rfMRI part corr ICA100 edge 290 link 6-36** | 0.55 | 15864 |
| **rfMRI full corr ICA100 edge 670 link 15-20** | 0.55 | 15864 |
| **rfMRI part corr ICA100 edge 1256 link 34-36** | 0.55 | 15864 |
| **rfMRI part corr ICA100 edge 1039 link 25-44** | 0.55 | 15864 |
| **ICVF Superior_cerebellar_peduncle_R** | 0.55 | 16540 |
| **hull_junction_length Morphologist SpC_right** | 0.55 | 16309 |
| **rfMRI full corr ICA100 edge 1413 link 43-49** | 0.55 | 15864 |
| **rfMRI full corr ICA100 edge 206 link 4-51** | 0.55 | 15864 |
| **rfMRI part corr ICA100 edge 1148 link 29-43** | 0.55 | 15864 |

| | | |
|---|---|---|
| **opening Morphologist SOTlatint_right** | 0.55 | 15621 |
| **rfMRI part corr ICA100 edge 954 link 22-52** | 0.55 | 15864 |
| **ThickAvg Destrieux Lat_Fis-post.lh** | 0.55 | 17126 |
| **rfMRI full corr ICA100 edge 1457 link 47-55** | 0.55 | 15864 |
| **rfMRI part corr ICA100 edge 1087 link 27-35** | 0.55 | 15864 |
| **rfMRI part corr ICA100 edge 770 link 17-43** | 0.56 | 15864 |
| **rfMRI full corr ICA100 edge 421 link 9-26** | 0.56 | 15863 |
| **rfMRI full corr ICA100 edge 1264 link 34-44** | 0.56 | 15864 |
| **rfMRI part corr ICA100 edge 913 link 21-44** | 0.56 | 15864 |
| **rfMRI part corr ICA100 edge 657 link 14-47** | 0.56 | 15864 |
| **rfMRI part corr ICA100 edge 1184 link 30-54** | 0.56 | 15864 |
| **rfMRI full corr ICA100 edge 89 link 2-37** | 0.56 | 15864 |
| **rfMRI part corr ICA100 edge 783 link 18-19** | 0.56 | 15864 |
| **ICVF Anterior_corona_radiata_L** | 0.56 | 16540 |
| **hull_junction_length Morphologist SOlf_right** | 0.56 | 18091 |
| **GrayVol Destrieux S_oc_sup_and_transversal.rh** | 0.56 | 17127 |
| **rfMRI part corr ICA25 edge 8 link 1-9** | 0.56 | 15864 |
| **ISOVF Pontine_crossing_tract-a_part_of_MCP** | 0.56 | 16527 |
| **rfMRI full corr ICA100 edge 984 link 23-50** | 0.56 | 15864 |
| **rfMRI part corr ICA100 edge 559 link 12-32** | 0.56 | 15864 |
| **rfMRI part corr ICA100 edge 351 link 7-49** | 0.56 | 15864 |
| **rfMRI part corr ICA100 edge 248 link 5-43** | 0.56 | 15864 |
| **rfMRI part corr ICA100 edge 760 link 17-33** | 0.56 | 15864 |
| **rfMRI part corr ICA100 edge 422 link 9-27** | 0.56 | 15864 |
| **rfMRI full corr ICA100 edge 1479 link 51-55** | 0.56 | 15864 |
| **ISOVF Body_of_corpus_callosum** | 0.56 | 16527 |
| **ThickAvg Destrieux G_temp_sup-Plan_polar.lh** | 0.56 | 17126 |
| **rfMRI full corr ICA100 edge 474 link 10-34** | 0.56 | 15864 |
| **surface Morphologist SFint_right** | 0.56 | 18100 |
| **rfMRI part corr ICA100 edge 791 link 18-27** | 0.56 | 15864 |
| **surface Morphologist SFpolairetr_left** | 0.56 | 18062 |
| **rfMRI full corr ICA100 edge 1030 link 25-35** | 0.56 | 15864 |
| **SurfArea Destrieux G_pariet_inf-Supramar.lh** | 0.56 | 17127 |
| **rfMRI full corr ICA25 edge 105 link 6-21** | 0.56 | 15864 |

| | | |
|---|---|---|
| **rfMRI part corr ICA100 edge 1421 link 44-46** | 0.56 | 15864 |
| **ThickAvg Destrieux G_and_S_frontomargin.rh** | 0.56 | 17126 |
| **rfMRI full corr ICA100 edge 188 link 4-33** | 0.56 | 15864 |
| **rfMRI part corr ICA100 edge 999 link 24-34** | 0.56 | 15864 |
| **rfMRI part corr ICA100 edge 1221 link 32-44** | 0.56 | 15864 |
| **rfMRI full corr ICA100 edge 243 link 5-38** | 0.56 | 15864 |
| **ThickAvg Desikan lateraloccipital.rh** | 0.56 | 17127 |
| **rfMRI part corr ICA100 edge 743 link 16-54** | 0.56 | 15864 |
| **rfMRI full corr ICA100 edge 862 link 20-27** | 0.56 | 15864 |
| **GrayVol Desikan posteriorcingulate.rh** | 0.56 | 17127 |
| **MD Anterior_limb_of_internal_capsule_R** | 0.56 | 16541 |
| **rfMRI part corr ICA100 edge 673 link 15-23** | 0.56 | 15864 |
| **surface Morphologist SCall_right** | 0.56 | 18086 |
| **rfMRI part corr ICA100 edge 189 link 4-34** | 0.56 | 15864 |
| **rfMRI part corr ICA100 edge 1485 link 54-55** | 0.56 | 15864 |
| **rfMRI full corr ICA25 edge 172 link 12-19** | 0.57 | 15864 |
| **hull_junction_length Morphologist STsterascant_le** | 0.57 | 17560 |
| **rfMRI part corr ICA100 edge 866 link 20-31** | 0.57 | 15864 |
| **rfMRI amplitude ICA100 component 29** | 0.57 | 15863 |
| **rfMRI part corr ICA100 edge 761 link 17-34** | 0.57 | 15864 |
| **rfMRI part corr ICA100 edge 1082 link 27-30** | 0.57 | 15864 |
| **rfMRI part corr ICA100 edge 61 link 2-9** | 0.57 | 15864 |
| **GM_thickness Morphologist SCall_left** | 0.57 | 18055 |
| **rfMRI full corr ICA25 edge 148 link 10-14** | 0.57 | 15864 |
| **rfMRI full corr ICA100 edge 709 link 16-20** | 0.57 | 15862 |
| **rfMRI full corr ICA100 edge 1267 link 34-47** | 0.57 | 15864 |
| **rfMRI full corr ICA100 edge 337 link 7-35** | 0.57 | 15864 |
| **rfMRI part corr ICA100 edge 926 link 22-24** | 0.57 | 15864 |
| **rfMRI full corr ICA100 edge 1225 link 32-48** | 0.57 | 15862 |
| **rfMRI part corr ICA100 edge 1342 link 38-48** | 0.57 | 15864 |
| **rfMRI part corr ICA100 edge 1424 link 44-49** | 0.57 | 15864 |
| **rfMRI part corr ICA25 edge 46 link 3-10** | 0.57 | 15864 |
| **rfMRI part corr ICA25 edge 93 link 6-9** | 0.57 | 15864 |
| **rfMRI full corr ICA100 edge 614 link 13-45** | 0.57 | 15864 |

| | | |
|---|---|---|
| **rfMRI full corr ICA25 edge 76 link 5-7** | 0.57 | 15864 |
| **OD Fornix-column_and_body_of_fornix** | 0.57 | 16541 |
| **rfMRI full corr ICA100 edge 158 link 3-54** | 0.57 | 15864 |
| **rfMRI full corr ICA100 edge 1331 link 37-54** | 0.57 | 15864 |
| **rfMRI part corr ICA100 edge 725 link 16-36** | 0.57 | 15864 |
| **rfMRI full corr ICA25 edge 119 link 7-21** | 0.57 | 15864 |
| **meandepth Morphologist SPoCsup_left** | 0.57 | 17973 |
| **rfMRI part corr ICA100 edge 864 link 20-29** | 0.57 | 15864 |
| **rfMRI full corr ICA100 edge 951 link 22-49** | 0.57 | 15863 |
| **rfMRI part corr ICA100 edge 1401 link 42-49** | 0.57 | 15864 |
| **rfMRI full corr ICA100 edge 662 link 14-52** | 0.57 | 15862 |
| **rfMRI amplitude ICA100 component 24** | 0.57 | 15863 |
| **rfMRI part corr ICA100 edge 157 link 3-53** | 0.57 | 15864 |
| **FA Posterior_thalamic_radiation-include_optic_rad** | 0.57 | 16541 |
| **surface Morphologist FIPrint2_right** | 0.57 | 13064 |
| **meandepth Morphologist SOTlatant_left** | 0.57 | 18076 |
| **rfMRI full corr ICA100 edge 856 link 20-21** | 0.57 | 15864 |
| **rfMRI full corr ICA100 edge 414 link 9-19** | 0.57 | 15863 |
| **rfMRI part corr ICA100 edge 1442 link 46-48** | 0.57 | 15864 |
| **hull_junction_length Morphologist INSULA_right** | 0.57 | 18100 |
| **rfMRI full corr ICA100 edge 883 link 20-48** | 0.57 | 15864 |
| **SurfArea Destrieux G_front_inf-Opercular.rh** | 0.57 | 17127 |
| **rfMRI part corr ICA100 edge 990 link 24-25** | 0.57 | 15864 |
| **maxdepth Morphologist FCalant-ScCal_right** | 0.57 | 18100 |
| **GrayVol Destrieux S_collat_transv_post.lh** | 0.57 | 17125 |
| **rfMRI full corr ICA100 edge 1385 link 41-46** | 0.57 | 15864 |
| **rfMRI part corr ICA100 edge 1479 link 51-55** | 0.57 | 15864 |
| **rfMRI part corr ICA100 edge 139 link 3-35** | 0.57 | 15864 |
| **rfMRI amplitude ICA25 component 3** | 0.57 | 15863 |
| **rfMRI part corr ICA100 edge 661 link 14-51** | 0.57 | 15864 |
| **rfMRI full corr ICA100 edge 1395 link 42-43** | 0.57 | 15864 |
| **rfMRI full corr ICA100 edge 978 link 23-44** | 0.57 | 15864 |
| **rfMRI part corr ICA100 edge 473 link 10-33** | 0.57 | 15864 |
| **maxdepth Morphologist SFmedian_left** | 0.57 | 18078 |

| | | |
|---|---|---|
| **SurfArea Destrieux S_front_middle.rh** | 0.57 | 17127 |
| **rfMRI full corr ICA100 edge 1402 link 42-50** | 0.57 | 15864 |
| **rfMRI part corr ICA100 edge 610 link 13-41** | 0.58 | 15864 |
| **rfMRI full corr ICA100 edge 922 link 21-53** | 0.58 | 15862 |
| **rfMRI full corr ICA100 edge 1156 link 29-51** | 0.58 | 15864 |
| **rfMRI part corr ICA100 edge 429 link 9-34** | 0.58 | 15864 |
| **rfMRI part corr ICA100 edge 118 link 3-14** | 0.58 | 15864 |
| **maxdepth Morphologist SpC_left** | 0.58 | 17043 |
| **rfMRI full corr ICA25 edge 42 link 3-6** | 0.58 | 15864 |
| **rfMRI part corr ICA100 edge 174 link 4-19** | 0.58 | 15864 |
| **rfMRI part corr ICA100 edge 272 link 6-18** | 0.58 | 15864 |
| **rfMRI full corr ICA100 edge 1230 link 32-53** | 0.58 | 15864 |
| **rfMRI full corr ICA100 edge 1077 link 26-53** | 0.58 | 15863 |
| **rfMRI part corr ICA100 edge 230 link 5-25** | 0.58 | 15864 |
| **rfMRI full corr ICA100 edge 181 link 4-26** | 0.58 | 15864 |
| **rfMRI full corr ICA100 edge 527 link 11-43** | 0.58 | 15861 |
| **GrayVol Destrieux G_pariet_inf-Angular.rh** | 0.58 | 17127 |
| **rfMRI full corr ICA100 edge 150 link 3-46** | 0.58 | 15864 |
| **rfMRI part corr ICA100 edge 848 link 19-48** | 0.58 | 15864 |
| **rfMRI full corr ICA100 edge 1362 link 39-52** | 0.58 | 15864 |
| **GrayVol Destrieux Lat_Fis-ant-Horizont.lh** | 0.58 | 17126 |
| **ThickAvg Desikan inferiortemporal.rh** | 0.58 | 17127 |
| **rfMRI part corr ICA100 edge 1304 link 36-45** | 0.58 | 15864 |
| **rfMRI full corr ICA25 edge 59 link 4-6** | 0.58 | 15864 |
| **rfMRI full corr ICA100 edge 1257 link 34-37** | 0.58 | 15864 |
| **rfMRI part corr ICA100 edge 658 link 14-48** | 0.58 | 15864 |
| **rfMRI part corr ICA100 edge 1140 link 29-35** | 0.58 | 15864 |
| **ThickAvg Destrieux G_orbital.rh** | 0.58 | 17127 |
| **opening Morphologist FIPPoCinf_right** | 0.58 | 18073 |
| **SurfArea Destrieux G_and_S_cingul-Mid-Post.rh** | 0.58 | 17127 |
| **rfMRI full corr ICA100 edge 1330 link 37-53** | 0.58 | 15864 |
| **GM_thickness Morphologist STsterascpost_left** | 0.58 | 17893 |
| **ICVF Fornix-cres-Stria_terminalis-not_resolved_w** | 0.58 | 16540 |
| **rfMRI part corr ICA100 edge 1375 link 40-50** | 0.58 | 15864 |

| | | |
|---|---|---|
| **rfMRI full corr ICA100 edge 355 link 7-53** | 0.58 | 15864 |
| **rfMRI part corr ICA100 edge 821 link 19-21** | 0.58 | 15864 |
| **rfMRI part corr ICA25 edge 63 link 4-10** | 0.58 | 15864 |
| **rfMRI part corr ICA100 edge 1301 link 36-42** | 0.58 | 15864 |
| **rfMRI part corr ICA100 edge 181 link 4-26** | 0.58 | 15864 |
| **rfMRI full corr ICA100 edge 360 link 8-11** | 0.58 | 15864 |
| **rfMRI full corr ICA100 edge 983 link 23-49** | 0.58 | 15864 |
| **GrayVol Desikan lateralorbitofrontal.rh** | 0.58 | 17127 |
| **ThickAvg Destrieux G_pariet_inf-Angular.rh** | 0.58 | 17127 |
| **ICVF Posterior_thalamic_radiation-include_optic_r** | 0.58 | 16540 |
| **rfMRI part corr ICA100 edge 43 link 1-44** | 0.58 | 15864 |
| **GrayVol Destrieux S_oc-temp_lat.lh** | 0.58 | 17127 |
| **rfMRI amplitude ICA25 component 13** | 0.58 | 15863 |
| **rfMRI full corr ICA100 edge 1409 link 43-45** | 0.58 | 15864 |
| **maxdepth Morphologist FCLrscant_left** | 0.58 | 4323 |
| **rfMRI full corr ICA100 edge 217 link 5-12** | 0.58 | 15864 |
| **rfMRI full corr ICA100 edge 278 link 6-24** | 0.59 | 15864 |
| **rfMRI part corr ICA100 edge 1114 link 28-35** | 0.59 | 15864 |
| **rfMRI full corr ICA100 edge 865 link 20-30** | 0.59 | 15864 |
| **rfMRI part corr ICA100 edge 1073 link 26-49** | 0.59 | 15864 |
| **rfMRI full corr ICA100 edge 795 link 18-31** | 0.59 | 15864 |
| **ThickAvg Desikan superiorfrontal.rh** | 0.59 | 17127 |
| **maxdepth Morphologist STsterascpost_right** | 0.59 | 18048 |
| **rfMRI part corr ICA100 edge 976 link 23-42** | 0.59 | 15864 |
| **rfMRI part corr ICA100 edge 1408 link 43-44** | 0.59 | 15864 |
| **rfMRI full corr ICA25 edge 141 link 9-18** | 0.59 | 15864 |
| **rfMRI part corr ICA25 edge 62 link 4-9** | 0.59 | 15864 |
| **rfMRI full corr ICA100 edge 723 link 16-34** | 0.59 | 15864 |
| **GrayVol Destrieux G_and_S_frontomargin.lh** | 0.59 | 17127 |
| **rfMRI full corr ICA100 edge 292 link 6-38** | 0.59 | 15864 |
| **hull_junction_length Morphologist SsP_right** | 0.59 | 18094 |
| **rfMRI full corr ICA100 edge 1070 link 26-46** | 0.59 | 15864 |
| **SurfArea Destrieux S_precentral-inf-part.lh** | 0.59 | 17127 |
| **rfMRI full corr ICA25 edge 68 link 4-15** | 0.59 | 15864 |

| | | |
|---|---|---|
| **rfMRI full corr ICA100 edge 71 link 2-19** | 0.59 | 15864 |
| **rfMRI part corr ICA25 edge 181 link 13-20** | 0.59 | 15864 |
| **SurfArea Desikan medialorbitofrontal.rh** | 0.59 | 17126 |
| **hull_junction_length Morphologist OCCIPITAL_rigl** | 0.59 | 18096 |
| **rfMRI full corr ICA25 edge 206 link 18-20** | 0.59 | 15864 |
| **hull_junction_length Morphologist SOTlatint_left** | 0.59 | 16594 |
| **rfMRI full corr ICA100 edge 693 link 15-43** | 0.59 | 15864 |
| **rfMRI part corr ICA100 edge 805 link 18-41** | 0.59 | 15864 |
| **rfMRI part corr ICA100 edge 984 link 23-50** | 0.59 | 15864 |
| **rfMRI part corr ICA100 edge 214 link 5-9** | 0.59 | 15864 |
| **rfMRI part corr ICA100 edge 480 link 10-40** | 0.59 | 15864 |
| **rfMRI full corr ICA100 edge 944 link 22-42** | 0.59 | 15864 |
| **rfMRI part corr ICA100 edge 845 link 19-45** | 0.59 | 15864 |
| **GrayVol Destrieux S_collat_transv_ant.rh** | 0.59 | 17125 |
| **rfMRI part corr ICA25 edge 19 link 1-20** | 0.59 | 15864 |
| **hull_junction_length Morphologist SForbitaire_rigl** | 0.59 | 17279 |
| **opening Morphologist SPasup_right** | 0.59 | 17400 |
| **rfMRI full corr ICA100 edge 722 link 16-33** | 0.59 | 15864 |
| **rfMRI full corr ICA100 edge 966 link 23-32** | 0.59 | 15863 |
| **rfMRI full corr ICA100 edge 21 link 1-22** | 0.59 | 15864 |
| **GM_thickness Morphologist FCalant-ScCal_right** | 0.59 | 18092 |
| **hull_junction_length Morphologist FCLrant_left** | 0.59 | 17708 |
| **rfMRI part corr ICA100 edge 1043 link 25-48** | 0.59 | 15864 |
| **maxdepth Morphologist SCLPC_right** | 0.59 | 5279 |
| **rfMRI full corr ICA100 edge 540 link 12-13** | 0.59 | 15863 |
| **rfMRI part corr ICA100 edge 41 link 1-42** | 0.6 | 15864 |
| **SurfArea Destrieux G_subcallosal.lh** | 0.6 | 17125 |
| **SurfArea Destrieux S_oc-temp_lat.rh** | 0.6 | 17127 |
| **rfMRI amplitude ICA100 component 46** | 0.6 | 15863 |
| **hull_junction_length Morphologist SPeCmarginal_** | 0.6 | 15613 |
| **hull_junction_length Morphologist SFsup_left** | 0.6 | 18101 |
| **hull_junction_length Morphologist SOTlatmed_rigl** | 0.6 | 17219 |
| **rfMRI full corr ICA100 edge 942 link 22-40** | 0.6 | 15863 |
| **ThickAvg Destrieux S_front_middle.lh** | 0.6 | 17127 |

| | | |
|---|---|---|
| **rfMRI full corr ICA25 edge 190 link 15-16** | 0.6 | 15864 |
| **maxdepth Morphologist SPeCinf_left** | 0.6 | 17300 |
| **rfMRI full corr ICA100 edge 328 link 7-26** | 0.6 | 15864 |
| **rfMRI part corr ICA100 edge 937 link 22-35** | 0.6 | 15864 |
| **rfMRI full corr ICA100 edge 318 link 7-16** | 0.6 | 15864 |
| **rfMRI part corr ICA100 edge 655 link 14-45** | 0.6 | 15864 |
| **rfMRI full corr ICA25 edge 25 link 2-7** | 0.6 | 15864 |
| **rfMRI part corr ICA100 edge 1466 link 49-51** | 0.6 | 15864 |
| **rfMRI amplitude ICA100 component 17** | 0.6 | 15863 |
| **MO Superior_longitudinal_fasciculus_L** | 0.6 | 16541 |
| **rfMRI full corr ICA100 edge 755 link 17-28** | 0.6 | 15864 |
| **ThickAvg Destrieux G_oc-temp_med-Parahip.rh** | 0.6 | 17127 |
| **ISOVF Posterior_corona_radiata_L** | 0.6 | 16527 |
| **rfMRI full corr ICA100 edge 518 link 11-34** | 0.6 | 15864 |
| **rfMRI amplitude ICA100 component 32** | 0.6 | 15863 |
| **rfMRI full corr ICA100 edge 634 link 14-24** | 0.6 | 15864 |
| **rfMRI full corr ICA100 edge 24 link 1-25** | 0.6 | 15864 |
| **surface Morphologist SForbitaire_right** | 0.6 | 17279 |
| **rfMRI full corr ICA100 edge 929 link 22-27** | 0.6 | 15864 |
| **rfMRI part corr ICA100 edge 345 link 7-43** | 0.6 | 15864 |
| **rfMRI full corr ICA25 edge 20 link 1-21** | 0.6 | 15864 |
| **rfMRI part corr ICA25 edge 151 link 10-17** | 0.6 | 15864 |
| **OD Anterior_corona_radiata_L** | 0.6 | 16541 |
| **rfMRI full corr ICA100 edge 93 link 2-41** | 0.6 | 15864 |
| **rfMRI part corr ICA100 edge 549 link 12-22** | 0.6 | 15864 |
| **SurfArea Desikan caudalanteriorcingulate.lh** | 0.6 | 17126 |
| **rfMRI full corr ICA100 edge 1236 link 33-37** | 0.6 | 15864 |
| **rfMRI full corr ICA100 edge 452 link 10-12** | 0.6 | 15864 |
| **GrayVol Destrieux G_orbital.rh** | 0.6 | 17127 |
| **GM_thickness Morphologist SPeCmedian_left** | 0.6 | 16793 |
| **rfMRI amplitude ICA100 component 10** | 0.6 | 15863 |
| **opening Morphologist FCLa_left** | 0.6 | 18059 |
| **ISOVF Fornix-cres-Stria_terminalis-not_resolved_** | 0.6 | 16527 |
| **rfMRI part corr ICA100 edge 834 link 19-34** | 0.6 | 15864 |

| | | |
|---|---|---|
| **rfMRI full corr ICA25 edge 209 link 19-21** | 0.6 | 15864 |
| **rfMRI part corr ICA100 edge 509 link 11-25** | 0.6 | 15864 |
| **SurfArea Desikan parsorbitalis.rh** | 0.6 | 17127 |
| **SurfArea Destrieux G_and_S_cingul-Ant.rh** | 0.6 | 17127 |
| **rfMRI part corr ICA100 edge 751 link 17-24** | 0.6 | 15864 |
| **ISOVF Splenium_of_corpus_callosum** | 0.6 | 16527 |
| **rfMRI part corr ICA100 edge 438 link 9-43** | 0.6 | 15864 |
| **MD Anterior_corona_radiata_L** | 0.6 | 16541 |
| **rfMRI part corr ICA100 edge 8 link 1-9** | 0.61 | 15864 |
| **rfMRI full corr ICA100 edge 1418 link 43-54** | 0.61 | 15864 |
| **rfMRI part corr ICA100 edge 371 link 8-22** | 0.61 | 15864 |
| **rfMRI part corr ICA100 edge 38 link 1-39** | 0.61 | 15864 |
| **hull_junction_length Morphologist SOr_right** | 0.61 | 18098 |
| **ThickAvg Desikan medialorbitofrontal.lh** | 0.61 | 17127 |
| **rfMRI part corr ICA100 edge 543 link 12-16** | 0.61 | 15864 |
| **rfMRI full corr ICA100 edge 692 link 15-42** | 0.61 | 15864 |
| **maxdepth Morphologist SPoCsup_left** | 0.61 | 17976 |
| **rfMRI part corr ICA100 edge 406 link 9-11** | 0.61 | 15864 |
| **rfMRI part corr ICA100 edge 1001 link 24-36** | 0.61 | 15864 |
| **rfMRI full corr ICA100 edge 1185 link 30-55** | 0.61 | 15864 |
| **rfMRI part corr ICA25 edge 172 link 12-19** | 0.61 | 15864 |
| **rfMRI part corr ICA100 edge 1213 link 32-36** | 0.61 | 15864 |
| **surface Morphologist SFsup_left** | 0.61 | 18101 |
| **rfMRI part corr ICA100 edge 1028 link 25-33** | 0.61 | 15864 |
| **rfMRI full corr ICA25 edge 16 link 1-17** | 0.61 | 15864 |
| **rfMRI part corr ICA100 edge 525 link 11-41** | 0.61 | 15864 |
| **rfMRI part corr ICA100 edge 519 link 11-35** | 0.61 | 15864 |
| **maxdepth Morphologist INSULA_left** | 0.61 | 18101 |
| **rfMRI full corr ICA100 edge 1259 link 34-39** | 0.61 | 15864 |
| **rfMRI part corr ICA100 edge 966 link 23-32** | 0.61 | 15864 |
| **rfMRI part corr ICA25 edge 80 link 5-11** | 0.61 | 15864 |
| **rfMRI part corr ICA100 edge 580 link 12-53** | 0.61 | 15864 |
| **ThickAvg Destrieux S_central.rh** | 0.61 | 17127 |
| **rfMRI part corr ICA100 edge 277 link 6-23** | 0.61 | 15864 |

| | | |
|---|---|---|
| **rfMRI part corr ICA100 edge 1346 link 38-52** | 0.61 | 15864 |
| **rfMRI full corr ICA100 edge 1318 link 37-41** | 0.61 | 15864 |
| **rfMRI full corr ICA100 edge 231 link 5-26** | 0.61 | 15863 |
| **rfMRI full corr ICA100 edge 82 link 2-30** | 0.61 | 15863 |
| **rfMRI part corr ICA100 edge 161 link 4-6** | 0.61 | 15864 |
| **rfMRI full corr ICA100 edge 439 link 9-44** | 0.61 | 15863 |
| **rfMRI part corr ICA100 edge 82 link 2-30** | 0.61 | 15864 |
| **rfMRI part corr ICA100 edge 135 link 3-31** | 0.61 | 15864 |
| **hull_junction_length Morphologist SFinf_right** | 0.61 | 18080 |
| **rfMRI part corr ICA100 edge 870 link 20-35** | 0.61 | 15864 |
| **SurfArea Desikan posteriorcingulate.lh** | 0.61 | 17126 |
| **rfMRI full corr ICA100 edge 1370 link 40-45** | 0.61 | 15864 |
| **rfMRI full corr ICA100 edge 731 link 16-42** | 0.61 | 15864 |
| **rfMRI part corr ICA100 edge 484 link 10-44** | 0.61 | 15864 |
| **rfMRI full corr ICA100 edge 631 link 14-21** | 0.61 | 15863 |
| **MD Uncinate_fasciculus_R** | 0.61 | 16541 |
| **rfMRI full corr ICA100 edge 981 link 23-47** | 0.61 | 15864 |
| **rfMRI part corr ICA100 edge 534 link 11-50** | 0.61 | 15864 |
| **rfMRI part corr ICA25 edge 88 link 5-19** | 0.61 | 15864 |
| **rfMRI full corr ICA100 edge 1338 link 38-44** | 0.61 | 15864 |
| **GM_thickness Morphologist SPeCinf_right** | 0.61 | 16595 |
| **rfMRI full corr ICA100 edge 767 link 17-40** | 0.61 | 15864 |
| **ISOVF Cingulum-hippocampus-R** | 0.62 | 16527 |
| **rfMRI full corr ICA100 edge 557 link 12-30** | 0.62 | 15864 |
| **rfMRI full corr ICA100 edge 1178 link 30-48** | 0.62 | 15864 |
| **rfMRI part corr ICA100 edge 1356 link 39-46** | 0.62 | 15864 |
| **rfMRI part corr ICA100 edge 312 link 7-10** | 0.62 | 15864 |
| **SurfArea Destrieux S_temporal_inf.lh** | 0.62 | 17127 |
| **rfMRI part corr ICA100 edge 1068 link 26-44** | 0.62 | 15864 |
| **maxdepth Morphologist SRinf_left** | 0.62 | 16697 |
| **GM_thickness Morphologist SForbitaire_left** | 0.62 | 17510 |
| **rfMRI full corr ICA100 edge 307 link 6-53** | 0.62 | 15856 |
| **rfMRI full corr ICA25 edge 51 link 3-15** | 0.62 | 15864 |
| **FA Sagittal_stratum-inf_longitidinal_fasci_and_inf_** | 0.62 | 16541 |

| | | |
|---|---|---|
| **maxdepth Morphologist SOr_right** | 0.62 | 18098 |
| **surface Morphologist SPoCsup_right** | 0.62 | 17842 |
| **MD Pontine_crossing_tract-a_part_of_MCP** | 0.62 | 16541 |
| **rfMRI full corr ICA100 edge 730 link 16-41** | 0.62 | 15864 |
| **rfMRI part corr ICA100 edge 1210 link 32-33** | 0.62 | 15864 |
| **rfMRI part corr ICA100 edge 366 link 8-17** | 0.62 | 15864 |
| **meandepth Morphologist SFinf_left** | 0.62 | 18101 |
| **rfMRI part corr ICA100 edge 1107 link 27-55** | 0.62 | 15864 |
| **rfMRI full corr ICA100 edge 1138 link 29-33** | 0.62 | 15864 |
| **rfMRI full corr ICA100 edge 349 link 7-47** | 0.62 | 15864 |
| **rfMRI full corr ICA100 edge 103 link 2-51** | 0.62 | 15864 |
| **rfMRI full corr ICA100 edge 161 link 4-6** | 0.62 | 15864 |
| **rfMRI part corr ICA100 edge 1358 link 39-48** | 0.62 | 15864 |
| **rfMRI part corr ICA100 edge 288 link 6-34** | 0.62 | 15864 |
| **rfMRI part corr ICA100 edge 1264 link 34-44** | 0.62 | 15864 |
| **surface Morphologist FCLa_left** | 0.62 | 18059 |
| **rfMRI full corr ICA100 edge 124 link 3-20** | 0.62 | 15864 |
| **rfMRI part corr ICA100 edge 854 link 19-54** | 0.62 | 15864 |
| **GrayVol Desikan precentral.lh** | 0.62 | 17127 |
| **rfMRI part corr ICA100 edge 898 link 21-29** | 0.62 | 15864 |
| **rfMRI full corr ICA100 edge 236 link 5-31** | 0.62 | 15864 |
| **rfMRI full corr ICA100 edge 657 link 14-47** | 0.62 | 15864 |
| **GrayVol Desikan entorhinal.rh** | 0.62 | 17125 |
| **rfMRI full corr ICA100 edge 1204 link 31-50** | 0.62 | 15864 |
| **rfMRI full corr ICA100 edge 1036 link 25-41** | 0.62 | 15864 |
| **maxdepth Morphologist STipost_right** | 0.62 | 18093 |
| **ThickAvg Destrieux S_oc_middle_and_Lunatus.lh** | 0.62 | 17126 |
| **meandepth Morphologist SFinter_left** | 0.62 | 18101 |
| **rfMRI part corr ICA100 edge 1286 link 35-46** | 0.62 | 15864 |
| **ISOVF Superior_fronto-occipital_fasciculus-part_c** | 0.62 | 16527 |
| **hull_junction_length Morphologist FIPPoCinf_left** | 0.62 | 18079 |
| **rfMRI full corr ICA100 edge 752 link 17-25** | 0.62 | 15863 |
| **rfMRI amplitude ICA100 component 16** | 0.63 | 15863 |
| **rfMRI full corr ICA100 edge 373 link 8-24** | 0.63 | 15864 |

| | | |
|---|---|---|
| **rfMRI part corr ICA100 edge 1451 link 47-49** | 0.63 | 15864 |
| **ThickAvg Destrieux S_circular_insula_ant.rh** | 0.63 | 17125 |
| **rfMRI part corr ICA100 edge 599 link 13-30** | 0.63 | 15864 |
| **meandepth Morphologist FCLa_right** | 0.63 | 18089 |
| **rfMRI full corr ICA25 edge 165 link 11-21** | 0.63 | 15864 |
| **surface Morphologist SOTlatant_right** | 0.63 | 18047 |
| **rfMRI full corr ICA100 edge 283 link 6-29** | 0.63 | 15864 |
| **maxdepth Morphologist SFinter_left** | 0.63 | 18101 |
| **MD Cingulum-cingulate_gyrus-L** | 0.63 | 16541 |
| **rfMRI amplitude ICA100 component 34** | 0.63 | 15863 |
| **rfMRI full corr ICA100 edge 831 link 19-31** | 0.63 | 15863 |
| **rfMRI full corr ICA100 edge 1344 link 38-50** | 0.63 | 15863 |
| **ThickAvg Desikan insula.lh** | 0.63 | 17127 |
| **rfMRI full corr ICA100 edge 782 link 17-55** | 0.63 | 15864 |
| **rfMRI part corr ICA100 edge 1340 link 38-46** | 0.63 | 15864 |
| **rfMRI full corr ICA100 edge 1439 link 45-54** | 0.63 | 15864 |
| **rfMRI full corr ICA100 edge 152 link 3-48** | 0.63 | 15864 |
| **rfMRI part corr ICA100 edge 269 link 6-15** | 0.63 | 15864 |
| **rfMRI full corr ICA100 edge 1463 link 48-54** | 0.63 | 15864 |
| **rfMRI amplitude ICA100 component 33** | 0.63 | 15863 |
| **rfMRI part corr ICA25 edge 42 link 3-6** | 0.63 | 15864 |
| **rfMRI full corr ICA100 edge 720 link 16-31** | 0.63 | 15864 |
| **MD Body_of_corpus_callosum** | 0.63 | 16541 |
| **SurfArea Destrieux S_orbital-H_Shaped.lh** | 0.63 | 17127 |
| **rfMRI full corr ICA100 edge 1361 link 39-51** | 0.63 | 15864 |
| **rfMRI full corr ICA25 edge 8 link 1-9** | 0.63 | 15864 |
| **rfMRI full corr ICA100 edge 424 link 9-29** | 0.63 | 15863 |
| **meandepth Morphologist SOr_left** | 0.63 | 18100 |
| **SurfArea Desikan fusiform.lh** | 0.63 | 17127 |
| **rfMRI part corr ICA100 edge 667 link 15-17** | 0.63 | 15864 |
| **rfMRI part corr ICA100 edge 1200 link 31-46** | 0.63 | 15864 |
| **rfMRI part corr ICA100 edge 440 link 9-45** | 0.63 | 15864 |
| **SurfArea Destrieux S_cingul-Marginalis.rh** | 0.63 | 17127 |
| **rfMRI part corr ICA100 edge 1290 link 35-50** | 0.63 | 15864 |

| | | |
|---|---|---|
| **rfMRI part corr ICA100 edge 1482 link 52-55** | 0.63 | 15864 |
| **rfMRI full corr ICA100 edge 304 link 6-50** | 0.63 | 15864 |
| **rfMRI part corr ICA100 edge 1448 link 46-54** | 0.63 | 15864 |
| **rfMRI part corr ICA100 edge 1147 link 29-42** | 0.63 | 15864 |
| **maxdepth Morphologist SPasup_left** | 0.63 | 16537 |
| **rfMRI full corr ICA100 edge 481 link 10-41** | 0.63 | 15864 |
| **rfMRI part corr ICA100 edge 904 link 21-35** | 0.64 | 15864 |
| **rfMRI full corr ICA100 edge 366 link 8-17** | 0.64 | 15864 |
| **rfMRI full corr ICA100 edge 1242 link 33-43** | 0.64 | 15862 |
| **rfMRI full corr ICA100 edge 691 link 15-41** | 0.64 | 15864 |
| **MO Posterior_corona_radiata_L** | 0.64 | 16541 |
| **rfMRI part corr ICA100 edge 1123 link 28-44** | 0.64 | 15864 |
| **rfMRI full corr ICA100 edge 1237 link 33-38** | 0.64 | 15864 |
| **rfMRI full corr ICA100 edge 227 link 5-22** | 0.64 | 15864 |
| **rfMRI part corr ICA100 edge 1419 link 43-55** | 0.64 | 15864 |
| **rfMRI part corr ICA100 edge 647 link 14-37** | 0.64 | 15864 |
| **SurfArea Destrieux S_oc-temp_lat.lh** | 0.64 | 17127 |
| **rfMRI full corr ICA100 edge 1474 link 50-54** | 0.64 | 15864 |
| **rfMRI full corr ICA100 edge 99 link 2-47** | 0.64 | 15864 |
| **rfMRI full corr ICA25 edge 170 link 12-17** | 0.64 | 15864 |
| **ICVF Uncinate_fasciculus_R** | 0.64 | 16540 |
| **rfMRI full corr ICA100 edge 1106 link 27-54** | 0.64 | 15864 |
| **rfMRI full corr ICA100 edge 810 link 18-46** | 0.64 | 15864 |
| **rfMRI full corr ICA100 edge 472 link 10-32** | 0.64 | 15864 |
| **rfMRI full corr ICA100 edge 1243 link 33-44** | 0.64 | 15864 |
| **rfMRI full corr ICA100 edge 202 link 4-47** | 0.64 | 15863 |
| **rfMRI part corr ICA100 edge 232 link 5-27** | 0.64 | 15864 |
| **SurfArea Desikan middletemporal.rh** | 0.64 | 17127 |
| **rfMRI part corr ICA100 edge 273 link 6-19** | 0.64 | 15864 |
| **rfMRI full corr ICA25 edge 203 link 17-20** | 0.64 | 15864 |
| **rfMRI full corr ICA100 edge 1169 link 30-39** | 0.64 | 15864 |
| **opening Morphologist SCall_right** | 0.64 | 18086 |
| **rfMRI part corr ICA100 edge 806 link 18-42** | 0.64 | 15864 |
| **rfMRI part corr ICA100 edge 110 link 3-6** | 0.64 | 15864 |

| | | |
|---|---|---|
| **GrayVol Destrieux G_Ins_lg_and_S_cent_ins.lh** | 0.64 | 17126 |
| **rfMRI part corr ICA100 edge 242 link 5-37** | 0.64 | 15864 |
| **rfMRI part corr ICA100 edge 836 link 19-36** | 0.64 | 15864 |
| **rfMRI part corr ICA100 edge 170 link 4-15** | 0.64 | 15864 |
| **rfMRI full corr ICA100 edge 141 link 3-37** | 0.64 | 15860 |
| **rfMRI part corr ICA100 edge 1261 link 34-41** | 0.64 | 15864 |
| **maxdepth Morphologist SForbitaire_right** | 0.64 | 17279 |
| **rfMRI full corr ICA100 edge 841 link 19-41** | 0.64 | 15864 |
| **rfMRI part corr ICA100 edge 970 link 23-36** | 0.64 | 15864 |
| **ThickAvg Destrieux S_front_sup.rh** | 0.64 | 17127 |
| **rfMRI part corr ICA100 edge 27 link 1-28** | 0.64 | 15864 |
| **rfMRI full corr ICA100 edge 1029 link 25-34** | 0.64 | 15862 |
| **rfMRI part corr ICA100 edge 150 link 3-46** | 0.64 | 15864 |
| **rfMRI part corr ICA100 edge 1223 link 32-46** | 0.64 | 15864 |
| **rfMRI part corr ICA25 edge 84 link 5-15** | 0.64 | 15864 |
| **rfMRI part corr ICA100 edge 1095 link 27-43** | 0.64 | 15864 |
| **rfMRI part corr ICA100 edge 561 link 12-34** | 0.64 | 15864 |
| **rfMRI part corr ICA100 edge 1212 link 32-35** | 0.64 | 15864 |
| **rfMRI full corr ICA100 edge 317 link 7-15** | 0.64 | 15864 |
| **rfMRI part corr ICA100 edge 562 link 12-35** | 0.64 | 15864 |
| **rfMRI full corr ICA100 edge 618 link 13-49** | 0.64 | 15864 |
| **ISOVF Posterior_limb_of_internal_capsule_R** | 0.64 | 16527 |
| **MD Superior_cerebellar_peduncle_L** | 0.64 | 16541 |
| **rfMRI part corr ICA100 edge 7 link 1-8** | 0.64 | 15864 |
| **rfMRI full corr ICA100 edge 887 link 20-52** | 0.64 | 15864 |
| **GrayVol Desikan lateraloccipital.lh** | 0.64 | 17127 |
| **rfMRI part corr ICA100 edge 617 link 13-48** | 0.64 | 15864 |
| **rfMRI part corr ICA100 edge 1410 link 43-46** | 0.64 | 15864 |
| **SurfArea Desikan transversetemporal.rh** | 0.65 | 17125 |
| **rfMRI part corr ICA100 edge 437 link 9-42** | 0.65 | 15864 |
| **rfMRI full corr ICA100 edge 271 link 6-17** | 0.65 | 15864 |
| **rfMRI full corr ICA100 edge 163 link 4-8** | 0.65 | 15864 |
| **rfMRI part corr ICA100 edge 818 link 18-54** | 0.65 | 15864 |
| **rfMRI full corr ICA25 edge 117 link 7-19** | 0.65 | 15863 |

| | | |
|---|---|---|
| **maxdepth Morphologist SOr_left** | 0.65 | 18100 |
| **rfMRI full corr ICA100 edge 1056 link 26-32** | 0.65 | 15864 |
| **rfMRI full corr ICA100 edge 607 link 13-38** | 0.65 | 15864 |
| **maxdepth Morphologist FCLrdiag_right** | 0.65 | 11519 |
| **rfMRI part corr ICA100 edge 435 link 9-40** | 0.65 | 15864 |
| **rfMRI part corr ICA100 edge 1278 link 35-38** | 0.65 | 15864 |
| **rfMRI part corr ICA100 edge 1190 link 31-36** | 0.65 | 15864 |
| **rfMRI part corr ICA100 edge 844 link 19-44** | 0.65 | 15864 |
| **GM_thickness Morphologist FCLrretroCtr_left** | 0.65 | 16820 |
| **rfMRI full corr ICA100 edge 1424 link 44-49** | 0.65 | 15864 |
| **rfMRI part corr ICA100 edge 268 link 6-14** | 0.65 | 15864 |
| **rfMRI part corr ICA100 edge 1329 link 37-52** | 0.65 | 15864 |
| **rfMRI part corr ICA100 edge 807 link 18-43** | 0.65 | 15864 |
| **ISOVF Genu_of_corpus_callosum** | 0.65 | 16527 |
| **rfMRI part corr ICA100 edge 669 link 15-19** | 0.65 | 15864 |
| **rfMRI full corr ICA100 edge 164 link 4-9** | 0.65 | 15864 |
| **rfMRI full corr ICA100 edge 29 link 1-30** | 0.65 | 15864 |
| **ThickAvg Destrieux G_cingul-Post-dorsal.lh** | 0.65 | 17125 |
| **ThickAvg Destrieux S_precentral-sup-part.rh** | 0.65 | 17127 |
| **meandepth Morphologist SPeCsup_left** | 0.65 | 17127 |
| **surface Morphologist SLipost_left** | 0.65 | 17919 |
| **rfMRI part corr ICA100 edge 358 link 8-9** | 0.65 | 15864 |
| **rfMRI part corr ICA100 edge 1472 link 50-52** | 0.65 | 15864 |
| **rfMRI full corr ICA100 edge 118 link 3-14** | 0.65 | 15864 |
| **rfMRI part corr ICA100 edge 1413 link 43-49** | 0.65 | 15864 |
| **rfMRI part corr ICA100 edge 875 link 20-40** | 0.65 | 15864 |
| **ISOVF Corticospinal_tract_R** | 0.65 | 16527 |
| **maxdepth Morphologist SsP_left** | 0.65 | 18098 |
| **hull_junction_length Morphologist SFsup_right** | 0.65 | 18100 |
| **rfMRI part corr ICA100 edge 797 link 18-33** | 0.65 | 15864 |
| **GM_thickness Morphologist OCCIPITAL_right** | 0.65 | 18088 |
| **rfMRI full corr ICA100 edge 632 link 14-22** | 0.65 | 15864 |
| **hull_junction_length Morphologist FCLrscpost_rig** | 0.65 | 16126 |
| **rfMRI full corr ICA100 edge 377 link 8-28** | 0.65 | 15864 |

| | | |
|---|---|---|
| **opening Morphologist FCLrasc_right** | 0.65 | 17670 |
| **maxdepth Morphologist SGSM_left** | 0.65 | 12712 |
| **rfMRI full corr ICA100 edge 803 link 18-39** | 0.65 | 15864 |
| **meandepth Morphologist SFinter_right** | 0.65 | 18099 |
| **rfMRI part corr ICA25 edge 72 link 4-19** | 0.65 | 15864 |
| **rfMRI full corr ICA100 edge 168 link 4-13** | 0.65 | 15864 |
| **SurfArea Desikan inferiorparietal.rh** | 0.65 | 17127 |
| **ISOVF Superior_cerebellar_peduncle_L** | 0.66 | 16527 |
| **rfMRI part corr ICA100 edge 701 link 15-51** | 0.66 | 15864 |
| **rfMRI part corr ICA100 edge 1397 link 42-45** | 0.66 | 15864 |
| **rfMRI full corr ICA100 edge 1 link 1-2** | 0.66 | 15864 |
| **GrayVol Desikan superiortemporal.lh** | 0.66 | 17127 |
| **rfMRI part corr ICA100 edge 1205 link 31-51** | 0.66 | 15864 |
| **maxdepth Morphologist FCMpost_left** | 0.66 | 18101 |
| **FA Cerebral_peduncle_R** | 0.66 | 16541 |
| **rfMRI part corr ICA100 edge 1367 link 40-42** | 0.66 | 15864 |
| **rfMRI part corr ICA100 edge 915 link 21-46** | 0.66 | 15864 |
| **surface Morphologist STipost_right** | 0.66 | 18093 |
| **rfMRI full corr ICA100 edge 57 link 2-5** | 0.66 | 15863 |
| **rfMRI full corr ICA100 edge 326 link 7-24** | 0.66 | 15864 |
| **GM_thickness Morphologist SCall_right** | 0.66 | 18079 |
| **rfMRI full corr ICA100 edge 825 link 19-25** | 0.66 | 15862 |
| **rfMRI full corr ICA100 edge 754 link 17-27** | 0.66 | 15864 |
| **rfMRI full corr ICA100 edge 561 link 12-34** | 0.66 | 15863 |
| **rfMRI full corr ICA100 edge 345 link 7-43** | 0.66 | 15864 |
| **surface Morphologist SFmedian_right** | 0.66 | 18080 |
| **rfMRI part corr ICA100 edge 819 link 18-55** | 0.66 | 15864 |
| **rfMRI part corr ICA100 edge 589 link 13-20** | 0.66 | 15864 |
| **GM_thickness Morphologist FCLa_right** | 0.66 | 18090 |
| **rfMRI part corr ICA100 edge 1415 link 43-51** | 0.66 | 15864 |
| **ThickAvg Desikan caudalanteriorcingulate.rh** | 0.66 | 17127 |
| **rfMRI part corr ICA100 edge 1076 link 26-52** | 0.66 | 15864 |
| **rfMRI part corr ICA100 edge 1297 link 36-38** | 0.66 | 15864 |
| **rfMRI full corr ICA25 edge 192 link 15-18** | 0.66 | 15864 |

| | | |
|---|---|---|
| **maxdepth Morphologist SFinf_left** | 0.66 | 18101 |
| **GM_thickness Morphologist SOTlatpost_right** | 0.66 | 18067 |
| **MO Posterior_thalamic_radiation-include_optic_ra** | 0.66 | 16541 |
| **rfMRI part corr ICA100 edge 1382 link 41-43** | 0.66 | 15864 |
| **rfMRI part corr ICA100 edge 1475 link 50-55** | 0.66 | 15864 |
| **rfMRI part corr ICA100 edge 533 link 11-49** | 0.66 | 15864 |
| **rfMRI full corr ICA25 edge 116 link 7-18** | 0.66 | 15864 |
| **rfMRI part corr ICA100 edge 1431 link 45-46** | 0.66 | 15864 |
| **rfMRI full corr ICA100 edge 241 link 5-36** | 0.66 | 15864 |
| **rfMRI full corr ICA25 edge 111 link 7-13** | 0.66 | 15864 |
| **rfMRI part corr ICA100 edge 1247 link 33-48** | 0.66 | 15864 |
| **rfMRI part corr ICA25 edge 29 link 2-11** | 0.66 | 15864 |
| **rfMRI full corr ICA100 edge 156 link 3-52** | 0.66 | 15863 |
| **rfMRI part corr ICA25 edge 5 link 1-6** | 0.66 | 15864 |
| **rfMRI full corr ICA100 edge 489 link 10-49** | 0.66 | 15863 |
| **rfMRI part corr ICA100 edge 138 link 3-34** | 0.67 | 15864 |
| **rfMRI full corr ICA100 edge 1421 link 44-46** | 0.67 | 15864 |
| **rfMRI full corr ICA100 edge 737 link 16-48** | 0.67 | 15864 |
| **rfMRI part corr ICA25 edge 117 link 7-19** | 0.67 | 15864 |
| **rfMRI full corr ICA100 edge 577 link 12-50** | 0.67 | 15864 |
| **Right-VentralDC** | 0.67 | 17127 |
| **GrayVol Desikan fusiform.lh** | 0.67 | 17127 |
| **SurfArea Destrieux S_calcarine.lh** | 0.67 | 17127 |
| **rfMRI part corr ICA100 edge 1298 link 36-39** | 0.67 | 15864 |
| **ThickAvg Destrieux S_collat_transv_ant.rh** | 0.67 | 17125 |
| **ThickAvg Destrieux G_front_sup.rh** | 0.67 | 17127 |
| **surface Morphologist SOTlatpost_right** | 0.67 | 18075 |
| **meandepth Morphologist INSULA_left** | 0.67 | 18101 |
| **rfMRI part corr ICA100 edge 321 link 7-19** | 0.67 | 15864 |
| **rfMRI full corr ICA25 edge 107 link 7-9** | 0.67 | 15864 |
| **rfMRI full corr ICA100 edge 327 link 7-25** | 0.67 | 15864 |
| **ThickAvg Destrieux S_precentral-inf-part.lh** | 0.67 | 17127 |
| **FA Corticospinal_tract_R** | 0.67 | 16541 |
| **rfMRI full corr ICA100 edge 343 link 7-41** | 0.67 | 15864 |

| | | |
|---|---|---|
| **rfMRI full corr ICA100 edge 533 link 11-49** | 0.67 | 15861 |
| **rfMRI full corr ICA100 edge 655 link 14-45** | 0.67 | 15864 |
| **SurfArea Desikan parsopercularis.lh** | 0.67 | 17127 |
| **rfMRI part corr ICA100 edge 209 link 4-54** | 0.67 | 15864 |
| **rfMRI full corr ICA100 edge 840 link 19-40** | 0.67 | 15864 |
| **rfMRI part corr ICA100 edge 1 link 1-2** | 0.67 | 15864 |
| **rfMRI full corr ICA100 edge 955 link 22-53** | 0.67 | 15864 |
| **GM_thickness Morphologist FIPPoCinf_left** | 0.67 | 18070 |
| **Left-Thalamus-Proper** | 0.67 | 17127 |
| **rfMRI part corr ICA100 edge 501 link 11-17** | 0.67 | 15864 |
| **MO Fornix-cres-Stria_terminalis-not_resolved_wit** | 0.67 | 16541 |
| **opening Morphologist SPoCsup_left** | 0.67 | 17976 |
| **opening Morphologist FCLrretroCtr_left** | 0.67 | 16831 |
| **rfMRI full corr ICA100 edge 1278 link 35-38** | 0.67 | 15864 |
| **rfMRI part corr ICA25 edge 163 link 11-19** | 0.67 | 15864 |
| **rfMRI full corr ICA100 edge 652 link 14-42** | 0.67 | 15864 |
| **rfMRI part corr ICA100 edge 1255 link 34-35** | 0.67 | 15864 |
| **rfMRI full corr ICA25 edge 5 link 1-6** | 0.67 | 15864 |
| **rfMRI full corr ICA100 edge 1261 link 34-41** | 0.67 | 15864 |
| **rfMRI full corr ICA100 edge 201 link 4-46** | 0.67 | 15864 |
| **rfMRI full corr ICA25 edge 100 link 6-16** | 0.67 | 15864 |
| **rfMRI full corr ICA100 edge 458 link 10-18** | 0.67 | 15864 |
| **ICVF Superior_longitudinal_fasciculus_R** | 0.67 | 16540 |
| **rfMRI part corr ICA100 edge 900 link 21-31** | 0.67 | 15864 |
| **GM_thickness Morphologist SPeCmarginal_left** | 0.68 | 15599 |
| **rfMRI full corr ICA25 edge 179 link 13-18** | 0.68 | 15864 |
| **rfMRI full corr ICA100 edge 102 link 2-50** | 0.68 | 15863 |
| **rfMRI part corr ICA100 edge 113 link 3-9** | 0.68 | 15864 |
| **rfMRI full corr ICA25 edge 108 link 7-10** | 0.68 | 15864 |
| **rfMRI amplitude ICA100 component 19** | 0.68 | 15863 |
| **rfMRI full corr ICA100 edge 868 link 20-33** | 0.68 | 15864 |
| **hull_junction_length Morphologist SOr_left** | 0.68 | 18100 |
| **rfMRI part corr ICA100 edge 1012 link 24-47** | 0.68 | 15864 |
| **rfMRI part corr ICA100 edge 60 link 2-8** | 0.68 | 15864 |

| | | |
|---|---|---|
| **rfMRI full corr ICA100 edge 735 link 16-46** | 0.68 | 15864 |
| **rfMRI part corr ICA100 edge 1339 link 38-45** | 0.68 | 15864 |
| **ThickAvg Destrieux S_orbital_med-olfact.lh** | 0.68 | 17126 |
| **rfMRI part corr ICA100 edge 402 link 8-53** | 0.68 | 15864 |
| **rfMRI full corr ICA100 edge 1308 link 36-49** | 0.68 | 15864 |
| **SurfArea Destrieux S_precentral-sup-part.lh** | 0.68 | 17127 |
| **rfMRI full corr ICA100 edge 500 link 11-16** | 0.68 | 15864 |
| **rfMRI part corr ICA100 edge 1361 link 39-51** | 0.68 | 15864 |
| **rfMRI full corr ICA100 edge 1020 link 24-55** | 0.68 | 15864 |
| **rfMRI full corr ICA100 edge 138 link 3-34** | 0.68 | 15864 |
| **rfMRI full corr ICA25 edge 134 link 9-11** | 0.68 | 15864 |
| **MD Inferior_cerebellar_peduncle_L** | 0.68 | 16541 |
| **hull_junction_length Morphologist FIP_right** | 0.68 | 18097 |
| **maxdepth Morphologist SFsup_left** | 0.68 | 18101 |
| **rfMRI part corr ICA100 edge 450 link 9-55** | 0.68 | 15864 |
| **rfMRI part corr ICA100 edge 192 link 4-37** | 0.68 | 15864 |
| **rfMRI full corr ICA25 edge 194 link 15-20** | 0.68 | 15864 |
| **rfMRI part corr ICA100 edge 1241 link 33-42** | 0.68 | 15864 |
| **rfMRI full corr ICA100 edge 555 link 12-28** | 0.68 | 15864 |
| **rfMRI part corr ICA100 edge 887 link 20-52** | 0.68 | 15864 |
| **rfMRI full corr ICA100 edge 1276 link 35-36** | 0.68 | 15864 |
| **rfMRI part corr ICA25 edge 109 link 7-11** | 0.68 | 15864 |
| **rfMRI full corr ICA100 edge 681 link 15-31** | 0.68 | 15864 |
| **GrayVol Destrieux S_front_inf.lh** | 0.68 | 17127 |
| **rfMRI full corr ICA100 edge 1069 link 26-45** | 0.68 | 15863 |
| **GM_thickness Morphologist OCCIPITAL_left** | 0.68 | 18088 |
| **rfMRI part corr ICA100 edge 726 link 16-37** | 0.68 | 15864 |
| **rfMRI full corr ICA100 edge 836 link 19-36** | 0.68 | 15864 |
| **rfMRI part corr ICA100 edge 542 link 12-15** | 0.68 | 15864 |
| **rfMRI part corr ICA100 edge 963 link 23-29** | 0.68 | 15864 |
| **surface Morphologist SOTlatint_left** | 0.68 | 16594 |
| **rfMRI part corr ICA100 edge 588 link 13-19** | 0.68 | 15864 |
| **rfMRI full corr ICA100 edge 27 link 1-28** | 0.68 | 15864 |
| **hull_junction_length Morphologist FCLrretroCtr_ri** | 0.68 | 16212 |

| | | |
|---|---|---|
| **rfMRI full corr ICA100 edge 1360 link 39-50** | 0.68 | 15864 |
| **rfMRI part corr ICA100 edge 686 link 15-36** | 0.68 | 15864 |
| **rfMRI full corr ICA100 edge 1417 link 43-53** | 0.68 | 15864 |
| **rfMRI full corr ICA100 edge 957 link 22-55** | 0.68 | 15864 |
| **surface Morphologist SPeCinter_right** | 0.69 | 18084 |
| **SurfArea Destrieux G_temp_sup-Plan_tempo.lh** | 0.69 | 17127 |
| **rfMRI part corr ICA100 edge 1380 link 40-55** | 0.69 | 15864 |
| **ThickAvg Destrieux S_temporal_transverse.rh** | 0.69 | 17124 |
| **rfMRI full corr ICA100 edge 1368 link 40-43** | 0.69 | 15864 |
| **rfMRI full corr ICA100 edge 1464 link 48-55** | 0.69 | 15864 |
| **ISOVF Corticospinal_tract_L** | 0.69 | 16527 |
| **ThickAvg Destrieux S_front_middle.rh** | 0.69 | 17127 |
| **maxdepth Morphologist SRh_right** | 0.69 | 17623 |
| **rfMRI full corr ICA25 edge 113 link 7-15** | 0.69 | 15864 |
| **rfMRI part corr ICA100 edge 649 link 14-39** | 0.69 | 15864 |
| **rfMRI full corr ICA100 edge 867 link 20-32** | 0.69 | 15864 |
| **rfMRI full corr ICA25 edge 146 link 10-12** | 0.69 | 15864 |
| **rfMRI part corr ICA100 edge 871 link 20-36** | 0.69 | 15864 |
| **opening Morphologist SPaint_right** | 0.69 | 18007 |
| **rfMRI full corr ICA25 edge 210 link 20-21** | 0.69 | 15864 |
| **rfMRI part corr ICA100 edge 1271 link 34-51** | 0.69 | 15864 |
| **rfMRI part corr ICA100 edge 594 link 13-25** | 0.69 | 15864 |
| **rfMRI part corr ICA100 edge 944 link 22-42** | 0.69 | 15864 |
| **rfMRI full corr ICA100 edge 877 link 20-42** | 0.69 | 15864 |
| **ICVF Cerebral_peduncle_L** | 0.69 | 16540 |
| **hull_junction_length Morphologist SFmarginal_left** | 0.69 | 18044 |
| **ThickAvg Desikan parstriangularis.lh** | 0.69 | 17127 |
| **rfMRI part corr ICA100 edge 1023 link 25-28** | 0.69 | 15864 |
| **rfMRI part corr ICA100 edge 282 link 6-28** | 0.69 | 15864 |
| **rfMRI part corr ICA100 edge 401 link 8-52** | 0.69 | 15864 |
| **hull_junction_length Morphologist FCLrasc_right** | 0.69 | 17670 |
| **rfMRI part corr ICA100 edge 934 link 22-32** | 0.69 | 15864 |
| **rfMRI part corr ICA100 edge 1093 link 27-41** | 0.69 | 15864 |
| **ThickAvg Desikan isthmuscingulate.lh** | 0.69 | 17126 |

| | | |
|---|---|---|
| **SurfArea Destrieux G_cingul-Post-dorsal.rh** | 0.69 | 17127 |
| **rfMRI full corr ICA25 edge 58 link 4-5** | 0.69 | 15864 |
| **rfMRI part corr ICA25 edge 86 link 5-17** | 0.69 | 15864 |
| **rfMRI full corr ICA100 edge 312 link 7-10** | 0.69 | 15864 |
| **FA Superior_cerebellar_peduncle_L** | 0.69 | 16541 |
| **rfMRI full corr ICA100 edge 1312 link 36-53** | 0.69 | 15864 |
| **rfMRI full corr ICA100 edge 1287 link 35-47** | 0.69 | 15864 |
| **rfMRI full corr ICA25 edge 54 link 3-18** | 0.7 | 15864 |
| **rfMRI part corr ICA100 edge 523 link 11-39** | 0.7 | 15864 |
| **rfMRI part corr ICA100 edge 877 link 20-42** | 0.7 | 15864 |
| **rfMRI full corr ICA25 edge 2 link 1-3** | 0.7 | 15858 |
| **rfMRI part corr ICA100 edge 300 link 6-46** | 0.7 | 15864 |
| **rfMRI part corr ICA100 edge 1368 link 40-43** | 0.7 | 15864 |
| **rfMRI part corr ICA100 edge 137 link 3-33** | 0.7 | 15864 |
| **rfMRI full corr ICA100 edge 719 link 16-30** | 0.7 | 15864 |
| **rfMRI full corr ICA100 edge 1380 link 40-55** | 0.7 | 15864 |
| **rfMRI full corr ICA100 edge 313 link 7-11** | 0.7 | 15861 |
| **rfMRI part corr ICA100 edge 436 link 9-41** | 0.7 | 15864 |
| **rfMRI part corr ICA100 edge 972 link 23-38** | 0.7 | 15864 |
| **GM_thickness Morphologist SCu_left** | 0.7 | 18048 |
| **rfMRI part corr ICA100 edge 697 link 15-47** | 0.7 | 15864 |
| **rfMRI part corr ICA100 edge 323 link 7-21** | 0.7 | 15864 |
| **rfMRI part corr ICA100 edge 448 link 9-53** | 0.7 | 15864 |
| **rfMRI full corr ICA100 edge 587 link 13-18** | 0.7 | 15863 |
| **GM_thickness Morphologist FCMant_right** | 0.7 | 17821 |
| **rfMRI part corr ICA100 edge 1327 link 37-50** | 0.7 | 15864 |
| **rfMRI part corr ICA100 edge 945 link 22-43** | 0.7 | 15864 |
| **meandepth Morphologist FCLp_left** | 0.7 | 18101 |
| **rfMRI part corr ICA100 edge 1284 link 35-44** | 0.7 | 15864 |
| **rfMRI part corr ICA100 edge 513 link 11-29** | 0.7 | 15864 |
| **rfMRI full corr ICA100 edge 586 link 13-17** | 0.7 | 15864 |
| **ThickAvg Destrieux S_collat_transv_post.lh** | 0.7 | 17125 |
| **rfMRI full corr ICA100 edge 1481 link 52-54** | 0.7 | 15864 |
| **rfMRI part corr ICA100 edge 1352 link 39-42** | 0.7 | 15864 |

| | | |
|---|---|---|
| rfMRI full corr ICA100 edge 1473 link 50-53 | 0.7 | 15864 |
| rfMRI part corr ICA100 edge 119 link 3-15 | 0.7 | 15864 |
| rfMRI part corr ICA100 edge 439 link 9-44 | 0.7 | 15864 |
| rfMRI full corr ICA100 edge 656 link 14-46 | 0.7 | 15864 |
| rfMRI part corr ICA100 edge 364 link 8-15 | 0.7 | 15864 |
| rfMRI part corr ICA100 edge 228 link 5-23 | 0.7 | 15864 |
| ThickAvg Destrieux S_oc-temp_lat.lh | 0.7 | 17127 |
| rfMRI part corr ICA100 edge 106 link 2-54 | 0.7 | 15864 |
| rfMRI full corr ICA100 edge 38 link 1-39 | 0.7 | 15864 |
| rfMRI full corr ICA100 edge 892 link 21-23 | 0.7 | 15864 |
| rfMRI full corr ICA100 edge 765 link 17-38 | 0.7 | 15864 |
| GrayVol Destrieux G_Ins_lg_and_S_cent_ins.rh | 0.7 | 17125 |
| ThickAvg Destrieux G_and_S_frontomargin.lh | 0.7 | 17127 |
| rfMRI part corr ICA100 edge 504 link 11-20 | 0.7 | 15864 |
| rfMRI part corr ICA100 edge 263 link 6-9 | 0.7 | 15864 |
| rfMRI full corr ICA100 edge 26 link 1-27 | 0.7 | 15864 |
| rfMRI full corr ICA100 edge 943 link 22-41 | 0.7 | 15864 |
| surface Morphologist STs_left | 0.7 | 18101 |
| rfMRI part corr ICA25 edge 165 link 11-21 | 0.71 | 15864 |
| rfMRI full corr ICA100 edge 1045 link 25-50 | 0.71 | 15864 |
| rfMRI full corr ICA100 edge 471 link 10-31 | 0.71 | 15864 |
| GM_thickness Morphologist SFmarginal_right | 0.71 | 17999 |
| rfMRI part corr ICA100 edge 430 link 9-35 | 0.71 | 15864 |
| rfMRI part corr ICA100 edge 94 link 2-42 | 0.71 | 15864 |
| MO Superior_fronto-occipital_fasciculus-part_of_a | 0.71 | 16541 |
| GM_thickness Morphologist SOr_right | 0.71 | 18089 |
| meandepth Morphologist SOTlatpost_right | 0.71 | 18075 |
| ICVF Pontine_crossing_tract-a_part_of_MCP | 0.71 | 16540 |
| opening Morphologist SOr_right | 0.71 | 18098 |
| rfMRI part corr ICA100 edge 555 link 12-28 | 0.71 | 15864 |
| rfMRI part corr ICA100 edge 1432 link 45-47 | 0.71 | 15864 |
| rfMRI part corr ICA100 edge 1217 link 32-40 | 0.71 | 15864 |
| rfMRI full corr ICA100 edge 881 link 20-46 | 0.71 | 15864 |
| rfMRI amplitude ICA100 component 36 | 0.71 | 15863 |

| | | |
|---|---|---|
| **rfMRI part corr ICA100 edge 1470 link 49-55** | 0.71 | 15864 |
| **rfMRI full corr ICA100 edge 788 link 18-24** | 0.71 | 15864 |
| **rfMRI part corr ICA100 edge 1294 link 35-54** | 0.71 | 15864 |
| **meandepth Morphologist SLipost_right** | 0.71 | 17975 |
| **maxdepth Morphologist SPeCinter_right** | 0.71 | 18084 |
| **rfMRI full corr ICA100 edge 769 link 17-42** | 0.71 | 15864 |
| **rfMRI part corr ICA100 edge 774 link 17-47** | 0.71 | 15864 |
| **CC_Central** | 0.71 | 17127 |
| **rfMRI full corr ICA100 edge 346 link 7-44** | 0.71 | 15863 |
| **rfMRI full corr ICA100 edge 901 link 21-32** | 0.71 | 15864 |
| **rfMRI full corr ICA100 edge 906 link 21-37** | 0.71 | 15864 |
| **rfMRI part corr ICA100 edge 634 link 14-24** | 0.71 | 15864 |
| **rfMRI part corr ICA100 edge 631 link 14-21** | 0.71 | 15864 |
| **rfMRI part corr ICA100 edge 365 link 8-16** | 0.71 | 15864 |
| **rfMRI part corr ICA100 edge 1042 link 25-47** | 0.71 | 15864 |
| **rfMRI full corr ICA100 edge 623 link 13-54** | 0.71 | 15863 |
| **rfMRI full corr ICA100 edge 10 link 1-11** | 0.71 | 15863 |
| **rfMRI part corr ICA25 edge 146 link 10-12** | 0.71 | 15864 |
| **rfMRI full corr ICA100 edge 144 link 3-40** | 0.71 | 15864 |
| **rfMRI part corr ICA100 edge 95 link 2-43** | 0.71 | 15864 |
| **rfMRI full corr ICA25 edge 103 link 6-19** | 0.71 | 15862 |
| **rfMRI full corr ICA100 edge 433 link 9-38** | 0.71 | 15864 |
| **maxdepth Morphologist FCLa_right** | 0.71 | 18097 |
| **GM_thickness Morphologist SFinter_right** | 0.72 | 18092 |
| **rfMRI part corr ICA100 edge 992 link 24-27** | 0.72 | 15864 |
| **rfMRI full corr ICA100 edge 1291 link 35-51** | 0.72 | 15864 |
| **opening Morphologist SCLPC_left** | 0.72 | 13421 |
| **Left-choroid-plexus** | 0.72 | 17127 |
| **rfMRI full corr ICA100 edge 928 link 22-26** | 0.72 | 15864 |
| **maxdepth Morphologist FCLrscant_right** | 0.72 | 5353 |
| **rfMRI part corr ICA100 edge 1075 link 26-51** | 0.72 | 15864 |
| **rfMRI part corr ICA100 edge 158 link 3-54** | 0.72 | 15864 |
| **rfMRI part corr ICA100 edge 837 link 19-37** | 0.72 | 15864 |
| **meandepth Morphologist FCMant_right** | 0.72 | 17828 |

| | | |
|---|---|---|
| **rfMRI part corr ICA25 edge 65 link 4-12** | 0.72 | 15864 |
| **opening Morphologist SCall_left** | 0.72 | 18064 |
| **opening Morphologist FCalant-ScCal_left** | 0.72 | 18100 |
| **rfMRI full corr ICA100 edge 1142 link 29-37** | 0.72 | 15863 |
| **rfMRI full corr ICA25 edge 106 link 7-8** | 0.72 | 15863 |
| **rfMRI part corr ICA100 edge 996 link 24-31** | 0.72 | 15864 |
| **rfMRI part corr ICA100 edge 527 link 11-43** | 0.72 | 15864 |
| **rfMRI part corr ICA100 edge 548 link 12-21** | 0.72 | 15864 |
| **ThickAvg Destrieux G_postcentral.rh** | 0.72 | 17127 |
| **rfMRI part corr ICA100 edge 1201 link 31-47** | 0.72 | 15864 |
| **rfMRI full corr ICA100 edge 1017 link 24-52** | 0.72 | 15864 |
| **Right-Lateral-Ventricle** | 0.72 | 17127 |
| **rfMRI part corr ICA100 edge 372 link 8-23** | 0.72 | 15864 |
| **rfMRI full corr ICA100 edge 727 link 16-38** | 0.72 | 15864 |
| **rfMRI part corr ICA100 edge 572 link 12-45** | 0.72 | 15864 |
| **rfMRI full corr ICA100 edge 454 link 10-14** | 0.72 | 15864 |
| **FA Fornix-cres-Stria_terminalis-not_resolved_with** | 0.72 | 16541 |
| **opening Morphologist SOp_right** | 0.72 | 17447 |
| **MD Cingulum-cingulate_gyrus-R** | 0.72 | 16541 |
| **ThickAvg Destrieux S_orbital-H_Shaped.lh** | 0.72 | 17127 |
| **surface Morphologist SPeCmedian_right** | 0.72 | 17074 |
| **rfMRI part corr ICA100 edge 48 link 1-49** | 0.72 | 15864 |
| **rfMRI part corr ICA100 edge 1308 link 36-49** | 0.72 | 15864 |
| **rfMRI part corr ICA100 edge 1056 link 26-32** | 0.72 | 15864 |
| **rfMRI full corr ICA100 edge 749 link 17-22** | 0.72 | 15864 |
| **rfMRI part corr ICA100 edge 1016 link 24-51** | 0.72 | 15864 |
| **ThickAvg Desikan parstriangularis.rh** | 0.72 | 17127 |
| **rfMRI part corr ICA100 edge 180 link 4-25** | 0.72 | 15864 |
| **rfMRI full corr ICA100 edge 368 link 8-19** | 0.72 | 15864 |
| **maxdepth Morphologist FCLrant_right** | 0.72 | 16205 |
| **rfMRI part corr ICA100 edge 802 link 18-38** | 0.72 | 15864 |
| **maxdepth Morphologist SFinter_right** | 0.72 | 18099 |
| **rfMRI full corr ICA100 edge 1265 link 34-45** | 0.72 | 15864 |
| **rfMRI part corr ICA100 edge 1428 link 44-53** | 0.73 | 15864 |

| | | |
|---|---|---|
| **rfMRI part corr ICA25 edge 12 link 1-13** | 0.73 | 15864 |
| **rfMRI full corr ICA100 edge 544 link 12-17** | 0.73 | 15864 |
| **rfMRI part corr ICA100 edge 969 link 23-35** | 0.73 | 15864 |
| **surface Morphologist SOTlatmed_left** | 0.73 | 17430 |
| **rfMRI full corr ICA100 edge 397 link 8-48** | 0.73 | 15863 |
| **surface Morphologist SRinf_left** | 0.73 | 16697 |
| **rfMRI full corr ICA25 edge 152 link 10-18** | 0.73 | 15864 |
| **rfMRI full corr ICA100 edge 1079 link 26-55** | 0.73 | 15864 |
| **GrayVol Desikan parsopercularis.rh** | 0.73 | 17127 |
| **rfMRI part corr ICA100 edge 659 link 14-49** | 0.73 | 15864 |
| **rfMRI part corr ICA25 edge 75 link 5-6** | 0.73 | 15864 |
| **rfMRI full corr ICA100 edge 1190 link 31-36** | 0.73 | 15864 |
| **rfMRI part corr ICA100 edge 1350 link 39-40** | 0.73 | 15864 |
| **rfMRI part corr ICA100 edge 129 link 3-25** | 0.73 | 15864 |
| **rfMRI full corr ICA100 edge 1223 link 32-46** | 0.73 | 15864 |
| **ThickAvg Destrieux Pole_occipital.lh** | 0.73 | 17127 |
| **rfMRI full corr ICA100 edge 1001 link 24-36** | 0.73 | 15864 |
| **rfMRI full corr ICA100 edge 95 link 2-43** | 0.73 | 15863 |
| **rfMRI full corr ICA25 edge 160 link 11-16** | 0.73 | 15864 |
| **rfMRI full corr ICA100 edge 372 link 8-23** | 0.73 | 15864 |
| **MO Fornix-column_and_body_of_fornix** | 0.73 | 16541 |
| **SurfArea Destrieux S_temporal_inf.rh** | 0.73 | 17127 |
| **rfMRI part corr ICA100 edge 315 link 7-13** | 0.73 | 15864 |
| **rfMRI part corr ICA100 edge 1270 link 34-50** | 0.73 | 15864 |
| **rfMRI full corr ICA100 edge 13 link 1-14** | 0.73 | 15864 |
| **SurfArea Desikan isthmuscingulate.rh** | 0.73 | 17127 |
| **GrayVol Destrieux S_precentral-sup-part.lh** | 0.73 | 17127 |
| **rfMRI part corr ICA100 edge 1319 link 37-42** | 0.73 | 15864 |
| **rfMRI full corr ICA100 edge 1219 link 32-42** | 0.73 | 15864 |
| **GrayVol Destrieux S_precentral-inf-part.lh** | 0.73 | 17127 |
| **rfMRI part corr ICA100 edge 80 link 2-28** | 0.73 | 15864 |
| **rfMRI part corr ICA100 edge 1383 link 41-44** | 0.73 | 15864 |
| **rfMRI full corr ICA100 edge 363 link 8-14** | 0.73 | 15864 |
| **rfMRI part corr ICA100 edge 1318 link 37-41** | 0.73 | 15864 |

| | | |
|---|---|---|
| **ICVF Inferior_cerebellar_peduncle_R** | 0.73 | 16540 |
| **opening Morphologist SFsup_left** | 0.73 | 18101 |
| **rfMRI part corr ICA100 edge 665 link 14-55** | 0.73 | 15864 |
| **rfMRI part corr ICA100 edge 577 link 12-50** | 0.73 | 15864 |
| **rfMRI full corr ICA100 edge 1128 link 28-49** | 0.73 | 15864 |
| **rfMRI full corr ICA100 edge 913 link 21-44** | 0.73 | 15861 |
| **rfMRI full corr ICA25 edge 22 link 2-4** | 0.73 | 15864 |
| **SurfArea Destrieux G_pariet_inf-Angular.rh** | 0.73 | 17127 |
| **GrayVol Destrieux G_subcallosal.rh** | 0.73 | 17126 |
| **rfMRI part corr ICA100 edge 826 link 19-26** | 0.73 | 15864 |
| **rfMRI full corr ICA100 edge 847 link 19-47** | 0.73 | 15864 |
| **rfMRI part corr ICA100 edge 1484 link 53-55** | 0.73 | 15864 |
| **rfMRI part corr ICA100 edge 428 link 9-33** | 0.73 | 15864 |
| **MD Inferior_cerebellar_peduncle_R** | 0.73 | 16541 |
| **rfMRI full corr ICA100 edge 1326 link 37-49** | 0.73 | 15864 |
| **ISOVF Middle_cerebellar_peduncle** | 0.74 | 16527 |
| **rfMRI part corr ICA100 edge 1386 link 41-47** | 0.74 | 15864 |
| **rfMRI full corr ICA100 edge 77 link 2-25** | 0.74 | 15864 |
| **rfMRI part corr ICA100 edge 1136 link 29-31** | 0.74 | 15864 |
| **rfMRI part corr ICA100 edge 586 link 13-17** | 0.74 | 15864 |
| **rfMRI part corr ICA100 edge 186 link 4-31** | 0.74 | 15864 |
| **ThickAvg Destrieux S_collat_transv_post.rh** | 0.74 | 17126 |
| **surface Morphologist SCsylvian_right** | 0.74 | 14992 |
| **rfMRI part corr ICA100 edge 521 link 11-37** | 0.74 | 15864 |
| **SurfArea Destrieux S_orbital_lateral.rh** | 0.74 | 17126 |
| **rfMRI full corr ICA100 edge 864 link 20-29** | 0.74 | 15864 |
| **ThickAvg Destrieux S_postcentral.lh** | 0.74 | 17127 |
| **rfMRI full corr ICA100 edge 1038 link 25-43** | 0.74 | 15864 |
| **rfMRI part corr ICA25 edge 111 link 7-13** | 0.74 | 15864 |
| **rfMRI part corr ICA25 edge 118 link 7-20** | 0.74 | 15864 |
| **MO Splenium_of_corpus_callosum** | 0.74 | 16541 |
| **ThickAvg Destrieux G_rectus.rh** | 0.74 | 17126 |
| **rfMRI full corr ICA100 edge 1187 link 31-33** | 0.74 | 15864 |
| **rfMRI full corr ICA100 edge 427 link 9-32** | 0.74 | 15864 |

| | | |
|---|---|---|
| **rfMRI part corr ICA100 edge 322 link 7-20** | 0.74 | 15864 |
| **ICVF Retrolenticular_part_of_internal_capsule_L** | 0.74 | 16540 |
| **GM_thickness Morphologist SFmedian_right** | 0.74 | 18073 |
| **rfMRI part corr ICA100 edge 786 link 18-22** | 0.74 | 15864 |
| **rfMRI full corr ICA100 edge 1286 link 35-46** | 0.74 | 15862 |
| **meandepth Morphologist FCLrant_right** | 0.74 | 16038 |
| **rfMRI full corr ICA25 edge 164 link 11-20** | 0.74 | 15864 |
| **rfMRI part corr ICA100 edge 175 link 4-20** | 0.74 | 15864 |
| **rfMRI part corr ICA25 edge 31 link 2-13** | 0.74 | 15864 |
| **rfMRI part corr ICA100 edge 1272 link 34-52** | 0.74 | 15864 |
| **rfMRI part corr ICA100 edge 1220 link 32-43** | 0.74 | 15864 |
| **ThickAvg Destrieux G_and_S_cingul-Ant.rh** | 0.74 | 17127 |
| **rfMRI part corr ICA100 edge 1427 link 44-52** | 0.74 | 15864 |
| **rfMRI full corr ICA100 edge 364 link 8-15** | 0.74 | 15864 |
| **rfMRI part corr ICA100 edge 829 link 19-29** | 0.74 | 15864 |
| **GM_thickness Morphologist SLipost_left** | 0.74 | 17910 |
| **rfMRI full corr ICA100 edge 1013 link 24-48** | 0.75 | 15864 |
| **rfMRI full corr ICA100 edge 702 link 15-52** | 0.75 | 15864 |
| **GrayVol Destrieux S_front_inf.rh** | 0.75 | 17127 |
| **rfMRI part corr ICA100 edge 1071 link 26-47** | 0.75 | 15864 |
| **GM_thickness Morphologist SOTlatmed_left** | 0.75 | 17419 |
| **Right-Hippocampus** | 0.75 | 17127 |
| **hull_junction_length Morphologist SOlf_left** | 0.75 | 18094 |
| **rfMRI full corr ICA100 edge 112 link 3-8** | 0.75 | 15864 |
| **meandepth Morphologist SRh_right** | 0.75 | 17622 |
| **hull_junction_length Morphologist STiant_right** | 0.75 | 18100 |
| **ICVF Superior_longitudinal_fasciculus_L** | 0.75 | 16540 |
| **GM_thickness Morphologist SFpolairetr_left** | 0.75 | 18053 |
| **rfMRI part corr ICA100 edge 550 link 12-23** | 0.75 | 15864 |
| **rfMRI full corr ICA100 edge 1110 link 28-31** | 0.75 | 15864 |
| **rfMRI full corr ICA100 edge 407 link 9-12** | 0.75 | 15864 |
| **hull_junction_length Morphologist FIPrint2_right** | 0.75 | 13064 |
| **rfMRI full corr ICA100 edge 406 link 9-11** | 0.75 | 15864 |
| **rfMRI full corr ICA100 edge 1039 link 25-44** | 0.75 | 15864 |

| | | |
|---|---|---|
| **rfMRI part corr ICA100 edge 1274 link 34-54** | 0.75 | 15864 |
| **rfMRI part corr ICA100 edge 1090 link 27-38** | 0.75 | 15864 |
| **opening Morphologist FPO_right** | 0.75 | 18097 |
| **rfMRI full corr ICA100 edge 846 link 19-46** | 0.75 | 15864 |
| **rfMRI full corr ICA25 edge 31 link 2-13** | 0.75 | 15860 |
| **opening Morphologist SsP_right** | 0.75 | 18094 |
| **hull_junction_length Morphologist FPO_left** | 0.75 | 18097 |
| **rfMRI part corr ICA25 edge 197 link 16-18** | 0.75 | 15864 |
| **rfMRI full corr ICA100 edge 1206 link 31-52** | 0.75 | 15864 |
| **rfMRI part corr ICA100 edge 1311 link 36-52** | 0.75 | 15864 |
| **rfMRI part corr ICA100 edge 1253 link 33-54** | 0.75 | 15864 |
| **rfMRI part corr ICA100 edge 14 link 1-15** | 0.75 | 15864 |
| **rfMRI part corr ICA100 edge 466 link 10-26** | 0.75 | 15864 |
| **rfMRI part corr ICA100 edge 1325 link 37-48** | 0.75 | 15864 |
| **rfMRI part corr ICA100 edge 1153 link 29-48** | 0.75 | 15864 |
| **GM_thickness Morphologist FPO_left** | 0.75 | 18088 |
| **rfMRI full corr ICA100 edge 110 link 3-6** | 0.75 | 15864 |
| **rfMRI full corr ICA25 edge 112 link 7-14** | 0.75 | 15864 |
| **rfMRI part corr ICA25 edge 153 link 10-19** | 0.75 | 15864 |
| **rfMRI full corr ICA100 edge 270 link 6-16** | 0.75 | 15864 |
| **rfMRI part corr ICA100 edge 1288 link 35-48** | 0.75 | 15864 |
| **meandepth Morphologist SFint_right** | 0.75 | 18100 |
| **rfMRI part corr ICA100 edge 1277 link 35-37** | 0.75 | 15864 |
| **rfMRI full corr ICA100 edge 1226 link 32-49** | 0.75 | 15864 |
| **rfMRI part corr ICA100 edge 1216 link 32-39** | 0.75 | 15864 |
| **rfMRI part corr ICA100 edge 1029 link 25-34** | 0.75 | 15864 |
| **rfMRI amplitude ICA25 component 1** | 0.75 | 15863 |
| **rfMRI full corr ICA100 edge 1050 link 25-55** | 0.75 | 15864 |
| **ThickAvg Destrieux S_front_inf.rh** | 0.76 | 17127 |
| **rfMRI part corr ICA100 edge 1289 link 35-49** | 0.76 | 15864 |
| **rfMRI full corr ICA100 edge 214 link 5-9** | 0.76 | 15863 |
| **rfMRI part corr ICA100 edge 620 link 13-51** | 0.76 | 15864 |
| **rfMRI full corr ICA100 edge 1047 link 25-52** | 0.76 | 15864 |
| **rfMRI full corr ICA100 edge 428 link 9-33** | 0.76 | 15864 |

| | | |
|---|---|---|
| **rfMRI full corr ICA100 edge 114 link 3-10** | 0.76 | 15864 |
| **rfMRI part corr ICA100 edge 360 link 8-11** | 0.76 | 15864 |
| **rfMRI part corr ICA100 edge 211 link 5-6** | 0.76 | 15864 |
| **rfMRI part corr ICA100 edge 929 link 22-27** | 0.76 | 15864 |
| **maxdepth Morphologist SCsylvian_right** | 0.76 | 14992 |
| **rfMRI full corr ICA100 edge 707 link 16-18** | 0.76 | 15864 |
| **rfMRI part corr ICA100 edge 715 link 16-26** | 0.76 | 15864 |
| **rfMRI full corr ICA100 edge 32 link 1-33** | 0.76 | 15864 |
| **ICVF Uncinate_fasciculus_L** | 0.76 | 16540 |
| **rfMRI part corr ICA100 edge 672 link 15-22** | 0.76 | 15864 |
| **SurfArea Destrieux G_and_S_cingul-Mid-Ant.lh** | 0.76 | 17127 |
| **ICVF Posterior_corona_radiata_R** | 0.76 | 16540 |
| **rfMRI part corr ICA25 edge 160 link 11-16** | 0.76 | 15864 |
| **GrayVol Desikan parsorbitalis.rh** | 0.76 | 17127 |
| **rfMRI part corr ICA25 edge 32 link 2-14** | 0.76 | 15864 |
| **rfMRI full corr ICA100 edge 251 link 5-46** | 0.76 | 15864 |
| **GrayVol Destrieux S_pericallosal.lh** | 0.76 | 17127 |
| **rfMRI full corr ICA100 edge 1198 link 31-44** | 0.76 | 15864 |
| **opening Morphologist SPeCsup_left** | 0.76 | 17143 |
| **rfMRI full corr ICA100 edge 64 link 2-12** | 0.76 | 15864 |
| **ThickAvg Destrieux S_orbital-H_Shaped.rh** | 0.76 | 17125 |
| **rfMRI full corr ICA100 edge 976 link 23-42** | 0.76 | 15864 |
| **ThickAvg Destrieux S_circular_insula_sup.rh** | 0.76 | 17126 |
| **ThickAvg Destrieux S_occipital_ant.lh** | 0.76 | 17127 |
| **rfMRI full corr ICA100 edge 1432 link 45-47** | 0.76 | 15864 |
| **ThickAvg Destrieux G_and_S_occipital_inf.lh** | 0.76 | 17127 |
| **rfMRI part corr ICA100 edge 1240 link 33-41** | 0.76 | 15864 |
| **rfMRI full corr ICA100 edge 463 link 10-23** | 0.76 | 15864 |
| **opening Morphologist STiant_left** | 0.76 | 18101 |
| **rfMRI full corr ICA100 edge 121 link 3-17** | 0.76 | 15864 |
| **rfMRI full corr ICA100 edge 1317 link 37-40** | 0.76 | 15864 |
| **rfMRI full corr ICA100 edge 1455 link 47-53** | 0.76 | 15864 |
| **rfMRI full corr ICA100 edge 927 link 22-25** | 0.76 | 15864 |
| **hull_junction_length Morphologist SPoCsup_right** | 0.76 | 17842 |

| | | |
|---|---|---|
| **rfMRI part corr ICA100 edge 891 link 21-22** | 0.76 | 15864 |
| **ThickAvg Destrieux G_occipital_middle.lh** | 0.76 | 17127 |
| **rfMRI part corr ICA25 edge 61 link 4-8** | 0.76 | 15864 |
| **rfMRI full corr ICA25 edge 138 link 9-15** | 0.76 | 15863 |
| **rfMRI full corr ICA100 edge 768 link 17-41** | 0.76 | 15864 |
| **rfMRI full corr ICA100 edge 222 link 5-17** | 0.76 | 15864 |
| **rfMRI part corr ICA100 edge 895 link 21-26** | 0.76 | 15864 |
| **rfMRI full corr ICA100 edge 245 link 5-40** | 0.76 | 15864 |
| **rfMRI full corr ICA100 edge 491 link 10-51** | 0.76 | 15864 |
| **rfMRI full corr ICA100 edge 465 link 10-25** | 0.76 | 15864 |
| **meandepth Morphologist FCLa_left** | 0.77 | 18009 |
| **rfMRI part corr ICA100 edge 318 link 7-16** | 0.77 | 15864 |
| **rfMRI full corr ICA100 edge 212 link 5-7** | 0.77 | 15864 |
| **maxdepth Morphologist STpol_left** | 0.77 | 18076 |
| **rfMRI full corr ICA100 edge 1363 link 39-53** | 0.77 | 15863 |
| **opening Morphologist SLipost_right** | 0.77 | 17978 |
| **SurfArea Destrieux G_rectus.lh** | 0.77 | 17127 |
| **rfMRI full corr ICA100 edge 596 link 13-27** | 0.77 | 15864 |
| **rfMRI full corr ICA25 edge 163 link 11-19** | 0.77 | 15864 |
| **rfMRI part corr ICA100 edge 1125 link 28-46** | 0.77 | 15864 |
| **rfMRI full corr ICA100 edge 1397 link 42-45** | 0.77 | 15864 |
| **surface Morphologist SFmarginal_left** | 0.77 | 18044 |
| **FA Posterior_corona_radiata_L** | 0.77 | 16541 |
| **rfMRI full corr ICA25 edge 197 link 16-18** | 0.77 | 15864 |
| **rfMRI part corr ICA100 edge 924 link 21-55** | 0.77 | 15864 |
| **opening Morphologist FCalant-ScCal_right** | 0.77 | 18100 |
| **surface Morphologist SPeCinter_left** | 0.77 | 18028 |
| **rfMRI part corr ICA100 edge 1414 link 43-50** | 0.77 | 15864 |
| **FA Cingulum-cingulate_gyrus-L** | 0.77 | 16541 |
| **rfMRI full corr ICA100 edge 140 link 3-36** | 0.77 | 15864 |
| **MO Genu_of_corpus_callosum** | 0.77 | 16541 |
| **surface Morphologist FCLrretroCtr_right** | 0.77 | 16212 |
| **rfMRI part corr ICA100 edge 582 link 12-55** | 0.77 | 15864 |
| **ThickAvg Desikan fusiform.rh** | 0.77 | 17127 |

| | | |
|---|---|---|
| **surface Morphologist STsterascpost_right** | 0.77 | 18048 |
| **rfMRI part corr ICA100 edge 266 link 6-12** | 0.77 | 15864 |
| **ThickAvg Destrieux G_insular_short.rh** | 0.77 | 17125 |
| **opening Morphologist FIP_left** | 0.77 | 18098 |
| **surface Morphologist SsP_left** | 0.77 | 18098 |
| **FA Anterior_corona_radiata_L** | 0.77 | 16541 |
| **ICVF Retrolenticular_part_of_internal_capsule_R** | 0.77 | 16540 |
| **rfMRI full corr ICA100 edge 1284 link 35-44** | 0.77 | 15864 |
| **rfMRI full corr ICA100 edge 1268 link 34-48** | 0.77 | 15864 |
| **rfMRI part corr ICA100 edge 935 link 22-33** | 0.77 | 15864 |
| **rfMRI part corr ICA100 edge 201 link 4-46** | 0.77 | 15864 |
| **rfMRI full corr ICA25 edge 34 link 2-16** | 0.77 | 15864 |
| **GM_thickness Morphologist SPeCsup_left** | 0.77 | 17131 |
| **rfMRI part corr ICA100 edge 539 link 11-55** | 0.77 | 15864 |
| **rfMRI part corr ICA100 edge 1280 link 35-40** | 0.77 | 15864 |
| **GM_thickness Morphologist SsP_left** | 0.77 | 18089 |
| **rfMRI part corr ICA100 edge 1208 link 31-54** | 0.77 | 15864 |
| **rfMRI part corr ICA100 edge 953 link 22-51** | 0.77 | 15864 |
| **rfMRI full corr ICA100 edge 436 link 9-41** | 0.77 | 15864 |
| **rfMRI full corr ICA100 edge 708 link 16-19** | 0.77 | 15861 |
| **surface Morphologist SOTlatmed_right** | 0.77 | 17219 |
| **rfMRI part corr ICA100 edge 831 link 19-31** | 0.77 | 15864 |
| **rfMRI part corr ICA25 edge 171 link 12-18** | 0.78 | 15864 |
| **rfMRI full corr ICA100 edge 824 link 19-24** | 0.78 | 15864 |
| **rfMRI full corr ICA100 edge 1315 link 37-38** | 0.78 | 15864 |
| **rfMRI full corr ICA100 edge 935 link 22-33** | 0.78 | 15863 |
| **rfMRI part corr ICA100 edge 682 link 15-32** | 0.78 | 15864 |
| **rfMRI part corr ICA100 edge 1193 link 31-39** | 0.78 | 15864 |
| **hull_junction_length Morphologist FCMant_right** | 0.78 | 17828 |
| **rfMRI full corr ICA25 edge 171 link 12-18** | 0.78 | 15864 |
| **rfMRI part corr ICA100 edge 863 link 20-28** | 0.78 | 15864 |
| **rfMRI part corr ICA100 edge 823 link 19-23** | 0.78 | 15864 |
| **ThickAvg Destrieux Lat_Fis-ant-Vertical.lh** | 0.78 | 17122 |
| **rfMRI part corr ICA100 edge 96 link 2-44** | 0.78 | 15864 |

| | | |
|---|---|---|
| **opening Morphologist FCLrretroCtr_right** | 0.78 | 16212 |
| **rfMRI part corr ICA100 edge 840 link 19-40** | 0.78 | 15864 |
| **FA Posterior_thalamic_radiation-include_optic_rad** | 0.78 | 16541 |
| **SurfArea Destrieux G_and_S_transv_frontopol.rh** | 0.78 | 17127 |
| **rfMRI full corr ICA25 edge 201 link 17-18** | 0.78 | 15864 |
| **rfMRI part corr ICA100 edge 336 link 7-34** | 0.78 | 15864 |
| **ThickAvg Destrieux S_oc-temp_med_and_Lingual.** | 0.78 | 17127 |
| **GrayVol Destrieux Pole_occipital.lh** | 0.78 | 17127 |
| **rfMRI part corr ICA100 edge 625 link 14-15** | 0.78 | 15864 |
| **rfMRI full corr ICA100 edge 405 link 9-10** | 0.78 | 15864 |
| **GM_thickness Morphologist FCLp_right** | 0.78 | 18093 |
| **rfMRI full corr ICA100 edge 664 link 14-54** | 0.78 | 15864 |
| **meandepth Morphologist STsterascpost_right** | 0.78 | 18045 |
| **rfMRI full corr ICA100 edge 821 link 19-21** | 0.78 | 15864 |
| **rfMRI part corr ICA100 edge 346 link 7-44** | 0.78 | 15864 |
| **rfMRI full corr ICA100 edge 1214 link 32-37** | 0.78 | 15864 |
| **GM_thickness Morphologist SPeCinf_left** | 0.78 | 17288 |
| **rfMRI full corr ICA100 edge 956 link 22-54** | 0.78 | 15864 |
| **GrayVol Desikan pericalcarine.lh** | 0.78 | 17127 |
| **rfMRI full corr ICA100 edge 445 link 9-50** | 0.78 | 15860 |
| **rfMRI full corr ICA100 edge 746 link 17-19** | 0.78 | 15864 |
| **meandepth Morphologist SFsup_left** | 0.78 | 18101 |
| **rfMRI part corr ICA100 edge 862 link 20-27** | 0.78 | 15864 |
| **MO Middle_cerebellar_peduncle** | 0.78 | 16541 |
| **rfMRI part corr ICA25 edge 201 link 17-18** | 0.78 | 15864 |
| **rfMRI part corr ICA100 edge 1384 link 41-45** | 0.78 | 15864 |
| **GM_thickness Morphologist SCu_right** | 0.78 | 18060 |
| **rfMRI full corr ICA100 edge 1104 link 27-52** | 0.78 | 15864 |
| **rfMRI part corr ICA100 edge 518 link 11-34** | 0.78 | 15864 |
| **rfMRI part corr ICA100 edge 332 link 7-30** | 0.78 | 15864 |
| **rfMRI part corr ICA100 edge 512 link 11-28** | 0.78 | 15864 |
| **rfMRI part corr ICA25 edge 124 link 8-13** | 0.78 | 15864 |
| **rfMRI full corr ICA100 edge 1428 link 44-53** | 0.78 | 15864 |
| **rfMRI amplitude ICA100 component 6** | 0.78 | 15863 |

| | | |
|---|---|---|
| rfMRI part corr ICA25 edge 192 link 15-18 | 0.78 | 15864 |
| rfMRI part corr ICA25 edge 49 link 3-13 | 0.79 | 15864 |
| rfMRI full corr ICA100 edge 462 link 10-22 | 0.79 | 15863 |
| ThickAvg Destrieux S_suborbital.rh | 0.79 | 17125 |
| rfMRI part corr ICA100 edge 957 link 22-55 | 0.79 | 15864 |
| opening Morphologist STipost_right | 0.79 | 18093 |
| rfMRI part corr ICA100 edge 159 link 3-55 | 0.79 | 15864 |
| rfMRI full corr ICA100 edge 2 link 1-3 | 0.79 | 15863 |
| OD Fornix-cres-Stria_terminalis-not_resolved_wit | 0.79 | 16541 |
| rfMRI part corr ICA100 edge 876 link 20-41 | 0.79 | 15864 |
| rfMRI full corr ICA100 edge 136 link 3-32 | 0.79 | 15864 |
| rfMRI full corr ICA100 edge 900 link 21-31 | 0.79 | 15864 |
| rfMRI part corr ICA100 edge 1326 link 37-49 | 0.79 | 15864 |
| GrayVol Destrieux G_front_inf-Orbital.rh | 0.79 | 17127 |
| rfMRI part corr ICA100 edge 849 link 19-49 | 0.79 | 15864 |
| rfMRI full corr ICA100 edge 677 link 15-27 | 0.79 | 15863 |
| rfMRI full corr ICA25 edge 198 link 16-19 | 0.79 | 15864 |
| ICVF Splenium_of_corpus_callosum | 0.79 | 16540 |
| rfMRI part corr ICA100 edge 1030 link 25-35 | 0.79 | 15864 |
| rfMRI full corr ICA100 edge 547 link 12-20 | 0.79 | 15864 |
| rfMRI part corr ICA100 edge 472 link 10-32 | 0.79 | 15864 |
| opening Morphologist INSULA_left | 0.79 | 18101 |
| MD Posterior_corona_radiata_L | 0.79 | 16541 |
| rfMRI part corr ICA100 edge 1119 link 28-40 | 0.79 | 15864 |
| rfMRI part corr ICA100 edge 1422 link 44-47 | 0.79 | 15864 |
| rfMRI full corr ICA100 edge 1136 link 29-31 | 0.79 | 15864 |
| rfMRI part corr ICA100 edge 708 link 16-19 | 0.79 | 15864 |
| ThickAvg Destrieux G_front_inf-Orbital.rh | 0.79 | 17127 |
| surface Morphologist FCLrretroCtr_left | 0.79 | 16831 |
| GrayVol Destrieux G_and_S_cingul-Ant.rh | 0.79 | 17127 |
| ThickAvg Desikan fusiform.lh | 0.79 | 17127 |
| rfMRI part corr ICA100 edge 812 link 18-48 | 0.79 | 15864 |
| rfMRI full corr ICA100 edge 774 link 17-47 | 0.79 | 15863 |
| rfMRI part corr ICA100 edge 956 link 22-54 | 0.79 | 15864 |

| | | |
|---|---|---|
| **surface Morphologist STipost_left** | 0.79 | 18096 |
| **rfMRI full corr ICA100 edge 1384 link 41-45** | 0.79 | 15864 |
| **rfMRI part corr ICA100 edge 520 link 11-36** | 0.79 | 15864 |
| **rfMRI part corr ICA25 edge 168 link 12-15** | 0.79 | 15864 |
| **rfMRI part corr ICA25 edge 59 link 4-6** | 0.79 | 15864 |
| **rfMRI full corr ICA100 edge 902 link 21-33** | 0.79 | 15864 |
| **hull_junction_length Morphologist SOTlatmed_left** | 0.79 | 17430 |
| **GM_thickness Morphologist SFinf_right** | 0.79 | 18073 |
| **GM_thickness Morphologist SFint_right** | 0.79 | 18093 |
| **rfMRI part corr ICA100 edge 460 link 10-20** | 0.79 | 15864 |
| **rfMRI full corr ICA100 edge 234 link 5-29** | 0.79 | 15864 |
| **rfMRI full corr ICA100 edge 1356 link 39-46** | 0.79 | 15864 |
| **rfMRI full corr ICA100 edge 261 link 6-7** | 0.79 | 15864 |
| **rfMRI full corr ICA100 edge 933 link 22-31** | 0.79 | 15864 |
| **Left-Lateral-Ventricle** | 0.79 | 17127 |
| **rfMRI full corr ICA25 edge 19 link 1-20** | 0.79 | 15864 |
| **rfMRI part corr ICA100 edge 1407 link 42-55** | 0.8 | 15864 |
| **OD Corticospinal_tract_R** | 0.8 | 16541 |
| **rfMRI part corr ICA25 edge 78 link 5-9** | 0.8 | 15864 |
| **rfMRI full corr ICA100 edge 1249 link 33-50** | 0.8 | 15864 |
| **rfMRI full corr ICA100 edge 1400 link 42-48** | 0.8 | 15864 |
| **SurfArea Destrieux S_subparietal.lh** | 0.8 | 17125 |
| **FA Sagittal_stratum-inf_longitidinal_fasci_and_inf_** | 0.8 | 16541 |
| **rfMRI full corr ICA100 edge 696 link 15-46** | 0.8 | 15864 |
| **rfMRI full corr ICA25 edge 72 link 4-19** | 0.8 | 15864 |
| **GrayVol Destrieux S_oc_middle_and_Lunatus.lh** | 0.8 | 17126 |
| **rfMRI full corr ICA100 edge 552 link 12-25** | 0.8 | 15864 |
| **rfMRI part corr ICA100 edge 1129 link 28-50** | 0.8 | 15864 |
| **rfMRI full corr ICA100 edge 1088 link 27-36** | 0.8 | 15862 |
| **rfMRI part corr ICA100 edge 511 link 11-27** | 0.8 | 15864 |
| **opening Morphologist FIPrint1_left** | 0.8 | 17282 |
| **GrayVol Desikan medialorbitofrontal.lh** | 0.8 | 17127 |
| **rfMRI full corr ICA100 edge 469 link 10-29** | 0.8 | 15864 |
| **opening Morphologist SFinf_left** | 0.8 | 18101 |

| | | |
|---|---|---|
| **rfMRI full corr ICA100 edge 1212 link 32-35** | 0.8 | 15863 |
| **rfMRI part corr ICA25 edge 30 link 2-12** | 0.8 | 15864 |
| **rfMRI part corr ICA25 edge 17 link 1-18** | 0.8 | 15864 |
| **rfMRI full corr ICA100 edge 387 link 8-38** | 0.8 | 15864 |
| **SurfArea Desikan posteriorcingulate.rh** | 0.8 | 17127 |
| **SurfArea Desikan lateraloccipital.lh** | 0.8 | 17127 |
| **GrayVol Destrieux S_orbital_lateral.rh** | 0.8 | 17126 |
| **rfMRI full corr ICA100 edge 1004 link 24-39** | 0.8 | 15864 |
| **rfMRI part corr ICA100 edge 909 link 21-40** | 0.8 | 15864 |
| **rfMRI full corr ICA25 edge 6 link 1-7** | 0.8 | 15864 |
| **rfMRI full corr ICA100 edge 1048 link 25-53** | 0.8 | 15864 |
| **rfMRI part corr ICA100 edge 795 link 18-31** | 0.8 | 15864 |
| **rfMRI part corr ICA100 edge 1222 link 32-45** | 0.8 | 15864 |
| **ThickAvg Destrieux G_orbital.lh** | 0.8 | 17127 |
| **rfMRI part corr ICA100 edge 1360 link 39-50** | 0.8 | 15864 |
| **hull_junction_length Morphologist SPoCsup_left** | 0.8 | 17976 |
| **rfMRI full corr ICA100 edge 990 link 24-25** | 0.8 | 15862 |
| **rfMRI part corr ICA100 edge 723 link 16-34** | 0.81 | 15864 |
| **rfMRI part corr ICA25 edge 100 link 6-16** | 0.81 | 15864 |
| **rfMRI full corr ICA100 edge 1160 link 29-55** | 0.81 | 15864 |
| **rfMRI full corr ICA100 edge 375 link 8-26** | 0.81 | 15864 |
| **rfMRI part corr ICA100 edge 627 link 14-17** | 0.81 | 15864 |
| **rfMRI full corr ICA100 edge 100 link 2-48** | 0.81 | 15864 |
| **GrayVol Desikan parsopercularis.lh** | 0.81 | 17127 |
| **rfMRI full corr ICA100 edge 213 link 5-8** | 0.81 | 15864 |
| **rfMRI full corr ICA100 edge 954 link 22-52** | 0.81 | 15864 |
| **rfMRI full corr ICA100 edge 994 link 24-29** | 0.81 | 15864 |
| **rfMRI full corr ICA100 edge 899 link 21-30** | 0.81 | 15864 |
| **rfMRI full corr ICA100 edge 1252 link 33-53** | 0.81 | 15864 |
| **rfMRI part corr ICA25 edge 115 link 7-17** | 0.81 | 15864 |
| **rfMRI full corr ICA100 edge 61 link 2-9** | 0.81 | 15864 |
| **rfMRI part corr ICA100 edge 931 link 22-29** | 0.81 | 15864 |
| **GrayVol Destrieux G_and_S_cingul-Ant.lh** | 0.81 | 17127 |
| **rfMRI full corr ICA100 edge 953 link 22-51** | 0.81 | 15864 |

| | | |
|---|---|---|
| **GrayVol Destrieux S_occipital_ant.lh** | 0.81 | 17127 |
| **rfMRI full corr ICA100 edge 589 link 13-20** | 0.81 | 15864 |
| **rfMRI full corr ICA100 edge 1207 link 31-53** | 0.81 | 15864 |
| **hull_junction_length Morphologist SsP_left** | 0.81 | 18098 |
| **rfMRI full corr ICA100 edge 904 link 21-35** | 0.81 | 15864 |
| **maxdepth Morphologist SC_right** | 0.81 | 18100 |
| **rfMRI full corr ICA100 edge 257 link 5-52** | 0.81 | 15864 |
| **rfMRI full corr ICA100 edge 918 link 21-49** | 0.81 | 15864 |
| **rfMRI full corr ICA100 edge 570 link 12-43** | 0.81 | 15864 |
| **rfMRI part corr ICA100 edge 537 link 11-53** | 0.81 | 15864 |
| **rfMRI full corr ICA100 edge 1116 link 28-37** | 0.81 | 15864 |
| **rfMRI part corr ICA100 edge 414 link 9-19** | 0.81 | 15864 |
| **rfMRI full corr ICA100 edge 559 link 12-32** | 0.81 | 15862 |
| **rfMRI full corr ICA100 edge 1263 link 34-43** | 0.81 | 15864 |
| **rfMRI full corr ICA100 edge 1129 link 28-50** | 0.81 | 15864 |
| **hull_junction_length Morphologist SOTlatint_right** | 0.81 | 15621 |
| **MO Anterior_corona_radiata_L** | 0.81 | 16541 |
| **rfMRI full corr ICA100 edge 1032 link 25-37** | 0.81 | 15864 |
| **rfMRI full corr ICA100 edge 738 link 16-49** | 0.81 | 15864 |
| **rfMRI part corr ICA25 edge 25 link 2-7** | 0.81 | 15864 |
| **rfMRI part corr ICA100 edge 1151 link 29-46** | 0.81 | 15864 |
| **rfMRI full corr ICA100 edge 1126 link 28-47** | 0.81 | 15864 |
| **rfMRI part corr ICA100 edge 1426 link 44-51** | 0.81 | 15864 |
| **GrayVol Desikan rostralanteriorcingulate.lh** | 0.81 | 17127 |
| **rfMRI full corr ICA100 edge 995 link 24-30** | 0.81 | 15864 |
| **rfMRI part corr ICA100 edge 424 link 9-29** | 0.81 | 15864 |
| **rfMRI full corr ICA100 edge 94 link 2-42** | 0.81 | 15864 |
| **opening Morphologist SFsup_right** | 0.81 | 18100 |
| **rfMRI part corr ICA25 edge 142 link 9-19** | 0.81 | 15864 |
| **maxdepth Morphologist SLipost_right** | 0.81 | 17978 |
| **opening Morphologist FCLrdiag_right** | 0.81 | 11519 |
| **rfMRI part corr ICA100 edge 1227 link 32-50** | 0.81 | 15864 |
| **rfMRI part corr ICA100 edge 384 link 8-35** | 0.81 | 15864 |
| **rfMRI part corr ICA100 edge 949 link 22-47** | 0.81 | 15864 |

| | | |
|---|---|---|
| **opening Morphologist FCLrasc_left** | 0.82 | 17784 |
| **Left-Inf-Lat-Vent** | 0.82 | 17127 |
| **surface Morphologist SFinf_right** | 0.82 | 18080 |
| **rfMRI full corr ICA100 edge 219 link 5-14** | 0.82 | 15864 |
| **rfMRI full corr ICA100 edge 314 link 7-12** | 0.82 | 15862 |
| **rfMRI full corr ICA100 edge 1292 link 35-52** | 0.82 | 15864 |
| **FA Superior_longitudinal_fasciculus_L** | 0.82 | 16541 |
| **rfMRI part corr ICA100 edge 494 link 10-54** | 0.82 | 15864 |
| **meandepth Morphologist SOTlatpost_left** | 0.82 | 18061 |
| **rfMRI part corr ICA100 edge 989 link 23-55** | 0.82 | 15864 |
| **rfMRI part corr ICA100 edge 1266 link 34-46** | 0.82 | 15864 |
| **rfMRI part corr ICA100 edge 387 link 8-38** | 0.82 | 15864 |
| **surface Morphologist SCsylvian_left** | 0.82 | 13814 |
| **rfMRI full corr ICA100 edge 633 link 14-23** | 0.82 | 15864 |
| **rfMRI part corr ICA100 edge 219 link 5-14** | 0.82 | 15864 |
| **rfMRI full corr ICA100 edge 993 link 24-28** | 0.82 | 15864 |
| **GM_thickness Morphologist SFint_left** | 0.82 | 18092 |
| **rfMRI full corr ICA25 edge 17 link 1-18** | 0.82 | 15864 |
| **rfMRI part corr ICA100 edge 203 link 4-48** | 0.82 | 15864 |
| **rfMRI part corr ICA100 edge 1034 link 25-39** | 0.82 | 15864 |
| **rfMRI full corr ICA100 edge 613 link 13-44** | 0.82 | 15861 |
| **rfMRI part corr ICA100 edge 330 link 7-28** | 0.82 | 15864 |
| **rfMRI full corr ICA100 edge 260 link 5-55** | 0.82 | 15861 |
| **rfMRI part corr ICA100 edge 1032 link 25-37** | 0.82 | 15864 |
| **FA Retrolenticular_part_of_internal_capsule_R** | 0.82 | 16541 |
| **GM_thickness Morphologist SFmarginal_left** | 0.82 | 18035 |
| **rfMRI full corr ICA100 edge 823 link 19-23** | 0.82 | 15864 |
| **rfMRI part corr ICA25 edge 134 link 9-11** | 0.82 | 15864 |
| **rfMRI part corr ICA100 edge 1312 link 36-53** | 0.82 | 15864 |
| **rfMRI full corr ICA100 edge 619 link 13-50** | 0.82 | 15864 |
| **rfMRI full corr ICA100 edge 362 link 8-13** | 0.82 | 15858 |
| **rfMRI part corr ICA25 edge 167 link 12-14** | 0.82 | 15864 |
| **rfMRI full corr ICA100 edge 601 link 13-32** | 0.82 | 15864 |
| **rfMRI part corr ICA100 edge 735 link 16-46** | 0.82 | 15864 |

| | | |
|---|---|---|
| **rfMRI full corr ICA100 edge 851 link 19-51** | 0.82 | 15863 |
| **ThickAvg Desikan lateraloccipital.lh** | 0.82 | 17127 |
| **rfMRI full corr ICA25 edge 4 link 1-5** | 0.82 | 15864 |
| **rfMRI full corr ICA100 edge 380 link 8-31** | 0.82 | 15864 |
| **rfMRI full corr ICA100 edge 1485 link 54-55** | 0.82 | 15864 |
| **rfMRI full corr ICA100 edge 838 link 19-38** | 0.82 | 15864 |
| **rfMRI part corr ICA100 edge 651 link 14-41** | 0.82 | 15864 |
| **MO Superior_cerebellar_peduncle_L** | 0.83 | 16541 |
| **rfMRI part corr ICA25 edge 9 link 1-10** | 0.83 | 15864 |
| **surface Morphologist SsP_right** | 0.83 | 18094 |
| **hull_junction_length Morphologist SPeCinter_left** | 0.83 | 18028 |
| **rfMRI full corr ICA100 edge 288 link 6-34** | 0.83 | 15864 |
| **rfMRI full corr ICA100 edge 839 link 19-39** | 0.83 | 15864 |
| **rfMRI part corr ICA25 edge 22 link 2-4** | 0.83 | 15864 |
| **ICVF Sagittal_stratum-inf_longitidinal_fasci_and_i** | 0.83 | 16540 |
| **SurfArea Destrieux S_suborbital.lh** | 0.83 | 17126 |
| **rfMRI part corr ICA25 edge 113 link 7-15** | 0.83 | 15864 |
| **rfMRI part corr ICA100 edge 1416 link 43-52** | 0.83 | 15864 |
| **SurfArea Desikan medialorbitofrontal.lh** | 0.83 | 17127 |
| **rfMRI full corr ICA100 edge 985 link 23-51** | 0.83 | 15864 |
| **rfMRI full corr ICA100 edge 617 link 13-48** | 0.83 | 15864 |
| **rfMRI full corr ICA100 edge 961 link 23-27** | 0.83 | 15864 |
| **rfMRI full corr ICA100 edge 1383 link 41-44** | 0.83 | 15864 |
| **ICVF Body_of_corpus_callosum** | 0.83 | 16540 |
| **rfMRI full corr ICA100 edge 705 link 15-55** | 0.83 | 15864 |
| **rfMRI full corr ICA100 edge 960 link 23-26** | 0.83 | 15864 |
| **rfMRI full corr ICA100 edge 308 link 6-54** | 0.83 | 15864 |
| **ThickAvg Desikan lateralorbitofrontal.lh** | 0.83 | 17127 |
| **rfMRI part corr ICA100 edge 235 link 5-30** | 0.83 | 15864 |
| **rfMRI part corr ICA100 edge 545 link 12-18** | 0.83 | 15864 |
| **rfMRI full corr ICA25 edge 186 link 14-18** | 0.83 | 15864 |
| **rfMRI full corr ICA100 edge 786 link 18-22** | 0.83 | 15864 |
| **rfMRI full corr ICA100 edge 715 link 16-26** | 0.83 | 15863 |
| **GM_thickness Morphologist SCsylvian_left** | 0.83 | 13807 |

| | | |
|---|---|---|
| **meandepth Morphologist SCu_left** | 0.83 | 18057 |
| **meandepth Morphologist SOlf_right** | 0.83 | 18091 |
| **rfMRI full corr ICA100 edge 49 link 1-50** | 0.83 | 15862 |
| **rfMRI full corr ICA100 edge 476 link 10-36** | 0.83 | 15864 |
| **rfMRI part corr ICA100 edge 479 link 10-39** | 0.83 | 15864 |
| **hull_junction_length Morphologist SCall_right** | 0.83 | 18086 |
| **rfMRI full corr ICA100 edge 912 link 21-43** | 0.83 | 15863 |
| **rfMRI part corr ICA100 edge 190 link 4-35** | 0.83 | 15864 |
| **ICVF Medial_lemniscus_L** | 0.83 | 16540 |
| **rfMRI amplitude ICA100 component 13** | 0.83 | 15863 |
| **FA Fornix-column_and_body_of_fornix** | 0.84 | 16541 |
| **meandepth Morphologist SOTlatmed_left** | 0.84 | 17419 |
| **rfMRI full corr ICA100 edge 159 link 3-55** | 0.84 | 15864 |
| **rfMRI part corr ICA100 edge 717 link 16-28** | 0.84 | 15864 |
| **rfMRI part corr ICA100 edge 988 link 23-54** | 0.84 | 15864 |
| **rfMRI part corr ICA25 edge 38 link 2-20** | 0.84 | 15864 |
| **GrayVol Desikan superiortemporal.rh** | 0.84 | 17127 |
| **GrayVol Desikan temporalpole.lh** | 0.84 | 17127 |
| **FA Posterior_corona_radiata_R** | 0.84 | 16541 |
| **surface Morphologist SOlf_left** | 0.84 | 18094 |
| **rfMRI full corr ICA100 edge 659 link 14-49** | 0.84 | 15864 |
| **rfMRI part corr ICA100 edge 1259 link 34-39** | 0.84 | 15864 |
| **rfMRI part corr ICA25 edge 133 link 9-10** | 0.84 | 15864 |
| **OD Posterior_thalamic_radiation-include_optic_ra** | 0.84 | 16541 |
| **rfMRI full corr ICA100 edge 426 link 9-31** | 0.84 | 15863 |
| **rfMRI part corr ICA100 edge 939 link 22-37** | 0.84 | 15864 |
| **meandepth Morphologist SPasup_right** | 0.84 | 17390 |
| **opening Morphologist FIP_right** | 0.84 | 18097 |
| **rfMRI amplitude ICA100 component 20** | 0.84 | 15863 |
| **rfMRI full corr ICA100 edge 568 link 12-41** | 0.84 | 15864 |
| **rfMRI full corr ICA100 edge 1217 link 32-40** | 0.84 | 15864 |
| **rfMRI part corr ICA100 edge 1008 link 24-43** | 0.84 | 15864 |
| **rfMRI part corr ICA100 edge 908 link 21-39** | 0.84 | 15864 |
| **rfMRI full corr ICA100 edge 316 link 7-14** | 0.84 | 15864 |

| | | |
|---|---|---|
| **rfMRI part corr ICA25 edge 157 link 11-13** | 0.84 | 15864 |
| **maxdepth Morphologist SRinf_right** | 0.84 | 17661 |
| **hull_junction_length Morphologist FCMant_left** | 0.84 | 17879 |
| **SurfArea Destrieux Lat_Fis-ant-Horizont.lh** | 0.84 | 17126 |
| **surface Morphologist SPeCinf_right** | 0.84 | 16608 |
| **rfMRI part corr ICA100 edge 1155 link 29-50** | 0.84 | 15864 |
| **rfMRI full corr ICA100 edge 1248 link 33-49** | 0.84 | 15864 |
| **rfMRI part corr ICA100 edge 942 link 22-40** | 0.84 | 15864 |
| **rfMRI full corr ICA100 edge 482 link 10-42** | 0.84 | 15864 |
| **rfMRI full corr ICA100 edge 1478 link 51-54** | 0.84 | 15864 |
| **rfMRI full corr ICA100 edge 1340 link 38-46** | 0.84 | 15864 |
| **rfMRI part corr ICA100 edge 245 link 5-40** | 0.84 | 15864 |
| **rfMRI full corr ICA100 edge 820 link 19-20** | 0.85 | 15862 |
| **meandepth Morphologist SFmarginal_left** | 0.85 | 18044 |
| **rfMRI part corr ICA100 edge 1050 link 25-55** | 0.85 | 15864 |
| **rfMRI part corr ICA25 edge 103 link 6-19** | 0.85 | 15864 |
| **opening Morphologist SCLPC_right** | 0.85 | 5279 |
| **rfMRI part corr ICA100 edge 1047 link 25-52** | 0.85 | 15864 |
| **rfMRI part corr ICA100 edge 1385 link 41-46** | 0.85 | 15864 |
| **rfMRI part corr ICA100 edge 827 link 19-27** | 0.85 | 15864 |
| **rfMRI part corr ICA100 edge 814 link 18-50** | 0.85 | 15864 |
| **rfMRI part corr ICA100 edge 689 link 15-39** | 0.85 | 15864 |
| **rfMRI full corr ICA100 edge 1341 link 38-47** | 0.85 | 15864 |
| **rfMRI full corr ICA100 edge 848 link 19-48** | 0.85 | 15863 |
| **rfMRI part corr ICA100 edge 296 link 6-42** | 0.85 | 15864 |
| **rfMRI part corr ICA25 edge 140 link 9-17** | 0.85 | 15864 |
| **rfMRI part corr ICA100 edge 134 link 3-30** | 0.85 | 15864 |
| **surface Morphologist FCMant_left** | 0.85 | 17879 |
| **rfMRI part corr ICA100 edge 1347 link 38-53** | 0.85 | 15864 |
| **rfMRI part corr ICA25 edge 207 link 18-21** | 0.85 | 15864 |
| **rfMRI part corr ICA100 edge 997 link 24-32** | 0.85 | 15864 |
| **rfMRI part corr ICA100 edge 536 link 11-52** | 0.85 | 15864 |
| **GM_thickness Morphologist SRinf_right** | 0.85 | 17654 |
| **rfMRI part corr ICA100 edge 452 link 10-12** | 0.85 | 15864 |

| | | |
|---|---|---|
| **rfMRI part corr ICA25 edge 206 link 18-20** | 0.85 | 15864 |
| **rfMRI part corr ICA25 edge 152 link 10-18** | 0.85 | 15864 |
| **surface Morphologist STiant_right** | 0.85 | 18100 |
| **CC_Mid_Posterior** | 0.85 | 17127 |
| **rfMRI part corr ICA25 edge 182 link 13-21** | 0.85 | 15864 |
| **rfMRI full corr ICA100 edge 1422 link 44-47** | 0.85 | 15864 |
| **rfMRI full corr ICA100 edge 1447 link 46-53** | 0.85 | 15864 |
| **rfMRI full corr ICA100 edge 1445 link 46-51** | 0.85 | 15864 |
| **rfMRI part corr ICA100 edge 560 link 12-33** | 0.85 | 15864 |
| **rfMRI part corr ICA100 edge 146 link 3-42** | 0.85 | 15864 |
| **rfMRI part corr ICA100 edge 1483 link 53-54** | 0.85 | 15864 |
| **rfMRI part corr ICA25 edge 24 link 2-6** | 0.85 | 15864 |
| **rfMRI full corr ICA100 edge 369 link 8-20** | 0.85 | 15864 |
| **ThickAvg Desikan frontalpole.lh** | 0.85 | 17127 |
| **rfMRI full corr ICA100 edge 139 link 3-35** | 0.85 | 15864 |
| **rfMRI part corr ICA100 edge 144 link 3-40** | 0.85 | 15864 |
| **rfMRI part corr ICA25 edge 143 link 9-20** | 0.85 | 15864 |
| **ThickAvg Destrieux G_temp_sup-Plan_polar.rh** | 0.85 | 17126 |
| **rfMRI part corr ICA25 edge 76 link 5-7** | 0.85 | 15864 |
| **rfMRI full corr ICA100 edge 1288 link 35-48** | 0.85 | 15864 |
| **MO Cingulum-cingulate_gyrus-R** | 0.85 | 16541 |
| **ISOVF Superior_corona_radiata_R** | 0.85 | 16527 |
| **opening Morphologist SCsylvian_left** | 0.86 | 13814 |
| **rfMRI full corr ICA100 edge 1392 link 41-53** | 0.86 | 15864 |
| **opening Morphologist OCCIPITAL_left** | 0.86 | 18097 |
| **rfMRI full corr ICA100 edge 1108 link 28-29** | 0.86 | 15864 |
| **Left-Pallidum** | 0.86 | 17127 |
| **rfMRI full corr ICA25 edge 101 link 6-17** | 0.86 | 15864 |
| **rfMRI full corr ICA100 edge 371 link 8-22** | 0.86 | 15864 |
| **rfMRI full corr ICA100 edge 247 link 5-42** | 0.86 | 15864 |
| **rfMRI full corr ICA100 edge 105 link 2-53** | 0.86 | 15864 |
| **ThickAvg Destrieux S_precentral-inf-part.rh** | 0.86 | 17127 |
| **rfMRI part corr ICA100 edge 1321 link 37-44** | 0.86 | 15864 |
| **rfMRI part corr ICA100 edge 1400 link 42-48** | 0.86 | 15864 |

| | | |
|---|---|---|
| **rfMRI part corr ICA100 edge 363 link 8-14** | 0.86 | 15864 |
| **surface Morphologist FCMant_right** | 0.86 | 17828 |
| **rfMRI part corr ICA100 edge 407 link 9-12** | 0.86 | 15864 |
| **rfMRI part corr ICA100 edge 1263 link 34-43** | 0.86 | 15864 |
| **rfMRI part corr ICA100 edge 1135 link 29-30** | 0.86 | 15864 |
| **GM_thickness Morphologist SFinf_left** | 0.86 | 18092 |
| **rfMRI part corr ICA100 edge 960 link 23-26** | 0.86 | 15864 |
| **rfMRI full corr ICA25 edge 115 link 7-17** | 0.86 | 15864 |
| **rfMRI full corr ICA100 edge 682 link 15-32** | 0.86 | 15864 |
| **MO Posterior_corona_radiata_R** | 0.86 | 16541 |
| **rfMRI part corr ICA100 edge 623 link 13-54** | 0.86 | 15864 |
| **rfMRI part corr ICA100 edge 123 link 3-19** | 0.86 | 15864 |
| **GM_thickness Morphologist FIPrint1_left** | 0.86 | 17273 |
| **meandepth Morphologist SPeCinter_left** | 0.86 | 18026 |
| **rfMRI full corr ICA100 edge 855 link 19-55** | 0.86 | 15864 |
| **rfMRI full corr ICA100 edge 1012 link 24-47** | 0.86 | 15864 |
| **rfMRI part corr ICA25 edge 90 link 5-21** | 0.86 | 15864 |
| **rfMRI part corr ICA100 edge 463 link 10-23** | 0.86 | 15864 |
| **rfMRI part corr ICA100 edge 1299 link 36-40** | 0.86 | 15864 |
| **rfMRI part corr ICA100 edge 1238 link 33-39** | 0.86 | 15864 |
| **rfMRI part corr ICA100 edge 244 link 5-39** | 0.86 | 15864 |
| **surface Morphologist SFmedian_left** | 0.86 | 18078 |
| **GM_thickness Morphologist FColl_right** | 0.86 | 18089 |
| **rfMRI full corr ICA100 edge 1028 link 25-33** | 0.86 | 15864 |
| **rfMRI part corr ICA100 edge 1181 link 30-51** | 0.86 | 15864 |
| **ThickAvg Destrieux G_and_S_subcentral.rh** | 0.87 | 17127 |
| **rfMRI full corr ICA25 edge 28 link 2-10** | 0.87 | 15864 |
| **rfMRI full corr ICA100 edge 417 link 9-22** | 0.87 | 15864 |
| **rfMRI part corr ICA100 edge 197 link 4-42** | 0.87 | 15864 |
| **rfMRI full corr ICA100 edge 1209 link 31-55** | 0.87 | 15864 |
| **rfMRI full corr ICA100 edge 420 link 9-25** | 0.87 | 15864 |
| **rfMRI part corr ICA100 edge 433 link 9-38** | 0.87 | 15864 |
| **rfMRI part corr ICA100 edge 1045 link 25-50** | 0.87 | 15864 |
| **rfMRI full corr ICA100 edge 170 link 4-15** | 0.87 | 15861 |

| | | |
|---|---|---|
| **rfMRI full corr ICA100 edge 1415 link 43-51** | 0.87 | 15864 |
| **rfMRI part corr ICA100 edge 1425 link 44-50** | 0.87 | 15864 |
| **rfMRI part corr ICA100 edge 73 link 2-21** | 0.87 | 15864 |
| **rfMRI part corr ICA100 edge 1458 link 48-49** | 0.87 | 15864 |
| **rfMRI part corr ICA100 edge 906 link 21-37** | 0.87 | 15864 |
| **rfMRI full corr ICA100 edge 60 link 2-8** | 0.87 | 15864 |
| **rfMRI full corr ICA100 edge 726 link 16-37** | 0.87 | 15864 |
| **rfMRI part corr ICA100 edge 1291 link 35-51** | 0.87 | 15864 |
| **rfMRI full corr ICA100 edge 1254 link 33-55** | 0.87 | 15864 |
| **rfMRI full corr ICA25 edge 38 link 2-20** | 0.87 | 15864 |
| **GM_thickness Morphologist FCMant_left** | 0.87 | 17870 |
| **rfMRI part corr ICA25 edge 149 link 10-15** | 0.87 | 15864 |
| **GM_thickness Morphologist SForbitaire_right** | 0.87 | 17271 |
| **rfMRI full corr ICA100 edge 1218 link 32-41** | 0.87 | 15864 |
| **surface Morphologist SPoCsup_left** | 0.87 | 17976 |
| **rfMRI part corr ICA100 edge 792 link 18-28** | 0.87 | 15864 |
| **rfMRI full corr ICA100 edge 1295 link 35-55** | 0.87 | 15864 |
| **rfMRI full corr ICA100 edge 119 link 3-15** | 0.87 | 15864 |
| **rfMRI part corr ICA100 edge 612 link 13-43** | 0.87 | 15864 |
| **FA Anterior_corona_radiata_R** | 0.87 | 16541 |
| **rfMRI part corr ICA100 edge 1396 link 42-44** | 0.87 | 15864 |
| **rfMRI part corr ICA25 edge 198 link 16-19** | 0.87 | 15864 |
| **rfMRI part corr ICA100 edge 1058 link 26-34** | 0.87 | 15864 |
| **rfMRI part corr ICA100 edge 1405 link 42-53** | 0.87 | 15864 |
| **meandepth Morphologist FCLrant_left** | 0.87 | 17561 |
| **rfMRI part corr ICA25 edge 4 link 1-5** | 0.87 | 15864 |
| **rfMRI part corr ICA100 edge 1169 link 30-39** | 0.88 | 15864 |
| **rfMRI part corr ICA100 edge 83 link 2-31** | 0.88 | 15864 |
| **ThickAvg Destrieux S_pericallosal.rh** | 0.88 | 17127 |
| **rfMRI part corr ICA100 edge 112 link 3-8** | 0.88 | 15864 |
| **GrayVol Destrieux S_front_middle.rh** | 0.88 | 17127 |
| **rfMRI part corr ICA100 edge 696 link 15-46** | 0.88 | 15864 |
| **rfMRI part corr ICA100 edge 916 link 21-47** | 0.88 | 15864 |
| **rfMRI full corr ICA100 edge 1393 link 41-54** | 0.88 | 15864 |

| | | |
|---|---|---|
| rfMRI full corr ICA100 edge 1203 link 31-49 | 0.88 | 15864 |
| rfMRI full corr ICA100 edge 441 link 9-46 | 0.88 | 15864 |
| opening Morphologist FCLrscpost_left | 0.88 | 13973 |
| rfMRI full corr ICA100 edge 1280 link 35-40 | 0.88 | 15864 |
| rfMRI part corr ICA100 edge 76 link 2-24 | 0.88 | 15864 |
| rfMRI full corr ICA100 edge 970 link 23-36 | 0.88 | 15863 |
| SurfArea Destrieux S_circular_insula_sup.lh | 0.88 | 17126 |
| rfMRI part corr ICA100 edge 34 link 1-35 | 0.88 | 15864 |
| rfMRI part corr ICA100 edge 349 link 7-47 | 0.88 | 15864 |
| rfMRI part corr ICA100 edge 748 link 17-21 | 0.88 | 15864 |
| rfMRI part corr ICA100 edge 339 link 7-37 | 0.88 | 15864 |
| rfMRI part corr ICA100 edge 739 link 16-50 | 0.88 | 15864 |
| rfMRI part corr ICA100 edge 985 link 23-51 | 0.88 | 15864 |
| GrayVol Destrieux Lat_Fis-ant-Horizont.rh | 0.88 | 17125 |
| rfMRI full corr ICA100 edge 712 link 16-23 | 0.88 | 15860 |
| rfMRI full corr ICA25 edge 53 link 3-17 | 0.88 | 15864 |
| GM_thickness Morphologist STsterascpost_right | 0.88 | 18041 |
| rfMRI part corr ICA100 edge 1059 link 26-35 | 0.88 | 15864 |
| maxdepth Morphologist SPoCsup_right | 0.88 | 17842 |
| hull_junction_length Morphologist STsterascpost_ | 0.88 | 18048 |
| rfMRI full corr ICA100 edge 734 link 16-45 | 0.88 | 15864 |
| OD Splenium_of_corpus_callosum | 0.88 | 16541 |
| rfMRI full corr ICA100 edge 300 link 6-46 | 0.88 | 15864 |
| rfMRI full corr ICA100 edge 339 link 7-37 | 0.88 | 15864 |
| rfMRI part corr ICA100 edge 1439 link 45-54 | 0.88 | 15864 |
| rfMRI full corr ICA100 edge 232 link 5-27 | 0.88 | 15864 |
| rfMRI part corr ICA100 edge 194 link 4-39 | 0.88 | 15864 |
| GrayVol Destrieux G_rectus.lh | 0.88 | 17127 |
| rfMRI full corr ICA100 edge 743 link 16-54 | 0.88 | 15861 |
| rfMRI full corr ICA100 edge 794 link 18-30 | 0.88 | 15863 |
| rfMRI part corr ICA100 edge 114 link 3-10 | 0.88 | 15864 |
| meandepth Morphologist FCalant-ScCal_right | 0.88 | 18100 |
| rfMRI full corr ICA100 edge 760 link 17-33 | 0.88 | 15862 |
| FA Superior_fronto-occipital_fasciculus-part_of_a | 0.88 | 16541 |

| | | |
|---|---|---|
| **rfMRI full corr ICA100 edge 116 link 3-12** | 0.88 | 15864 |
| **rfMRI full corr ICA100 edge 1200 link 31-46** | 0.88 | 15864 |
| **surface Morphologist FIPrint2_left** | 0.88 | 16244 |
| **rfMRI amplitude ICA100 component 44** | 0.88 | 15863 |
| **rfMRI part corr ICA100 edge 670 link 15-20** | 0.88 | 15864 |
| **rfMRI part corr ICA100 edge 1302 link 36-43** | 0.88 | 15864 |
| **rfMRI full corr ICA100 edge 1342 link 38-48** | 0.88 | 15864 |
| **MD Fornix-column_and_body_of_fornix** | 0.88 | 16541 |
| **rfMRI full corr ICA100 edge 19 link 1-20** | 0.88 | 15864 |
| **rfMRI part corr ICA100 edge 168 link 4-13** | 0.88 | 15864 |
| **surface Morphologist SFinf_left** | 0.88 | 18101 |
| **rfMRI part corr ICA100 edge 281 link 6-27** | 0.88 | 15864 |
| **rfMRI full corr ICA100 edge 689 link 15-39** | 0.89 | 15864 |
| **SurfArea Destrieux S_oc_middle_and_Lunatus.lh** | 0.89 | 17126 |
| **rfMRI part corr ICA100 edge 97 link 2-45** | 0.89 | 15864 |
| **ICVF Sagittal_stratum-inf_longitidinal_fasci_and_i** | 0.89 | 16540 |
| **maxdepth Morphologist STipost_left** | 0.89 | 18096 |
| **rfMRI full corr ICA100 edge 394 link 8-45** | 0.89 | 15864 |
| **rfMRI full corr ICA100 edge 1347 link 38-53** | 0.89 | 15864 |
| **rfMRI part corr ICA25 edge 47 link 3-11** | 0.89 | 15864 |
| **rfMRI full corr ICA25 edge 97 link 6-13** | 0.89 | 15864 |
| **maxdepth Morphologist SFmarginal_right** | 0.89 | 18006 |
| **MD Posterior_corona_radiata_R** | 0.89 | 16541 |
| **MO Superior_fronto-occipital_fasciculus-part_of_** | 0.89 | 16541 |
| **SurfArea Destrieux S_collat_transv_ant.rh** | 0.89 | 17125 |
| **rfMRI part corr ICA100 edge 1006 link 24-41** | 0.89 | 15864 |
| **surface Morphologist STsterascant_left** | 0.89 | 17560 |
| **rfMRI full corr ICA100 edge 1269 link 34-49** | 0.89 | 15864 |
| **maxdepth Morphologist SFsup_right** | 0.89 | 18100 |
| **opening Morphologist SOlf_left** | 0.89 | 18094 |
| **rfMRI full corr ICA25 edge 74 link 4-21** | 0.89 | 15864 |
| **rfMRI part corr ICA100 edge 799 link 18-35** | 0.89 | 15864 |
| **rfMRI part corr ICA100 edge 794 link 18-30** | 0.89 | 15864 |
| **meandepth Morphologist SOlf_left** | 0.89 | 18094 |

| | | |
|---|---|---|
| **ISOVF Anterior_limb_of_internal_capsule_R** | 0.89 | 16527 |
| **rfMRI part corr ICA100 edge 506 link 11-22** | 0.89 | 15864 |
| **hull_junction_length Morphologist STipost_right** | 0.89 | 18093 |
| **rfMRI part corr ICA100 edge 251 link 5-46** | 0.89 | 15864 |
| **hull_junction_length Morphologist SC_right** | 0.89 | 18100 |
| **rfMRI part corr ICA100 edge 131 link 3-27** | 0.89 | 15864 |
| **SurfArea Destrieux G_and_S_cingul-Ant.lh** | 0.89 | 17127 |
| **rfMRI part corr ICA25 edge 20 link 1-21** | 0.89 | 15864 |
| **rfMRI full corr ICA100 edge 802 link 18-38** | 0.89 | 15864 |
| **rfMRI part corr ICA100 edge 962 link 23-28** | 0.89 | 15864 |
| **rfMRI full corr ICA100 edge 1221 link 32-44** | 0.89 | 15864 |
| **rfMRI full corr ICA100 edge 513 link 11-29** | 0.89 | 15864 |
| **rfMRI part corr ICA100 edge 755 link 17-28** | 0.89 | 15864 |
| **rfMRI part corr ICA25 edge 53 link 3-17** | 0.9 | 15864 |
| **meandepth Morphologist INSULA_right** | 0.9 | 18100 |
| **surface Morphologist SOr_left** | 0.9 | 18100 |
| **rfMRI full corr ICA25 edge 168 link 12-15** | 0.9 | 15864 |
| **meandepth Morphologist SPeCinter_right** | 0.9 | 18083 |
| **rfMRI part corr ICA25 edge 178 link 13-17** | 0.9 | 15864 |
| **rfMRI part corr ICA100 edge 333 link 7-31** | 0.9 | 15864 |
| **rfMRI part corr ICA100 edge 1245 link 33-46** | 0.9 | 15864 |
| **rfMRI full corr ICA100 edge 566 link 12-39** | 0.9 | 15864 |
| **rfMRI part corr ICA100 edge 510 link 11-26** | 0.9 | 15864 |
| **rfMRI full corr ICA100 edge 1043 link 25-48** | 0.9 | 15864 |
| **rfMRI part corr ICA100 edge 1031 link 25-36** | 0.9 | 15864 |
| **rfMRI part corr ICA100 edge 1191 link 31-37** | 0.9 | 15864 |
| **rfMRI part corr ICA100 edge 630 link 14-20** | 0.9 | 15864 |
| **rfMRI full corr ICA25 edge 142 link 9-19** | 0.9 | 15864 |
| **rfMRI part corr ICA100 edge 69 link 2-17** | 0.9 | 15864 |
| **rfMRI full corr ICA100 edge 988 link 23-54** | 0.9 | 15864 |
| **rfMRI amplitude ICA25 component 21** | 0.9 | 15863 |
| **rfMRI full corr ICA100 edge 669 link 15-19** | 0.9 | 15864 |
| **GM_thickness Morphologist FCLrdiag_left** | 0.9 | 8547 |
| **rfMRI full corr ICA100 edge 495 link 10-55** | 0.9 | 15864 |

| | | |
|---|---|---|
| **rfMRI full corr ICA100 edge 740 link 16-51** | 0.9 | 15864 |
| **rfMRI part corr ICA100 edge 938 link 22-36** | 0.9 | 15864 |
| **rfMRI full corr ICA100 edge 1153 link 29-48** | 0.9 | 15863 |
| **rfMRI part corr ICA100 edge 556 link 12-29** | 0.9 | 15864 |
| **rfMRI part corr ICA100 edge 187 link 4-32** | 0.9 | 15864 |
| **rfMRI part corr ICA100 edge 271 link 6-17** | 0.9 | 15864 |
| **GM_thickness Morphologist FCalant-ScCal_left** | 0.9 | 18091 |
| **rfMRI full corr ICA25 edge 128 link 8-17** | 0.9 | 15864 |
| **rfMRI full corr ICA25 edge 49 link 3-13** | 0.9 | 15864 |
| **surface Morphologist SFsup_right** | 0.9 | 18100 |
| **opening Morphologist INSULA_right** | 0.9 | 18100 |
| **rfMRI full corr ICA100 edge 1247 link 33-48** | 0.9 | 15864 |
| **ThickAvg Destrieux S_orbital_med-olfact.rh** | 0.9 | 17125 |
| **rfMRI part corr ICA100 edge 1204 link 31-50** | 0.9 | 15864 |
| **rfMRI full corr ICA100 edge 311 link 7-9** | 0.9 | 15863 |
| **rfMRI part corr ICA100 edge 1402 link 42-50** | 0.9 | 15864 |
| **rfMRI full corr ICA100 edge 310 link 7-8** | 0.9 | 15864 |
| **rfMRI part corr ICA100 edge 1189 link 31-35** | 0.9 | 15864 |
| **ThickAvg Desikan parsorbitalis.rh** | 0.9 | 17127 |
| **rfMRI full corr ICA100 edge 1044 link 25-49** | 0.9 | 15864 |
| **rfMRI part corr ICA100 edge 664 link 14-54** | 0.9 | 15864 |
| **rfMRI full corr ICA100 edge 242 link 5-37** | 0.9 | 15863 |
| **rfMRI full corr ICA25 edge 45 link 3-9** | 0.9 | 15863 |
| **rfMRI full corr ICA100 edge 733 link 16-44** | 0.9 | 15863 |
| **rfMRI part corr ICA100 edge 292 link 6-38** | 0.9 | 15864 |
| **rfMRI part corr ICA100 edge 1447 link 46-53** | 0.9 | 15864 |
| **rfMRI part corr ICA100 edge 1106 link 27-54** | 0.91 | 15864 |
| **rfMRI full corr ICA100 edge 871 link 20-36** | 0.91 | 15864 |
| **rfMRI full corr ICA100 edge 192 link 4-37** | 0.91 | 15863 |
| **rfMRI full corr ICA100 edge 400 link 8-51** | 0.91 | 15864 |
| **rfMRI part corr ICA25 edge 108 link 7-10** | 0.91 | 15864 |
| **rfMRI full corr ICA100 edge 1352 link 39-42** | 0.91 | 15864 |
| **rfMRI part corr ICA25 edge 35 link 2-17** | 0.91 | 15864 |
| **meandepth Morphologist FCLrretroCtr_right** | 0.91 | 16148 |

| | | |
|---|---|---|
| **opening Morphologist FIPPoCinf_left** | 0.91 | 18079 |
| **rfMRI part corr ICA100 edge 22 link 1-23** | 0.91 | 15864 |
| **rfMRI part corr ICA25 edge 89 link 5-20** | 0.91 | 15864 |
| **rfMRI full corr ICA25 edge 12 link 1-13** | 0.91 | 15864 |
| **rfMRI full corr ICA100 edge 1180 link 30-50** | 0.91 | 15864 |
| **rfMRI part corr ICA100 edge 259 link 5-54** | 0.91 | 15864 |
| **rfMRI full corr ICA100 edge 534 link 11-50** | 0.91 | 15863 |
| **surface Morphologist FCLrscpost_left** | 0.91 | 13973 |
| **rfMRI full corr ICA100 edge 748 link 17-21** | 0.91 | 15864 |
| **ThickAvg Desikan rostralmiddlefrontal.lh** | 0.91 | 17127 |
| **rfMRI full corr ICA100 edge 1148 link 29-43** | 0.91 | 15864 |
| **rfMRI part corr ICA100 edge 325 link 7-23** | 0.91 | 15864 |
| **rfMRI part corr ICA100 edge 1258 link 34-38** | 0.91 | 15864 |
| **rfMRI part corr ICA100 edge 1180 link 30-50** | 0.91 | 15864 |
| **rfMRI part corr ICA100 edge 308 link 6-54** | 0.91 | 15864 |
| **opening Morphologist FCLrscant_right** | 0.91 | 5353 |
| **SurfArea Destrieux S_occipital_ant.lh** | 0.91 | 17127 |
| **ThickAvg Desikan rostralmiddlefrontal.rh** | 0.91 | 17127 |
| **GrayVol Destrieux G_cingul-Post-dorsal.rh** | 0.91 | 17127 |
| **meandepth Morphologist FCLrdiag_right** | 0.91 | 11366 |
| **ThickAvg Destrieux S_front_inf.lh** | 0.91 | 17127 |
| **rfMRI part corr ICA100 edge 1187 link 31-33** | 0.91 | 15864 |
| **GrayVol Desikan caudalanteriorcingulate.rh** | 0.91 | 17127 |
| **rfMRI part corr ICA100 edge 140 link 3-36** | 0.91 | 15864 |
| **SurfArea Desikan caudalanteriorcingulate.rh** | 0.91 | 17127 |
| **rfMRI full corr ICA100 edge 422 link 9-27** | 0.91 | 15863 |
| **rfMRI full corr ICA100 edge 31 link 1-32** | 0.91 | 15864 |
| **rfMRI part corr ICA100 edge 71 link 2-19** | 0.91 | 15864 |
| **rfMRI part corr ICA100 edge 265 link 6-11** | 0.91 | 15864 |
| **rfMRI full corr ICA100 edge 511 link 11-27** | 0.91 | 15864 |
| **rfMRI part corr ICA100 edge 567 link 12-40** | 0.91 | 15864 |
| **maxdepth Morphologist SFinf_right** | 0.91 | 18080 |
| **rfMRI full corr ICA100 edge 1111 link 28-32** | 0.91 | 15864 |
| **rfMRI part corr ICA100 edge 1469 link 49-54** | 0.91 | 15864 |

| | | |
|---|---|---|
| **rfMRI part corr ICA100 edge 1317 link 37-40** | 0.92 | 15864 |
| **rfMRI full corr ICA100 edge 853 link 19-53** | 0.92 | 15864 |
| **rfMRI part corr ICA100 edge 1452 link 47-50** | 0.92 | 15864 |
| **rfMRI part corr ICA100 edge 685 link 15-35** | 0.92 | 15864 |
| **rfMRI part corr ICA100 edge 1072 link 26-48** | 0.92 | 15864 |
| **rfMRI full corr ICA100 edge 1365 link 39-55** | 0.92 | 15864 |
| **rfMRI part corr ICA100 edge 1234 link 33-35** | 0.92 | 15864 |
| **rfMRI amplitude ICA25 component 5** | 0.92 | 15863 |
| **rfMRI full corr ICA100 edge 750 link 17-23** | 0.92 | 15860 |
| **GM_thickness Morphologist SOr_left** | 0.92 | 18089 |
| **rfMRI full corr ICA100 edge 16 link 1-17** | 0.92 | 15864 |
| **rfMRI full corr ICA100 edge 895 link 21-26** | 0.92 | 15864 |
| **rfMRI part corr ICA100 edge 355 link 7-53** | 0.92 | 15864 |
| **rfMRI part corr ICA100 edge 10 link 1-11** | 0.92 | 15864 |
| **rfMRI part corr ICA100 edge 409 link 9-14** | 0.92 | 15864 |
| **ThickAvg Desikan rostralanteriorcingulate.rh** | 0.92 | 17127 |
| **hull_junction_length Morphologist SCall_left** | 0.92 | 18064 |
| **rfMRI part corr ICA100 edge 1096 link 27-44** | 0.92 | 15864 |
| **rfMRI full corr ICA100 edge 1476 link 51-52** | 0.92 | 15864 |
| **rfMRI full corr ICA100 edge 701 link 15-51** | 0.92 | 15864 |
| **rfMRI part corr ICA25 edge 154 link 10-20** | 0.92 | 15864 |
| **rfMRI part corr ICA25 edge 58 link 4-5** | 0.92 | 15864 |
| **ThickAvg Destrieux S_central.lh** | 0.92 | 17127 |
| **FA External_capsule_R** | 0.92 | 16541 |
| **rfMRI part corr ICA25 edge 83 link 5-14** | 0.92 | 15864 |
| **rfMRI full corr ICA100 edge 1241 link 33-42** | 0.92 | 15864 |
| **rfMRI full corr ICA100 edge 86 link 2-34** | 0.92 | 15864 |
| **rfMRI part corr ICA100 edge 503 link 11-19** | 0.92 | 15864 |
| **rfMRI part corr ICA100 edge 1176 link 30-46** | 0.92 | 15864 |
| **rfMRI part corr ICA100 edge 328 link 7-26** | 0.92 | 15864 |
| **rfMRI full corr ICA100 edge 486 link 10-46** | 0.92 | 15864 |
| **rfMRI part corr ICA100 edge 444 link 9-49** | 0.92 | 15864 |
| **rfMRI part corr ICA25 edge 16 link 1-17** | 0.92 | 15864 |
| **rfMRI part corr ICA100 edge 495 link 10-55** | 0.92 | 15864 |

| | | |
|---|---|---|
| **rfMRI full corr ICA100 edge 687 link 15-37** | 0.92 | 15864 |
| **rfMRI part corr ICA100 edge 729 link 16-40** | 0.92 | 15864 |
| **rfMRI part corr ICA100 edge 1120 link 28-41** | 0.92 | 15864 |
| **rfMRI full corr ICA100 edge 672 link 15-22** | 0.92 | 15862 |
| **rfMRI part corr ICA100 edge 1249 link 33-50** | 0.92 | 15864 |
| **rfMRI part corr ICA100 edge 1171 link 30-41** | 0.92 | 15864 |
| **GM_thickness Morphologist SpC_left** | 0.92 | 17034 |
| **rfMRI full corr ICA100 edge 665 link 14-55** | 0.92 | 15864 |
| **rfMRI part corr ICA25 edge 150 link 10-16** | 0.93 | 15864 |
| **SurfArea Desikan parsorbitalis.lh** | 0.93 | 17126 |
| **rfMRI part corr ICA100 edge 1024 link 25-29** | 0.93 | 15864 |
| **rfMRI full corr ICA100 edge 1321 link 37-44** | 0.93 | 15864 |
| **ThickAvg Destrieux G_oc-temp_med-Parahip.lh** | 0.93 | 17127 |
| **rfMRI full corr ICA100 edge 888 link 20-53** | 0.93 | 15864 |
| **GM_thickness Morphologist SFinfant_right** | 0.93 | 18035 |
| **rfMRI part corr ICA100 edge 1079 link 26-55** | 0.93 | 15864 |
| **rfMRI part corr ICA100 edge 500 link 11-16** | 0.93 | 15864 |
| **rfMRI amplitude ICA100 component 37** | 0.93 | 15863 |
| **rfMRI part corr ICA100 edge 1207 link 31-53** | 0.93 | 15864 |
| **rfMRI full corr ICA100 edge 627 link 14-17** | 0.93 | 15864 |
| **rfMRI full corr ICA100 edge 22 link 1-23** | 0.93 | 15864 |
| **rfMRI part corr ICA100 edge 25 link 1-26** | 0.93 | 15864 |
| **rfMRI part corr ICA100 edge 1070 link 26-46** | 0.93 | 15864 |
| **4th-Ventricle** | 0.93 | 17127 |
| **rfMRI full corr ICA25 edge 195 link 15-21** | 0.93 | 15864 |
| **rfMRI full corr ICA100 edge 196 link 4-41** | 0.93 | 15864 |
| **rfMRI part corr ICA100 edge 1468 link 49-53** | 0.93 | 15864 |
| **rfMRI part corr ICA100 edge 1344 link 38-50** | 0.93 | 15864 |
| **hull_junction_length Morphologist SPeCmedian_ri** | 0.93 | 17074 |
| **rfMRI part corr ICA100 edge 128 link 3-24** | 0.93 | 15864 |
| **rfMRI part corr ICA100 edge 54 link 1-55** | 0.93 | 15864 |
| **rfMRI full corr ICA100 edge 244 link 5-39** | 0.93 | 15864 |
| **rfMRI full corr ICA100 edge 958 link 23-24** | 0.93 | 15864 |
| **maxdepth Morphologist SPeCsup_left** | 0.93 | 17143 |

| | | |
|---|---|---|
| **rfMRI full corr ICA100 edge 1348 link 38-54** | 0.93 | 15864 |
| **ThickAvg Destrieux S_collat_transv_ant.lh** | 0.93 | 17127 |
| **ThickAvg Destrieux G_and_S_cingul-Ant.lh** | 0.93 | 17127 |
| **rfMRI full corr ICA100 edge 909 link 21-40** | 0.93 | 15864 |
| **rfMRI part corr ICA100 edge 243 link 5-38** | 0.93 | 15864 |
| **rfMRI full corr ICA100 edge 1293 link 35-53** | 0.93 | 15864 |
| **rfMRI part corr ICA100 edge 62 link 2-10** | 0.93 | 15864 |
| **rfMRI part corr ICA100 edge 1069 link 26-45** | 0.93 | 15864 |
| **surface Morphologist FCLrscant_right** | 0.93 | 5353 |
| **rfMRI part corr ICA100 edge 107 link 2-55** | 0.93 | 15864 |
| **MD Middle_cerebellar_peduncle** | 0.93 | 16541 |
| **rfMRI part corr ICA100 edge 1197 link 31-43** | 0.93 | 15864 |
| **rfMRI full corr ICA100 edge 480 link 10-40** | 0.93 | 15863 |
| **rfMRI part corr ICA100 edge 400 link 8-51** | 0.93 | 15864 |
| **rfMRI full corr ICA100 edge 145 link 3-41** | 0.93 | 15859 |
| **rfMRI part corr ICA100 edge 1478 link 51-54** | 0.93 | 15864 |
| **rfMRI part corr ICA100 edge 307 link 6-53** | 0.93 | 15864 |
| **rfMRI full corr ICA100 edge 1423 link 44-48** | 0.93 | 15864 |
| **FA Superior_fronto-occipital_fasciculus-part_of_a** | 0.93 | 16541 |
| **rfMRI full corr ICA100 edge 934 link 22-32** | 0.93 | 15864 |
| **rfMRI part corr ICA100 edge 326 link 7-24** | 0.94 | 15864 |
| **rfMRI part corr ICA100 edge 1443 link 46-49** | 0.94 | 15864 |
| **rfMRI full corr ICA100 edge 1327 link 37-50** | 0.94 | 15864 |
| **opening Morphologist FCLa_right** | 0.94 | 18097 |
| **rfMRI full corr ICA25 edge 26 link 2-8** | 0.94 | 15863 |
| **GrayVol Desikan medialorbitofrontal.rh** | 0.94 | 17126 |
| **rfMRI part corr ICA100 edge 1214 link 32-37** | 0.94 | 15864 |
| **MO Anterior_corona_radiata_R** | 0.94 | 16541 |
| **opening Morphologist FCLrscant_left** | 0.94 | 4323 |
| **rfMRI full corr ICA100 edge 341 link 7-39** | 0.94 | 15864 |
| **rfMRI full corr ICA25 edge 24 link 2-6** | 0.94 | 15864 |
| **rfMRI part corr ICA100 edge 1365 link 39-55** | 0.94 | 15864 |
| **rfMRI part corr ICA100 edge 145 link 3-41** | 0.94 | 15864 |
| **rfMRI full corr ICA100 edge 1414 link 43-50** | 0.94 | 15864 |

| | | |
|---|---|---|
| **rfMRI part corr ICA100 edge 1324 link 37-47** | 0.94 | 15864 |
| **rfMRI part corr ICA100 edge 998 link 24-33** | 0.94 | 15864 |
| **maxdepth Morphologist FIPrint1_left** | 0.94 | 17282 |
| **rfMRI full corr ICA100 edge 1245 link 33-46** | 0.94 | 15864 |
| **rfMRI part corr ICA100 edge 191 link 4-36** | 0.94 | 15864 |
| **GrayVol Desikan transversetemporal.rh** | 0.94 | 17125 |
| **rfMRI part corr ICA100 edge 983 link 23-49** | 0.94 | 15864 |
| **rfMRI part corr ICA100 edge 250 link 5-45** | 0.94 | 15864 |
| **ISOVF Anterior_corona_radiata_L** | 0.94 | 16527 |
| **rfMRI part corr ICA25 edge 3 link 1-4** | 0.94 | 15864 |
| **rfMRI full corr ICA25 edge 93 link 6-9** | 0.94 | 15864 |
| **rfMRI part corr ICA100 edge 93 link 2-41** | 0.94 | 15864 |
| **rfMRI part corr ICA100 edge 1203 link 31-49** | 0.94 | 15864 |
| **surface Morphologist FPO_left** | 0.94 | 18097 |
| **rfMRI full corr ICA100 edge 784 link 18-20** | 0.94 | 15864 |
| **rfMRI full corr ICA100 edge 487 link 10-47** | 0.94 | 15864 |
| **rfMRI part corr ICA100 edge 652 link 14-42** | 0.94 | 15864 |
| **rfMRI full corr ICA100 edge 711 link 16-22** | 0.94 | 15861 |
| **ThickAvg Desikan transversetemporal.rh** | 0.94 | 17125 |
| **rfMRI part corr ICA100 edge 1423 link 44-48** | 0.94 | 15864 |
| **ICVF Cingulum-cingulate_gyrus-L** | 0.94 | 16540 |
| **GM_thickness Morphologist SLipost_right** | 0.94 | 17969 |
| **rfMRI full corr ICA100 edge 125 link 3-21** | 0.94 | 15864 |
| **rfMRI part corr ICA25 edge 6 link 1-7** | 0.94 | 15864 |
| **rfMRI part corr ICA100 edge 342 link 7-40** | 0.94 | 15864 |
| **rfMRI part corr ICA100 edge 955 link 22-53** | 0.94 | 15864 |
| **ThickAvg Destrieux G_front_inf-Triangul.lh** | 0.95 | 17127 |
| **rfMRI full corr ICA100 edge 175 link 4-20** | 0.95 | 15863 |
| **rfMRI full corr ICA100 edge 1329 link 37-52** | 0.95 | 15864 |
| **meandepth Morphologist STpol_right** | 0.95 | 18062 |
| **rfMRI part corr ICA100 edge 1315 link 37-38** | 0.95 | 15864 |
| **rfMRI part corr ICA100 edge 1373 link 40-48** | 0.95 | 15864 |
| **rfMRI part corr ICA100 edge 1338 link 38-44** | 0.95 | 15864 |
| **rfMRI part corr ICA100 edge 1111 link 28-32** | 0.95 | 15864 |

| | | |
|---|---|---|
| **rfMRI full corr ICA100 edge 1379 link 40-54** | 0.95 | 15864 |
| **FA Cerebral_peduncle_L** | 0.95 | 16541 |
| **rfMRI part corr ICA100 edge 681 link 15-31** | 0.95 | 15864 |
| **rfMRI part corr ICA100 edge 1138 link 29-33** | 0.95 | 15864 |
| **rfMRI part corr ICA25 edge 57 link 3-21** | 0.95 | 15864 |
| **rfMRI part corr ICA100 edge 167 link 4-12** | 0.95 | 15864 |
| **SurfArea Desikan lateralorbitofrontal.rh** | 0.95 | 17127 |
| **rfMRI part corr ICA100 edge 568 link 12-41** | 0.95 | 15864 |
| **rfMRI full corr ICA100 edge 323 link 7-21** | 0.95 | 15864 |
| **rfMRI part corr ICA100 edge 713 link 16-24** | 0.95 | 15864 |
| **rfMRI full corr ICA100 edge 742 link 16-53** | 0.95 | 15863 |
| **rfMRI part corr ICA100 edge 1334 link 38-40** | 0.95 | 15864 |
| **rfMRI amplitude ICA100 component 12** | 0.95 | 15863 |
| **rfMRI part corr ICA100 edge 633 link 14-23** | 0.95 | 15864 |
| **maxdepth Morphologist FCMant_left** | 0.95 | 17879 |
| **rfMRI part corr ICA100 edge 261 link 6-7** | 0.95 | 15864 |
| **rfMRI full corr ICA100 edge 1171 link 30-41** | 0.95 | 15864 |
| **meandepth Morphologist SRinf_right** | 0.95 | 17661 |
| **rfMRI part corr ICA25 edge 128 link 8-17** | 0.95 | 15864 |
| **rfMRI part corr ICA100 edge 1110 link 28-31** | 0.95 | 15864 |
| **rfMRI full corr ICA100 edge 521 link 11-37** | 0.95 | 15864 |
| **rfMRI part corr ICA25 edge 10 link 1-11** | 0.95 | 15864 |
| **rfMRI full corr ICA25 edge 46 link 3-10** | 0.95 | 15864 |
| **rfMRI full corr ICA100 edge 1322 link 37-45** | 0.95 | 15864 |
| **rfMRI part corr ICA100 edge 70 link 2-18** | 0.95 | 15864 |
| **rfMRI part corr ICA100 edge 745 link 17-18** | 0.95 | 15864 |
| **rfMRI full corr ICA100 edge 309 link 6-55** | 0.95 | 15861 |
| **rfMRI full corr ICA100 edge 542 link 12-15** | 0.95 | 15864 |
| **rfMRI full corr ICA100 edge 208 link 4-53** | 0.95 | 15864 |
| **rfMRI full corr ICA100 edge 493 link 10-53** | 0.95 | 15864 |
| **rfMRI full corr ICA100 edge 446 link 9-51** | 0.95 | 15864 |
| **GrayVol Destrieux S_suborbital.lh** | 0.95 | 17126 |
| **rfMRI full corr ICA100 edge 354 link 7-52** | 0.95 | 15864 |
| **rfMRI part corr ICA100 edge 1036 link 25-41** | 0.96 | 15864 |

| | | |
|---|---|---|
| **rfMRI part corr ICA100 edge 413 link 9-18** | 0.96 | 15864 |
| **GM_thickness Morphologist SFinter_left** | 0.96 | 18092 |
| **rfMRI amplitude ICA100 component 28** | 0.96 | 15863 |
| **GM_thickness Morphologist FIPrint2_right** | 0.96 | 13056 |
| **rfMRI full corr ICA100 edge 1274 link 34-54** | 0.96 | 15864 |
| **rfMRI full corr ICA100 edge 430 link 9-35** | 0.96 | 15862 |
| **rfMRI part corr ICA100 edge 121 link 3-17** | 0.96 | 15864 |
| **rfMRI full corr ICA25 edge 85 link 5-16** | 0.96 | 15864 |
| **rfMRI part corr ICA100 edge 980 link 23-46** | 0.96 | 15864 |
| **SurfArea Destrieux Pole_occipital.lh** | 0.96 | 17127 |
| **meandepth Morphologist SFinf_right** | 0.96 | 18078 |
| **rfMRI part corr ICA100 edge 825 link 19-25** | 0.96 | 15864 |
| **rfMRI part corr ICA100 edge 677 link 15-27** | 0.96 | 15864 |
| **ISOVF Uncinate_fasciculus_R** | 0.96 | 16527 |
| **rfMRI full corr ICA100 edge 875 link 20-40** | 0.96 | 15864 |
| **rfMRI part corr ICA100 edge 11 link 1-12** | 0.96 | 15864 |
| **rfMRI part corr ICA100 edge 15 link 1-16** | 0.96 | 15864 |
| **meandepth Morphologist FPO_left** | 0.96 | 18097 |
| **maxdepth Morphologist SPasup_right** | 0.96 | 17400 |
| **rfMRI part corr ICA100 edge 176 link 4-21** | 0.96 | 15864 |
| **rfMRI full corr ICA100 edge 190 link 4-35** | 0.96 | 15864 |
| **rfMRI part corr ICA100 edge 858 link 20-23** | 0.96 | 15864 |
| **ThickAvg Destrieux G_temp_sup-G_T_transv.rh** | 0.96 | 17125 |
| **hull_junction_length Morphologist SFinf_left** | 0.96 | 18101 |
| **rfMRI part corr ICA100 edge 853 link 19-53** | 0.96 | 15864 |
| **MO Superior_cerebellar_peduncle_R** | 0.96 | 16541 |
| **rfMRI full corr ICA100 edge 661 link 14-51** | 0.96 | 15864 |
| **rfMRI full corr ICA100 edge 1191 link 31-37** | 0.96 | 15864 |
| **rfMRI full corr ICA100 edge 133 link 3-29** | 0.96 | 15864 |
| **ICVF Posterior_corona_radiata_L** | 0.96 | 16540 |
| **GrayVol Destrieux G_occipital_sup.rh** | 0.96 | 17127 |
| **rfMRI part corr ICA100 edge 1328 link 37-51** | 0.96 | 15864 |
| **rfMRI full corr ICA100 edge 924 link 21-55** | 0.96 | 15864 |
| **rfMRI full corr ICA100 edge 574 link 12-47** | 0.96 | 15863 |

| | | |
|---|---|---|
| **surface Morphologist SC_left** | 0.96 | 18101 |
| **rfMRI full corr ICA100 edge 684 link 15-34** | 0.97 | 15864 |
| **GM_thickness Morphologist SFmedian_left** | 0.97 | 18069 |
| **MD Superior_longitudinal_fasciculus_R** | 0.97 | 16541 |
| **rfMRI full corr ICA100 edge 931 link 22-29** | 0.97 | 15864 |
| **rfMRI full corr ICA100 edge 516 link 11-32** | 0.97 | 15863 |
| **opening Morphologist SPasup_left** | 0.97 | 16537 |
| **rfMRI part corr ICA100 edge 392 link 8-43** | 0.97 | 15864 |
| **rfMRI part corr ICA25 edge 1 link 1-2** | 0.97 | 15864 |
| **rfMRI part corr ICA100 edge 1364 link 39-54** | 0.97 | 15864 |
| **surface Morphologist FCLa_right** | 0.97 | 18097 |
| **rfMRI part corr ICA100 edge 1007 link 24-42** | 0.97 | 15864 |
| **GM_thickness Morphologist SCLPC_right** | 0.97 | 5274 |
| **rfMRI part corr ICA100 edge 1219 link 32-42** | 0.97 | 15864 |
| **rfMRI full corr ICA100 edge 1006 link 24-41** | 0.97 | 15864 |
| **rfMRI full corr ICA100 edge 1466 link 49-51** | 0.97 | 15864 |
| **rfMRI part corr ICA100 edge 1254 link 33-55** | 0.97 | 15864 |
| **rfMRI full corr ICA100 edge 1425 link 44-50** | 0.97 | 15863 |
| **SurfArea Destrieux S_temporal_transverse.rh** | 0.97 | 17124 |
| **rfMRI part corr ICA100 edge 699 link 15-49** | 0.97 | 15864 |
| **rfMRI full corr ICA100 edge 1438 link 45-53** | 0.97 | 15864 |
| **GM_thickness Morphologist SOp_left** | 0.97 | 15266 |
| **rfMRI full corr ICA100 edge 818 link 18-54** | 0.97 | 15864 |
| **maxdepth Morphologist FCLrdiag_left** | 0.97 | 8551 |
| **rfMRI full corr ICA100 edge 1068 link 26-44** | 0.97 | 15864 |
| **SurfArea Destrieux S_calcarine.rh** | 0.97 | 17127 |
| **rfMRI part corr ICA100 edge 169 link 4-14** | 0.97 | 15864 |
| **rfMRI full corr ICA100 edge 1412 link 43-48** | 0.97 | 15864 |
| **rfMRI part corr ICA100 edge 1348 link 38-54** | 0.97 | 15864 |
| **rfMRI part corr ICA100 edge 995 link 24-30** | 0.97 | 15864 |
| **MD Splenium_of_corpus_callosum** | 0.97 | 16541 |
| **rfMRI full corr ICA100 edge 1062 link 26-38** | 0.97 | 15864 |
| **rfMRI full corr ICA100 edge 1188 link 31-34** | 0.97 | 15864 |
| **rfMRI full corr ICA25 edge 202 link 17-19** | 0.97 | 15864 |

| | | |
|---|---|---|
| **rfMRI part corr ICA100 edge 370 link 8-21** | 0.97 | 15864 |
| **opening Morphologist SLipost_left** | 0.97 | 17919 |
| **rfMRI full corr ICA100 edge 37 link 1-38** | 0.97 | 15864 |
| **rfMRI full corr ICA100 edge 724 link 16-35** | 0.97 | 15861 |
| **rfMRI full corr ICA100 edge 1420 link 44-45** | 0.97 | 15864 |
| **rfMRI part corr ICA100 edge 1445 link 46-51** | 0.97 | 15864 |
| **rfMRI full corr ICA100 edge 660 link 14-50** | 0.97 | 15862 |
| **rfMRI full corr ICA100 edge 625 link 14-15** | 0.97 | 15864 |
| **GrayVol Destrieux G_temp_sup-Lateral.lh** | 0.97 | 17127 |
| **maxdepth Morphologist FPO_left** | 0.97 | 18097 |
| **rfMRI full corr ICA25 edge 99 link 6-15** | 0.97 | 15864 |
| **rfMRI part corr ICA100 edge 706 link 16-17** | 0.97 | 15864 |
| **ThickAvg Destrieux S_precentral-sup-part.lh** | 0.97 | 17127 |
| **rfMRI full corr ICA100 edge 12 link 1-13** | 0.97 | 15864 |
| **rfMRI part corr ICA100 edge 86 link 2-34** | 0.97 | 15864 |
| **rfMRI part corr ICA25 edge 99 link 6-15** | 0.97 | 15864 |
| **rfMRI part corr ICA100 edge 89 link 2-37** | 0.98 | 15864 |
| **SurfArea Destrieux G_orbital.rh** | 0.98 | 17127 |
| **opening Morphologist STiant_right** | 0.98 | 18100 |
| **rfMRI full corr ICA25 edge 140 link 9-17** | 0.98 | 15864 |
| **hull_junction_length Morphologist SPeCinter_righ** | 0.98 | 18084 |
| **rfMRI part corr ICA25 edge 15 link 1-16** | 0.98 | 15864 |
| **rfMRI full corr ICA100 edge 863 link 20-28** | 0.98 | 15864 |
| **Left-vessel** | 0.98 | 17127 |
| **rfMRI full corr ICA100 edge 1196 link 31-42** | 0.98 | 15864 |
| **rfMRI full corr ICA100 edge 199 link 4-44** | 0.98 | 15864 |
| **ISOVF Uncinate_fasciculus_L** | 0.98 | 16527 |
| **rfMRI part corr ICA100 edge 1330 link 37-53** | 0.98 | 15864 |
| **rfMRI full corr ICA100 edge 325 link 7-23** | 0.98 | 15864 |
| **ThickAvg Destrieux G_insular_short.lh** | 0.98 | 17126 |
| **rfMRI full corr ICA100 edge 1431 link 45-46** | 0.98 | 15864 |
| **rfMRI part corr ICA100 edge 547 link 12-20** | 0.98 | 15864 |
| **rfMRI full corr ICA100 edge 890 link 20-55** | 0.98 | 15864 |
| **rfMRI part corr ICA25 edge 69 link 4-16** | 0.98 | 15864 |

| | | |
|---|---|---|
| **rfMRI part corr ICA100 edge 817 link 18-53** | 0.98 | 15864 |
| **rfMRI full corr ICA100 edge 562 link 12-35** | 0.98 | 15864 |
| **rfMRI full corr ICA100 edge 690 link 15-40** | 0.98 | 15864 |
| **rfMRI full corr ICA100 edge 183 link 4-28** | 0.98 | 15864 |
| **rfMRI full corr ICA100 edge 1325 link 37-48** | 0.98 | 15864 |
| **hull_junction_length Morphologist SPeCinf_right** | 0.98 | 16608 |
| **rfMRI part corr ICA100 edge 343 link 7-41** | 0.98 | 15864 |
| **rfMRI part corr ICA100 edge 882 link 20-47** | 0.98 | 15864 |
| **rfMRI part corr ICA100 edge 738 link 16-49** | 0.98 | 15864 |
| **ISOVF Posterior_corona_radiata_R** | 0.98 | 16527 |
| **rfMRI full corr ICA25 edge 27 link 2-9** | 0.98 | 15864 |
| **maxdepth Morphologist SOTlatmed_left** | 0.98 | 17430 |
| **rfMRI full corr ICA100 edge 1419 link 43-55** | 0.98 | 15864 |
| **rfMRI part corr ICA100 edge 64 link 2-12** | 0.98 | 15864 |
| **rfMRI part corr ICA100 edge 769 link 17-42** | 0.98 | 15864 |
| **MO Cingulum-cingulate_gyrus-L** | 0.98 | 16541 |
| **rfMRI part corr ICA100 edge 489 link 10-49** | 0.98 | 15864 |
| **rfMRI part corr ICA100 edge 309 link 6-55** | 0.98 | 15864 |
| **rfMRI full corr ICA100 edge 1337 link 38-43** | 0.98 | 15864 |
| **rfMRI full corr ICA25 edge 178 link 13-17** | 0.98 | 15864 |
| **rfMRI part corr ICA25 edge 186 link 14-18** | 0.98 | 15864 |
| **rfMRI part corr ICA100 edge 767 link 17-40** | 0.98 | 15864 |
| **surface Morphologist STpol_left** | 0.98 | 18076 |
| **ISOVF Anterior_limb_of_internal_capsule_L** | 0.99 | 16527 |
| **rfMRI full corr ICA100 edge 290 link 6-36** | 0.99 | 15864 |
| **ThickAvg Destrieux S_intrapariet_and_P_trans.rh** | 0.99 | 17127 |
| **rfMRI full corr ICA25 edge 23 link 2-5** | 0.99 | 15864 |
| **rfMRI part corr ICA100 edge 1048 link 25-53** | 0.99 | 15864 |
| **GrayVol Destrieux S_suborbital.rh** | 0.99 | 17125 |
| **rfMRI full corr ICA25 edge 174 link 12-21** | 0.99 | 15863 |
| **GM_thickness Morphologist SsP_right** | 0.99 | 18087 |
| **rfMRI part corr ICA100 edge 124 link 3-20** | 0.99 | 15864 |
| **rfMRI part corr ICA25 edge 70 link 4-17** | 0.99 | 15864 |
| **rfMRI full corr ICA100 edge 1262 link 34-42** | 0.99 | 15864 |

| | | |
|---|---|---|
| **GM_thickness Morphologist STsterascant_left** | 0.99 | 17549 |
| **hull_junction_length Morphologist SFpolairetr_left** | 0.99 | 18062 |
| **ISOVF Fornix-column_and_body_of_fornix** | 0.99 | 16527 |
| **rfMRI amplitude ICA25 component 6** | 0.99 | 15863 |
| **rfMRI part corr ICA100 edge 565 link 12-38** | 0.99 | 15864 |
| **rfMRI full corr ICA25 edge 47 link 3-11** | 0.99 | 15864 |
| **rfMRI full corr ICA100 edge 54 link 1-55** | 0.99 | 15864 |
| **rfMRI full corr ICA100 edge 1336 link 38-42** | 0.99 | 15864 |
| **maxdepth Morphologist FCLrscpost_right** | 0.99 | 16126 |
| **hull_junction_length Morphologist SFmedian_left** | 0.99 | 18078 |
| **SurfArea Destrieux S_oc_sup_and_transversal.rh** | 0.99 | 17127 |
| **rfMRI full corr ICA100 edge 1358 link 39-48** | 0.99 | 15864 |
| **opening Morphologist SForbitaire_left** | 0.99 | 17520 |
| **surface Morphologist FIP_left** | 0.99 | 18098 |
| **rfMRI full corr ICA100 edge 792 link 18-28** | 0.99 | 15862 |
| **rfMRI part corr ICA100 edge 492 link 10-52** | 0.99 | 15864 |
| **rfMRI full corr ICA100 edge 419 link 9-24** | 0.99 | 15864 |
| **MD Corticospinal_tract_L** | 0.99 | 16541 |
| **rfMRI part corr ICA100 edge 1149 link 29-44** | 0.99 | 15864 |
| **FA External_capsule_L** | 0.99 | 16541 |
| **rfMRI full corr ICA25 edge 3 link 1-4** | 0.99 | 15864 |
| **rfMRI full corr ICA25 edge 175 link 13-14** | 0.99 | 15864 |
| **opening Morphologist FCLrdiag_left** | 0.99 | 8551 |
| **hull_junction_length Morphologist FCLp_left** | 0.99 | 18101 |
| **meandepth Morphologist SLipost_left** | 1.0 | 17917 |
| **rfMRI part corr ICA25 edge 97 link 6-13** | 1.0 | 15864 |
| **rfMRI full corr ICA100 edge 34 link 1-35** | 1.0 | 15864 |
| **rfMRI full corr ICA100 edge 1297 link 36-38** | 1.0 | 15864 |
| **rfMRI full corr ICA100 edge 423 link 9-28** | 1.0 | 15864 |
| **rfMRI full corr ICA100 edge 729 link 16-40** | 1.0 | 15864 |
| **rfMRI part corr ICA100 edge 1020 link 24-55** | 1.0 | 15864 |
| **rfMRI part corr ICA25 edge 123 link 8-12** | 1.0 | 15864 |
| **ThickAvg Destrieux G_temporal_inf.rh** | 1.0 | 17127 |
| **rfMRI part corr ICA100 edge 724 link 16-35** | 1.0 | 15864 |

| | | |
|---|---|---|
| **rfMRI full corr ICA100 edge 506 link 11-22** | 1.0 | 15864 |
| **rfMRI part corr ICA25 edge 106 link 7-8** | 1.0 | 15864 |
| **rfMRI full corr ICA100 edge 1289 link 35-49** | 1.0 | 15864 |
| **rfMRI part corr ICA100 edge 453 link 10-13** | 1.0 | 15864 |
| **rfMRI part corr ICA100 edge 671 link 15-21** | 1.0 | 15864 |
| **rfMRI full corr ICA100 edge 911 link 21-42** | 1.0 | 15864 |
| **rfMRI part corr ICA25 edge 74 link 4-21** | 1.0 | 15864 |
| **rfMRI full corr ICA100 edge 1416 link 43-52** | 1.0 | 15864 |
| **GM_thickness Morphologist FCLrscpost_left** | 1.0 | 13956 |
| **rfMRI full corr ICA100 edge 1118 link 28-39** | 1.0 | 15863 |
| **rfMRI part corr ICA100 edge 890 link 20-55** | 1.0 | 15864 |
| **rfMRI part corr ICA100 edge 918 link 21-49** | 1.0 | 15864 |
| **GrayVol Destrieux G_occipital_sup.lh** | 1.0 | 17127 |
